# Mapping neurogenetic characteristics of psychopathological procrastination using normative modeling in a prospective twin cohort

**DOI:** 10.1101/2025.06.06.658269

**Authors:** Yuanyuan Hu, Yancheng Tang, Wei Li, Dang Zheng, Yuening Jin, Qingchen Fan, Jan K Buitelaar, Yuan Zhou, Zhiyi Chen

**Affiliations:** State Key Laboratory of Cognitive Science and Mental Health, Institute of Psychology, Chinese Academy of Sciences, Beijing 100101, China; Department of Psychology, University of Chinese Academy of Sciences, Beijing, 100049, China; Experimental Research Center of Medical and Psychological Science, School of Psychology, Third Military Medical University, Chongqing, 400038, China; Department of Early Childhood Education, China National Children’s Center, Beijing, 100035, China; Department of Cognitive Neuroscience, Radboud University Medical Center, Nijmegen, Netherlands; Donders Institute for Brain, Cognition and Behaviour, Nijmegen, Netherlands; The National Clinical Research Center for Mental Disorders & Beijing Key Laboratory of Mental Disorders, Beijing Anding Hospital, Capital Medical University Beijing, China; Faculty of Psychology, Southwest University, Chongqing, 400038, China; Key Laboratory of Cognition and Personality, Ministry of Education, 400038 China

**Author notes:** Corresponding at Zhiyi Chen and Yuan Zhou. These authors contributed equally to this work (Yuanyuan Hu, Yancheng Tang).

## Abstract

Procrastination, affecting over 70% global population, is pervasively incurring negative outcomes in human society. This has long been studied as a bad daily habit, but it loses in delineating neurogenetic substrates underlying its psychopathological phenotyping. Using a prospective twin adolescent cohort, we demonstrate moderate heritability of this subclinical condition - PPS. Neuroimaging normative modeling analysis, further reveals that neurodevelopmental deviations in nucleus accumbens during adolescence, are predictive of PPS in adulthood, while such deviations-PPS mappings were highly genetically shared. Beyond to regional anomalies, PPS-specific whole-brain deviation patterns, notably in the default mode network, are neurobiologically enriched with changes in cortical manifolds (gradients) and neurotransmitter systems. Integrating these neuroimaging markers with transcriptomic atlas, we capture significant PPS-specific neurogenetic signatures associated with molecular transport system, neuroimmune responses, and neuroinflammation, particularly in serotonergic and dopaminergic pathways. These findings shed light on the multisystem neurogenetic architecture underlying PPS, providing evidence to theoretically conceptualize this psychopathological phenotype as a subclinical “brain disorder”.

**Teaser:** Psychopathological procrastination is not a bad daily habit solely, but a “brain disorder” associated with multiscale neurodevelopment.

## Introduction

Procrastination — the pervasive tendency to delay tasks despite adverse consequences — has surged in prevalence over recent decades, now affecting 15-20% of adults globally, a sharp rise from 4-5% in the 1970s.(*1*) Among students, over 75% report academic or personal procrastination,(*2–4*) with cross-cultural studies confirming its 5-year stability across Asian, European, and American populations.(*5–8*) Beyond its ubiquity, procrastination correlates with profound societal and individual costs, including diminished academic/occupational performance, strained relationships, and reduced socioeconomic status and well-being.(*9–12*) Moreover, chronic procrastination is strongly associated with lifelong physical and mental health risks, such as cardiovascular diseases, hypertension and internalizing psychiatric symptoms.(*13–17*) Despite alarming statistics and consequences, procrastination has historically been dismissed as a maladaptive behavioral habit rather than as a psychopathological entity, leading to an underestimation of its (sub)clinical significance.

Recently psychopathological procrastination (PPS) has been proposed as a psychopathological condition characterized by persistent, irrational and pervasive task postponement leading to significantly uncontrollable distress and panic for ≥ 6 months.(*18*) This framework re-conceptualizes procrastination as a potential subclinical manifestation involving symptomatology of mood and personality disorders.(*18, 19*) Cumulative evidence have identified “persistent and pervasive postponements” as a transdiagnostic marker linking to neurodevelopmental disorders,(*20*) internalizing disorders,(*16, 17, 21*) externalizing disorders(*22, 23*) and panic symptomatology.(*24*) As reinforced by theoretical models, reframing procrastination through the lens of subclinical psychopathology opens a new pathway to understand its etiology.(*25–27*) Despite these compelling arguments, psychopathological substrates of PPS remain largely uncharacterized — a critical gap limiting mechanistic and translational interpretability.

The Research Domain Criteria (RDoC) framework provides a hierarchical and multiscale approach to decipher psychopathology, integrating symptomatologic, genetic, and neurobiological data across cellular, molecular, circuit-level system.(*28–30*) Unlike symptom-based classifications, RDoC conceptualizes mental illnesses as “brain disorders”, emphasizing the neurogenetic foundations to elucidate their etiologies.(*31, 32*) This research framework has demonstrated superior psychopathological interpretability in mood disorders (e.g., depression),(*33*) suicide(*34*) and neurodevelopmental conditions.(*35*) Although no direct evidence currently links PPS to a known brain disorder, leveraging the RDoC framework, procrastination traits are found to be intimately correlated with dysfunctions in brain networks involved in self-control, emotional regulation and episodic thinking,(*36, 37*) and particularly with neuroanatomical abnormalities in prefrontal-limbic circuits (e.g., prefrontal cortex, anterior cingulate cortex, insula, orbital frontal cortex).(*36, 38–40*) Twin studies further corroborate a genetic basis for procrastination, with moderate heritability (*h*^2^ = 0.28 - 0.46) observed across independent cohorts.(*41–43*) Thus, RDoC’s integrative framework—bridging symptomatology, genetics and multiscale brain ontologies—paves an unique pathway to further unravel the neural and genetic mechanisms underlying PPS.

Despite strengths of the RDoC framework in parsing the multiscale psychopathology of PPS, a critical challenge remains: the notable individual heterogeneity in brain phenotyping.(*44, 45*) To address this issue, normative modeling has emerged as a powerful method, mapping individual deviations from population-based healthy brain norm rather than relying on group-averaged statistics.(*46–48*) This approach overcomes the limitations of traditional case-control analyses—which prioritize group differences— by capturing individualized brain features that mitigate statistical biases caused by inter-individual variability in neuroimaging phenotyping.(*49, 50*) Accumulating evidence demonstrates that normative deviations identify robust neurobiological biomarkers and clinical biotypes, particularly in heterogeneous psychopathological populations.(*51–56*) By integrating normative modeling with RDoC’s conceptualization of mental illnesses as “brain disorders”, it enables a more precise delineation for the PPS’s neurogenetic architecture.

In this study, leveraging the integrative RDoC framework and normative modeling, we investigated the neurogenetic substrates underlying PPS spanning multiscale large-scale neurobiological datasets. We began by analyzing a 8-year longitudinal twin cohort (*N* = 140; 37 monozygotic-twin [MZ] pairs and 33 dizygotic-twins [DZ] pairs), building an ACE (Additive genetic, Common environmental, and Unique environmental) structural equation model to estimate the heritability of PPS, as phenotyped via the Pathological Procrastination Diagnostic Questionnaire (PDQ; **Figure 1a, Supplemental Methods 1, Table S1**). Upon confirming PPS as a heritable phenotype, we used the ENIGMA lifespan working group dataset (*N* = 37,407; 19,826 females; age range: 3-90 years) for normative modeling to evaluate individual neurodevelopmental deviations in cortical thickness (CT), surface areas (SA) and subcortical volume (SV).(*57*) This enabled individual-level analysis of univariate neurogenetic associations between regional neuroanatomical deviations and PPS **(Figure 1b)**. Moving beyond univariate associations, we employed multivariate intersubject representation similarity analysis (IS-RSA), identifying whole-brain deviation pattern of specifically representing PPS **(Figure 1c)**. Aligning these patterns with 40+ multiscale normative neurobiological maps (e.g., metabolism, neurotransmitters, electrophysiology) via the Neuromaps toolbox,(*58*) it allows us to characterize PPS’s brain ontologies **(Figure 1d)**. Finally, integrating normative transcriptomic maps from Allen Human Brain Atlas (AHBA; *N* = 6 donors) and BrainSpan Developmental Atlas (BSDA; *N* = 42 donors; prenatal weeks to 40 years), we systematically examined upstream PPS-specific gene expression patterns and delineated their downstream genetically neurobiological correlates, particularly in molecular and cellular enrichment **(Figure 1d)**.

**Figure 1.**
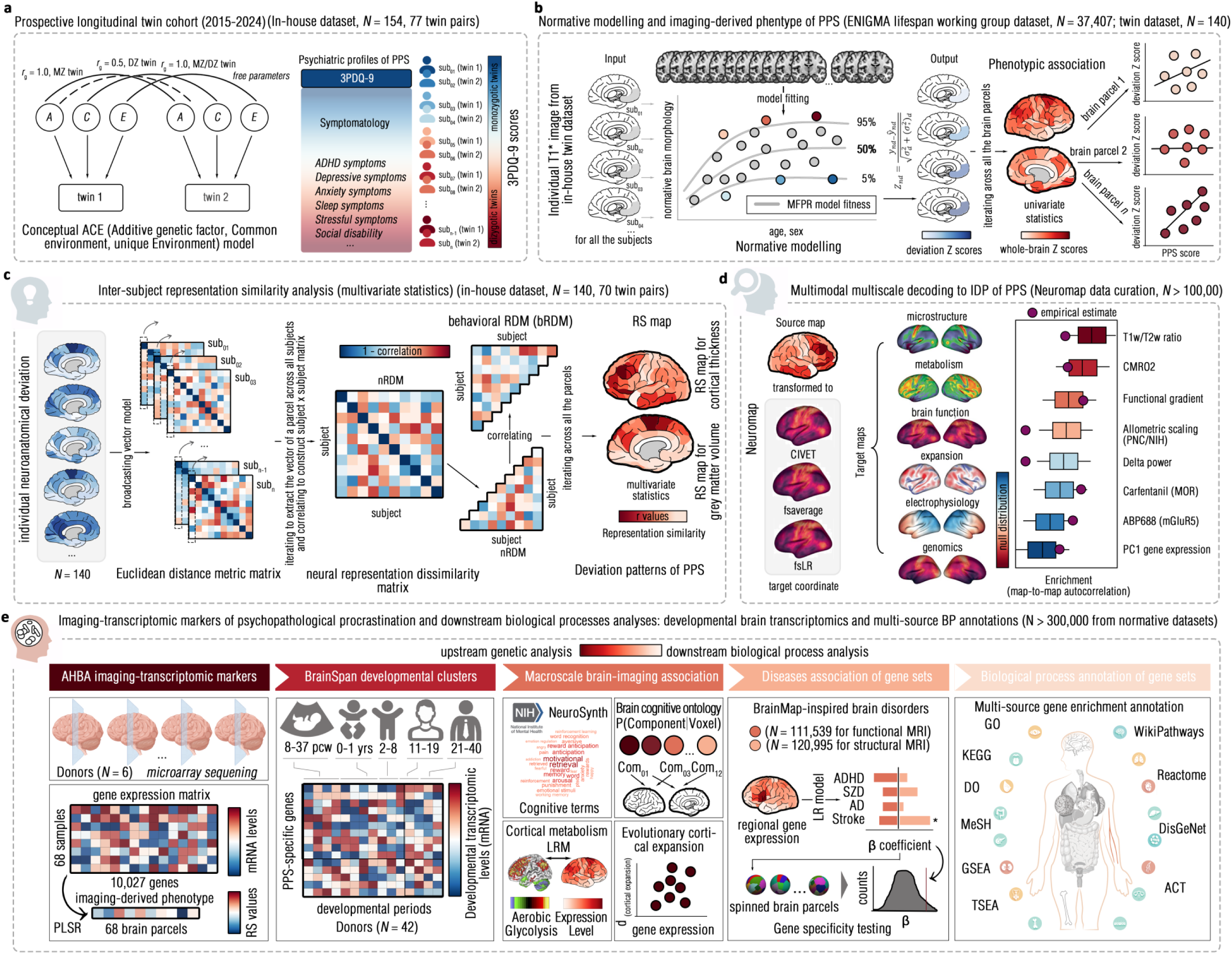
Analytic flowchart of the present study. **a**, described conceptual computations to twin ACE (Addictive genetic factor, Common environment, unique Environment) model and featured core symptoms that the 3PDQ-9 (Pathological Procrastination Diagnostic Questionnaire - 9 items) measured. Colorbar indicated 3PDQ-9 scores. **b**, illustrated main steps of estimating deviation Z scores from normative modeling method. The pre-trained normative model established from ENIGMA lifespan working group (N = 37,047) was used to calculate brain morphometric deviation of these twins in each brain region that parceled by Desikan-Killiany and Aseg Atlas. The phenotypic association between such deviations and pathological procrastination tendency that measured by the 3PDQ-9 scores was calculated for each brain region by generalized linear mixed-effect regression model (GLMM). **c**, pictured main steps of conducting inter-subject representation similarity analysis (RSA). This method established regional neural representation dissimilarity matrices (nRDMs) and inter-subject behavioral RDM by using Euclidean distance, and further estimated Spearman’s correlation between the bRDM and nRDM across all the brain parcels, with Benjamini-Hochberg correction for multiple comparisons. **d**, depicted brain functional decoding from RS patterns. Once the RS values for all the brain regions are calculated, they are deployed as “source map” to correlate to spatial patterns of normative multimodal multiscale brain atlas, for delving into what normative brain features were enriched by the RS patterns. **e**, detailed downstream analyses from imaging-transcriptomic association. By using the Allen Human Brain Atlas (AHBA) normative gene expression data, the partial least square regression (PLSR) was used to find imaging-transcriptomic markers of RS values. Furthermore, the BrainSpan dataset was used to probe into the brain developmental clusters of these imaging-transcriptomic markers. Finally, several normative neurobiological datasets were used to annotate macroscale, microscale, molecular, cellular biological processes and other phenotype (e.g., diseases).

## Results

### Schematic Summary of Main Findings

Using the RDoC framework, this 8-year longitudinal twin cohort study first revealed a moderate heritability for PPS (*h*^2^ = 0.47), confirming a genetic contribution to this subclinical phenotype. Building on this genetic foundation, we combined normative modeling of a population-based neuroimaging dataset and bivariate Cholesky decomposition methods to elucidate PPS’s neurogenetic underpinnings. Results revealed that morphological deviations in nucleus accumbens (NAcc) during adolescence were predictive of the PPS phenotyped in adulthood, with significant brain-symptomatology associations (*r_ph_* = 0.18, *95%CI* = [0.02, 0.35]) and shared genetic influences (*r_g_* = 0.89, *95%CI* = [0.89, 1]). Beyond univariate regional associations, multivariate representation similarity model identified whole-brain deviation patterns specifically represented PPS, localized predominantly within the default mode network. These representation patterns exhibited spatial enrichment in cortical manifolds and catecholaminergic neurotransmitter systems (e.g., dopamine D1 receptors, serotonin transporters). Integrating these genetic, brain, and multiscale neurobiological mappings via partial least square (PLS) modeling, we identified PPS-specific transcriptomic markers, notably *FRMD8*, *MPLKIP* and *DEPP1*, biologically implicated in molecular membrane trafficking (e.g., vesicle-mediated transport), neuroinflammatory responses, and neuroimmune pathways (e.g., microRNA families and TNFα). In summary, these findings contributed empirical support for conceptualizing PPS as a neurogenetically driven “brain disorder”, arising from dysregulated neurotransmitter (dopamine/serotonin) and neuroimmune systems involved in reward processing.

### Moderate Heritability of PPS: Twin ACE Modeling

To investigate genetic contributions underlying PPS, we first carried out parametric phenotypic association analyses. After adjusting for demographic covariates (e.g., age, sex; **Supplemental Results 1**), we found significant within-pair correlations for PPS in MZ twins (*r* = 0.52, *p* < .001), but not in DZ twins (*r* = −0.27, *p* = 0.12). Intra-class correlation (ICC) analysis further reinforced this observation, showing that the within-pair correlations twins were significantly higher in MZ twins than those in DZ ones (*F* = 6.48, *p* < .0001) (**Supplemental Results 1**), indicating a potential genetic influence on PPS. Thus, leveraging nested AE structural equation modeling (**see *Methods* and Supplemental Methods 2, Table S2**), the univariate heritability (*h*²) of PPS was estimated, revealing a statistically moderate effect in this cohort (*h²* = 0.47, 95% Confidence Interval (CI): 0.02 - 0.50) (**Figure 2a and Supplemental Results 1, Table S3**). These results jointly support this notion that PPS represents a moderately heritable subclinical phenotype.

**Figure 2.**
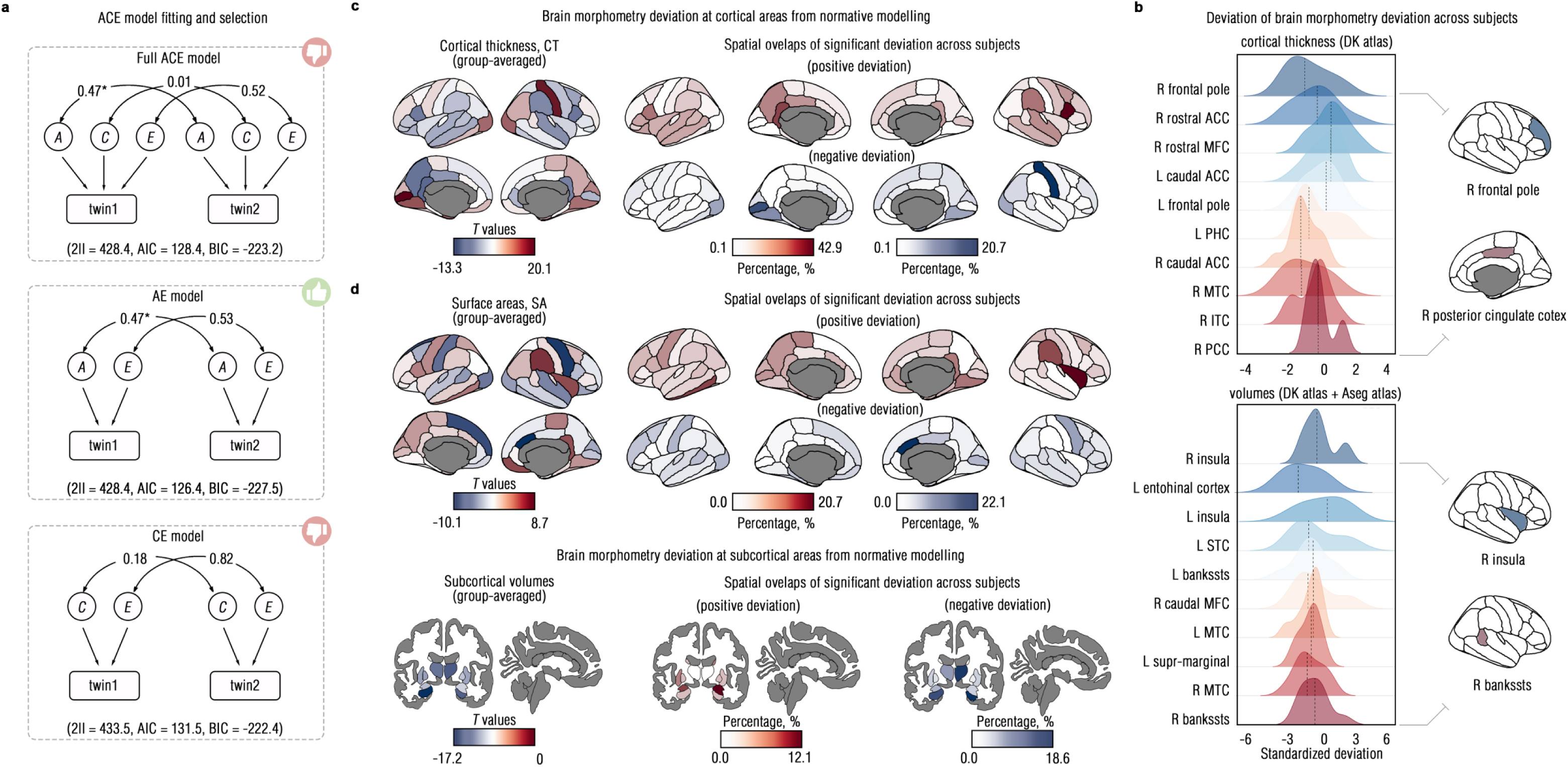
ACE models and brain morphological deviations from normative modeling methods. **a**, described goodness-of-fitting of the full ACE model and its stratified nested alternatives (i.e., −2ll, Log-Likelihood * −2; AIC, Akaike Information Criterion; BIC, Bayesian Information Criterion). The icon of “applaud” indicated the best-fitting twin model. **b**, exemplified distributions of deviation Z scores for brain regions that measured by cortical thickness (Top) and surface areas (Bottom). **c**, presented cortical thickness deviations of each brain region, with the left panel for showing group-averaged deviations. By calculating the normalized deviation Z score of each brain region across all the participants, the proportions of showing statistically significant deviations across all the participants (Z ≥ 1.96 or Z ≤ 1.96) are presented in the right panel. **d**, illustrated similar layouts mentioned above, with the top panel for showing surface areas deviations and with the bottom panel for showing subcortical volume deviations.

### Brain Morphological Deviations From Normative Modeling

Upon confirming the PPS’s genetic basis, we applied normative modeling to estimate individual-level morphological deviations and examine brain-PPS neurogenetic associations. By referencing twin-specific neuroanatomical measures against a population-based neuroimaging dataset (*N* = 37,407), we quantified regional cortical thickness (CT), surface areas (SA) and subcortical volume (SV) deviations as standardized *Z-*scores using the Desikan-Killiany (DK) atlas (68 cortical regions) and the Aseg atlases (14 subcortical regions). Kernel density estimation (KDE; ***see Methods***) revealed approximately Gaussian distributions across almost all the brain parcels **(Figure 2b, Supplemental Results 2, Figure S3-5 Table S4-5)**, validating their statistical suitability for phenotypic association analyses.

For CT, group-level deviations were found statistically significant in the precuneus, lingual, anterior cingulate cortex (ACC) and inferior/orbital frontal cortex (*q* < .05) **(Figure 2b, Supplemental Results 2, Table S6)**. For SA and SV, group-level deviations dominantly localized to the posterior cingulate cortex (PCC), ACC, insula, orbital frontal cortex (OFC), hippocampus, and NAcc (*q* < .05) **(Figure 2d, Supplemental Table S7-8)**. At the individual level, 11.4 - 42.8% of participants exhibited significant regional deviations on cortical thickness (*Z* ≥ 1.96 or *Z* ≤ −1.96) in the lingual cortex, pericalcarine cortex, and middle temporal cortex (MTC) **(Figure 2c, Supplemental Table S6)**. Over 10% of twins showed significant individual deviations (Z ≥ 1.96 or Z ≤ −1.96) in the NAcc, ACC and insula, as quantified by either SA or SV **(Fig. 2d, Supplemental Tab. S7-8)**. Taken together, these findings confirmed individual brain deviations in this cohort indeed, particularly in precuneus, temporal cortex and NAcc, suggesting their potentially neurodevelopmental differences.

### Shared Regional Neurogenetic Markers of PPS

After confirming statistical suitability and neurodevelopmental deviations in this cohort, we employed generalized linear mixed-effect models (GLMM) to test predictive of such deviations to PPS, region-by-region, adjusting for zygosity, sex and age. In twins, CT deviations in the precuneus during adolescence significantly predicted PPS phenotyped in their adulthood (*β* = 2.9, *SE* = 0.7, *R*^2^ = 0.1, *t* = 4.1, *q* = .006, adjusted by Benjamini-Hochberg correction; **Figure 3a, Supplemental Results 3, Table S9-10)**. For modeling SA deviations into this prediction, inferior frontal gyrus and Heschl’s gyrus showed possibly predictive of PPS (*p* _uncorrected_ = .02-.007). Despite reaching marginal statistical significance, SV deviations in the NAcc still exhibited suggestive associations with PPS (left: *β* = 1.2, *p* _uncorrected_ = .03; right: *β* = 1.5, *p* _uncorrected_ = .01)**(Figure 3a, Supplemental Results 3, Table S11)**. More importantly, the AE model posing best fit identified significantly shared genetic variants between right NAcc SV and PPS (bivariate *h*^2^ = 0.60, 95% CI = [0.56 - 0.63]) only, with significant phenotypic (*r*_ph_ = 0.18, *95% CI* = [0.02-0.34]) and genetic correlations (*r*_g_ = 0.89, *95% CI* = [0.89, 1]) **(Figure 3b)**. Full results for bivariate AE models can be found at **Supplemental Results 4, Table S12-14.**

**Figure 3.**
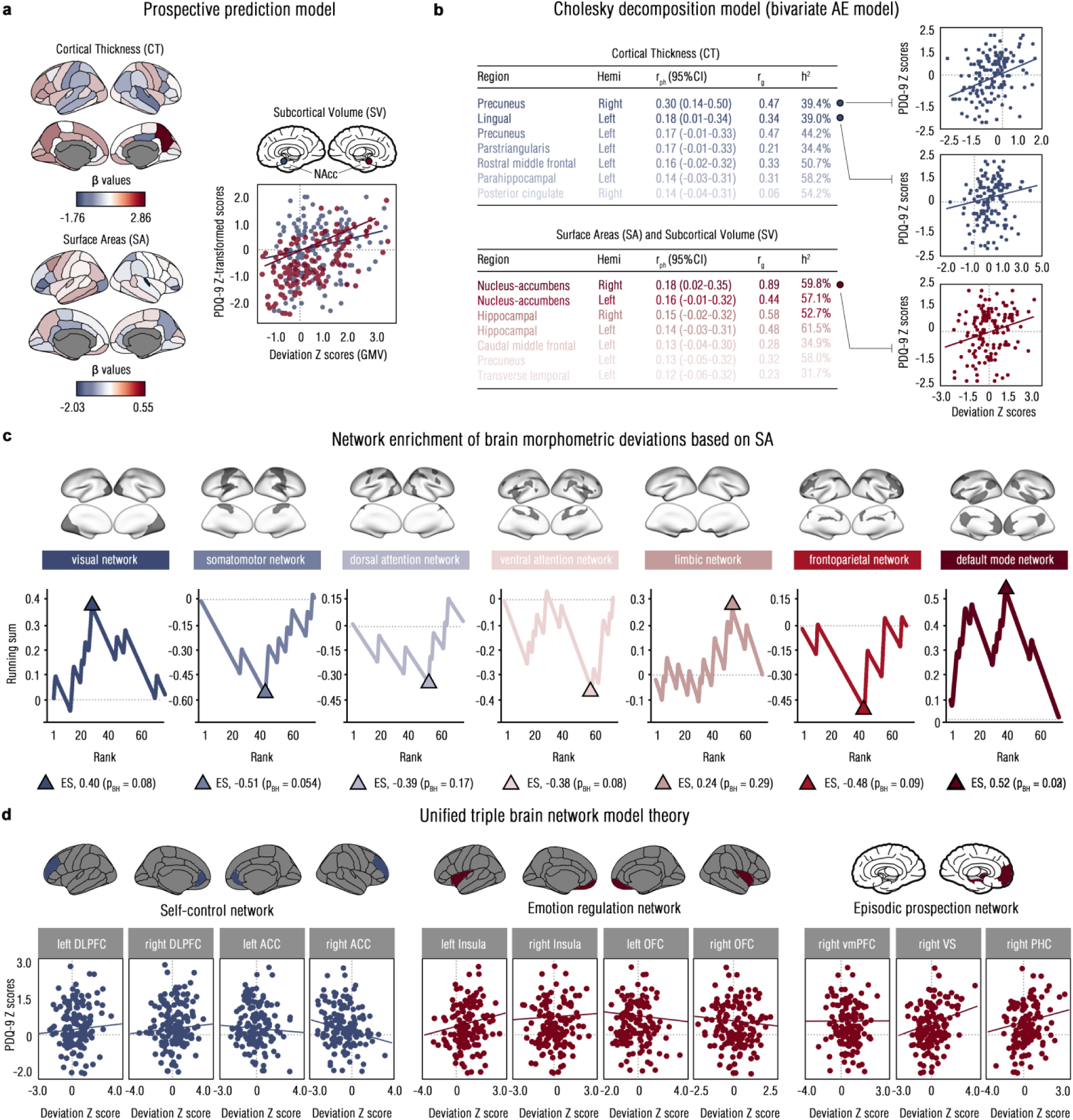
Prospective prediction of brain morphological deviations to psychopathological procrastination (PPS). **a**, presented prospectively predictive effects (*β* coefficient) of brain morphological deviations (*Z* scores) to pathological procrastination scores using mixed-effect generalized linear model, with top layout showing cortical thickness deviation and with the bottom layout showing surface areas. In the right panel, the scatter plots illustrated the linear association between subcortical volumes in bilateral nucleus accumbens (NAcc) and PPS measured by PDQ-9, with the left NAcc colored by blue and with the right NAcc colored by red. **b**, reported results of bivariate AE model (r_ph_ = phenotypic correlation; r_g_ = genetic correlation; h^2^ = heritability), and illustrated scatter plots showing statistically significant deviation-PPS phenotypic associations. **c**, showed network-wise enrichment from predictive effects (*β*) into brain systems defined by Yeo-7 atlas, and the triangles indicated enrichment score (ES). P values were estimated by the self-constrain spinning permutation test and were adjusted by Benjamini-Hochberg (BH) correction. **d**, detailed linear association between the deviations and specific brain regions that were defined by triple brain network model of procrastination trait (Chen & Feng, 2022), incorporated by 11 brain regions.

To assess spatial enrichment of deviation-PPS predictive mappings over regional associations, we conducted network enrichment significance testing (NEST, Weinstein et al., 2024) based on the Yeo-7 atlas (***see Methods***). Despite high variability across regions (*β* = −2.1 to 2.8), these predictive associations were significantly enriched within the default mode network (DMN, *q* < .05) phenotyped by CT **(Figure 3c, Supplemental Results 5, Table S15-16)**. To test the specificity of these findings, we further examined whether deviation-PPS associations mapped onto the theoretical triple brain network of procrastination trait (i.e., self-control, emotional regulation and episodic thinking networks; Chen & Feng, 2022). However, no theory-specific networks significantly enriched, irrespective of phenotyping from CT, SA or SV **(Figure 3d, Supplemental Results 5, Table S17)**. Collectively, these finding may establish morphological deviations, particularly in NAcc and DMN, as uniquely neurogenetic scaffolding of PPS, supporting its conceptualization as a subclinical “brain disorder”.

### Multivariate Morphological Representation Patterns and Multiscale Neurobiological Ontologies of PPS

Beyond univariate brain-symptomatology mappings, multivariate intersubject representation similarity (RS) analysis characterized whole-brain and region-specific morphological deviation patterns underlying PPS, respectively **(*see Methods*)**. Estimating the spatial alignment between the neural representation dissimilarity matrix (neural RDM) of whole-brain morphological deviations and behavioral RDM of the PPS **(Figure 4a)**, whole-brain deviation patterns were found significantly correlated with PPS, including CT-derived (*r* = 0.03, *95% CI* = [0.01 - 0.05], *q* = .006), SA-derived (*r* _SA_ = 0.02; 95% *CI* = [0.01 - 0.04], *q* = .04), and SV-derived (*r* _sv_ = 0.03; 95% CI = [0.01 - 0.05], *q* = .006) representations **(Figure 4b)**. Using spatial Euclidean distances **(*see Methods*)**, region-specific RS analysis mapped PPS to deviation patterns in the superior temporal sulcus (STS), ACC, NAcc, hippocampus, and prefrontal regions (*q* < .05) **(Figure 4c and Supplemental Results 6, Table S18-20)**. Nevertheless, these representation patterns were not yet enriched in known intrinsic functional networks **(Figure 4d and Supplemental Results 6, Table S21-22)**. Despite unfeasibility of estimating shared heritability in the RS design, these results allowed us identifying whole-brain representation patterns of PPS, thus enabling multiscale neurobiological mappings.

**Figure 4.**
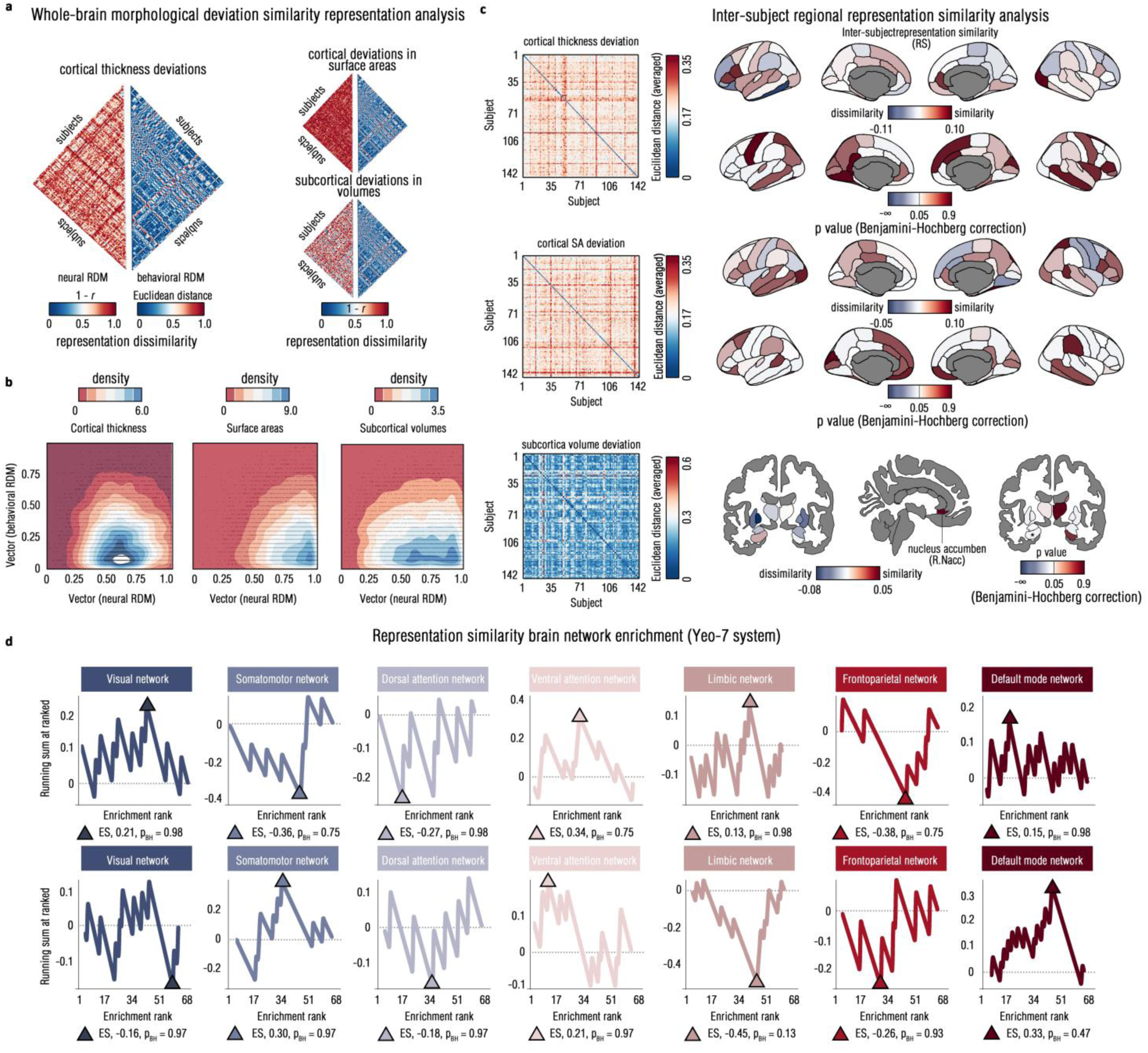
Representation similarity of brain morphological deviations to psychopathological procrastination. **a**, drew upper triangular matrix for whole-brain representation dissimilarity (RDM) that was calculated by the 1 - (inter-regional correlations) for cortical thickness (CT) deviation, surface areas (SA) deviation and subcortical volume (SV) deviation, respectively. **b**, pictured the density scatterplot to show the linear matrix-wise association between neural RDM and behavioral RDM constructed by inter-subject Euclidean distance to PPS scores. **c**, visualized inter-subject representation dissimilarity matrices for CT, SA and SV, respectively. In the right panel, it depicted inter-subject representation similarity of brain morphological deviations to psychopathological procrastination across all the regions, colored by both representation similarity (RS, *r*) values (top layout) and corresponding *q* values (bottom layout). **d**, illustrated network-wise enrichment of RS values to predefined brain systems (Yeo-7 networks), as phenotyped by CT (top) and SA (bottom).

By estimating neurobiological mappings across over 40 multiscale normative ontologies (e.g., neuronal receptors, electrophysiology, microstructure; ***see Methods***), spatial enrichment analysis clarified their alignment with cortical manifold (fourth gradient, G4, *r* = 0.20, *p*_spin_ = 0.003; **Figure 5a and Supplemental Results 7, Table S23**) and catecholaminergic neurotransmissions, including the serotonin transporter (5-HTT; *r* = - 0.34, *p*_spin_ < 0.001), dopamine transporter (DAT; *r* = −0.24, *p*_spin_ = 0.01) and dopamine type-1 receptor (D1; *r* = −0.19, *p*_spin_ = 0.03; spin permutation test; **Figure 5b-c and Supplemental Results 7, Table S24**). Despite no significant spatial correlations with other macroscale/microscale maps, these findings bridged PPS-specific morphological deviation representations to dysregulated neurofunctional gradients and molecular pathways, reinforcing this conceptualization of positioning PPS as a “brain disorder” involving in multisystematic psychopathology.

**Figure 5.**
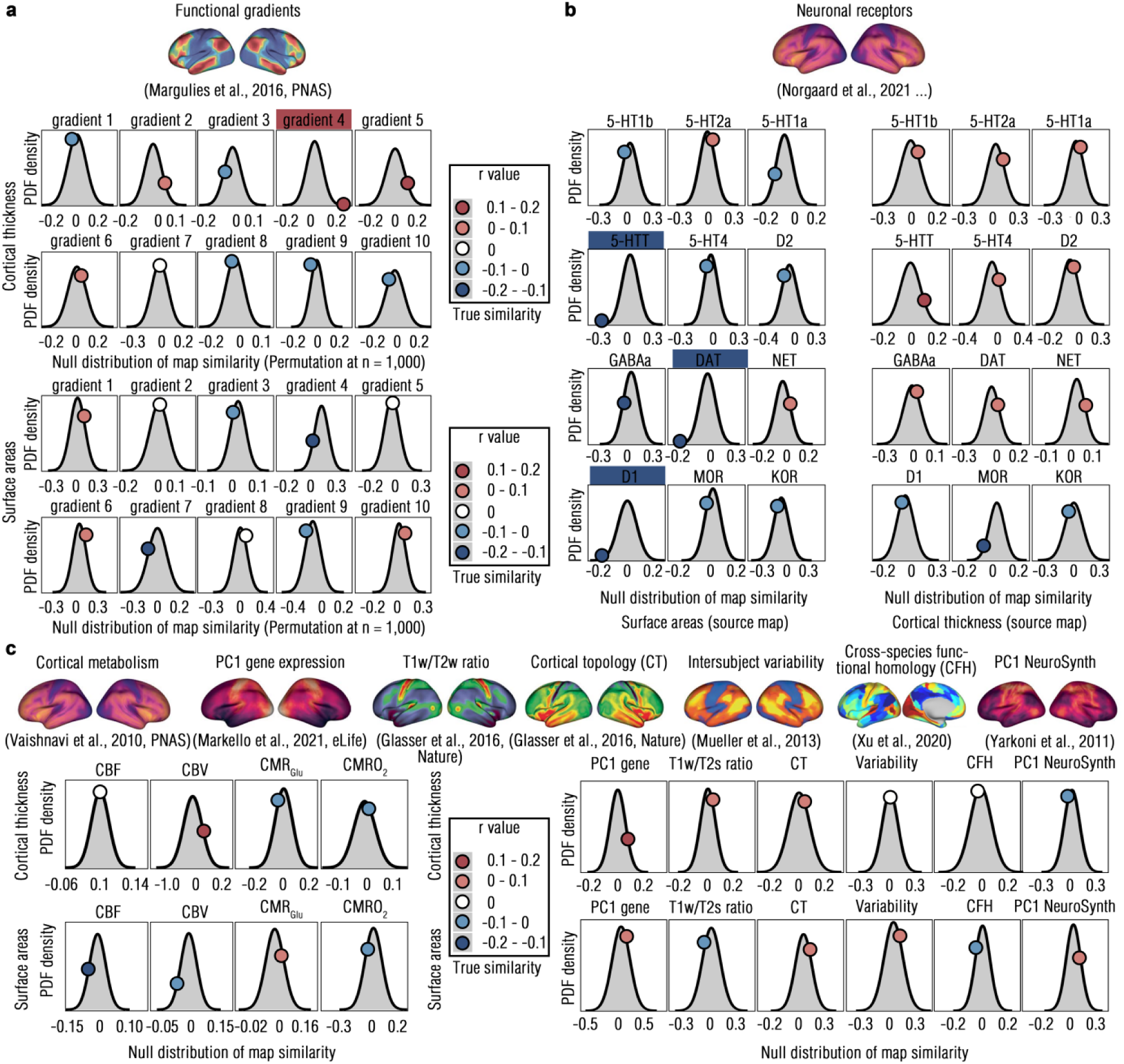
Spatial similarity of representation similarity (RS) pattern to multiscale normative maps. **a**, provided null distributions of spatial similarity between source map (RS patterns) and target maps (functional gradients) from spin permutation (n = 1,000) using Probability Density Function (PDF) plots. The top of this panel exemplified a target map that referred from the technical paper (Markello et al., 2022). The colored shallow box in names of these brain ontologies indicated this spatial similarity reached statistical significance (*p* < .05), with dark red for indicating the positive enrichment and with the dark blue for indicating the dark red. **b**, illustrated such spatial similarities from neuronal receptors by the same plotting characteristics. **c**, detailed spatial similarity of RS patterns to cortical metabolism, gene expression, microstructure (T1w/T2w), cortical topology, inter-subject variability, cross-species functional homology and cognitive ontology, respectively, with the same plotting characteristics.

### Transcriptomic Markers and Their Multiscale Neurobiological Enrichment to PPS

While genetic, neural, and molecular markers associated with PPS have been identified separately, a holistic understanding of its multiscale neurogenetic underpinnings remains not yet looped. We thus integrated whole-brain gene expression data from the Allan Human Brain Atlas (AHBA) to capture deviation-transcriptomic markers for PPS. Using partial least squares (PLS) regression, we mapped gene expression profiles to PPS-specific deviation patterns characterized by SA, explaining 22.5% of the variances in the second component (PLS2, *p* < .01) **(Figure 6a)**. PLS2+ (78 overexpressed genes, *q* < 0.005) and PLS2-(226 underexpressed genes, *q* < 0.005) were identified enriching in loci *FRMD8*, *MPLKIP, DEPP1* (PLS2+) and *CSTB*, *SCRN1, C9orf16* (PLS2-), respectively **(Figure 6b-c and Supplemental Results 8, Table S25-26)**. In the NeuroSynth decoding, these gene expressions were mapped into cognitive ontologies involving in psychopathological processes, particularly in memory loading, control process and behavioral responses **(Figure 6d)**. No statistically significant predictions were found when deviations were phenotyped by either CT or SV. Thus, we identified molecular neurogenetic markers potentially implicated in PPS, looping RDoC research frameworks by bridging molecular gene expression patterns and its brain deviations Considering prospective prediction from brain deviations to PPS, we further curated longitudinal transcriptomic data from the BrainSpan Developmental Atlas (BSDA) for identifying neurodevelopmental trajectories of these PPS-specific gene expression patterns. Using overlapping genes between this dataset and PLS2 set (273 genes) across six brain regions identified in RSA analysis **(Figure 7a*, see Methods*)**, we mapped their temporal expression profiles from the fetal period (8 post-conception weeks) to adulthood (40 years old) for aligning PPS-specific deviation patterns. This analysis demonstrated statistically significant longitudinal trajectories across 30 neurodevelopmental periods in each region (*p* < .001, Permutation test; **Figure 7b and Supplemental Methods 9, Table S27**). By building upon dominance model **(*see Methods*)**, *ZNF599*, *PAFH1B3*, and *ARMCX6* were found dominantly expressing in the dorsolateral prefrontal cortex (DLPFC) **(Figure 7c and Supplemental Methods 9, Table S28)**, with similar dominance patterns observed in temporal cortex (*LPIN1, S100A1, TARSL2*) and caudal anterior cingulate cortex (*LRRC61, BASP1, ZNF599*; **Figure 7c and Supplemental Results 9, Table S28-33**). To interpret molecular pathways of these gene expression trajectories, we performed fuzzy C-means clustering on the neurodevelopmental trajectories, followed by Gene Ontology (GO) enrichment analysis (***see Methods***). These analyses identified their enrichment into molecular transports, particularly in “pleural mesothelioma”, “transport of small molecules” and “vesicle-mediated transport” **(Figure 7d, Supplemental Results 10, Table S34-35)**, thereby enriching close-loop neurobiological understanding of PPS’s neurodevelopmental psychopathology.

**Figure 6.**
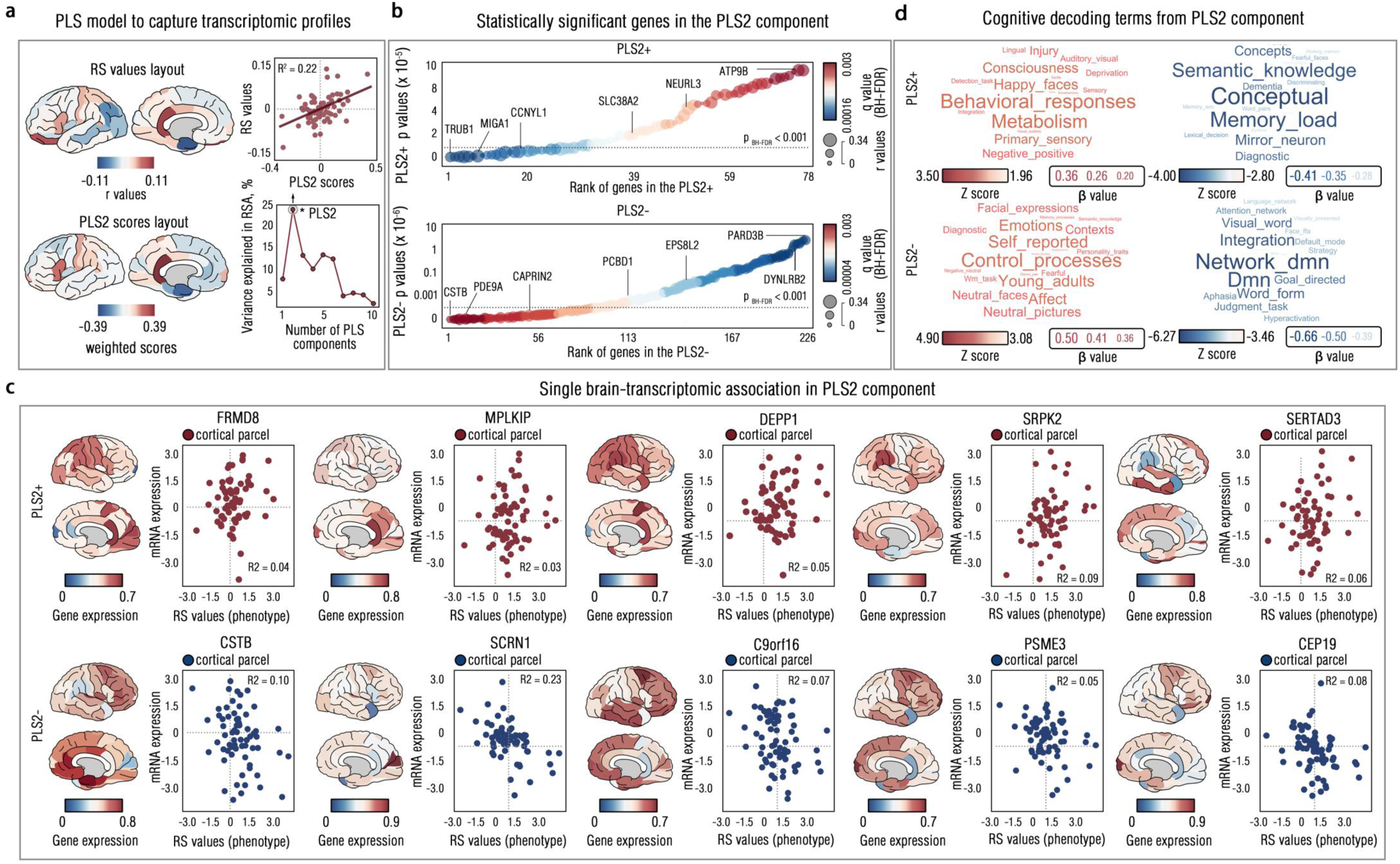
Imaging-transcriptomic markers and their brain ontology annotations. **a**, showed the brain maps to deviations-PPS representation similarity (RS) patterns and illustrated brain layout showing the second partial least square component (PLS2) pattern (left), as well the model fitting performance of PLS regression (right). **b**, illustrated the gene set that extracted from the PLS2 component, which reached statistical significance for overrepresented gene set (PLS2+, top) and underrepresented gene set (PLS2-, bottom) (Bootstrapping test at n = 1,000; *q* < .001, Benjamini-Hochberg correction). **c**, detailed linear association between gene expression (mRNA level) and neuroimaging phenotype (RS values) for top 5 genes with highest positive (PLS2+) and negative (PLS2-) weights. R^2^ indicated the explained variances from correlation models.

**Figure 7.**
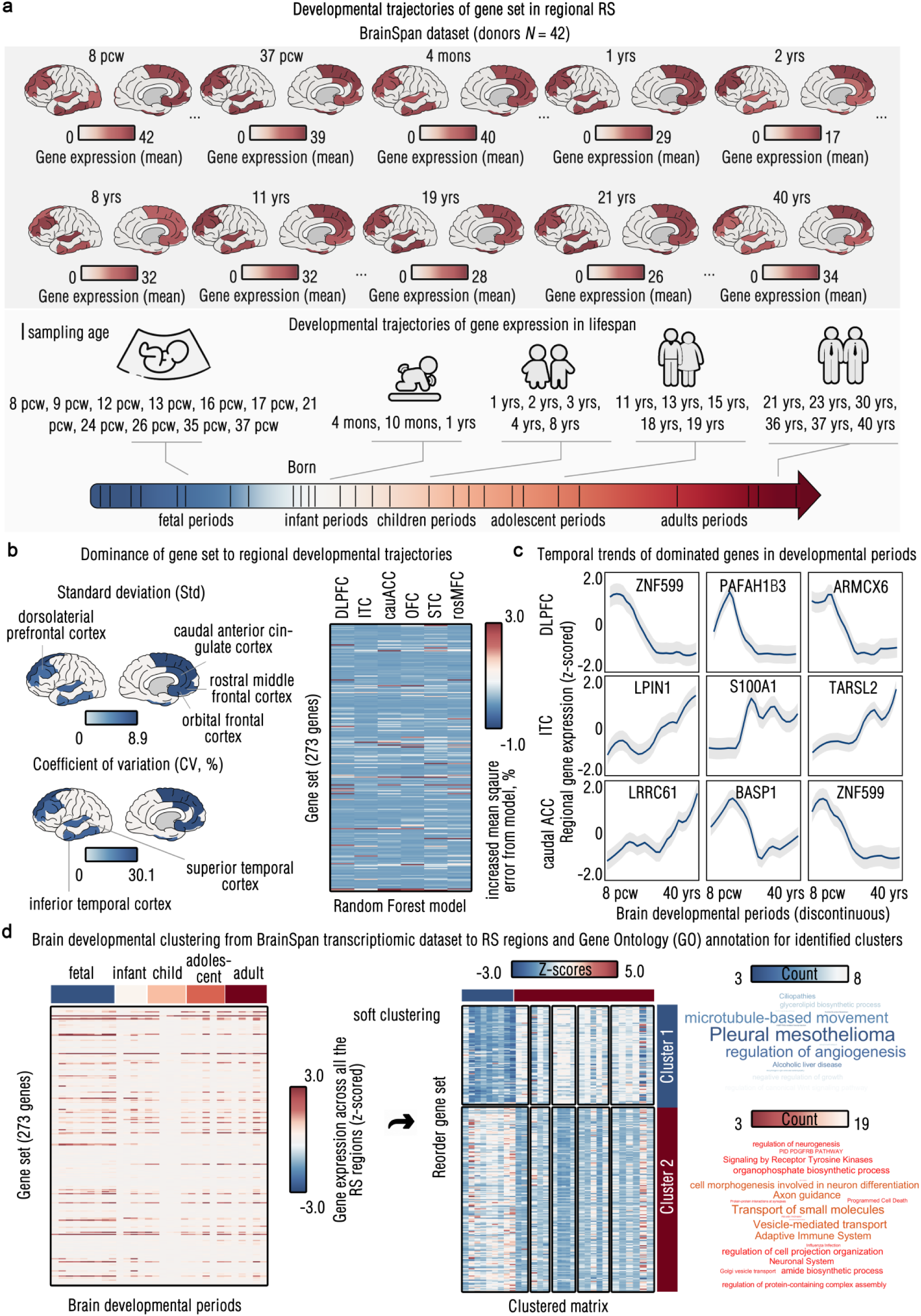
Brain transcriptomic developmental trajectories of PLS2 component. **a**, showed averaged gene expression of the PLS2 gene set (Top). Given that the sampling ages in the BrainSpan atlas were not continuous, we visualized all the sampling periods from these 42 donors to show the relative timestamps for the neurodevelopmental (Bottom). **b**, showed the temporal variants of gene expressions in six brain regions identified in the RSA, using standard deviations (SD) and coefficient of variations (left). In the right panel, relative contributions to the temporal predictive roles of these genes in brain regions by the random forest model were presented as quantified by increased mean square error from model (%). **c**, exemplified temporal trends of most contributing this prediction model for these brain regions, with shallow areas for indicating 95% Confidence Interval (CI) from Bootstrapping method (*n* = 1,000). **d**, presented identified clusters of the gene set x brain developmental trajectories (8pcw-40yrs) and their Gene Ontology (GO) enrichment.

Moving to multiscale neurobiological enrichment, we performed systematic enrichment analyses for this PPS-specific gene set - PLS2 component **(*see Methods*)**. Results revealed significant enrichment in vesicle-mediated transport (R-HSA-5653656) and small molecules transport (R-HSA-382551) (*q* < 5 × 10 ^-6^; **Figure 8a, Supplemental Results 11, Table S36**). These pathways were further observed in interconnecting with other GO categories related to molecular transports and cortical neurodevelopment **(Figure 8b; Supplemental Results 11, Table S37)**. Neurobiological annotation using the Molecular Signatures Database (MSigDB) indicated that these genes were significantly enriched in molecular pathways regulating both neuroinflammation and neuroimmunity, such as TNF-α, IFN-α and IL-6 **(Figure 8c and Supplemental Information 2)**. At the cellular level, significant enrichment was observed in midbrain neurons, particularly within the dorsal raphe nucleus, which plays a key role in serotonergic transports **(Supplemental Results 11, Table S38-39)**. In addition to such multiscale enrichment, bridging to psychopathological and clinical outcomes, these PPS-specific genes were found significantly enriching into psychiatric disorders and neurodegenerative diseases, particularly in cognitive impairment, dementia, and depression (*p* < .05, spin permutation test) **(Supplemental Results 11, Figure S6 and Table S40-41)**. Collectively, within a close-loop RDoC research framework, these findings delineated complicated multiscale neurogenetic mechanisms underlying PPS, suggesting to conceptualize PPS as a “brain disorder” implicating dysregulated molecular transport, neuroimmune response, and neuroinflammation.

**Figure 8.**
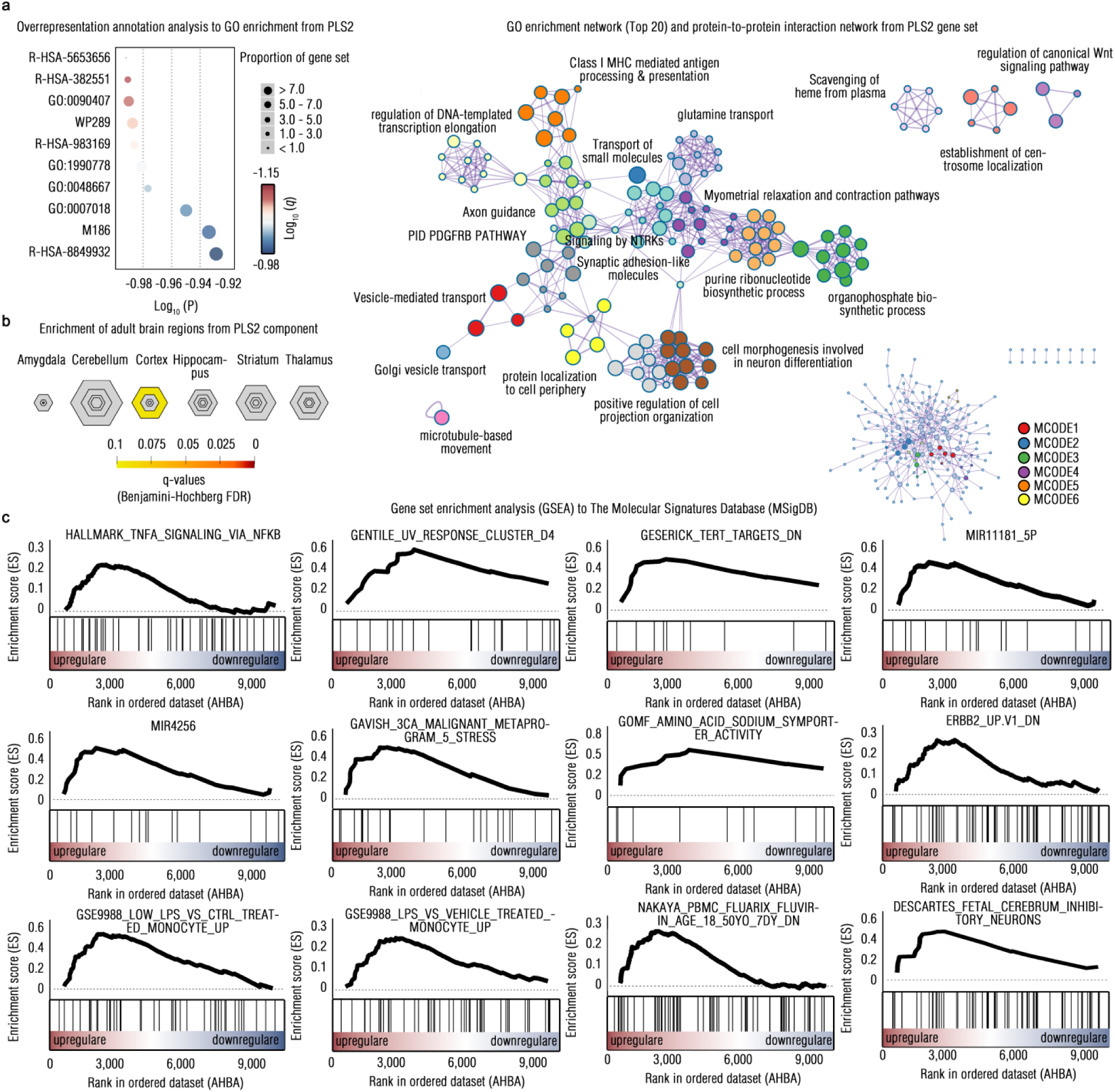
Enrichment analysis to PPS-specific gene set (PLS2). **a**, presented enrichment results of Gene Ontology (GO) at the Metascape, showing top 10 GO terms at the left panel and GO-to-Go/protein-to-protein interaction network at the right panel. Here, each node represents an enriched term and is colored first by its cluster descriptions. The Molecular Complex Detection (MCODE) algorithm was used to capture independent components (MCODE1 - 6). **b**, showed GSEA of PLS2 gene set to adult brain developments. The colorbar indicated the Fisher’s Exact p values, and the rhombus size indicated pSI threshold, ranging from 0.05 to 0.0001. **c**, illustrated the GSEA results from Molecular Signatures Database (MSigDB). Here, the enrichment plots enriched terms that reached stringent statistical thresholds (p < .01, FWE (Family-Wise Error) correction) are presented.

**Figure 9.**
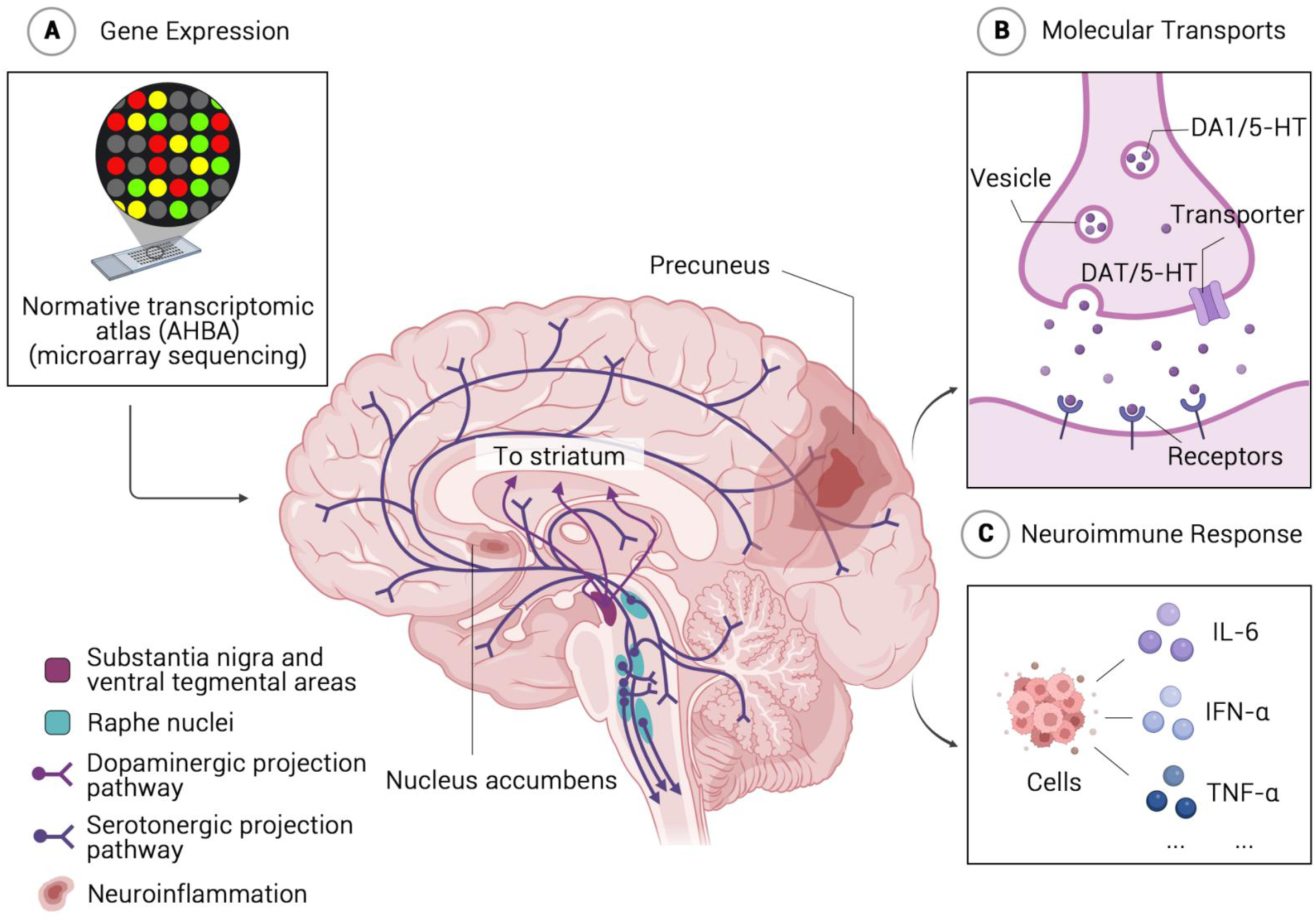
Theoretical schematic of neurogenetic dysregulation model for conceptualizing PPS as a “brain disorder”. **a**, described the microarray sequencing techniques to curate Allen Human Brain Atlas (AHBA) for disentangling brain-transcriptomic associations, particularly in identifying neurogenetic deviations localized in precuneus and NAcc. **b**, showed significant neurobiological enrichment of these PPS-specific neurogenetic characterizations, indicating dysregulations in the molecular transports implicating reward processing (e.g., DA1/5-HT) as well in neuroimmune and neuroinformmation systems (e.g.,TNF-α, IL-6).

### Neurogenetic Dysregulation Model: Theoretical Conceptualization of PPS as a “Brain Disorder”

To synthesize multiscale neurobiological markers and elucidate the neurogenetic mechanisms underlying PPS, we propose a conceptual framework-the neurogenetic dysregulation model. This model conceptualizes PPS as a “brain disorder” arising from neurogenetic dysregulations in neurotransmitter and neuroimmune systems, particularly those involved in reward processing. At the macroscale level, PPS is associated with neurogenetic variations in neurodevelopmental morphologies, particularly in reward-related structures such as NAcc and DMN **(Figure 8a)**. At the microscale and molecular levels, these morphological deviations may, in turn, reflect dysregulation within key dopaminergic and serotonergic pathways (e.g., D1, DAT, 5-HTT; **Figure 8b**) as well as neuroimmune responses linked to heightened neuroinflammation (e.g., TNF-α, IFN-α; **Figure 8c**). Supporting this argument, these PPS-specific neurogenetic markers are closely associated with known psychiatric or clinical conditions implicating in reward processing deficits. On balance, as integrated from multiscale neurobiological markers, this neurogenetic dysregulation model theoretically conceptualized PPS as a subclinical “brain disorder” resulting from dysregulations in neurotransmitter and neuroimmune systems involving into reward systems.

## Discussion

This study leveraged an 8-year longitudinal twin cohort and brain normative modeling approach to systematically investigate the neurogenetic substrates underlying psychopathological procrastination (PPS), an understudied subclinical phenotype. Within the RDoC research framework, our findings revealed moderate heritability of PPS and identified neural predictors, with adolescent morphological deviations forecasting PPS phenotyping in their adulthood. Notably, these brain-symptomatology mappings shared genetic factors. Integrating PPS-specific neural representation patterns into genetic and multiscale neurobiological maps, we clarified PPS’s gene expression markers and identified their enrichment into molecular transports in serotonergic/dopaminergic pathways, neuroimmune and neuroinflammation response. These findings offer a close-loop framework bridging genetic associations to neurobiological mechanisms, supporting conceptualization of PPS as a “brain disorder” characterized by dysregulated neurotransmitter systems and neuroimmune processes in reward system.

A key finding of this study is that psychopathological procrastination (PPS) exhibits moderate heritability. While previous research has established the heritability of procrastination as a trait,(*1, 41–43*) our findings provide the empirical evidence supporting a genetic basis for procrastination within the context of subclinical psychiatric psychopathology. While procrastination, as a personality trait, has shown to be heritable, these estimates have largely varied, ranging from 0.12 to 0.46 in adults.(*41, 42, 59*) This heterogeneity suggests the latent subgroups that may confound phenotyping in procrastination. Supporting this notion, analogous patterns have been consistently observed in psychiatric disorders such as major depressive disorder and post-traumatic stress disorder, where subclinical pieces within cohorts can show significant variability in psychopathological mechanisms and treatment responses.(*60–62*) Furthermore, compared to personality traits, which typically exhibit higher heritability, neuropsychiatric and neurological disorders show moderate heritability due to complex and multifactorial etiologies.(*63–66*) Collectively, our findings support the conceptualization of PPS as a heritable subclinical phenotype, rather than merely a personality trait.

Another key finding of this study is the identification of neurogenetic markers underlying PPS, with substantial shared genetic variations between neurodevelopmental deviations and this subclinical phenotype. Normative modeling, a state-of-the-art neuroimaging approach, quantified individual morphological deviations compared to a population-based brain atlas, offering a more precise measure of neurodevelopmental differences.(*47, 49, 50, 67*) This method is particularly effective for identifying neuromarkers in heterogeneous populations, such as those with depression, schizophrenia and autism spectrum disorder (Parkes et al., 2021; Sun et al., 2023; Wang, 2022).(*51, 68–70*) Using this approach, our results show the prediction of PPS by neurodevelopmental morphological deviations, suggesting that they could serve as “precision biomarkers” for early identification of individuals at risk for this subclinical phenotype.

Rather than attributing mental disorders solely to case-control brain differences, we conceptualize PPS as deviations from normative brain developments.(*49, 67*) By doing so, we found that neuroanatomical deviations in the nucleus accumbens (NAcc) and default mode network (DMN) were uniquely associated with PPS, distinguishing it from general procrastination traits.(*37, 39, 71*) While procrastination traits are often linked to brain areas involved in self-control, emotional regulation and episodic thinking, such as DLPFC, ACC and hippocampus,(*37*) PPS is more specifically tied to reward systems, as reflected in the NAcc. As a key node in the mesolimbic system, brain abnormalities in the NAcc have disentangled to function on inhibiting reward computational processing and consequently perpetuating devaluation on action motivation.(*72–74*) This aligns with the core concept of procrastination: individuals irrationally delay tasks when they undervalue rewards or punishments, leading to insufficient motivation to act.(*40, 75*) In addition to involvements into reward processing neurally represented by NAcc, the DMN has been well-studied in supporting reward learning by encoding task-related motivation through its top-down projections to the limbic system.(*76–78*) Thus, rather than stemming from cognitive vulnerability such as deficient self-control or episodic thinking solely, PPS may reflect inherent psychopathological dysfunctions in the reward systems, as our findings suggested.

Our genetic analysis further links these brain deviations to PPS, bridging neurogenetic markers with psychopathological mechanisms. Procrastination has been theorized to share a genetic basis with impulsivity, reflecting an evolutionary adaptation of impulsivity to modern societal demands.(*1, 41, 42*) Standing for this point, brain regions associated with procrastination traits are largely overlapped into neocortex involving into impulsivity, particularly in the DLPFC.(*79–81*) However, we found that PPS is genetically mapped to subcortical areas, particularly the NAcc, which are less implicated in impulsivity and more directly involved in reward processing, showing neural representation distinction between individual trait and subclinical condition. This aligns with growing evidence linking cognitive traits to psychiatric disorders.(*82, 83*) Although common genetic and phenotypic underpinnings exist between behavioral tendencies and psychiatric disorders, these pairs often display divergent neural representations, with psychiatric disorders showing more pronounced neuropsychopathological abnormalities than their behavioral counterparts.(*84, 85*) This collectively suggests that PPS may be governed by distinct genetic pathways, setting it apart from impulsivity and reinforcing the idea that PPS is a unique subclinical “brain disorder” underpinned by neurogenetic dysregulation in reward-related circuits.

A significant contribution of this study is the identification of serotonergic and dopaminergic pathway dysfunctions, alongside their interaction with neuroimmune responses and neuroinflammation, as key mechanisms underlying PPS. We identified significant mappings from these genetic markers to neurotransmitter systems, particularly in DAT, D1 and 5-HTT. As the psychopathological factor (*p* factor) theory delineated, key symptomatology of procrastination, such as irrational postponement and inattentiveness, may represent a transdiagnostic factor across externalizing problems,(*43, 86, 87*) with significant genetic vulnerabilities and molecular dysregulation in NAcc-regulated dopamine and serotonin systems.(*88–90*) Moreover, the interplay between serotonergic and dopaminergic dysregulation and neuroimmune responses points to a neuroinflammatory mechanism in PPS. There is ample evidence that hyperactivation of these neurotransmitter systems modulate neuroimmune and autoimmune responses, leading to the accumulation of neuroinflammatory factors,(*91, 92*) which contribute to the pathophysiology of various neuropsychiatric externalizing symptoms.(*93–96*) More importantly, the reciprocal interaction between neurotransmitter’s dysregulation and neuroinflammation suggests a downward spiral: hyperactivation of these neurotransmitter systems exacerbates neuroinflammatory sensitization, while chronic neuroinflammation perpetuates serotonergic and dopaminergic overexpression, further amplifying psychiatric psychopathology.(*97–101*) Further supporting this, previous studies have linked catechol-O-methyltransferase (COMT) polymorphism to chronic procrastination, a genetic moderator role in neuroimmune-driven neuroinflammatory responses.(*102–104*) These findings may create a new insight into the psychopathology of PPS, suggesting interactions of neuroimmune and neuroinflammatory responses in its neurogenetic mechanisms.

Despite its contributions, this study has several inevitable limitations. First, PPS has not yet been formally categorized within established diagnostic systems, such as DSM-5 or ICD-10. As a result, while this study provides valuable empirical insights into the neurogenetic mechanisms underlying this subclinical condition, these findings are not directly relevant to clinical implications. Another limitation is the restricted external generalizability of the findings. Due to the inherent difficulties of curating longitudinal twin neuroimaging cohorts, the sample size is relatively limited, and no independent replication samples were included. This emphasizes the need for future research with larger and more diverse populations to validate and extend these findings. Finally, while this study employed the RDoC framework and robust normative neurobiological datasets to investigate the multiscale psychopathologies of PPS, the connections between neurogenetic markers and molecular, cellular and other biological ontologies remain indirect. Therefore, these findings should be interpreted within a conceptualized framework, avoiding overstatements or unwarranted generalizations. To tackle with this point, further empirical studies directly examining the multiscale neurobiological associations of PPS are urgently needed.

In conclusion, using a 8-year longitudinal twin cohort and population-based normative modeling approach, we identified unique neurogenetic markers of PPS implicating psychopathological processes within reward systems. Leveraging the RDoC framework, we mapped these markers to multiscale neurobiological systems, demonstrating significant enrichment in serotonergic and dopaminergic neurotransmitter pathways, as well as their potential interactions to neuroimmune and neuroinflammatory responses.

Synthesizing these insights into a unified “neurogenetic dysregulation model”, we conceptually propose PPS as a subclinical “brain disorder” characterized by dysregulated neurotransmitter and neuroimmune/neuroinflammatory process within reward systems.

Collectively, this study provides both empirical evidence and a theoretical framework for illuminating the psychopathology of PPS, supporting its classification as a subclinical condition posing unique neurogenetic foundation.

## Materials and Methods

### Participants and Ethic Statement

This 8-year prospective twin neuroimaging cohort was drawn from the Beijing Twins Brain-Behavior Association Dataset.(*105, 106*) The cohort encompassing 162 twin pairs (*N* = 324) with within-pair same sex was established in 2015 summer, where 77 twin paris (*N* = 154, 55.8% female pairs, 40 monozygotic twins [MZ] pairs; 37 dizygotic [DZ] pairs) were successfully followed up for the phenotyping of psychopathological procrastination (PPS) in 2024 spring. For genetic brain-behavior prediction analysis, six pairs were excluded due to neuroimaging data incompleteness (*see below*). Thus, the final sample for investigating the genetic neuromarkers of PPS consisted of 71 twin pairs (*N* = 142, 56.3% female pairs, 38 MZ pairs, 33 DZ pairs; mean age = 20.0 years, SD = 2.4).

None of the included participants had a history of psychiatric or neurological disorders, nor a family history of genetic diseases. Additionally, all twins reported no craniocerebral injury, surgeries, or traumatic life events at the time of recruitment and follow-up. The study protocol and analytic pipeline were approved by the Institutional Review Board (IRB) of the Institute of Psychology (Chinese Academy of Sciences, China, #H20037) and the IRB of the Faculty of Psychology (Southwest University, China, #H24121).

### Measurements of Psychopathological Procrastination

The 3PDQ-9, a DSM-based subclinical psychopathological procrastination questionnaire, was used to assess PPS symptomatology.(*19*) This nine-item scale was developed to phenotype subclinical symptoms of PPS using a 5-point Likert scale, with higher total scores indicating greater severity of symptoms. The questionnaire integrates core constructs of procrastination and its associated psychiatric symptomatology, such as items of “*I cannot change or control my procrastination behavior by myself.*” (difficulty in self-regulation); “*I procrastinate tasks despite that I know they are very important to me.*” (pervasive irrational postponement); “*The thought of ‘Put off this task’ always irrupts in my mind when I begin to do this task or I am currently working for this task.*” (ADHD-like symptoms); and “*I always feel bored and depressive for many activities.*” (depressive symptoms) (**Supplemental Methods 1, Table S1**). The 3PDQ-9 has demonstrated strong diagnostic performance in identifying subclinical psychopathological procrastinators (Chen et al., 2024).

### Neuroimaging Data Acquisition and Processing

All included twins underwent T1-weighted structural imaging using a GE 3.0T MRI scanner (GE Healthcare, Waukesha, US). Brain structural images were acquired with a magnetization-prepared rapid acquisition gradient echo (MPRAGE) sequence using the following scanning parameters: repetition time (TR) = 2520 msec, echo time (TE) = 3.37 msec, flip angle = 7^°^, inversion time (T1) = 1100 msec, number of slices = 144, and slice thickness = 1.33 mm. To minimize head motion, participants’ heads were stabilized with sMRI-compatible foam padding within a 8-channel head coil.

T1-weighted neuroimaging data were preprocessed using FreeSurfer (Version 7.2.0; https://surfer.nmr.mgh.harvard.edu). We followed the standardized automatic “recon-all” pipeline, which includes multiple steps for cortical and subcortical morphometric estimation. First, head motion correction and slice-time correction were applied using an iterative affine registration algorithm.(*107*) Intensity normalization was then performed to correct for bias field inhomogeneities using a Gaussian class model.(*108*) Skull stripping was conducted using a hybrid watershed/surface deformation method to isolate brain tissue.(*109*) Subcortical structures were segmented and labeled using a probabilistic atlas-based Bayesian framework incorporating manually labeled training data.(*110*) For cortical surface reconstruction, white matter was segmented, tessellated, and refined using deformable surface algorithms with topology correction.(*111*) The reconstructed surfaces were then registered to a spherical atlas for inter-subject alignment, optimizing sulcal and gyral feature correspondence.(*111*) Finally, cortical surfaces were parcellated into the 68 anatomical regions based on the Desikan-Killiany atlas,(*112*) from which cortical thickness (CT) and surface areas (SA) were computed, region-by-region. Moreover, subcortical volume (SV) were calculated from 14 subcortical regions using the Automatic Segmentation (Aseg) atlas.

### Heritability and Bivariate Genetic Association Estimation

#### Additive Genetic - Common Environment - unique Environment (ACE) Model

To estimate the heritability of PPS in this twin cohort, we employed a univariate ACE model, a reliable statistical approach in behavioral genetics, implemented via the *OpenMx* package (v2.18.1).(*113*) This mode decomposes the total phenotypic variances (*V*_p_) of PPS into three ingredients: addictive genetic variance (*A*), shared environmental variance (*C*) and non-shared environmental variance (*E*), where *A* represents the heritability estimate (*h*^2^). Both MZ and DZ twin pairs were included in this analysis, enabling ACE modeling under the standard assumption that MZ twins shared 100% of their genes material (*r* = 1), whereas DZ twins share, on average, 50% (*r* = 0.5). Given this statistical genetic relatedness, any excess within-pair similarity in MZ twins relative to DZ twins is attributed to additive genetic influences, allowing for the estimation of phenotypic heritability.(*114*)

Maximum likelihood estimation (MLE) was used to fit the model by leveraging the covariance structure of the phenotype across twin pairs. This approach quantified the relative contributions of *A*, *C*, and *E* as proportions of the total phenotypic variance (*V*_p_ = *A* + *C* + *E*), yielding estimates for heritability (*a*^2^ = *h*^2^ = *A*/*V*_p_), shared environmental influence (*c*^2^ = *C*/*V*_p_) and non-shared environmental influence (*e*^2^ = *E*/*V*_p_). To identify the most parsimonious model, nested submodels (e.g., AE, CE, E) were compared against the full ACE model using likelihood ratio tests and model fit indices, including −2 Log Likelihood (−2LL), Akaike Information Criterion (AIC) and Bayesian Information Criterion (BIC). The model showing best fits would be determined the final one for drawing statistical conclusions (**Supplemental Methods 2, Table S2**). Prior to ACE modeling, all phenotypic measures were adjusted for age and sex to account for potential confounding effects.

### Bivariate ACE Model and Cholesky Decomposition

To estimate the genetic associations (*r*_g_) between brain morphological deviations and PPS, we capitalized on a bivariate ACE model using the *OpenMx* package. This model decomposes the total phenotypic variance and covariance of PPS and morphological deviations into three latent components (**see *Equation 1***) of ACE. We applied Cholesky decomposition to partition these components, extracting their relative contributions to the variance of each phenotype and their covariance structure. This was achieved by reformulating the covariance matrix into a lower triangular form, processed sequentially. In this decomposition, latent factors were structured hierarchically: the first layer (*A*_1_, *C*_1_, *E*_1_) captures shared variance jointly influencing both PPS and brain morphological deviations, while the second layer (*A*_2_, *C*_2_, *E*_2_) accounts for residual variance specific to brain morphological deviations. Once the Cholesky decomposition was performed, MLE was used to estimate these latent factors from the covariance matrices, yielding statistical coefficients for genetic correlation (*r*_A_ = *r*_g_, **see *Equation 2***), shared environmental correlation (*r*_C_, **see *Equation 3***) and non-shared environmental correlation (*r*_E_, *see* ***Equation 4***):

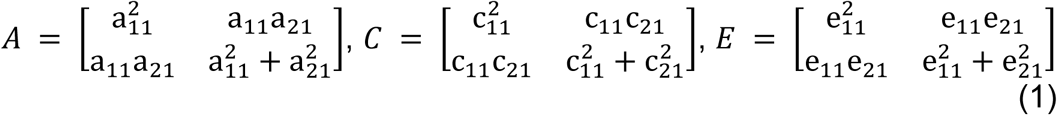

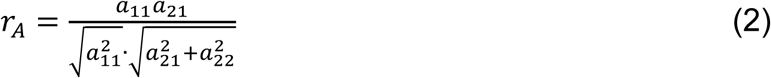

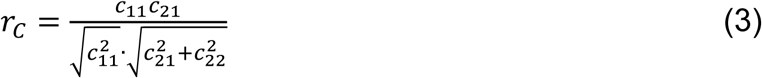

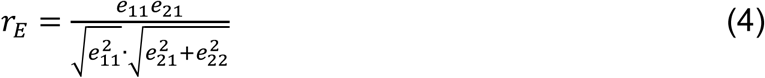

In addition to estimating genetic associations, we further assessed the phenotypic correlations (*r*_p_) between PPS and brain morphological deviations, as well as their heritability (*h*^2^). These estimates were derived by partitioning additive variance into contributions from A, C and E respectively (**see *Equation 5***):

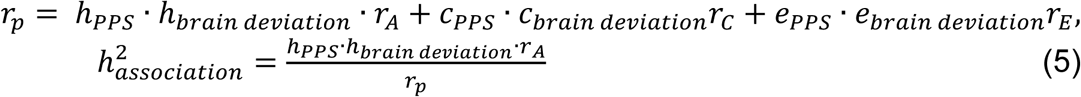

Model evaluation and selection followed the same approach as in the univariate model. Model fit was assessed using likelihood ratio tests, comparing the full ACE model with nested alternatives models (e.g., AE, CE, E) based on −2LL, AIC, and BIC. The statistical significance of the genetic correlations was determined using likelihood-based confidence intervals. To control for potential confounding effects, phenotypic data were adjusted for age and sex prior to analysis.

### Normative Modeling and Brain-Behavior Prospective Prediction

Using the population-based neuroimaging cohort curated by the ENIGMA Lifespan Working Group (*N* = 37407, 19826 females, aged range: 3-90 years), normative morphometric models for CT, SA and SV were pretrained using the multivariate fractional polynomial regression (MFPR) algorithm.(*57*) These models accounted for sex and age-related effects on regional morphometric characteristic, generating lifespan neurodevelopmental chart categorized into percentiles.(*115*) To estimate individual morphometric deviations from these normative neurodevelopmental charts, twins with eligible brain morphological data were analyzed to determine the extent of overdevelopment or underdevelopment in specific regions. Standardized Z-scores (*Z*_nd_) were then computed to quantify these neurodevelopmental deviations and adjust for inter-subject heterogeneity (**see *Equation 6***), where *y*_nd_ represents the observed morphometric values, while ⋀*y*_nd_ denotes the predicted values generated by the normative model. The denominator corresponds to the model’s root mean square error (RMSE). For each participant, Z-scored morphometric deviations were estimated iteratively across the entire brain, encompassing all cortical and subcortical regions. A positive *Z* scores indicates overdevelopment, whereas a negative Z-score reflects underdevelopment relative to the normative chart. Statistically significant neurodevelopmental anomalies were defined as Z-scores exceeding ±1.96 (corresponding to p < .05 in a Gaussian distribution).

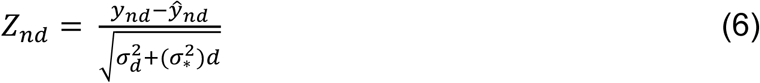

To identify brain-behavior mappings, we implemented generalized linear mixed-effects models (GLMMs) for investigating predictive of regional morphometric deviations to PPS phenotype, region-by-region, using the *lme4* package (version 1.1.24). Age, sex, zygosity, and individual intercepts were included as no interests of covariates. All *p* values from GLMM analyses were corrected for multiple comparisons using the Benjamini-Hochberg procedure, with a significance threshold of *q* < .05.

To interpret the neurofunctional ontologies underlying multiple regional deviations, we applied brain network enrichment significance analysis (NESA), a statistical framework designed to test whether regional brain-behavior associations are enriched within known brain networks as parceled by the Yeo-7 atlas.(*116*) Specifically, based on the *β* coefficients representing brain-behavior associations, we first ranked the regions in descending order to generate a reference map (*T*_v_). Using this map, each region was assigned a running score depending on whether it belonged to a given network. This iterative process across all regions yielded a running sum scores for each network. The enrichment score (ES) for a given network from reference map was then computed as the maximum deviation from zero. For each brain network, ES values quantified the degree of spatial enrichment, serving as the observed metric. To assess statistical significance, a self-constrained null model was generated by randomly shuffling participants with permutation process (*n* = 5,000). *P*-values were calculated as the proportion of permuted ES values exceeding the observed ES value. Multiple comparisons were controlled using the Benjamini-Hochberg correction to mitigate inflated false-positive rates.

### Inter-Subject Representation Similarity Analysis

Beyond univariate brain-behavior predictions, we conducted inter-subject representation similarity analysis (IS-RSA) to capture the multivariate neural representation similarity (RS) pattern of the PPS phenotype. Specifically, whole-brain deviations were first arranged into a 1 × 68 vector (representing cortical parcels) for each participant and then correlated across participants to generate a 70 × 70 inter-subject neural representation similarity matrix (nRSM), where each element represents a correlation coefficient (*r*). To quantify individual variability in whole-brain deviation patterns, we computed a neural representation dissimilarity matrix (nRDM) by subtracting the nRSM values from 1.

Subsequently, to establish a multivariate phenotypic pattern of PPS, we calculated inter-subject 2-dimensional Euclidean distance between PPS scores across all subjects, generating a 70 × 70 inter-subject behavioral representation similarity matrix (bRSM). As in line with construction of nRDM, we derived the bRDM to illustrate multivariate patterns in PPS patterns within the cohort. Next, we extracted the upper triangular portion of both nRDM and bRDM (excluding the diagonal) and vectorized them into a 1 × 2,415 nRDM vector and a 1 × 2415 bRDM vector. The whole-brain multivariate neural representation of PPS was then quantified by computing Spearman’s rank correlation between these vectors. Separate multivariate neural representation matrices were constructed for CT, SA and SV. In addition to whole-brain deviations, regional deviations were used to construct nRDMs by computing inter-regional Euclidean distances, following the same approach described above. Each region was vectorized into a 1 × 2,415 nRDM vector and correlated with the 1 × 2,415 bRDM vector, region-by-region. A statistically significant positive *r* value for a given region demonstrated its multivariate neural pattern similar to represent the PPS phenotype. Iterating this process across all regions generated a whole-brain RS pattern, mapped by region-wise PPS *r* values. For all the correlations, age, sex and zygosity of twins were inputted as covariates of no interests for statistical corrections. All the *p* values have been adjusted for multiple comparisons by the Benjamini-Hochberg correction at *q* < .05.

### Biological Enrichment of Neural Representation Pattern Underlying the PPS

After generating a whole-brain RS map representing PPS, we applied Neuromap (https://github.com/netneurolab/neuromaps), a cutting-edge annotation method, to decode neurobiological ontologies enriched by this PPS-specific neural representation pattern.(*58*) Specifically, we transformed the whole-brain map from its native coordinate system into multiple standardized coordinate systems to ensure comparability with over 40 reference maps, which span various neurobiological domains, including microstructure (e.g., T1w/T2w ratio), metabolism (e.g., cerebral blood flow), neurofunction (e.g., gradients), neuroexpansion (e.g., evolutionary expansion), electrophysiology (e.g., delta power), neurotransmitters (e.g., 5-HT) and genomics (e.g., gene expression).(*117–120*) To assess whether the PPS-specific neural representation pattern was spatially enriched in these reference maps, we computed Pearson’s correlations between the whole-brain map and each reference map individually. Statistical significance was determined using a spin permutation test, which accounts for spatial autocorrelation.(*121*) All *p* values underwent statistical corrections using Benjamini-Hochberg method, accounting for multiple comparisons.

### Gene Expression Profiles and Close-loop Multiscale Neurobiological Enrichment Allen Human Brain Atlas and BrainSpan Lifespan Transcriptomic Dataset

To capture neurogenetic markers underlying the PPS in a close-loop RDoC research framework, we incorporated normative transcriptomic data curated from the Allen Human Brain Atlas (AHBA), a comprehensive dataset mapping the transcriptional landscape of the human brain.(*122*) This dataset includes six neurotypical donors (*N* = 6), with 3,702 anatomical tissues for sequencing mRNA levels from microarray. Preprocessing followed established protocols,(*123*) with modifications to align the data with the DK atlas, resulting in a normative gene-brain matrix of 68 (parcels) × 10,027 (genes) (**see Supplemental Methods 3**). Furthermore, the PPS-specific whole-brain representation map was transformed into a 68 (parcels) × 1 (*r* values) vector, representing multivariate representation similarity between regional brain deviations and PPS phenotype identified earlier. To extract gene expression profiles associated with this PPS-specific neural representation pattern, we applied a partial least square (PLS) regression model, fitting the gene-brain matrix (68 × 10027) to the neural vector (68 × 1). Statistical significance of latent components decomposed from this model was assessed using permutation testing (*p* < .05, *n* = 5,000). Since the second latent component (PLS2) reached statistical significance, we further calculated *Z*-scored weights and corresponding *p* values for each gene within this component using a bootstrapping approach (*N* = 5000). To adjust for multiple comparisons, only genes passing the significance threshold (*p* < 5 × 10^-3^; *Z* > 3.7 for PLS2+, *Z* < −3.7 for PLS2-) were retained following Benjamini-Hochberg correction.

Considering longitudinal predictions from brain deviations to PPS, we further investigated the neurodevelopmental trajectories of these transcriptomic profiles (i.e., genes identified in PLS2). Using the BrainSpan lifespan gene expression dataset, which includes 42 donors spanning from 8 post-conception weeks (pcw) to 40 years (*N* = 42), (*124*) we first constructed a 52,377 gene × 30 age groups × 524 brain samples (regions) matrix to quantify neurodevelopmental pattern of gene expression. To align this dataset with PLS2 gene set and brain regions identified to representing PPS phenotype, we extracted a subset of 273 overlapping genes across 30 age groups in 6 cortical regions, resulting in a PPS-specific longitudinal matrix (273 × 30 × 6). Transcriptomic levels would be averaged if multiple samples sequenced within the same age group or cortical parcels.(*125*) We constructed a random forest model (RFM, tree constrain = 1000) to fit gene expression profiles (273 genes × 30 ages) to neurodevelopmental trajectories (1 × 30 ages).

Bootstrapping (*N* = 5000) was used to estimate statistical significance (*p* < .05) and 95% confidence interval (CI). Following model training, we utilized the Increased Mean Square Error (IMSE) algorithm to assess the relative dominance of each gene in explaining neurodevelopmental transcriptomic trajectories. Specifically, in this RFM, we recorded the mean squared error (MSE) for each gene. Then, we randomly shuffled the transcriptomic levels of each gene and refitted the model, one-by-one. The new MSE was then compared to the original MSE, and the difference (IMSE) quantified the gene’s relative contribution to the model’s predictive performance, with higher IMSE values indicating greater dominance in explaining neurodevelopmental trajectories.(*126*) To further ensure statistical reliability, we calculated the proportion of change between the new and original IMSE values. All analyses were conducted using the “randomForest” and “randomForestExplainer” packages in R (version 4.1.2, R Core Team, 2021).

Upon estimating dominance of all 273 genes, we calculated *Z*-scored IMSE proportions and averaged them for clustering analysis to enhance the interpretability of these transcriptomic trajectories. Rather than using conventional clustering algorithms with rigid boundary (e.g., *k*-means), we employed the fuzzy clustering algorithm “Mfuzz” to identify modules with overlapping temporal gene expression patterns.(*127*) This method is particularly suited for clustering microarray genetic data, as it incorporated soft boundary that allow genes to belong to multiple clusters, effectively capturing temporal autocorrelations in gene expression.(*127*) A two-cluster solution was determined as optimal, as it maximized within-cluster variances (**Supplemental Methods 4, Figure S1-2**). Although a three-cluster structure showed comparable variance, it contained only three unique genes, limiting its neurobiological interpretability.

### Macoscale Brain-Imaging and Disease Associations and Multiscale Neurobiological Ontologies Enrichment Analysis

To establish a close-loop framework of understanding these PPS-specific neurogenetic characteristics identified above, we capitalized on the Gene Annotation by the Macroscale Brainimaging Association (GAMBA, http://dutchconnectomelab.nl/GAMBA/) dataset, the Metascape dataset (https://metascape.org/gp/index.html), Cell-type and Tissue-type Specific Expression Analysis datasets (CSEA and TSEA, http://doughertytools.wustl.edu/), CellMarker dataset (http://xteam.xbio.top/CellMarker/index.jsp), Annotation of Cell Types (ACT, http://xteam.xbio.top/ACT/) dataset and Molecular Signature Database (https://www.gsea-msigdb.org/gsea/msigdb/index.jsp). These datasets curated normative macroscale brain maps for examine brain-behavior associations at a systems level, and provided both molecular and cellular biological ontology sets for gene set enrichment analysis (GSEA).(*128*)

Using the GAMBA dataset, we inputted PLS2+ and PLS2-gene set with descending order based on the *Z*-scored weights respectively. This approach allowed us to spatially regress the whole-brain expression patterns of PPS-specific genes onto normative macroscale brain biological ontologies and disease-associated gene expression maps. These included intrinsic brain network (e.g., functional connectome within default mode network), brain cognitive ontologies (e.g., theory of mind), cognitive terms (e.g., control process), cortical metabolism (e.g., cerebral blood flow, CBF), evolutionary cortical expansion and neuropsychiatric/neurological disorders (e.g., ADHD, stroke).(*129*) To ensure statistical specificity and robustness, we implemented spatial general linear models (GLM) with a spin null-brain-gene permutation test to assess statistical significance (**Supplemental Methods 5**). *P* values inflated from multiple comparisons were corrected using the Bonferroni-Holm method.

To extend our analysis from macroscale brain ontologies to molecular and cellular enrichment, we applied the full PLS2 gene set, reordered using the same ranking rule, to the “Metascape” data platform. Metascape is a widely-used tool for decoding neurobiological ontologies via over-representation analysis (ORA) across multiple normative datasets (e.g., Gene Ontology, GO; Kyoto Encyclopedia of Genes and Genomes, KEGG), enabling the identification of enrichment in known molecular and cellular functions. For obviating statistical overestimation, we further decoded the PLS2 gene set using the GSEA and TSEA assessing gene enrichment specificity across cell type and tissues, respectively. The specificity index probability (pSI) was calculated to determine the likelihood that a given gene set was significantly enriched in a particular biological category relative to background distributions across multiple statistical thresholds.(*130, 131*) All the *p* values for the gene enrichment were adjusted by the Benjamini-Hochberg correction. To incorporate the latest knowledge on cellular markers, we further mapped this gene set onto the CellMarker and ACT datasets, which provide the newest normative cell type markers, with a statistically robust weighted hypergeometric enrichment analysis model.(*132*) Finally, using the Molecular Signatures Database (MSigDB) including more than ten-thousands biological ontologies, we conducted the gene set enrichment analysis (GSEA) by the GSEA software (Version 4.3.3 released).

Consistent with prior enrichment analyses, statistical significance was estimated via permutation test (*N* = 1,000) (**Supplemental Methods 6**).

## Acknowledgments

We do appreciate Dr. Xuerong Liu for assistance on proofreading and thank to Prof. Tingyong Feng for scientific comments on theoretical conceptualization.

## Funding

National Natural Science Foundation of China (32300907, 72033006, CZY, ZY)

Chongqing Natural Science Foundation (General Project, CZY)

PLA Talent Program Foundation (2022160258, CZY)

AMU-RD Scholar Foundation (202211001, CZY)

## Author contributions

Conceptualization and Writing—original draft: HYY, YCT, CZY

Methodology: HYY, ZD, JYN

Investigation: HYY, FQC

Visualization: CZY, HYY, LW

Supervision: CZY, ZY, JKB

Writing—review & editing: JKB, LW, ZY, CZY

## Competing interests

Authors declare that they have no competing interests

## Data and materials availability

Data supporting reproducibility of this study are available in the main text and the supplementary materials. The raw genetic neuroimaging data are available under restricted access for data legal restrictions. Specifically, these genetic neuroimaging data are collected by the Beijing Twin Data Collection Project, which has not yet completed. Thus, given biometric data that are undergoing collection in an ongoing project, the legal censorship to share them from data regulation authorities is not available at this time, until this project is fully completed. For scientific collaborations or research purposes, any researchers can obtain access by reaching out Dr. Yuan Zhou.

## Notes

### Competing Interest Statement

The authors have declared no competing interest.

