## Supplementary Materials for "Mapping neurogenetic characteristics of psychopathological procrastination using normative modeling in a prospective twin cohort"

Yuanyuan Hu *et al.*

*Zhiyi Chen,; Yuan Zhou,

**This PDF file includes:**

Supplementary Text
Figs. S1 to S6
Tables S1 to S41
References (1 to 3)

**Supplementary Text**

**Supplemental Methods**

**1. Measurements of Psychopathological Procrastination**

Psychopathological procrastination (PPS) was assessed using the Pathological Procrastination Diagnostic Questionnaire (3PDQ-9; Chen et al., 2024), a standardized nine-item instrument designed to phenotype PPS within a subclinical psychiatric framework. The 3PDQ-9 integrates core procrastination constructs with psychiatric symptomatology, capturing pervasive and irrational task postponement alongside associated emotional and cognitive dysfunctions. Each item is rated on a five-point Likert scale (1 = strongly disagree to 5 = strongly agree), with higher total scores indicating greater PPS severity. In the present study, PPS was operationalized as a continuous variable, with standardized scores used for all subsequent statistical analyses to mitigate scale distribution biases. Full items of this scale can be found in the **Supplemental Table S1**.

| Items | 1 | 2 | 3 | 4 | 5 |
|---|---|---|---|---|---|
| I cannot change or control my procrastination behavior by myself. | | | | | |
| The thought of "Put off this task" always irrupts in my mind when I begin to do this task or I am currently working for this task. | | | | | |
| I procrastinate tasks despite that I know they are very important to me. | | | | | |
| I have prominent deficits in sense of responsibility and time management. | | | | | |
| I always feel bored and depressive for many activities. | | | | | |
| I put off taking actions to do tasks despite I have already made decisions to do so. | | | | | |
| I am of quite remorse for my procrastination behaviors. | | | | | |
| I cannot complete my social tasks (e.g., personal works, homework or learning) well because I am hard to follow instructions or guidelines of these tasks. | | | | | |
| I have a poor sleep quality. | | | | | |

**Table S1. Pathological Procrastination Diagnostic Questionnaire.**

**2. ACE Model Comparison and Selection**

To estimate the heritability of psychopathological procrastination (PPS), an Additive Genetic - Common Environmental - Non-shared Environmental (ACE) structural equation model was implemented within a classical twin study framework. This model decomposes the total phenotypic variance into three latent components: additive genetic variance (A), shared environmental variance (C), and non-shared environmental variance (E). The additive genetic component represents heritable influences, the shared environmental component accounts for environmental factors common to both twins, such as socioeconomic background and family upbringing, while the non-shared environmental componen captures individual-specific environmental influences and measurement error.

To determine the most parsimonious model, nested submodels were compared against the full ACE model using maximum likelihood estimation (MLE) via the OpenMx package (v2.18.1). The model fit was evaluated based on multiple statistical criteria, including the -2 Log Likelihood (-2LL), Akaike Information Criterion (AIC), and Bayesian Information Criterion (BIC). The -2LL statistic was used as a primary likelihood measure, with lower values indicating better model fit. The AIC was applied to assess the trade-off between model fit and complexity, with lower values suggesting a more optimal balance. The BIC, which incorporates sample size adjustments, served as an additional index to prevent overfitting and improve generalizability. To statistically assess model fit differences, likelihood ratio tests were performed to compare the full ACE model with reduced versions that systematically excluded either the additive genetic component (AE model), the shared environmental component (CE model), or both (E model). These comparisons allowed for an evaluation of whether excluding specific variance components resulted in a significant loss of explanatory power.

The model comparison results indicated that the AE model provided the best fit, as evidenced by its lowest AIC and BIC values **(Supplemental Table S2)**. The exclusion of the shared environmental component did not significantly degrade the model fit ($p > 0.05$), suggesting a minimal contribution of shared environmental influences to PPS. In contrast, the removal of the additive genetic component resulted in a significantly poorer fit ($p < 0.001$), reinforcing the role of genetic factors in shaping PPS phenotypic variation. Given these findings, the AE model was selected as the optimal model, yielding a heritability estimate of $h^2$=0.38 (95% CI: [0.03 – 0.66]).

| | **ACE model** | **AE model** | **CE model** | **E model** |
|---|---|---|---|---|
| Akaike Information Criterion, AIC | 128.38 | **126.38** | 131.46 | 132.03 |
| Bayesian Information Criterion, BIC | -223.19 | **-227.53** | -222.44 | -224.23 |
| -2 Log Likelihood , -2LL | 428.38 | **428.37** | 433.47 | 436.03 |

**Table S2. Model fits of ACE model and nested submodels.** The best-fitting model is in bold.

**3. Preprocessing of Gene Transcriptomic Datasets**

To examine the neurogenetic basis of psychopathological procrastination (PPS), we utilized the Allen Human Brain Atlas (AHBA) dataset, a spatially resolved whole-brain transcriptional atlas developed by the Allen Institute for Brain Science (AIBS) (http://human.brain-map.org, RRID: SCR_007416). This dataset provides gene expression profiles from six adult postmortem donors (*mean age = 42.50 ± 13.38 years*, *one female*), including two whole-brain samples and four left-hemisphere-only samples. All donors were confirmed to be free of neuropsychiatric and neurological conditions, including traumatic brain injury, epilepsy, and substance use disorders. Postmortem brains were processed within 30 hours after death, with each brain dissected into approximately 500 anatomically localized tissue samples per hemisphere. Gene expression levels were quantified using high-throughput DNA microarray sequencing, and the resulting data were subsequently mapped onto 3D MRI coordinates in the standard MNI space using high-resolution T1-weighted neuroimaging data. The complete dataset comprised 3,072 samples, with qualified expression data from 58,692 probes.

To ensure rigorous preprocessing and standardization, we followed a validated pipeline (https://github.com/BMHLab/AHBAprocessing), which involved six key processing steps: First, gene re-annotation was performed to update probe-to-gene mappings and address outdated annotations. Using the re-annotator toolbox, probe sequences (*n = 45,821*) were aligned to the hg38 sequencing database, resulting in the identification of 20,232 uniquely annotated genes. Second, an intensity-based filter (IBF) was applied to remove low-expression background noise, retaining only genes with expression levels exceeding background thresholds (*final gene set: n = 10,190*). Third, to resolve instances where multiple probes mapped to the same gene, a probe selection procedure was conducted, wherein the probe with the highest RNA-seq correlation was chosen to represent each gene. Fourth, for connectome-transcriptional alignment, tissue samples were registered to the Desikan-Killiany (D-K) cortical and subcortical parcellation atlas, which consists of 68 anatomically defined brain regions (34 per hemisphere). To ensure precise spatial correspondence, samples were retained only if their centroid Euclidean distance to the assigned D-K region was ≤ 2 mm. This quality control step resulted in the inclusion of 1,184 high-quality samples, covering all 68 brain regions and 10,027 genes for downstream analyses. Fifth, to correct for inter-sample and inter-donor variability, a scaled robust sigmoid normalization was applied. Intra-sample normalization was first performed across probes within each sample, followed by inter-donor normalization to ensure expression comparability across brains.Finally, gene selection and regional aggregation were conducted, in which gene expression levels were averaged across all six donors within each of the 68 regions of the D-K atlas. This produced a final region-gene expression matrix of 68 × 10,027, which was subsequently used for brain-transcriptomic association analysis.

**4. Fuzzy Clustering Model**

Fzzy (soft) clustering analysis was employed to identify co-expression modules of genes spanning all the 25 neurodevelopmental periods for these 273 PPS-specific genes. Unlike traditional clustering methods that assign genes to a single discrete category, fuzzy clustering allows for soft partitioning, acknowledging the reality that many genes participate in multiple biological processes simultaneously. This flexibility is particularly advantageous for transcriptomic analyses, where gene

expression patterns often exhibit overlapping regulatory influences over the time (Wang et al., 2024).
The clustering was performed using the "Mfuzz" package (*v2.56.0*) in R (https://bioconductor.org/packages/release/bioc/html/Mfuzz.html), a well-established tool for analyzing gene expression dynamics. To do so, the gene expressions were averaged across all the regions firstly (Wang et al., 2024). Before clustering, Z-score normalization was applied to the gene-age expression matrix (273 × 25) to standardize expression levels. This step ensured comparability across genes and mitigated the impact of absolute expression differences on clustering outcomes. Following normalization, fuzzy c-means clustering was implemented to group genes based on similarity in expression profiles. The algorithm operates by iteratively minimizing within-cluster variance while maximizing separation between clusters. Instead of discrete assignments, it calculates a membership score ranging from 0 to 1, which quantifies the degree to which a gene belongs to each cluster. A higher membership score indicates a stronger association between a gene and a given cluster, while lower scores suggest a more diffuse expression pattern.
Determining the optimal number of clusters ($c$) was critical to ensuring biologically meaningful partitioning. Rather than relying on arbitrary selection, a systematic evaluation was conducted by varying $c$ from 2 to 10, assessing within-cluster variances. By doing so, we plotted the association between $c$ and within-cluster variance, which revealed a characteristic elbow point = 3, indicating the threshold beyond which increasing $c$ yielded diminishing variance reduction **(Supplemental Figure S1)**. However, considering this case that the third cluster includes three genes only, we determined the elbow point = 2 as the optimal balance between resolution and biological interpretability **(Supplemental Figure S2)**.

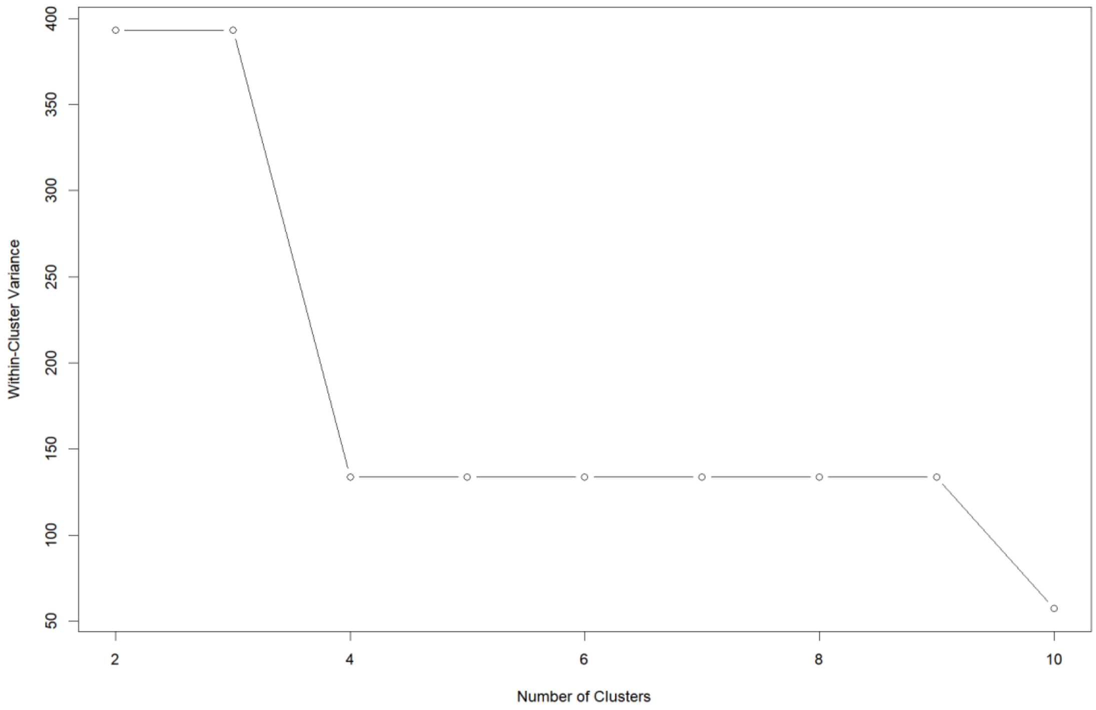


**Figure S1. Clustering solution selection from with-cluster variance across the**

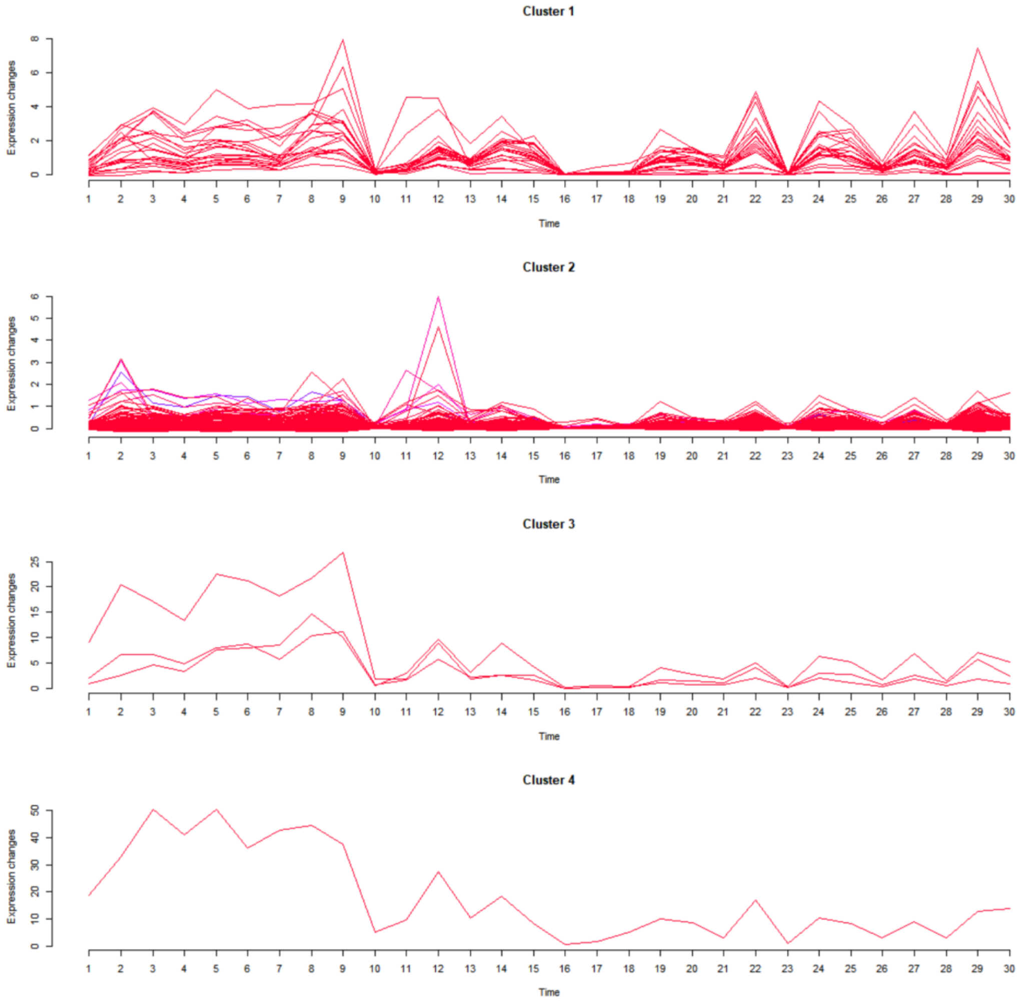

**number of clusters**.
**Figure S2. Gene co-expression trajectories across all the neurodevelopmental ages**.

### 5. GAMBA Enrichment Analysis

The Gene Annotation by Macroscale Brain Imaging Association (GAMBA) platform was utilized to decode the association between identified gene set (i.e., PLS2) and PPS-related neuroimaging phenotypes, employing a spatially resolved regression-based framework (http://dutchconnectomelab.nl/GAMBA/). The detailed methodology for constructing the gene list and its integration with neuroimaging features is presented in the main text. Here, we provide the original statistical formulations and computational procedures applied in GAMBA for assessing gene-neuroimaging associations.

The relationship between cortical gene expression profiles and neuroimaging-derived phenotypes was quantified using a linear regression model, formulated as follows:

$$y_i = \beta_i \beta 1 X j + \varepsilon$$

where $Y_i$ represents the normalized expression profile of gene $i$, or the average expression profile of a gene set $i$; $X^j$ denotes the normalized cortical profile of neuroimaging phenotype $j$; and *cov* accounts for normalized covariates included in the model. Normalization was performed by subtracting the mean from each value and dividing by the standard deviation, ensuring that all variables were standardized prior to regression analysis. The GAMBA pipeline estimated the standardized regression coefficient $\beta_i$ and the corresponding p-value, reflecting the spatial association between gene expression and the given neuroimaging phenotype.
In addition to assessing direct gene-neuroimaging associations, GAMBA implemented multiple null models to test the spatial specificity and gene specificity of the observed relationships. These included:
**Null-Spin Model –** Preserves the spatial autocorrelation structure of neuroimaging data while randomly rotating cortical gene expression maps.
**Null-Random Model –** Randomly shuffles regional gene expression profiles across brain regions, breaking any spatial organization.
**Null-Coexpression Model –** Maintains spatial patterns but replaces individual gene profiles with randomly selected co-expressed genes, ensuring that effects are not driven by broad co-expression patterns.
**Null-Brain Model –** Tests whether observed gene-phenotype associations exceed those expected under random neurobiological constraints.
For each gene-neuroimaging association, GAMBA performed a z-test to determine whether the observed effect size ($\beta_i$) exceeded the distribution of effect sizes across null models:

$$z = \frac{\beta_1 - u}{\sigma}$$

where $\mu$ and $\sigma$ represent the mean and standard deviation of $\beta_i$ across null models, respectively.

**6. Gene Set Enrichment Analysis**

To elucidate the biological functions, molecular pathways, and regulatory networks underlying psychopathological procrastination (PPS)-specific transcriptomic signatures, we conducted gene set enrichment analysis (GSEA) using the Gene Set Enrichment Analysis (GSEA) software (v4.3.3) (https://www.gsea-msigdb.org/gsea/). This computational method determines whether a predefined set of genes exhibits significant, coordinated differences in expression within a given biological context, providing insight into the molecular pathways governing PPS-related neurogenetic mechanisms.

For the enrichment analysis, the full transcriptomic profile (10,027 genes) was ranked based on Z-scored expression weights derived from the partial least squares (PLS) regression analysis, which linked PPS-specific neuroanatomical deviations to gene expression levels. The ranking metric was calculated as follows:

$$R_{(g)} = -\log_{10}(p) \times sign(\beta)$$

where $R(_{g})$ represents the gene ranking score, $p$ is the unadjusted p-value from PLS regression, and $\beta$ is the gene-wise regression coefficient. This approach ensures that genes positively correlated with PPS-specific neuroanatomical deviations receive high positive ranks, whereas genes negatively correlated are assigned lower ranks, preserving directional biological information in the analysis.

GSEA was performed using multiple curated gene set collections from the Molecular Signatures Database (MSigDB, v7.5.1), capturing diverse biological categories relevant to neuropsychiatric disorders, neurogenetic dysregulation, and neuroinflammation. To assess statistical significance, enrichment scores were calculated for each gene set based on a weighted Kolmogorov-Smirnov (K-S) statistic, which quantifies the degree to which gene sets exhibit concordant expression shifts in the ranked transcriptome. The normalized enrichment score (NES) was computed to account for gene set size differences, ensuring comparability across gene sets. Statistical significance was determined using empirical p-values ($p < 0.05$) derived from 1,000 random permutations, controlling for false positives arising from stochastic expression fluctuations.Permutation test was conducted as following parameters: Gene set-based permutation ($N = 1{,}000$); Minimum gene set size: 15; Maximum gene set size: 500; Metric for ranking genes: Signal-to-noise ratio; Normalization mode: Meandiv. To correct for multiple comparisons, the False Discovery Rate (FDR) correction was applied, with an adjusted threshold of $q < 0.05$ to identify statistically robust gene set enrichments. Results meeting this threshold were considered biologically significant, while non-significant enrichments were excluded from further interpretation.

**Supplemental Results**

**1. AE Model to Estimate Heritability**

To quantify the heritability of psychopathological procrastination (PPS), we implemented Additive Genetic – Unique Environmental (AE) modeling, a parsimonious structural equation model that accounts for the proportion of phenotypic variance attributable to additive genetic factors (A) and non-shared environmental influences (E). This model was selected based on model comparison results, where the full Additive Genetic – Common Environmental – Unique Environmental (ACE) model was compared to reduced models (AE, CE, and E) using likelihood ratio tests (LRTs) and information criteria-based selection. As reported in Supplemental Methods, the AE model demonstrated the best fit, with the lowest Akaike Information Criterion (AIC) and Bayesian Information Criterion (BIC) values, confirming the negligible contribution of shared environmental factors.

Upon this AE model, we estimate the variants explained by A and E, respectively, which has been tabulated into the **Table S3**.

| | $a^2(h^2)$ | $c^2$ | $e^2$ | *p*_ACE_AE | *p*_ACE_CE | *p*_ACE_E | *p*_AE_E |
|---|---|---|---|---|---|---|---|
| ACE | 0.47 | 5.9E-13 | 0.52 | 0.05 | 0.08 | 0.01 | 0.009 |
| CE | 0 | 0.18 | 0.81 | | | | |
| AE | 0.4721 | 0 | 0.52 | 1 | 0.02 | 0.02 | 0.005 |

**Table S3. Estimates of ACE model and nested submodels.** *P* values indicate statistical significance of likelihood ratio tests (LRTs).

For the parametric modeling, we firstly conducted linear correlation analysis for within-pair phenotypic association on MZ and DZ, respectively. For the zero-rank correlation analysis, phenotypic associations in the MZ ($r = 0.53, p < .001$) are larger than in DZ ($r = -0.21, p = .25$). Also, for the intra-class correlation (ICC) model to PPS scores, ICC in the MZ were found statistically larger than in the DZ as well ($F = 5.62, p < .0001$). Therefore, phenotypic associations were not varied from statistical models in modeling the differences between MZ and DZ, indicating statistical robustness.

**2. Data Distribution of Brain Morphological Deviations**

To evaluate the distributional properties of cortical thickness (CT), surface area (SA), and subcortical volume (SV), we employed Gaussian kernel density estimation (KDE) using custom MATLAB scripts (R2023b). This non-parametric density estimation technique provided a smoothed representation of data distribution without assuming an underlying parametric model, ensuring a more precise characterization of the morphological variability across brain regions. For cortical thickness and surface area, KDE was applied separately to each of the 68 cortical regions defined by the Desikan-Killiany (D-K) atlas. For subcortical volume, KDE analysis was performed for 14 subcortical structures identified using the Aseg atlas. This ensured that all structural MRI-derived features were examined within their respective anatomical parcellation frameworks. To optimize the estimation process, Gaussian kernels with adaptive bandwidth selection were used to balance resolution and smoothness in the probability density function. The Silverman's rule of thumb was applied to determine bandwidth parameters, ensuring that the density curves retained fine-grained distributional details while avoiding over-smoothing or excessive fragmentation of the data. By doing so, we found no significant deviations from the Gaussian normality to morphological deviations in this cohort on CT **(Figure S3)**, SA **(Figure S4)** and SV **(Figure S5)**. For interpretability, the full names of brain regions numbered in these figures were tabulated into the **Table S4-5**.

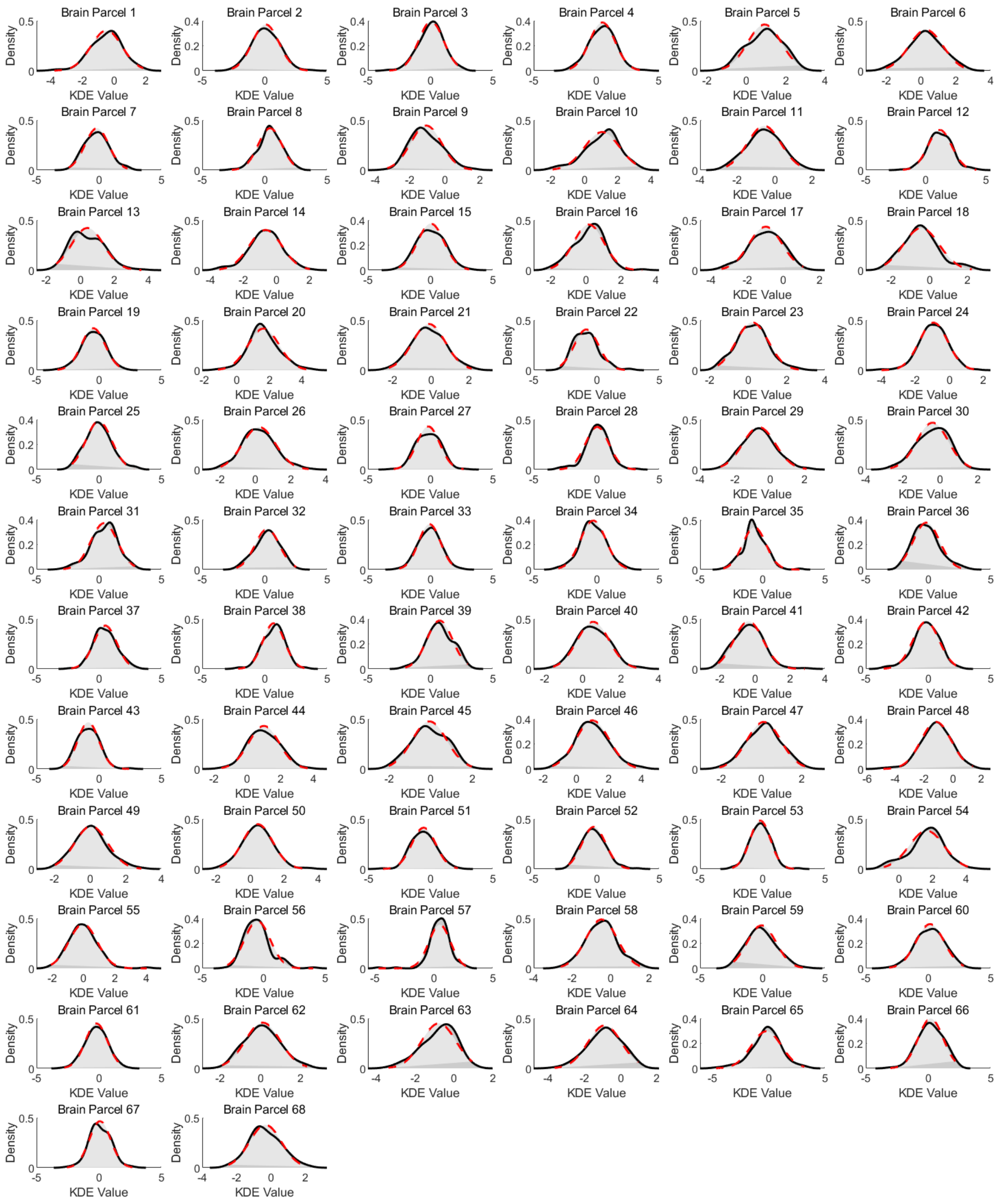

**Figure S3. Kernel Density Estimation (KDE) for cortical thickness deviations in this cohort and standard normal distribution curves across 68 brain cortical regions.** The figure displays the probability density functions (PDFs) for each of the 68 brain regions, estimated using Kernel Density Estimation (KDE). The KDE curves (solid black lines) represent the empirical distribution of the data, while the standard normal distribution curves (red dashed lines) are overlaid for comparison. The area under the KDE curves is shaded in light gray, and the area under the standard normal distribution curves is shaded in lighter gray. Each subplot corresponds to a specific brain region, illustrating the similarity or deviation of the empirical distribution from a normal distribution. The x-axis represents the values, and the y-axis represents the probability density.

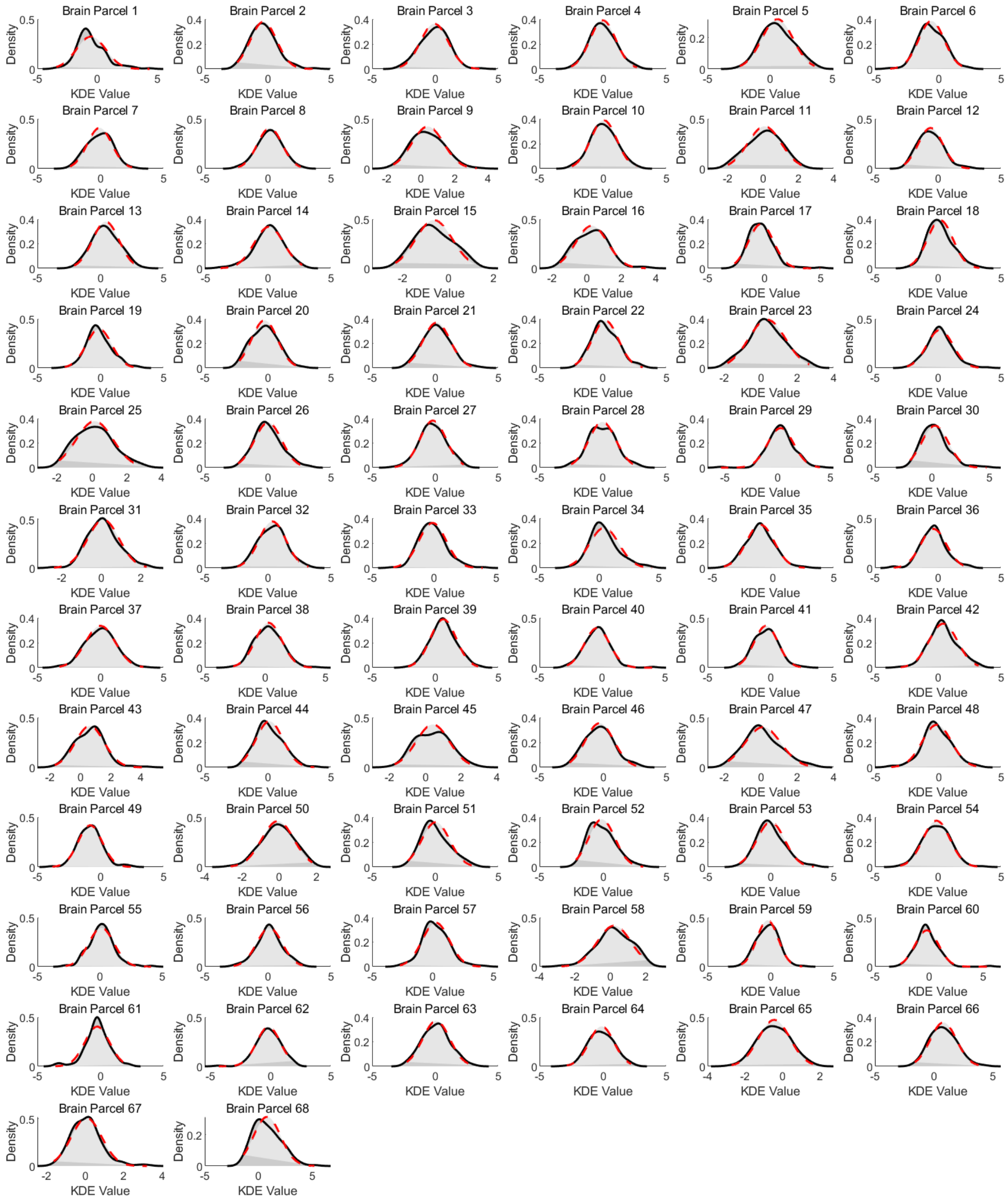

**Figure S4. Kernel Density Estimation (KDE) for surface area deviations in this cohort and standard normal distribution curves across 68 brain cortical regions.**

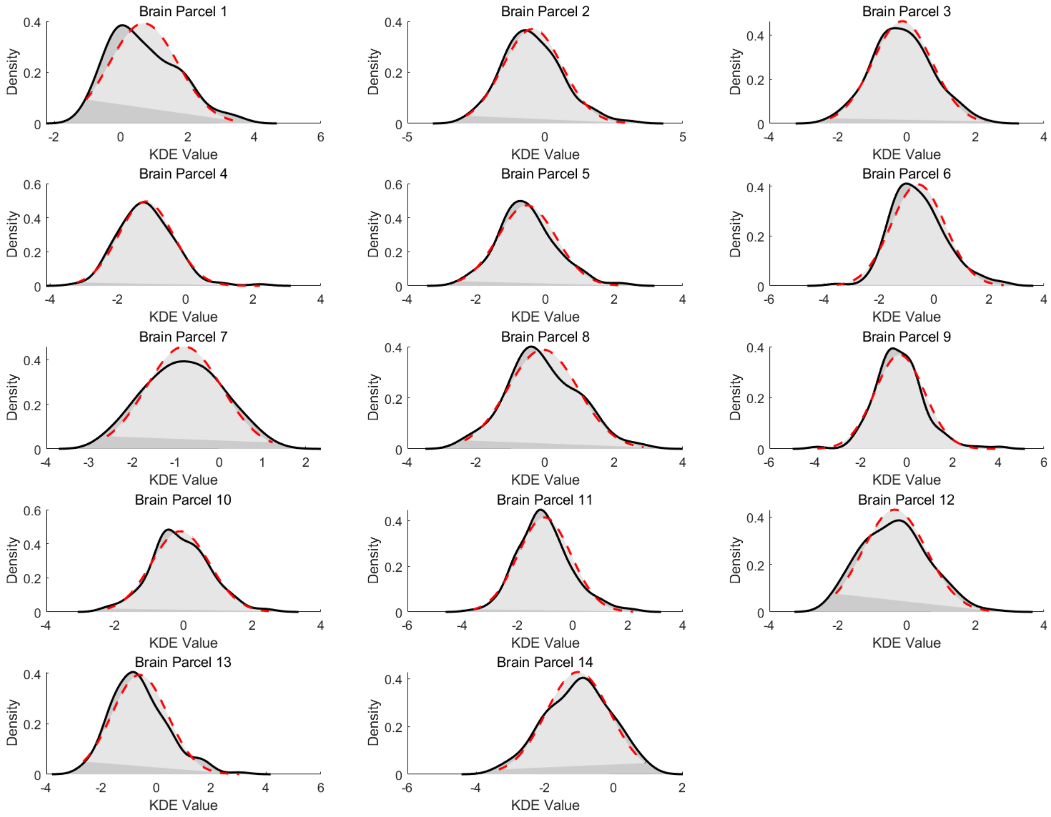


**Figure S5. Kernel Density Estimation (KDE) for subcortical areas deviations in this cohort and standard normal distribution curves across 14 brain subcortical regions.**

| Number | Full names |
|---|---|
| Brain parcel 1 | Left banks of the superior temporal sulcus |
| Brain parcel 2 | Left caudal anterior cingulate |
| Brain parcel 3 | Left caudal middle frontal |
| Brain parcel 4 | Left cuneus |
| Brain parcel 5 | Left entorhinal cortex |
| Brain parcel 6 | Left fusiform gyrus |
| Brain parcel 7 | Left inferior parietal lobule |
| Brain parcel 8 | Left inferior temporal gyrus |
| Brain parcel 9 | Left isthmus of the cingulate cortex |
| Brain parcel 10 | Left lateral occipital cortex |
| Brain parcel 11 | Left lateral orbitofrontal cortex |
| Brain parcel 12 | Left lingual gyrus |
| Brain parcel 13 | Left medial orbitofrontal cortex |
| Brain parcel 14 | Left middle temporal gyrus |
| Brain parcel 15 | Left parahippocampal gyrus |
| Brain parcel 16 | Left paracentral lobule |

| | |
|---|---|
| Brain parcel 17 | Left pars opercularis |
| Brain parcel 18 | Left pars orbitalis |
| Brain parcel 19 | Left pars triangularis |
| Brain parcel 20 | Left pericalcarine cortex |
| Brain parcel 21 | Left postcentral gyrus |
| Brain parcel 22 | Left posterior cingulate cortex |
| Brain parcel 23 | Left precentral gyrus |
| Brain parcel 24 | Left precuneus |
| Brain parcel 25 | Left rostral anterior cingulate cortex |
| Brain parcel 26 | Left rostral middle frontal cortex |
| Brain parcel 27 | Left superior frontal gyrus |
| Brain parcel 28 | Left superior parietal lobule |
| Brain parcel 29 | Left superior temporal gyrus |
| Brain parcel 30 | Left supramarginal gyrus |
| Brain parcel 31 | Left frontal pole |
| Brain parcel 32 | Left temporal pole |
| Brain parcel 33 | Left transverse temporal gyrus |
| Brain parcel 34 | Left insular cortex |
| Brain parcel 35 | Right banks of the superior temporal sulcus |
| Brain parcel 36 | Right caudal anterior cingulate |
| Brain parcel 37 | Right caudal middle frontal |
| Brain parcel 38 | Right cuneus |
| Brain parcel 39 | Right entorhinal cortex |
| Brain parcel 40 | Right fusiform gyrus |
| Brain parcel 41 | Right inferior parietal lobule |
| Brain parcel 42 | Right inferior temporal gyrus |
| Brain parcel 43 | Right isthmus of the cingulate cortex |
| Brain parcel 44 | Right lateral occipital cortex |
| Brain parcel 45 | Right lateral orbitofrontal cortex |
| Brain parcel 46 | Right lingual gyrus |
| Brain parcel 47 | Right medial orbitofrontal cortex |
| Brain parcel 48 | Right middle temporal gyrus |
| Brain parcel 49 | Right parahippocampal gyrus |
| Brain parcel 50 | Right paracentral lobule |
| Brain parcel 51 | Right pars opercularis |
| Brain parcel 52 | Right pars orbitalis |
| Brain parcel 53 | Right pars triangularis |
| Brain parcel 54 | Right pericalcarine cortex |
| Brain parcel 55 | Right postcentral gyrus |
| Brain parcel 56 | Right posterior cingulate cortex |
| Brain parcel 57 | Right precentral gyrus |
| Brain parcel 58 | Right precuneus |
| Brain parcel 59 | Right rostral anterior cingulate cortex |
| Brain parcel 60 | Right rostral middle frontal cortex |
| Brain parcel 61 | Right superior frontal gyrus |
| Brain parcel 62 | Right superior parietal lobule |
| Brain parcel 63 | Right superior temporal gyrus |
| Brain parcel 64 | Right supramarginal gyrus |
| Brain parcel 65 | Right frontal pole |
| Brain parcel 66 | Right temporal pole |

| | |
|---|---|
| Brain parcel 67 | Right transverse temporal gyrus |
| Brain parcel 68 | Right insular cortex |

**Table S4. Full names to brain parcel numbers.**

| Number | Full names |
|---|---|
| Brain parcel 1 | Left nucleus accumbens |
| Brain parcel 2 | Left amygdala |
| Brain parcel 3 | Left caudate nucleus |
| Brain parcel 4 | Left hippocampus |
| Brain parcel 5 | Left globus pallidus |
| Brain parcel 6 | Left putamen |
| Brain parcel 7 | Left thalamus |
| Brain parcel 8 | Right nucleus accumbens |
| Brain parcel 9 | Right amygdala |
| Brain parcel 10 | Right caudate nucleus |
| Brain parcel 11 | Right hippocampus |
| Brain parcel 12 | Right globus pallidus |
| Brain parcel 13 | Right putamen |
| Brain parcel 14 | Right thalamus |

**Table S5. Full names to brain parcel numbers.**

Furthermore, we conducted one-sample t-tests comparing the mean deviation scores against a theoretical mean of zero, for assess whether regional morphological deviations in cortical thickness (CT), surface area (SA), and subcortical volume (SV) significantly differed from baseline expectations. The analysis was performed separately for each cortical region (n = 68, Desikan-Killiany atlas) and each subcortical structure (n = 14, Aseg atlas) to determine whether observed deviations significantly diverged from expected neurodevelopmental patterns. All t-tests were implemented using MATLAB's statistical toolbox (R2023b, ttest function), ensuring numerical precision and consistency across regions. To correct for multiple comparisons across all brain regions, False Discovery Rate (FDR) correction was applied at $q<0.05$, following the Benjamini-Hochberg (BH) procedure. In this cohort, for the CT morphmetry, 18 cortical regions significantly overdeveloped, while 25 ones in underdeveloped **(Table S6)**. For the SA morphmetry, 18 cortical regions significantly overdeveloped, while16 ones in underdeveloped in this cohort **(Table S7)**. Finally, for the subcortical volumes, 11 subcortical regions significantly underdeveloped in this cohort **(Table S8)**. Moreover, for the individual-level analysis, we predefined deviation Z scores with ≥ (≤) 1.96 as threshold to determine individual brain morphological overdevelopment (underdevelopment). By doing so, we presented the proportion of identified participants showing overdevelopment (underdevelopment) to total number of participants ($N$ = 140) in this cohort. These statistics were tabulated into **Table S6-8**.

| Brain Parcel | T value | P value | BH q value | Proportion |
|---|---|---|---|---|
| L_pericalcarine | 20.485 | 0 | 0 | 0.0000 |
| R_pericalcarine | 19.054 | 0 | 0 | 0.0000 |
| L_lingual | 14.187 | 0 | 0 | 0.0000 |
| R_lateraloccipital | 13.034 | 0 | 0 | 0.0000 |
| L_entorhinal | 12.789 | 0 | 0 | 0.0000 |
| L_lateraloccipital | 12.267 | 0 | 0 | 0.0000 |
| R_lingual | 11.684 | 0 | 0 | 0.0000 |
| R_cuneus | 9.923 | 0 | 0 | 0.7143 |
| R_entorhinal | 8.654 | 0 | 0 | 0.7143 |
| R_fusiform | 7.872 | 0 | 0 | 0.0000 |
| R_precentral | 6.792 | 0 | 0 | 1.4286 |
| R_caudalmiddlefrontal | 6.546 | 0 | 0 | 0.7143 |
| R_paracentral | 6.022 | 0 | 0 | 0.0000 |
| L_cuneus | 5.908 | 0 | 0 | 0.7143 |
| L_medialorbitofrontal | 5.497 | 0 | 0 | 0.0000 |
| L_inferiortemporal | 5.474 | 0 | 0 | 1.4286 |
| L_frontalpole | 4.663 | 0 | 0 | 2.8571 |
| L_fusiform | 3.582 | 0 | 0 | 1.4286 |
| L_precentral | 3.077 | 0.003 | 0.054 | 0.0000 |
| L_rostralmiddlefrontal | 2.676 | 0.008 | 0.144 | 0.7143 |
| L_temporalpole | 2.604 | 0.01 | 0.15 | 1.4286 |
| R_temporalpole | 2.427 | 0.016 | 0.224 | 2.1429 |
| L_paracentral | 2.37 | 0.019 | 0.2327 | 1.4286 |
| R_medialorbitofrontal | 1.782 | 0.077 | 0.514 | 0.7143 |
| R_superiorparietal | 1.729 | 0.086 | 0.515 | 0.0000 |
| R_rostralmiddlefrontal | 1.639 | 0.104 | 0.52 | 4.2857 |
| R_transversetemporal | 1.311 | 0.192 | 0.576 | 0.7143 |
| L_caudalanteriorcingulate | 0.942 | 0.348 | 0.696 | 2.1429 |
| L_superiorparietal | 0.853 | 0.395 | 0.79 | 4.2857 |
| L_caudalmiddlefrontal | 0.819 | 0.414 | 0.828 | 2.1429 |
| R_parahippocampal | 0.768 | 0.444 | 0.888 | 0.7143 |
| L_parahippocampal | 0.486 | 0.628 | 1 | 2.1429 |
| L_rostralanteriorcingulate | -0.07 | 0.944 | 0.94 | 2.1429 |
| R_postcentral | -0.323 | 0.747 | 0.77 | 1.4286 |
| R_lateralorbitofrontal | -0.649 | 0.518 | 0.55 | 0.7143 |
| R_rostralanteriorcingulate | -0.685 | 0.494 | 0.54 | 3.5714 |
| L_transversetemporal | -0.823 | 0.412 | 0.47 | 1.4286 |
| L_postcentral | -1.348 | 0.18 | 0.21 | 1.4286 |
| R_caudalanteriorcingulate | -1.562 | 0.121 | 0.15 | 2.8571 |
| R_frontalpole | -1.669 | 0.097 | 0.12 | 7.8571 |
| R_inferiortemporal | -1.842 | 0.068 | 0.09 | 6.4286 |
| L_inferiorparietal | -1.942 | 0.054 | 0.07 | 0.7143 |

| | | | | |
|---|---|---|---|---|
| L_superiorfrontal | -2.087 | 0.039 | 0.05 | 1.4286 |
| R_parstriangularis | -2.482 | 0.014 | 0.02 | 2.1429 |
| R_parsorbitalis | -2.498 | 0.014 | 0.02 | 0.7143 |
| R_superiorfrontal | -2.578 | 0.011 | 0.02 | 2.1429 |
| L_insula | -2.936 | 0.004 | 0.01 | 6.4286 |
| R_insula | -3.908 | 0 | 0.00 | 2.8571 |
| R_inferiorparietal | -4.995 | 0 | 0.00 | 1.4286 |
| L_parsorbitalis | -5.268 | 0 | 0.00 | 3.5714 |
| L_parstriangularis | -5.557 | 0 | 0.00 | 7.1429 |
| R_posteriorcingulate | -5.581 | 0 | 0.00 | 5.0000 |
| L_supramarginal | -5.634 | 0 | 0.00 | 4.2857 |
| L_bankssts | -5.7 | 0 | 0.00 | 5.7143 |
| R_precuneus | -6.62 | 0 | 0.00 | 2.8571 |
| R_parsopercularis | -6.632 | 0 | 0.00 | 4.2857 |
| L_lateralorbitofrontal | -7.403 | 0 | 0.00 | 7.8571 |
| L_middletemporal | -7.595 | 0 | 0.00 | 7.1429 |
| L_superiortemporal | -7.664 | 0 | 0.00 | 7.1429 |
| R_bankssts | -8.261 | 0 | 0.00 | 8.5714 |
| L_posteriorcingulate | -9.584 | 0 | 0.00 | 8.5714 |
| R_superiortemporal | -10.297 | 0 | 0.00 | 9.2857 |
| R_supramarginal | -11.024 | 0 | 0.00 | 11.4286 |
| R_isthmuscingulate | -11.673 | 0 | 0.00 | 7.8571 |
| L_parsopercularis | -12.445 | 0 | 0.00 | 11.4286 |
| R_middletemporal | -12.842 | 0 | 0.00 | 20.7143 |
| L_precuneus | -13.089 | 0 | 0.00 | 10.0000 |
| L_isthmuscingulate | -14.243 | 0 | 0 | 15.7143 |

**Table S6. Group-averaged deviations on CT for each region in this cohort.** T values indicate statistics from the one-sample t test, and BH q values suggest corrected p values from original ones using Benjamini-Hochberg method.

| Brain Parcel | T value | P value | BH q value | Proportion |
|---|---|---|---|---|
| R_supramarginal | 8.722 | 0 | 0.0000 | 0.0000 |
| R_medialorbitofrontal | 7.729 | 0 | 0.0000 | 0.0000 |
| R_insula | 7.667 | 0 | 0.0000 | 0.0000 |
| R_isthmuscingulate | 7.488 | 0 | 0.0000 | 0.7143 |
| L_inferiortemporal | 5.641 | 0 | 0.0000 | 1.4286 |
| R_paracentral | 5.42 | 0 | 0.0000 | 0.0000 |
| L_parsopercularis | 5.368 | 0 | 0.0000 | 0.0000 |
| L_precentral | 4.806 | 0 | 0.0000 | 0.7143 |
| R_lingual | 4.472 | 0 | 0.0000 | 1.4286 |
| L_frontalpole | 4.291 | 0 | 0.0000 | 1.4286 |
| L_precuneus | 4.167 | 0 | 0.0000 | 0.7143 |

| | | | | |
|---|---|---|---|---|
| L_cuneus | 4.11 | 0 | 0.0000 | 0.0000 |
| L_superiorparietal | 3.763 | 0 | 0.0000 | 0.0000 |
| R_fusiform | 3.298 | 0.001 | 0.0028 | 1.4286 |
| L_paracentral | 2.996 | 0.003 | 0.0078 | 0.7143 |
| R_caudalmiddlefrontal | 2.911 | 0.004 | 0.0098 | 1.4286 |
| R_bankssts | 2.861 | 0.005 | 0.0115 | 2.1429 |
| L_pericalcarine | 2.786 | 0.006 | 0.0130 | 0.0000 |
| L_caudalanteriorcingulate | 2.24 | 0.027 | 0.0554 | 0.7143 |
| L_supramarginal | 2.119 | 0.036 | 0.0702 | 1.4286 |
| L_middletemporal | 1.999 | 0.048 | 0.0891 | 2.1429 |
| L_entorhinal | 1.859 | 0.065 | 0.1152 | 1.4286 |
| L_isthmuscingulate | 1.808 | 0.073 | 0.1238 | 0.0000 |
| L_posteriorcingulate | 1.719 | 0.088 | 0.1430 | 0.7143 |
| R_temporalpole | 1.592 | 0.114 | 0.1725 | 3.5714 |
| R_frontalpole | 1.586 | 0.115 | 0.1725 | 1.4286 |
| R_parahippocampal | 1.463 | 0.146 | 0.2075 | 0.7143 |
| R_pericalcarine | 1.453 | 0.149 | 0.2075 | 0.0000 |
| L_insula | 1.356 | 0.177 | 0.2380 | 1.4286 |
| L_parstriangularis | 1.322 | 0.188 | 0.2415 | 0.7143 |
| R_transversetemporal | 1.312 | 0.192 | 0.2415 | 0.0000 |
| R_parstriangularis | 0.991 | 0.323 | 0.3937 | 2.8571 |
| L_rostralmiddlefrontal | 0.865 | 0.388 | 0.4585 | 5.7143 |
| L_parsorbitalis | 0.845 | 0.4 | 0.4588 | 2.8571 |
| R_precuneus | 0.471 | 0.639 | 0.7120 | 0.7143 |
| L_lingual | 0.31 | 0.757 | 0.8201 | 1.4286 |
| R_superiorfrontal | 0.191 | 0.849 | 0.8949 | 1.4286 |
| L_inferiorparietal | 0.066 | 0.948 | 0.9550 | 2.1429 |
| R_entorhinal | 0.057 | 0.955 | 0.9550 | 3.5714 |
| L_parahippocampal | -0.356 | 0.723 | 0.7230 | 5.0000 |
| L_fusiform | -0.623 | 0.534 | 0.5531 | 5.0000 |
| L_lateralorbitofrontal | -0.737 | 0.462 | 0.4962 | 2.1429 |
| R_precentral | -0.913 | 0.363 | 0.4049 | 1.4286 |
| R_rostralanteriorcingulate | -0.917 | 0.361 | 0.4049 | 1.4286 |
| R_rostralmiddlefrontal | -1.155 | 0.25 | 0.3021 | 2.1429 |
| L_rostralanteriorcingulate | -1.185 | 0.238 | 0.3001 | 2.1429 |
| R_superiorparietal | -1.194 | 0.235 | 0.3001 | 3.5714 |
| L_temporalpole | -1.226 | 0.222 | 0.3001 | 2.1429 |
| R_postcentral | -1.233 | 0.22 | 0.3001 | 5.7143 |
| L_medialorbitofrontal | -1.862 | 0.065 | 0.0992 | 4.2857 |
| R_parsorbitalis | -1.965 | 0.051 | 0.0822 | 2.8571 |
| R_inferiortemporal | -2.061 | 0.041 | 0.0699 | 2.8571 |
| R_parsopercularis | -2.695 | 0.008 | 0.0145 | 7.1429 |
| L_transversetemporal | -2.901 | 0.004 | 0.0077 | 3.5714 |
| L_superiortemporal | -2.914 | 0.004 | 0.0077 | 3.5714 |

| | | | | |
|---|---|---|---|---|
| R_middletemporal | -3.068 | 0.003 | 0.0067 | 4.2857 |
| R_inferiorparietal | -3.229 | 0.002 | 0.0048 | 1.4286 |
| L_caudalmiddlefrontal | -3.791 | 0 | 0.0000 | 5.0000 |
| R_cuneus | -4.052 | 0 | 0.0000 | 2.8571 |
| R_lateralorbitofrontal | -4.421 | 0 | 0.0000 | 3.5714 |
| R_lateraloccipital | -4.662 | 0 | 0.0000 | 5.0000 |
| L_bankssts | -4.855 | 0 | 0.0000 | 6.4286 |
| R_superiortemporal | -5.94 | 0 | 0.0000 | 2.8571 |
| L_lateraloccipital | -6.597 | 0 | 0.0000 | 7.1429 |
| L_postcentral | -7.079 | 0 | 0.0000 | 7.1429 |
| L_superiorfrontal | -8.96 | 0 | 0.0000 | 3.5714 |
| R_posteriorcingulate | -10.058 | 0 | 0.0000 | 9.2857 |
| R_caudalanteriorcingulate | -10.577 | 0 | 0.0000 | 22.1429 |

**Table S7. Group-averaged deviations on SA for each region in this cohort.** T values indicate statistics from the one-sample t test, and BH q values suggest corrected p values from original ones using Benjamini-Hochberg method.

| Brain Parcel | T value | P value | BH q value | Proportion |
|---|---|---|---|---|
| Lthal | -11.037 | 0 | 0.0000 | 9.2857 |
| Rthal | -13.083 | 0 | 0.0000 | 17.8571 |
| Lhippo | -17.194 | 0 | 0.0000 | 15.7143 |
| Rput | -6.97 | 0 | 0.0000 | 5.0000 |
| Rhippo | -12.559 | 0 | 0.0000 | 18.5714 |
| Lpal | -7.388 | 0 | 0.0000 | 3.5714 |
| Rpal | -4.435 | 0 | 0.0000 | 2.8571 |
| Lput | -7.08 | 0 | 0.0000 | 5.0000 |
| Lamyg | -4.982 | 0 | 0.0000 | 7.1429 |
| Laccumb | 7.959 | 0 | 0 | 0.0000 |

**Table S8. Group-averaged deviations on SV for each subcortical region in this cohort.** T values indicate statistics from the one-sample t test, and BH q values suggest corrected p values from original ones using Benjamini-Hochberg method.

**3. Predictive Mappings from Brain Deviations to PPS**

Generalized linear mixed-effect models (GLMMs) were implemented using the "lme4" package in R (v4.1.2) to assess the predictive roles of regional neuroanatomical deviations to PPS scores, while accounting for individual/twin variability, age effects, zygosity, and sex differences. Separate models were constructed for each cortical region ($N = 68$, Desikan-Killiany atlas) and subcortical structure ($N = 14$, Aseg atlas), with PPS scores as the dependent variable and regional morphological deviations (cortical thickness, surface area, or subcortical volume) as the primary predictor, one-by-one.

The model specification was defined as:

$$PPS\_i = \beta_0 + \beta_1 \times BrainDeviation_i + Age + Zygosity + Sex + (1 \mid SubjectID) + (1 \mid TwinID) + \varepsilon_i$$

where $\beta_1$ represents the fixed effect of regional morphological deviations on PPS, (1|SubjectID) and (1|TwinID) accounts for individual-level and twin-level variability.

Model fitting was performed using the "lmer" function in the "lme4" package, with Restricted Maximum Likelihood Estimation (REML) to obtain unbiased variance component estimates. To control for multiple comparisons across all brain regions, False Discovery Rate (FDR) correction ($q < 0.05$) was applied using the "p.adjust" function in R, ensuring that significant associations were not driven by chance, for each morphmetric deviation (e.g., CT, SA and SV).

By doing so, we found the statistically significant predictive of brain deviations in CT localized into precuneus for PPS scores ($\beta = 2.9$， $q = .006$). Full results can be found at the **Table S9**.

| Brain Parcel | Beta | Std.Error | T value | P value | BH Q value |
|---|---|---|---|---|---|
| R_precuneus | 2.860377658 | 0.706154792 | 4.050638318 | 8.42E-05 | 0.0057256 |
| L_precuneus | 1.524182971 | 0.718076343 | 2.122591818 | 0.035548093 | 0.443199695 |
| L_lingual | 1.224105836 | 0.597298362 | 2.049404309 | 0.042289699 | 0.443199695 |
| L_parstriangularis | 1.174051603 | 0.636076118 | 1.84577218 | 0.067037583 | 0.468907987 |
| L_superiorfrontal | 1.112074713 | 0.648608878 | 1.71455364 | 0.088639892 | 0.468907987 |
| L_rostralmiddlefrontal | 1.086187673 | 0.643571957 | 1.687748606 | 0.093686079 | 0.468907987 |
| L_parahippocampal | 1.032130866 | 0.564871857 | 1.827194705 | 0.069800149 | 0.468907987 |
| L_posteriorcingulate | 0.954143227 | 0.61833803 | 1.543077056 | 0.125069464 | 0.469817741 |
| R_medialorbitofrontal | 0.946277744 | 0.71107206 | 1.330776158 | 0.185426022 | 0.527454396 |
| L_pericalcarine | 0.926401179 | 0.632241283 | 1.465265246 | 0.145090773 | 0.469817741 |
| R_caudalanteriorcingulate | 0.925710156 | 0.571445907 | 1.619943627 | 0.107494767 | 0.468907987 |
| L_parsopercularis | 0.918010575 | 0.660533995 | 1.389800648 | 0.166795457 | 0.515549594 |
| R_posteriorcingulate | 0.84984268 | 0.576426893 | 1.474328644 | 0.142638263 | 0.469817741 |

| | | | | | |
|---|---|---|---|---|---|
| R_pericalcarine | 0.77342522 | 0.5821606 | 1.328542708 | 0.186160375 | 0.527454396 |
| L_lateraloccipital | 0.757083666 | 0.580447879 | 1.304309472 | 0.194268326 | 0.528409847 |
| L_inferiorparietal | 0.746190865 | 0.637294689 | 1.170872561 | 0.243639084 | 0.591694918 |
| L_insula | 0.720566418 | 0.595189054 | 1.210651326 | 0.22806894 | 0.574395849 |
| R_insula | 0.666046926 | 0.646441057 | 1.030328934 | 0.304632333 | 0.670355049 |
| L_lateralorbitofrontal | 0.600821964 | 0.672953657 | 0.892813282 | 0.373489255 | 0.750719443 |
| L_entorhinal | 0.585702926 | 0.706992017 | 0.828443479 | 0.408829488 | 0.750719443 |
| L_precentral | 0.554404537 | 0.721000505 | 0.768937793 | 0.443225508 | 0.750719443 |
| R_cuneus | 0.523597466 | 0.695233553 | 0.753124563 | 0.452639664 | 0.750719443 |
| L_cuneus | 0.478462089 | 0.591448837 | 0.80896615 | 0.419907274 | 0.750719443 |
| R_inferiorparietal | 0.477272122 | 0.725292647 | 0.658040757 | 0.511592537 | 0.818444719 |
| L_bankssts | 0.469893854 | 0.613896084 | 0.765428981 | 0.445304607 | 0.750719443 |
| L_caudalanteriorcingulate | 0.431230387 | 0.566723825 | 0.760918048 | 0.447985736 | 0.750719443 |
| L_parsorbitalis | 0.390312124 | 0.640466565 | 0.609418423 | 0.543234876 | 0.820888257 |
| L_medialorbitofrontal | 0.375909443 | 0.641160854 | 0.586295062 | 0.558621346 | 0.825788077 |
| L_fusiform | 0.339962232 | 0.625924175 | 0.543136446 | 0.587899573 | 0.850578106 |
| R_parsorbitalis | 0.333090728 | 0.641536438 | 0.519207808 | 0.604435663 | 0.856283856 |
| R_superiorfrontal | 0.280118167 | 0.672688871 | 0.416415641 | 0.677743593 | 0.869557817 |
| R_paracentral | 0.225551888 | 0.674430369 | 0.334433173 | 0.738553197 | 0.91167627 |
| L_inferiortemporal | 0.196979284 | 0.619003371 | 0.31822005 | 0.750792222 | 0.91167627 |
| R_lateraloccipital | 0.158821171 | 0.658716656 | 0.241106961 | 0.80982497 | 0.924689516 |
| R_rostralmiddlefrontal | 0.107231816 | 0.541849946 | 0.197899469 | 0.843410781 | 0.925031179 |
| R_rostralanteriorcingulate | 0.027655006 | 0.523398969 | 0.052837333 | 0.957936835 | 0.986965224 |
| R_isthmuscingulate | 0.024162126 | 0.709612187 | 0.034049762 | 0.972885968 | 0.987406654 |
| L_paracentral | 0.008049656 | 0.693742678 | 0.011603231 | 0.990758688 | 0.990758688 |

| | | | | | |
|---|---|---|---|---|---|
| L_isthmuscingulate | -0.067525564 | 0.68183491 | -0.099035064 | 0.921252093 | 0.96377142 |
| R_parahippocampal | -0.084270603 | 0.655723763 | -0.128515402 | 0.897925543 | 0.954045889 |
| R_fusiform | -0.100559349 | 0.712827058 | -0.141071173 | 0.888016499 | 0.954045889 |
| R_lingual | -0.124933758 | 0.591962254 | -0.211050209 | 0.833154986 | 0.925031179 |
| L_frontalpole | -0.142293375 | 0.56585654 | -0.25146546 | 0.801823002 | 0.924689516 |
| L_temporalpole | -0.145102127 | 0.622069651 | -0.233257043 | 0.815902514 | 0.924689516 |
| R_precentral | -0.168799722 | 0.663799795 | -0.254293122 | 0.799642232 | 0.924689516 |
| L_rostralanteriorcingulate | -0.190989683 | 0.591919271 | -0.322661707 | 0.747432827 | 0.91167627 |
| L_caudalmiddlefrontal | -0.259837081 | 0.597179876 | -0.435106894 | 0.664154846 | 0.869557817 |
| R_parsopercularis | -0.271102082 | 0.633213998 | -0.42813659 | 0.669209596 | 0.869557817 |
| R_parstriangularis | -0.317853389 | 0.734246022 | -0.432897666 | 0.665755285 | 0.869557817 |
| R_lateralorbitofrontal | -0.354678352 | 0.715349115 | -0.495811548 | 0.620804813 | 0.861525047 |
| R_frontalpole | -0.374992452 | 0.454088916 | -0.825812829 | 0.410315327 | 0.750719443 |
| L_transversetemporal | -0.428836112 | 0.694467531 | -0.617503473 | 0.537905783 | 0.820888257 |
| R_superiorparietal | -0.453977108 | 0.699743902 | -0.648776084 | 0.517545925 | 0.818444719 |
| R_supramarginal | -0.579254852 | 0.656272871 | -0.882643299 | 0.37894206 | 0.750719443 |

| | | | | | |
|---|---|---|---|---|---|
| R_inferiortemporal | -0.59211076 | 0.575840264 | -1.028255224 | 0.305603037 | 0.670355049 |
| R_transversetemporal | -0.675932433 | 0.710956101 | -0.950737229 | 0.343376212 | 0.729674451 |
| L_superiorparietal | -0.701074604 | 0.650218125 | -1.078214489 | 0.282792292 | 0.663099167 |
| R_temporalpole | -0.74588283 | 0.607173617 | -1.22845066 | 0.221338928 | 0.574395849 |
| R_entorhinal | -0.941121891 | 0.584476323 | -1.610196776 | 0.110331291 | 0.468907987 |
| R_bankssts | -0.970452563 | 0.656678035 | -1.477820958 | 0.141701891 | 0.469817741 |
| L_supramarginal | -1.041460366 | 0.70742873 | -1.472177086 | 0.143217533 | 0.469817741 |
| L_superiortemporal | -1.052671851 | 0.642846389 | -1.637516939 | 0.103768271 | 0.468907987 |
| R_caudalmiddlefrontal | -1.083736781 | 0.655000448 | -1.654558839 | 0.100254266 | 0.468907987 |
| L_middletemporal | -1.10484268 | 0.602323923 | -1.834299848 | 0.068732622 | 0.468907987 |
| L_postcentral | -1.399513357 | 0.693911281 | -2.016847679 | 0.045623498 | 0.443199695 |
| R_middletemporal | -1.414740069 | 0.563079151 | -2.512506573 | 0.013122091 | 0.223075547 |
| R_postcentral | -1.712521419 | 0.658476365 | -2.600733314 | 0.010301238 | 0.223075547 |
| R_superiortemporal | -1.786465665 | 0.668524006 | -2.672253575 | 0.008447579 | 0.223075547 |

**Table S9. Model estimates from generalized linear mixed-effect model (GLMM).** CT deviation in each region is modeled as the independent variable, while the PPS score was inputted as dependent variable, one-by-one. Beta is not yet standardized by *Z* scores to strengthen interpretability. BH q values indicate the corrected p values by Benjamini-Hochberg method.

Furthermore, for the SA morphmetry, no statistically significant predictors from regional brain deviations remained after Benjamini-Hochberg correction **(Table S10)**.

| Brain Parcel | Beta | Std.Error | T value | P value | BH Q value |
|---|---|---|---|---|---|
| L_caudalmiddlefrontal | 0.913997022 | 0.551170794 | 1.658282754 | 0.099499313 | 0.959866644 |
| L_entorhinal | 0.633168472 | 0.480851792 | 1.316764297 | 0.190069057 | 0.959866644 |
| L_inferiortemporal | 0.753539204 | 0.62065815 | 1.214097011 | 0.226754738 | 0.959866644 |
| L_middletemporal | 0.622139821 | 0.51556909 | 1.206705044 | 0.229580799 | 0.959866644 |
| L_transversetemporal | 0.527390553 | 0.552264155 | 0.954960681 | 0.341243597 | 0.959866644 |
| L_precuneus | 0.561702646 | 0.590182017 | 0.951744766 | 0.34286668 | 0.959866644 |
| L_medialorbitofrontal | 0.54546691 | 0.577710183 | 0.944187807 | 0.346700296 | 0.959866644 |
| L_precentral | 0.547828919 | 0.602106714 | 0.90985353 | 0.364463529 | 0.959866644 |
| R_cuneus | 0.485691693 | 0.550870163 | 0.881680886 | 0.379460626 | 0.959866644 |
| L_inferiorparietal | 0.505708844 | 0.607798195 | 0.832034133 | 0.406806655 | 0.959866644 |
| R_bankssts | 0.426399124 | 0.524117063 | 0.813557036 | 0.417280258 | 0.959866644 |
| R_entorhinal | 0.42849474 | 0.594578761 | 0.720669437 | 0.472314543 | 0.959866644 |
| L_fusiform | 0.40257804 | 0.589122779 | 0.683351679 | 0.495514015 | 0.959866644 |
| L_parahippocampal | 0.444840731 | 0.749752839 | 0.593316501 | 0.55392667 | 0.959866644 |
| L_insula | 0.268611505 | 0.493704729 | 0.544073187 | 0.587256552 | 0.959866644 |
| R_middletemporal | 0.227612475 | 0.523090768 | 0.435129979 | 0.66413813 | 0.959866644 |
| R_posteriorcingulate | 0.240528411 | 0.595708177 | 0.40376886 | 0.686998764 | 0.959866644 |
| R_frontalpole | 0.280062125 | 0.722391955 | 0.387687215 | 0.698836316 | 0.959866644 |
| R_paracentral | 0.269574052 | 0.705269424 | 0.382228469 | 0.702871512 | 0.959866644 |
| R_superiorfrontal | 0.215363038 | 0.619576826 | 0.347596988 | 0.728664903 | 0.959866644 |
| L_cuneus | 0.20056681 | 0.598521349 | 0.335103853 | 0.738048326 | 0.959866644 |
| L_superiortemporal | 0.164371854 | 0.494086009 | 0.332678624 | 0.739874515 | 0.959866644 |

| | | | | | |
|---|---|---|---|---|---|
| L_postcentral | 0.18745485 | 0.566104276 | 0.331131309 | 0.741040413 | 0.959866644 |
| R_caudalmiddlefrontal | 0.121284013 | 0.512957589 | 0.236440625 | 0.813436358 | 0.959866644 |
| R_pericalcarine | 0.132766168 | 0.568242757 | 0.233643397 | 0.815603127 | 0.959866644 |
| R_rostralmiddlefrontal | 0.118665503 | 0.566836142 | 0.209347101 | 0.834481603 | 0.959866644 |
| L_supramarginal | 0.105173727 | 0.518274639 | 0.202930491 | 0.839483994 | 0.959866644 |
| L_superiorparietal | 0.105320821 | 0.566886869 | 0.185788076 | 0.852879868 | 0.959866644 |
| R_insula | 0.085386007 | 0.48017235 | 0.177823665 | 0.859118499 | 0.959866644 |
| L_bankssts | 0.087555269 | 0.499313048 | 0.175351455 | 0.861056842 | 0.959866644 |
| R_lateraloccipital | 0.084077599 | 0.565126335 | 0.148776643 | 0.881943969 | 0.9619333 |
| R_medialorbitofrontal | 0.084583809 | 0.617258968 | 0.137031317 | 0.89120291 | 0.9619333 |
| R_lateralorbitofrontal | 0.06317758 | 0.655370998 | 0.096399718 | 0.923340974 | 0.981049785 |
| L_rostralanteriorcingulate | 0.038112318 | 0.578822298 | 0.065844592 | 0.947595486 | 0.98168373 |
| R_precentral | 0.032280358 | 0.544506426 | 0.059283705 | 0.952810679 | 0.98168373 |
| R_parsopercularis | -0.002911058 | 0.545031077 | -0.005341086 | 0.995746054 | 0.995746054 |
| R_postcentral | -0.017090123 | 0.600969969 | -0.028437566 | 0.977353661 | 0.991941029 |
| L_superiorfrontal | -0.110799767 | 0.580224518 | -0.190960161 | 0.848833455 | 0.959866644 |
| R_isthmuscingulate | -0.133841395 | 0.635270533 | -0.210684092 | 0.833440128 | 0.959866644 |
| R_superiortemporal | -0.148421822 | 0.564760034 | -0.262805109 | 0.793087148 | 0.959866644 |
| R_inferiorparietal | -0.198528588 | 0.632664263 | -0.313797696 | 0.754141768 | 0.959866644 |
| R_lingual | -0.216096743 | 0.541477769 | -0.399087008 | 0.690437194 | 0.959866644 |
| R_supramarginal | -0.271253677 | 0.621167159 | -0.436683866 | 0.66301338 | 0.959866644 |

| | | | | | |
|---|---|---|---|---|---|
| L_temporalpole | -0.269180395 | 0.572448593 | -0.470226319 | 0.638925028 | 0.959866644 |
| L_caudalanteriorcingulate | -0.288261486 | 0.574690181 | -0.501594591 | 0.616740591 | 0.959866644 |
| L_rostralmiddlefrontal | -0.285326978 | 0.561302905 | -0.508329772 | 0.612022147 | 0.959866644 |
| R_inferiortemporal | -0.276800268 | 0.538014283 | -0.514484981 | 0.607724187 | 0.959866644 |
| R_parstriangularis | -0.294906132 | 0.551831194 | -0.534413667 | 0.593903021 | 0.959866644 |
| R_parahippocampal | -0.347562551 | 0.639372863 | -0.54359916 | 0.587581904 | 0.959866644 |
| L_lingual | -0.340112472 | 0.623839514 | -0.545192256 | 0.586488804 | 0.959866644 |
| L_pericalcarine | -0.379469311 | 0.593251711 | -0.639643011 | 0.523450026 | 0.959866644 |
| L_paracentral | -0.430832143 | 0.663573819 | -0.64926031 | 0.517233872 | 0.959866644 |
| R_caudalanteriorcingulate | -0.410145015 | 0.600471671 | -0.683038077 | 0.495711546 | 0.959866644 |
| L_frontalpole | -0.534789988 | 0.755274607 | -0.708073571 | 0.480076774 | 0.959866644 |
| R_fusiform | -0.43084894 | 0.607070939 | -0.709717616 | 0.479059655 | 0.959866644 |
| R_rostralanteriorcingulate | -0.53132547 | 0.722032065 | -0.735875172 | 0.463037793 | 0.959866644 |
| L_lateralorbitofrontal | -0.558546973 | 0.643950772 | -0.867375267 | 0.387220689 | 0.959866644 |
| L_isthmuscingulate | -0.758936828 | 0.642471831 | -1.181276426 | 0.239495733 | 0.959866644 |
| L_parsopercularis | -0.652105016 | 0.546536033 | -1.19316015 | 0.234824763 | 0.959866644 |

| | | | | | |
|---|---|---|---|---|---|
| L_posteriorcingulate | -0.800482962 | 0.587301138 | -1.362985547 | 0.175075245 | 0.959866644 |
| R_superiorparietal | -0.86159522 | 0.609113204 | -1.414507541 | 0.159433118 | 0.959866644 |
| R_precuneus | -0.93260303 | 0.628011456 | -1.485009582 | 0.139789481 | 0.959866644 |
| L_parsorbitalis | -1.011707959 | 0.59205225 | -1.708815328 | 0.089701114 | 0.959866644 |
| L_lateraloccipital | -1.060047054 | 0.589465875 | -1.798317934 | 0.074281307 | 0.959866644 |
| R_temporalpole | -1.022013748 | 0.540438605 | -1.891082053 | 0.060679423 | 0.959866644 |
| L_parstriangularis | -1.329195497 | 0.588726645 | -2.257746458 | 0.025508698 | 0.578197155 |
| R_parsorbitalis | -1.284254416 | 0.565880992 | -2.269477918 | 0.024829582 | 0.578197155 |
| R_transversetemporal | -2.03289925 | 0.753284716 | -2.698712991 | 0.007818104 | 0.531631072 |

**Table S10. Model estimates from generalized linear mixed-effect model (GLMM).** SA deviation in each region is modeled as the independent variable, while the PPS score was inputted as dependent variable, one-by-one. Beta is not yet standardized by *Z* scores to strengthen interpretability. BH *q* values indicate the corrected p values by Benjamini-Hochberg method.

Finally, for the SV morphometry, there are two regions reaching marginal statistical significance, including bilateral NAcc. Full results have been tabulated into the **Table S11**.

| Brain Parcel | Beta | Std.Error | T value | P value | BH Q value |
|---|---|---|---|---|---|
| Raccumb | 1.494782823 | 0.573874108 | 2.604722538 | 0.010187664 | 0.142627296 |
| Laccumb | 1.242925155 | 0.580393091 | 2.141523003 | 0.033963081 | 0.237741567 |
| Lhippo | 1.368782617 | 0.750623197 | 1.823528267 | 0.07035638 | 0.254258152 |
| Rhippo | 1.120245109 | 0.628477232 | 1.782475246 | 0.076839109 | 0.254258152 |

| | | | | | |
|---|---|---|---|---|---|
| Lput | 1.055066068 | 0.619460571 | 1.703201331 | 0.090806483 | 0.254258152 |
| Lamyg | 0.603197557 | 0.564995546 | 1.067614712 | 0.287531845 | 0.579202731 |
| Rthal | 0.695229838 | 0.654355616 | 1.062464845 | 0.289853986 | 0.579202731 |
| Ramyg | 0.516914696 | 0.560946883 | 0.92150382 | 0.3583727 | 0.579202731 |
| Rput | 0.455828705 | 0.594363236 | 0.766919414 | 0.444420785 | 0.579202731 |
| Lcaud | 0.492456981 | 0.703126043 | 0.700382223 | 0.48485094 | 0.579202731 |
| Lthal | 0.475849166 | 0.69787826 | 0.681851252 | 0.496459484 | 0.579202731 |
| Lpal | 0.2413306 | 0.705284924 | 0.342174619 | 0.732732621 | 0.789096669 |
| Rcaud | 0.111381045 | 0.711700359 | 0.156499914 | 0.875864441 | 0.875864441 |
| Rpal | -0.536312236 | 0.650487839 | -0.824476961 | 0.411071089 | 0.579202731 |

**Table S11. Model estimates from generalized linear mixed-effect model (GLMM).** SV deviation in each region is modeled as the independent variable, while the PPS score was inputted as dependent variable, one-by-one. Beta is not yet standardized by *Z* scores to strengthen interpretability. BH *q* values indicate the corrected p values by Benjamini-Hochberg method.

**4. Bivariate AE model to estimate shared neurogenetic substrates**

The bivariate Additive Genetic – Common Environmental – Unique Environmental (ACE) model was initially fitted to decompose the covariance between PPS and regional neuroanatomical deviations into three latent components: additive genetic effects (A), shared environmental influences (C), and unique environmental influences (E). This analysis was performed separately for associations of PPS to identified brain regions using the OpenMx package (v2.18.1) in R. As we mentioned above, model performance was evaluated using likelihood ratio tests (LRTs), Akaike Information Criterion (AIC), and Bayesian Information Criterion (BIC), comparing the full ACE model against the reduced AE and E models. Across most brain regions, the AE model demonstrated superior fit, as indicated by lower AIC and BIC values, while the shared environmental component (C) contributed negligibly to the covariance between PPS and neuroanatomical variations. The AE model was selected as the final model, confirming that shared genetic and unique environmental factors primarily explained the associations between PPS and brain deviations. Full results of modelling genetic factors on the bivariate correlation between brain deviations and PPS can be found at **Table S12-14**.

| Region | $r_g$ | $r_g$l | $r_g$u | $r_{ph}$ | $r_{ph}$l | $r_{ph}$u | $h^2$ | $h^2$l | $h^2$u |
|---|---|---|---|---|---|---|---|---|---|
| Laccumb | 0.4384 | 0.4384 | 1.0000 | 0.1587 | -0.0131 | 0.3217 | 0.5708 | 0.4838 | 0.5701 |
| Lamyg | 0.3559 | 0.3559 | 1.0000 | 0.0710 | -0.1027 | 0.2429 | 0.5356 | 0.4072 | 0.5302 |
| Lcaud | 0.0504 | -0.2730 | 0.0504 | 0.0697 | -0.1077 | 0.2439 | 0.6242 | 0.6469 | 0.5742 |
| Lhippo | 0.4781 | 0.0623 | 1.0000 | 0.1409 | -0.0318 | 0.3082 | 0.6154 | 0.5554 | 0.5973 |
| Lpal | 0.1469 | 0.1469 | 0.6610 | 0.0305 | -0.1459 | 0.2064 | 0.5515 | 0.4912 | 0.5245 |
| Lput | 0.1935 | 0.1935 | 0.8588 | 0.1062 | -0.0709 | 0.2763 | 0.6807 | 0.7041 | 0.6329 |
| Lthal | 0.1332 | 0.1332 | 1.0000 | 0.0011 | -0.1786 | 0.1786 | 0.5465 | 0.4762 | 0.5197 |
| **Raccumb** | **0.8899** | **0.8899** | **1.0000** | **0.1846** | **0.0174** | **0.3455** | **0.5980** | **0.5594** | **0.6289** |
| Ramyg | 0.2485 | 0.2485 | 0.9268 | 0.0620 | -0.1138 | 0.2364 | 0.5138 | 0.3835 | 0.5061 |
| Rcaud | 0.0349 | 0.0349 | 0.4607 | 0.0111 | -0.1665 | 0.1879 | 0.5912 | 0.5965 | 0.5467 |
| Rhippo | 0.5844 | 0.5844 | 1.0000 | 0.1547 | -0.0167 | 0.3207 | 0.5265 | 0.3538 | 0.5415 |
| Rpal | -0.0431 | -0.2025 | 0.5019 | -0.0744 | -0.2475 | 0.1032 | 0.4951 | 0.4419 | 0.4706 |
| Rput | 0.2016 | 0.2016 | 1.0000 | 0.0347 | -0.1419 | 0.2080 | 0.6511 | 0.6583 | 0.6099 |
| Rthal | 0.1952 | 0.1952 | 1.0000 | 0.0356 | -0.1393 | 0.2074 | 0.5487 | 0.4747 | 0.5305 |

**Table S12. Bivariate AE model estimations for brain deviations on SV.** $r_g$ = genetic association, $r_g$l ($r_g$u) = lower (upper) bound of genetic association; $r_{ph}$ = phenotypic association, $r_{ph}$l ($r_{ph}$u) = lower (upper) bound of phenotypic association;

$h2$ = heritability, $h^2$l ($h^2$u) = lower (upper) bound of heritability. Statistically significant shared neurogenetic associations are bold.

| **Region** | $r_g$ | $r_g$**l** | $r_g$**u** | $r_{ph}$ | $r_{ph}$**l** | $r_{ph}$**u** | $h^2$ | $h^2$**l** |
|---|---|---|---|---|---|---|---|---|
| L_bankssts | 1.0000 | -1.0000 | | 0.0683 | -0.0988 | 0.2344 | 0.3263 | 0.049 |
| L_caudalanteriorcingulate | 0.1188 | -0.1824 | 0.7427 | 0.0645 | -0.1093 | 0.2362 | 0.4232 | 0.219 |
| L_caudalmiddlefrontal | -0.3389 | -1.0000 | 1.0000 | -0.0189 | -0.1906 | 0.1513 | 0.2253 | 0.037 |
| L_cuneus | 0.1551 | -1.0000 | 1.0000 | 0.0699 | -0.0983 | 0.2343 | 0.2454 | 0.031 |
| L_entorhinal | -0.5135 | -1.0000 | 1.0000 | 0.0812 | -0.0905 | 0.2458 | 0.1527 | 0.037 |
| L_fusiform | -0.2163 | -1.0000 | 1.0000 | 0.0572 | -0.1152 | 0.2233 | 0.2224 | 0.046 |
| L_inferiorparietal | 0.0149 | -0.8097 | 0.6197 | 0.0925 | -0.0818 | 0.2595 | 0.3487 | 0.089 |
| L_inferiortemporal | 0.0453 | -1.0000 | 1.0000 | 0.0520 | -0.1175 | 0.2180 | 0.2630 | 0.032 |
| L_isthmuscingulate | 0.0554 | -0.3094 | 0.6608 | 0.0136 | -0.1642 | 0.1909 | 0.4888 | 0.383 |
| L_lateraloccipital | -0.0656 | -0.6638 | -0.0656 | 0.0642 | -0.1157 | 0.2370 | 0.4473 | 0.342 |
| L_lateralorbitofrontal | 0.5399 | -0.0576 | 1.0000 | 0.1044 | -0.0679 | 0.2746 | 0.4732 | 0.218 |
| L_lingual | 0.3364 | -1.0000 | 1.0000 | 0.1841 | 0.0145 | 0.3447 | 0.3902 | 0.036 |
| L_medialorbitofrontal | 0.5355 | | 1.0000 | 0.0422 | -0.1252 | 0.2091 | 0.3020 | 0.033 |
| L_middletemporal | -0.3714 | -1.0000 | 0.3489 | -0.1427 | -0.3066 | 0.0275 | 0.2390 | 0.046 |
| L_parahippocampal | 0.3051 | 0.3051 | 0.9450 | 0.1438 | -0.0334 | 0.3142 | 0.5822 | 0.507 |
| L_paracentral | 0.3271 | -0.3656 | 1.0000 | 0.0042 | -0.1700 | 0.1843 | 0.4439 | 0.165 |
| L_parsopercularis | 0.4862 | -0.3525 | 1.0000 | 0.1248 | -0.0480 | 0.2964 | 0.4362 | 0.079 |
| L_parsorbitalis | 0.1222 | 0.0509 | 0.8329 | 0.0380 | -0.1365 | 0.2113 | 0.4251 | 0.208 |
| L_parstriangularis | 0.2117 | -1.0000 | 1.0000 | 0.1652 | -0.0047 | 0.3289 | 0.3436 | |
| L_pericalcarine | 0.5230 | | 1.0000 | 0.1153 | -0.0551 | 0.2842 | 0.3757 | |
| L_postcentral | -0.4648 | -1.0000 | -0.4648 | -0.1464 | -0.3150 | 0.0279 | 0.3200 | 0.299 |
| L_posteriorcingulate | 0.1409 | -0.7536 | 0.1409 | 0.1345 | -0.0417 | 0.3062 | 0.4429 | 0.218 |
| L_precentral | 0.2771 | -0.5380 | 1.0000 | 0.0602 | -0.1124 | 0.2330 | 0.3972 | 0.055 |
| L_precuneus | 0.4664 | -0.1766 | 1.0000 | 0.1655 | -0.0043 | 0.3286 | 0.4415 | 0.159 |
| L_rostralanteriorcingulate | 0.4448 | -1.0000 | 1.0000 | -0.0245 | -0.1902 | 0.1438 | 0.2767 | 0.029 |
| L_rostralmiddlefrontal | 0.3276 | 0.3276 | 1.0000 | 0.1558 | -0.0183 | 0.3231 | 0.5067 | 0.335 |
| L_superiorfrontal | 0.4371 | 0.4371 | 1.0000 | 0.1315 | -0.0422 | 0.3001 | 0.4935 | 0.270 |
| L_superiorparietal | -0.2755 | -0.9248 | 0.5118 | -0.1211 | -0.2937 | 0.0544 | 0.3227 | 0.125 |
| L_superiortemporal | -0.5331 | -1.0000 | 0.0342 | -0.1362 | -0.3029 | 0.0354 | 0.2290 | 0.141 |
| L_supramarginal | -0.2223 | -0.8892 | -0.2196 | -0.1117 | -0.2810 | 0.0631 | 0.3422 | 0.173 |
| L_frontalpole | 0.1009 | -0.2749 | 0.8327 | -0.0179 | -0.1904 | 0.1610 | 0.3954 | 0.132 |
| L_temporalpole | -0.6365 | -1.0000 | 1.0000 | 0.0104 | -0.1583 | 0.1767 | 0.1138 | |
| L_transversetemporal | -0.1914 | -0.9889 | 0.5179 | -0.0573 | -0.2289 | 0.1160 | 0.3186 | 0.108 |
| L_insula | 0.1160 | -0.5815 | 0.1160 | 0.0966 | -0.0808 | 0.2720 | 0.4871 | 0.351 |
| R_bankssts | 0.0077 | -1.0000 | 1.0000 | -0.1273 | -0.2911 | 0.0453 | 0.3052 | 0.040 |
| R_caudalanteriorcingulate | 1.0000 | -1.0000 | | 0.1349 | -0.0319 | 0.2978 | 0.3324 | 0.058 |
| R_caudalmiddlefrontal | -0.1193 | | 0.7008 | -0.1468 | -0.3160 | 0.0287 | 0.4155 | 0.293 |

| | | | | | | | | |
|---|---|---|---|---|---|---|---|---|
| R_cuneus | 0.3241 | -0.3463 | 1.0000 | 0.0641 | -0.1091 | 0.2354 | 0.4301 | 0.139 |
| R_entorhinal | -0.0977 | -1.0000 | 1.0000 | -0.1433 | -0.3066 | 0.0295 | 0.3057 | 0.039 |
| R_fusiform | 0.4532 | -0.1501 | 1.0000 | 0.0120 | -0.1614 | 0.1887 | 0.4751 | 0.226 |
| R_inferiorparietal | -0.0402 | -1.0000 | 1.0000 | 0.0260 | -0.1493 | 0.1962 | 0.3015 | |
| R_inferiortemporal | -1.0000 | | 1.0000 | -0.0998 | -0.2612 | 0.0663 | 0.1487 | |
| R_isthmuscingulate | 0.1100 | -0.1902 | 0.5899 | 0.0190 | -0.1551 | 0.1962 | 0.4580 | 0.304 |
| R_lateraloccipital | 0.2931 | -0.3582 | 1.0000 | 0.0058 | -0.1657 | 0.1771 | 0.4023 | 0.094 |
| R_lateralorbitofrontal | 0.2670 | -1.0000 | 1.0000 | -0.0528 | -0.2198 | 0.1197 | 0.3299 | |
| R_lingual | 0.4269 | -1.0000 | 1.0000 | -0.0194 | -0.1873 | 0.1535 | 0.3303 | |
| R_medialorbitofrontal | 0.1117 | -0.6704 | 1.0000 | 0.1129 | -0.0637 | 0.2794 | 0.3712 | 0.064 |
| R_middletemporal | -0.0842 | -1.0000 | 1.0000 | -0.1790 | -0.3372 | -0.0105 | 0.2459 | 0.037 |
| R_parahippocampal | -0.0076 | -0.6419 | 0.1484 | -0.0220 | -0.1956 | 0.1541 | 0.4186 | 0.270 |
| R_paracentral | 0.0379 | -0.6261 | 0.7775 | 0.0002 | -0.1726 | 0.1729 | 0.3888 | 0.172 |
| R_parsopercularis | -0.9925 | | 1.0000 | -0.0605 | -0.2258 | 0.1059 | 0.1164 | |
| R_parsorbitalis | 0.1349 | -1.0000 | 1.0000 | 0.0350 | -0.1373 | 0.2061 | 0.3505 | 0.036 |
| R_parstriangularis | -0.0579 | -0.8007 | 0.6884 | -0.0557 | -0.2291 | 0.1220 | 0.3808 | 0.157 |
| R_pericalcarine | 0.1322 | -0.1937 | 0.5803 | 0.1131 | -0.0641 | 0.2843 | 0.5125 | 0.411 |
| R_postcentral | -0.2535 | -0.9094 | -0.2535 | -0.1872 | -0.3530 | -0.0102 | 0.4340 | 0.408 |
| R_posteriorcingulate | 0.0562 | -0.5492 | 0.1215 | 0.1363 | -0.0420 | 0.3068 | 0.5420 | 0.498 |
| R_precentral | 0.0869 | -0.4291 | 0.7351 | -0.0295 | -0.2045 | 0.1490 | 0.4863 | 0.365 |
| R_precuneus | 0.4675 | -1.0000 | 1.0000 | 0.3008 | 0.1387 | 0.4499 | 0.3939 | |
| R_rostralanteriorcingulate | 0.2060 | | 1.0000 | 0.0067 | -0.1610 | 0.1740 | 0.2586 | 0.023 |
| R_rostralmiddlefrontal | 0.0682 | -0.5023 | 0.6484 | 0.0262 | -0.1562 | 0.2084 | 0.5258 | 0.442 |
| R_superiorfrontal | 0.2487 | -1.0000 | 1.0000 | 0.0359 | -0.1363 | 0.2079 | 0.3713 | 0.032 |
| R_superiorparietal | -0.4185 | -1.0000 | -0.4185 | -0.0543 | -0.2271 | 0.1191 | 0.2812 | 0.173 |
| R_superiortemporal | -0.5914 | -1.0000 | 0.0671 | -0.2276 | -0.3876 | -0.0588 | 0.1895 | 0.052 |
| R_supramarginal | -0.2105 | -1.0000 | 0.4124 | -0.0414 | -0.2116 | 0.1304 | 0.3068 | 0.110 |
| R_frontalpole | -0.2858 | -1.0000 | 0.3967 | -0.0686 | -0.2413 | 0.1049 | 0.3025 | 0.107 |
| R_temporalpole | -1.0000 | | -0.1033 | -0.1129 | -0.2776 | 0.0550 | 0.0374 | 0.044 |
| R_transversetemporal | 0.1429 | -0.5217 | 1.0000 | -0.0783 | -0.2463 | 0.0970 | 0.3770 | 0.095 |
| R_insula | 0.6284 | -0.1102 | 1.0000 | 0.0730 | -0.0978 | 0.2433 | 0.4217 | 0.095 |

**Table S13. Bivariate AE model estimations for brain deviations on CT.** $r_g$ = genetic association, $r_g$l ($r_g$u) = lower (upper) bound of genetic association; $r_{ph}$ = phenotypic association, $r_{ph}$l ($r_{ph}$u) = lower (upper) bound of phenotypic association; $h2$ = heritability, $h^2$l ($h^2$u) = lower (upper) bound of heritability. Statistically significant shared neurogenetic associations are bold. Empty indicates that this estimate falls to be modeled with acceptable goodness-of-fits.

| Region | $r_g$ | $r_g$l | $r_g$u | $r_{ph}$ | $r_{ph}$l | $r_{ph}$u | $h^2$ | $h^2$l | $h$ |
|---|---|---|---|---|---|---|---|---|---|
| L_bankssts | -0.4741 | -1.0000 | 0.2737 | -0.0016 | -0.1709 | 0.1661 | 0.1715 | 0.0209 | 0. |

| | | | | | | | | | |
|---|---|---|---|---|---|---|---|---|---|
| L_caudalanteriorcingulate | -0.1248 | -1.0000 | 1.0000 | -0.0460 | -0.2142 | 0.1239 | 0.2561 | | |
| L_caudalmiddlefrontal | 0.2804 | -1.0000 | 1.0000 | 0.1330 | -0.0372 | 0.2957 | 0.3491 | 0.0174 | 0.[illegible] |
| L_cuneus | -0.0573 | -1.0000 | 1.0000 | 0.0392 | -0.1326 | 0.2067 | 0.2781 | | |
| L_entorhinal | -1.0000 | | 1.0000 | 0.1149 | -0.0539 | 0.2750 | 0.1204 | | |
| L_fusiform | -1.0000 | | | 0.0667 | -0.1018 | 0.2296 | 0.1492 | | |
| L_inferiorparietal | 0.3674 | | 1.0000 | 0.0642 | -0.1047 | 0.2331 | 0.3093 | 0.0424 | 0.[illegible] |
| L_inferiortemporal | -0.0052 | -1.0000 | | 0.0854 | -0.0816 | 0.2480 | 0.2030 | | |
| L_isthmuscingulate | -0.1816 | -0.8193 | 0.4057 | -0.0926 | -0.2644 | 0.0828 | 0.3884 | 0.2696 | 0.[illegible] |
| L_lateraloccipital | -0.0182 | -1.0000 | 1.0000 | -0.1538 | -0.3162 | 0.0157 | 0.2924 | 0.0151 | 0.[illegible] |
| L_lateralorbitofrontal | -0.1685 | -1.0000 | 1.0000 | -0.0793 | -0.2464 | 0.0911 | 0.2628 | | |
| L_lingual | -0.1912 | -0.9106 | -0.1912 | -0.0627 | -0.2361 | 0.1114 | 0.3299 | 0.1236 | 0.[illegible] |
| L_medialorbitofrontal | 0.6488 | -1.0000 | 1.0000 | 0.0759 | -0.0941 | 0.2452 | 0.3880 | 0.0496 | 0.[illegible] |
| L_middletemporal | 0.1302 | -1.0000 | 1.0000 | 0.0726 | -0.0977 | 0.2399 | 0.3153 | | |
| L_parahippocampal | 0.5777 | -0.2789 | 1.0000 | 0.0404 | -0.1288 | 0.2100 | 0.3767 | 0.0404 | 0.[illegible] |
| L_paracentral | 0.0657 | -1.0000 | 1.0000 | -0.0639 | -0.2318 | 0.1080 | 0.3299 | 0.0323 | 0.[illegible] |
| L_parsopercularis | -0.0988 | -0.8296 | 0.4777 | -0.0635 | -0.2386 | 0.1166 | 0.4377 | 0.2728 | 0.[illegible] |
| L_parsorbitalis | 0.2408 | -1.0000 | 1.0000 | -0.1419 | -0.3025 | 0.0292 | 0.2751 | 0.0428 | 0.[illegible] |
| L_parstriangularis | -0.0377 | -0.6456 | 0.9070 | -0.1727 | -0.3348 | 0.0013 | 0.3522 | 0.1158 | 0.[illegible] |
| L_pericalcarine | -0.0245 | -0.0245 | 0.8161 | -0.0565 | -0.2333 | 0.1210 | 0.4276 | 0.2627 | 0.[illegible] |
| L_postcentral | -0.1266 | -1.0000 | 0.6599 | 0.0022 | -0.1700 | 0.1731 | 0.3034 | 0.0386 | 0.[illegible] |
| L_posteriorcingulate | 0.6629 | -0.1606 | 1.0000 | -0.0888 | -0.2557 | 0.0882 | 0.4008 | 0.1027 | 0.[illegible] |
| L_precentral | -0.1074 | -1.0000 | 1.0000 | 0.0906 | -0.0798 | 0.2544 | 0.2262 | 0.0364 | 0.[illegible] |
| L_precuneus | 0.3160 | -0.1957 | 0.7651 | 0.1300 | -0.0569 | 0.3175 | 0.5802 | 0.4540 | 0.[illegible] |
| L_rostralanteriorcingulate | -0.3513 | -1.0000 | 0.3178 | 0.0013 | -0.1705 | 0.1710 | 0.2396 | 0.0427 | 0.[illegible] |
| L_rostralmiddlefrontal | 0.9480 | -1.0000 | | -0.0123 | -0.1777 | 0.1571 | 0.2676 | | |

| | | | | | | | | | |
|---|---|---|---|---|---|---|---|---|---|
| L_superiorfrontal | -0.3960 | -1.0000 | 1.0000 | -0.0047 | -0.1770 | 0.1652 | 0.2093 | 0.0344 | 0.[illegible] |
| L_superiorparietal | -0.1116 | -1.0000 | 1.0000 | 0.0170 | -0.1521 | 0.1850 | 0.2440 | 0.0282 | 0.[illegible] |
| L_superiortemporal | -0.2998 | -1.0000 | 1.0000 | -0.0024 | -0.1705 | 0.1648 | 0.1823 | | |
| L_supramarginal | 0.0469 | -1.0000 | 1.0000 | 0.0073 | -0.1631 | 0.1769 | 0.3159 | 0.0336 | 0.[illegible] |
| L_frontalpole | -1.0000 | | 1.0000 | -0.0542 | -0.2214 | 0.1127 | 0.1034 | | |
| L_temporalpole | -0.1587 | -1.0000 | 0.3682 | -0.0105 | -0.1848 | 0.1630 | 0.3387 | 0.1204 | 0.[illegible] |
| L_transversetemporal | 0.2252 | -1.0000 | 1.0000 | 0.1161 | -0.0536 | 0.2801 | 0.3172 | | |
| L_insula | -0.2216 | -1.0000 | 0.4198 | 0.0502 | -0.1232 | 0.2183 | 0.2813 | 0.0670 | 0.[illegible] |
| R_bankssts | 0.3855 | -1.0000 | 1.0000 | 0.0500 | -0.1197 | 0.2192 | 0.3626 | 0.0382 | 0.[illegible] |
| R_caudalanteriorcingulate | 0.0746 | -1.0000 | 1.0000 | -0.0506 | -0.2182 | 0.1223 | 0.3208 | 0.0429 | 0.[illegible] |
| R_caudalmiddlefrontal | 0.3150 | -1.0000 | 1.0000 | 0.0352 | -0.1338 | 0.2034 | 0.3221 | | |
| R_cuneus | 0.0778 | -1.0000 | 1.0000 | 0.0886 | -0.0839 | 0.2546 | 0.3197 | | |
| R_entorhinal | -0.3569 | -1.0000 | 1.0000 | 0.0760 | -0.0937 | 0.2402 | 0.1790 | 0.0270 | 0.[illegible] |
| R_fusiform | -1.0000 | | -0.9978 | -0.1523 | -0.3259 | 0.0217 | 0.0783 | 0.1058 | 0.[illegible] |
| R_inferiorparietal | 1.0000 | -1.0000 | | -0.0314 | -0.1952 | 0.1350 | 0.2116 | | |
| R_inferiortemporal | -0.5133 | -1.0000 | -0.5133 | -0.0465 | -0.2136 | 0.1218 | 0.1538 | 0.0216 | 0.[illegible] |
| R_isthmuscingulate | 0.2979 | -0.3999 | 1.0000 | -0.0124 | -0.1815 | 0.1586 | 0.3645 | 0.0333 | 0.[illegible] |
| R_lateraloccipital | -0.4829 | -1.0000 | 1.0000 | 0.0019 | -0.1757 | 0.1731 | 0.1977 | 0.0737 | 0.[illegible] |
| R_lateralorbitofrontal | 0.2205 | -1.0000 | 1.0000 | 0.0084 | -0.1592 | 0.1762 | 0.2731 | 0.0296 | 0.[illegible] |
| R_lingual | -0.0005 | -0.7013 | -0.0005 | -0.0144 | -0.1878 | 0.1613 | 0.4088 | 0.2277 | 0.[illegible] |
| R_medialorbitofrontal | 0.6687 | -0.1184 | 1.0000 | 0.0255 | -0.1465 | 0.2017 | 0.4246 | 0.1018 | 0.[illegible] |
| R_middletemporal | 0.3545 | 0.3545 | 0.9371 | 0.0245 | -0.1541 | 0.2070 | 0.5622 | 0.4589 | 0.[illegible] |
| R_parahippocampal | 0.1089 | -0.5007 | 1.0000 | -0.0550 | -0.2233 | 0.1173 | 0.3617 | 0.1024 | 0.[illegible] |
| R_paracentral | 0.0243 | -1.0000 | 1.0000 | 0.0432 | -0.1274 | 0.2125 | 0.3034 | | |
| R_parsopercularis | -0.1866 | -1.0000 | 1.0000 | 0.0097 | -0.1605 | 0.1780 | 0.2264 | 0.0346 | 0.[illegible] |

| | | | | | | | | | |
|---|---|---|---|---|---|---|---|---|---|
| R_parsorbitalis | -0.5083 | -1.0000 | 1.0000 | -0.2159 | -0.3759 | -0.0473 | 0.1945 | 0.0422 | 0.[illegible] |
| R_parstriangularis | 0.0459 | -1.0000 | 1.0000 | -0.0334 | -0.2006 | 0.1365 | 0.2654 | 0.0343 | 0.[illegible] |
| R_pericalcarine | 0.1283 | -0.5868 | 1.0000 | 0.0251 | -0.1482 | 0.1961 | 0.3685 | 0.0568 | 0.[illegible] |
| R_postcentral | -0.1718 | -0.8058 | 0.1009 | -0.0381 | -0.2137 | 0.1359 | 0.3633 | 0.2010 | 0.[illegible] |
| R_posteriorcingulate | 0.6903 | -1.0000 | 1.0000 | 0.0503 | -0.1175 | 0.2178 | 0.3350 | | |
| R_precentral | 0.0193 | -0.2939 | 0.7718 | 0.0191 | -0.1551 | 0.1903 | 0.3903 | 0.1940 | 0.[illegible] |
| R_precuneus | -0.4159 | -1.0000 | 1.0000 | -0.1336 | -0.3001 | 0.0370 | 0.2119 | 0.0473 | 0.[illegible] |
| R_rostralanteriorcingulate | -1.0000 | | -0.1130 | -0.0636 | -0.2323 | 0.1055 | 0.0120 | 0.0150 | 0.[illegible] |
| R_rostralmiddlefrontal | 0.1340 | 0.1340 | 0.8713 | -0.0556 | -0.2299 | 0.1253 | 0.4707 | 0.3243 | 0.[illegible] |
| R_superiorfrontal | 0.0094 | -0.8730 | 0.2782 | 0.0425 | -0.1303 | 0.2124 | 0.3399 | 0.0632 | 0.[illegible] |
| R_superiorparietal | 0.3138 | -0.2591 | 1.0000 | -0.0856 | -0.2592 | 0.0996 | 0.4806 | 0.2609 | 0.[illegible] |
| R_superiortemporal | -0.1276 | -1.0000 | 1.0000 | -0.0423 | -0.2126 | 0.1285 | 0.2801 | | |
| R_supramarginal | -0.3265 | -1.0000 | 1.0000 | -0.0887 | -0.2575 | 0.0824 | 0.2441 | 0.0476 | 0.[illegible] |
| R_frontalpole | 0.4681 | -1.0000 | 1.0000 | 0.0239 | -0.1469 | 0.1977 | 0.3960 | 0.0620 | 0.[illegible] |
| R_temporalpole | -0.9564 | | 1.0000 | -0.1598 | -0.3199 | 0.0062 | 0.0634 | | |
| R_transversetemporal | -1.0000 | | 1.0000 | -0.2080 | -0.3622 | -0.0445 | 0.1152 | | |
| R_insula | -0.2000 | -0.9997 | 0.4123 | 0.0093 | -0.1649 | 0.1802 | 0.3196 | 0.1318 | 0.[illegible] |

**Table S14. Bivariate AE model estimations for brain deviations on SA.** $r_g$ = genetic association, $r_g$l ($r_g$u) = lower (upper) bound of genetic association; $r_{ph}$ = phenotypic association, $r_{ph}$l ($r_{ph}$u) = lower (upper) bound of phenotypic association; *h*2 = heritability, $h^2$l ($h^2$u) = lower (upper) bound of heritability. Statistically significant shared neurogenetic associations are bold. Empty indicates that this estimate falls to be modeled with acceptable goodness-of-fits.

**5. Network Enrichment Significance Testing and Theory-Driven Specificity**

Network Enrichment Significance Testing (NEST) was applied to determine whether brain morphometric deviations associated with psychopathological procrastination (PPS) exhibited significant spatial enrichment within large-scale brain networks (Weinstein et al., 2024). Leveraging the Yeo-7 functional atlas, this analysis assessed whether PPS-related deviations were non-randomly distributed across intrinsic brain networks. Regional effect sizes (herein the β values) quantifying PPS-related deviations were ranked to generate a reference distribution, from which each region was iteratively assigned a running enrichment score (ES) based on its network affiliation. Brain deviations phenotyped by CT and SA have been modeled separately. The maximum deviation from zero in this cumulative scoring process defined the ES for each network, quantifying the degree of spatial concentration of PPS-associated deviations. Statistical significance was evaluated through 5,000 permutation tests, wherein regional PPS associations were randomly reassigned across the brain to generate a null distribution of ES values. The observed ES was compared against this null distribution, and multiple comparisons for seven networks were corrected using the Benjamini-Hochberg false discovery rate (FDR) correction ($q < 0.05$, one-side). Results demonstrated statistically significant enrichment into the default mode network when brain deviations were phenotyped by the CT (**$q < .05$; Table S15-16)**.

| Brain network | Enrichment Scores | P value | BH Q value |
|---|---|---|---|
| Ventral Attention | -0.375 | 0.0485 | 0.084875 |
| Limbic | 0.238 | 0.258 | 0.258 |
| Frontoparietal | -0.478 | 0.064 | 0.0896 |
| Visual | 0.406 | 0.0445 | 0.084875 |
| Dorsal Attention | -0.392 | 0.1415 | 0.1650 |
| **Default Mode** | **0.521** | **0.004** | **0.028** |
| Somatomotor | -0.508 | 0.0155 | 0.05425 |

**Table S15. Network enrichment of brain CT deviation-PPS mappings.** Enrichment score reaching statistical significance for a given network is bold.

| Brain network | Enrichment Scores | P value | BH Q value |
|---|---|---|---|
| Ventral Attention | 0.292 | 0.1565 | 0.4545 |
| Limbic | -0.399 | 0.0205 | 0.1435 |
| Frontoparietal | 0.296 | 0.3255 | 0.4545 |
| Visual | 0.214 | 0.3685 | 0.4545 |
| Dorsal Attention | -0.220 | 0.4545 | 0.4545 |
| Default Mode | -0.198 | 0.3475 | 0.4545 |
| Somatomotor | 0.200 | 0.441 | 0.4545 |

**Table S16. Network enrichment of brain SA/SV deviation-PPS mappings.** Enrichment score reaching statistical significance for a given network is bold.

For examining specificity of this network enrichment, based on the unified triple brain network theory (Chen & Feng, 2022), we modeled deviations-PPS mappings from triple brain networks identified to represent procrastination trait. Brain regions derived from this theoretical model, including dorsolateral prefrontal cortex (labeled rostral middle frontal in D-K atlas), anterior cingulate cortex (labeled rostral anterior cingulate in D-K atlas), insular, orbital frontal cortex (labeled lateral orbitofrontal in D-K atlas), ventromedial prefrontal cortex (labeled medial orbitofrontal in D-K atlas), ventral striatum (labeled Nucleus accumbens in Aseg atlas) and parahippocampus (labeled Hippocampus in Aseg atlas). Despite some regional deviation-PPS associations, no network-wise enrichment reached statistical significance ($q < 0.05$) **(Table S17).**

| Brain Regions | CT | | SA/SV | |
|---|---|---|---|---|
| | R value | BH Q value | R value | BH Q value |
| L_lateralorbitofrontal | 0.09 | 0.27 | -0.078 | 0.364 |
| L_medialorbitofrontal | 0.04 | 0.61 | 0.07 | 0.413 |
| L_rostralanteriorcingulate | -0.03 | 0.75 | 0.007 | 0.935 |
| L_rostralmiddlefrontal | 0.15 | 0.08 | -0.016 | 0.85 |
| L_insula | 0.09 | 0.31 | 0.055 | 0.519 |
| R_lateralorbitofrontal | -0.06 | 0.52 | 0.007 | 0.939 |
| R_medialorbitofrontal | 0.12 | 0.17 | 0.008 | 0.921 |
| R_rostralanteriorcingulate | 0 | 0.97 | -0.061 | 0.477 |
| R_rostralmiddlefrontal | 0.02 | 0.8 | 0.022 | 0.8 |
| R_insula | 0.07 | 0.44 | 0.018 | 0.831 |
| Laccumb | | | 0.172 | 0.043 |
| Lhippo | | | 0.151 | 0.077 |
| Raccumb | | | 0.212 | 0.012 |
| Rhippo | | | 0.154 | 0.07 |

**Table S17. Theory-driven deviation-PPS associations.**

**6. Representation Similarity Patterns of Brain Deviations to PPS**

We capitalized on the representation similarity analysis (IS-RSA) to decode the multivariate morphological deviation patterns underlying PPS. As described above, these individual deviation patterns were vectorized into whole-brain morphological deviation maps (70 participant × 68/14 cortical/subcortical parcels), and inter-subject correlations were estimated to generate a 70 × 70 neural representation dissimilarity matrix (nRDM) for each morphometric features (CT, SA, SV) by subtracting from 1. For the regional RSA, the Euclidean distance was used to phenotype intersubject nRDM, region-by-region. By iterating each regional nRDM to correlate with the behavioral representation dissimilarity matrix (bRDM) derived from PPS phenotyping, we obtained representation similarity (RS, positive r values) and representation dissimilarity (RDS, negative r values), with statistical significance determined at $q < .05$ using Benjamini-Hochberg correction. Thus, for the whole-brain deviation mappings, 1 × 2,415 (70 × 67 / 2) nRDM was correlated with 1 x 2,415 bRDM ($r = 0.03$, *95% CI* = [0.01 - 0.05], $q = .006$) when the brain deviations were phenotyped by CT. SA-derived and SV-derived nRDMs were found positively correlated with bRDM of PPS, respectively ($r_{SA} = 0.02$; 95% *CI* = [0.01 - 0.04], $q = .04$; $r_{sv} = 0.03$; 95% CI = [0.01 - 0.05], $q = .006$). For regional representation patterns, these results have been tabulated into **Table S18-20**.

| Brain Parcels | RS values | P value | BH Q value |
|---|---|---|---|

| | | | |
|---|---|---|---|
| **L_bankssts** | **0.042305002** | **2.29E-05** | **0.000155974** |
| **L_caudalanteriorcingulate** | **0.039601988** | **7.39E-05** | **0.000358839** |
| **R_lateraloccipital** | **0.106712546** | **0** | **0** |
| **L_parsopercularis** | **0.093409504** | **0** | **0** |
| **R_precuneus** | **0.088284706** | **0** | **0** |
| **L_parstriangularis** | **0.085523412** | **0** | **0** |
| **L_superiortemporal** | **0.047079939** | **2.45E-06** | **2.08E-05** |
| **L_insula** | **0.041112434** | **3.88E-05** | **0.000239628** |
| **R_middletemporal** | **0.038112842** | **0.000136575** | **0.000619142** |
| **R_lingual** | **0.035061572** | **0.000450259** | **0.001913601** |
| **L_superiorfrontal** | **0.034660295** | **0.000523286** | **0.002093144** |
| **L_parahippocampal** | **0.033600259** | **0.000772673** | **0.002918987** |
| **L_lateraloccipital** | **0.031479943** | **0.001632075** | **0.005549056** |
| **L_posteriorcingulate** | **0.026030626** | **0.00919789** | **0.023165055** |
| **L_postcentral** | **0.025285563** | **0.011405108** | **0.02769812** |
| **R_medialorbitofrontal** | **0.025070512** | **0.012124131** | **0.028428996** |
| L_caudalmiddlefrontal | 0.020635071 | 0.038960512 | 0.082791088 |
| R_inferiorparietal | 0.020448142 | 0.040766512 | 0.084003721 |
| L_temporalpole | 0.018762874 | 0.060483853 | 0.114247279 |
| R_temporalpole | 0.017377152 | 0.082107818 | 0.146929779 |
| L_entorhinal | 0.015999157 | 0.109443823 | 0.190825126 |
| R_superiortemporal | 0.015546061 | 0.119860203 | 0.198792532 |
| R_supramarginal | 0.015270427 | 0.126567005 | 0.204918009 |
| L_pericalcarine | 0.009334214 | 0.350387252 | 0.489952747 |
| L_transversetemporal | 0.00928267 | 0.353054185 | 0.489952747 |
| R_parahippocampal | 0.008803971 | 0.378432457 | 0.514668141 |
| L_precuneus | 0.007117033 | 0.476455674 | 0.602871252 |
| L_middletemporal | 0.006088225 | 0.542466903 | 0.65870981 |
| R_posteriorcingulate | 0.005345696 | 0.592787741 | 0.707185375 |
| R_insula | 0.003478445 | 0.727845195 | 0.836567279 |
| L_fusiform | 0.003341645 | 0.738147599 | 0.836567279 |
| R_inferiortemporal | 0.00250781 | 0.801900775 | 0.885446256 |
| L_lingual | 0.00113387 | 0.909685499 | 0.937251727 |
| L_isthmuscingulate | 0.000691507 | 0.944846232 | 0.944846232 |
| L_precentral | -0.000913149 | 0.927211523 | 0.941050501 |
| R_caudalanteriorcingulate | -0.001936537 | 0.846382451 | 0.885446256 |
| R_parstriangularis | -0.002052811 | 0.837284212 | 0.885446256 |
| R_rostralanteriorcingulate | -0.002161148 | 0.82882646 | 0.885446256 |
| R_cuneus | -0.002420608 | 0.80865311 | 0.885446256 |
| L_superiorparietal | -0.00423647 | 0.671690595 | 0.787499318 |
| R_parsorbitalis | -0.006732304 | 0.500613946 | 0.618940878 |
| L_cuneus | -0.007080036 | 0.4787507 | 0.602871252 |
| L_inferiorparietal | -0.007605026 | 0.446754607 | 0.584217563 |
| L_rostralanteriorcingulate | -0.008004211 | 0.423262494 | 0.564349992 |

| | | | |
|---|---|---|---|
| R_fusiform | -0.009775391 | 0.328085275 | 0.474676569 |
| R_superiorfrontal | -0.01089076 | 0.275902135 | 0.40785533 |
| R_parsopercularis | -0.011473182 | 0.251033193 | 0.379339047 |
| R_precentral | -0.012994466 | 0.193582577 | 0.299173073 |
| L_paracentral | -0.013858469 | 0.165593581 | 0.261868919 |
| R_isthmuscingulate | -0.015718522 | 0.115807481 | 0.196872717 |
| R_postcentral | -0.017453172 | 0.080777636 | 0.146929779 |
| R_transversetemporal | -0.018921148 | 0.058346135 | 0.113358206 |
| L_medialorbitofrontal | -0.019943711 | 0.045997424 | 0.091994847 |
| R_frontalpole | -0.021456289 | 0.031810184 | 0.069777177 |
| **R_pericalcarine** | **-0.023591482** | **0.018251346** | **0.041369718** |
| **L_supramarginal** | **-0.026524526** | **0.007953284** | **0.020800898** |
| **R_bankssts** | **-0.027177252** | **0.006540399** | **0.017789885** |
| **R_rostralmiddlefrontal** | **-0.027833045** | **0.005352332** | **0.01516494** |
| **R_lateralorbitofrontal** | **-0.028833709** | **0.003911702** | **0.011565031** |
| **R_caudalmiddlefrontal** | **-0.030323598** | **0.002410578** | **0.007450877** |
| **R_paracentral** | **-0.030752201** | **0.002089168** | **0.006764926** |
| **L_frontalpole** | **-0.032179219** | **0.001281356** | **0.004585905** |
| **L_parsorbitalis** | **-0.039610801** | **7.36E-05** | **0.000358839** |
| **R_entorhinal** | **-0.040434027** | **5.19E-05** | **0.000294259** |
| **L_rostralmiddlefrontal** | **-0.043254675** | **1.50E-05** | **0.000113003** |
| **R_superiorparietal** | **-0.061612629** | **6.84E-10** | **6.64E-09** |
| **L_lateralorbitofrontal** | **-0.064136554** | **1.34E-10** | **1.51E-09** |
| **L_inferiortemporal** | **-0.095979815** | **0** | **0** |

**Table S18 Representation similarity from regional multivariate neural representation dissimilarity matrix (nRDM) phenotyped by CT to behavioral dissimilarity matrix.** Regional representation patterns reaching statistical significance ($q < .05$) were in bold.The presentation of these brain regions was reordered from similarity to dissimilarity.

| Brain Parcels | RS values | P value | BH Q value |
|---|---|---|---|
| **R_fusiform** | **0.108956424** | **0** | **0** |
| **R_superiorfrontal** | **0.10360051** | **0** | **0** |
| **L_lateraloccipital** | **0.099931742** | **0** | **0** |
| **R_temporalpole** | **0.098076845** | **0** | **0** |
| **R_superiorparietal** | **0.0898476** | **0** | **0** |
| **R_middletemporal** | **0.086859238** | **0** | **0** |
| **L_middletemporal** | **0.070653064** | **1.47E-12** | **1.43E-11** |
| **R_rostralmiddlefrontal** | **0.068973136** | **4.90E-12** | **4.16E-11** |
| **R_parsorbitalis** | **0.068697305** | **5.95E-12** | **4.50E-11** |
| **L_posteriorcingulate** | **0.067782475** | **1.13E-11** | **7.68E-11** |
| **L_superiortemporal** | **0.063000438** | **2.81E-10** | **1.74E-09** |
| **R_lateraloccipital** | **0.062212184** | **4.67E-10** | **2.64E-09** |

| | | | |
|---|---|---|---|
| **R_precuneus** | **0.058687574** | **4.19E-09** | **2.19E-08** |
| **L_parsopercularis** | **0.057959718** | **6.50E-09** | **3.16E-08** |
| **L_paracentral** | **0.051837937** | **2.11E-07** | **8.96E-07** |
| **L_parstriangularis** | **0.050606538** | **4.06E-07** | **1.62E-06** |
| **R_postcentral** | **0.049090277** | **8.92E-07** | **3.37E-06** |
| **R_precentral** | **0.045983862** | **4.17E-06** | **1.42E-05** |
| **L_inferiortemporal** | **0.045285753** | **5.82E-06** | **1.80E-05** |
| **R_parstriangularis** | **0.041521279** | **3.24E-05** | **9.59E-05** |
| **L_superiorparietal** | **0.039347746** | **8.22E-05** | **0.000223512** |
| **L_temporalpole** | **0.038888065** | **9.94E-05** | **0.000260099** |
| **L_entorhinal** | **0.036021338** | **0.000312352** | **0.000786665** |
| **R_superiortemporal** | **0.03578887** | **0.000341555** | **0.000829491** |
| **L_precuneus** | **0.032295067** | **0.001230468** | **0.00278906** |
| **R_entorhinal** | **0.031183564** | **0.00180581** | **0.003911196** |
| **R_transversetemporal** | **0.031127441** | **0.001840563** | **0.003911196** |
| **L_lingual** | **0.030967076** | **0.001943281** | **0.004004337** |
| **R_caudalmiddlefrontal** | **0.027779983** | **0.005440661** | **0.010570426** |
| **L_precentral** | **0.026255585** | **0.00861092** | **0.016102433** |
| **L_parsorbitalis** | **0.023758049** | **0.017446996** | **0.031220941** |
| **L_isthmuscingulate** | **0.022901241** | **0.021940424** | **0.038255099** |
| R_paracentral | 0.0153239 | 0.125243546 | 0.181203428 |
| L_rostralmiddlefrontal | 0.011676573 | 0.242728806 | 0.336848139 |
| L_supramarginal | 0.009810658 | 0.326343114 | 0.443826635 |
| R_cuneus | 0.0095777 | 0.337962445 | 0.450616594 |
| L_frontalpole | 0.009082984 | 0.363506873 | 0.475355141 |
| R_isthmuscingulate | 0.007737967 | 0.438849728 | 0.563052482 |
| L_caudalmiddlefrontal | 0.005643344 | 0.572360374 | 0.682586091 |
| L_superiorfrontal | 0.004383555 | 0.660992102 | 0.724959079 |
| R_inferiortemporal | 0.004110146 | 0.680932493 | 0.734974754 |
| L_cuneus | -8.11E-05 | 0.993528978 | 0.993528978 |
| R_supramarginal | -0.001240271 | 0.901252015 | 0.914703538 |
| L_medialorbitofrontal | -0.001586592 | 0.873884175 | 0.900365513 |
| R_medialorbitofrontal | -0.001926317 | 0.847183203 | 0.886283966 |
| L_bankssts | -0.003019213 | 0.762613761 | 0.810277121 |
| L_pericalcarine | -0.00473637 | 0.635613375 | 0.708552614 |
| L_fusiform | -0.00520897 | 0.602281845 | 0.682586091 |
| L_rostralanteriorcingulate | -0.005261552 | 0.598622565 | 0.682586091 |
| L_lateralorbitofrontal | -0.005295827 | 0.596242663 | 0.682586091 |
| L_caudalanteriorcingulate | -0.005657285 | 0.571411841 | 0.682586091 |
| R_inferiorparietal | -0.006730825 | 0.500708027 | 0.619057197 |
| R_parsopercularis | -0.00752126 | 0.451776731 | 0.568904032 |
| L_parahippocampal | -0.013938399 | 0.163166714 | 0.231152845 |
| L_transversetemporal | -0.015383583 | 0.123779155 | 0.181203428 |
| R_lateralorbitofrontal | -0.0154817 | 0.121400794 | 0.181203428 |

| | | | |
|---|---|---|---|
| R_parahippocampal | -0.016331888 | 0.102260597 | 0.158039104 |
| L_insula | -0.018427441 | 0.065229254 | 0.103153239 |
| L_inferiorparietal | -0.020640613 | 0.038908023 | 0.062993942 |
| R_insula | -0.021150121 | 0.034332264 | 0.056941316 |
| **R_bankssts** | **-0.021811447** | **0.029085106** | **0.049444681** |
| **L_postcentral** | **-0.026196552** | **0.008761618** | **0.016102433** |
| **R_frontalpole** | **-0.030030027** | **0.00265619** | **0.005312379** |
| **R_rostralanteriorcingulate** | **-0.033998948** | **0.000668156** | **0.001566711** |
| **R_pericalcarine** | **-0.040918599** | **4.22E-05** | **0.000119453** |
| **R_caudalanteriorcingulate** | **-0.045336815** | **5.68E-06** | **1.80E-05** |
| **R_posteriorcingulate** | **-0.047843439** | **1.68E-06** | **6.00E-06** |
| **R_lingual** | **-0.054415197** | **5.10E-08** | **2.31E-07** |

**Table S19 Representation similarity from regional multivariate neural representation dissimilarity matrix (nRDM) phenotyped by SA to behavioral dissimilarity matrix.** Regional representation patterns reaching statistical significance ($q < .05$) were in bold.The presentation of these brain regions was reordered from similarity to dissimilarity.

| Brain Parcels | RS values | P value | BH Q value |
|---|---|---|---|
| **Lpal** | **-0.085372211** | **0** | **0** |
| **Lput** | **-0.056024434** | **2.03E-08** | **1.25E-07** |
| **Rput** | **-0.055550474** | **2.67E-08** | **1.25E-07** |
| **Rpal** | **-0.045588121** | **5.04E-06** | **1.41E-05** |
| **Ramyg** | **-0.032055952** | **0.00133763** | **0.003121137** |
| **Lamyg** | **-0.024149502** | **0.0156779** | **0.027436325** |
| Lcaud | -0.017551292 | 0.079086672 | 0.123023711 |
| Lthal | -0.016793812 | 0.092915831 | 0.130082163 |
| Rhippo | -0.005209802 | 0.602223909 | 0.694690782 |
| Rthal | 0.002234215 | 0.823133401 | 0.823133401 |
| Laccumb | 0.004604237 | 0.645070012 | 0.694690782 |
| Rcaud | 0.008726132 | 0.382662749 | 0.487025317 |
| **Lhippo** | **0.027933623** | **0.005188454** | **0.010376907** |
| **Raccumb** | **0.052246724** | **1.69E-07** | **5.92E-07** |

**Table S20 Representation similarity from regional multivariate neural representation dissimilarity matrix (nRDM) phenotyped by SV to behavioral dissimilarity matrix.** Regional representation patterns reaching statistical significance ($q < .05$) were in bold.The presentation of these brain regions was reordered from similarity to dissimilarity.

However, no statistical enrichment was found from these multivariate brain deviation patterns in any known network parceled by the Yeo-7 network **(Table S21-22)**.

| Brain network | Enrichment Scores | P value | BH Q value |
|---|---|---|---|
| Ventral Attention | 0.339285714285714 | 0.158 | 0.751333333 |
| Limbic | 0.125 | 0.984 | 0.984 |
| Frontoparietal | -0.376344086 | 0.322 | 0.751333333 |
| Visual | 0.213793103448276 | 0.732 | 0.984 |
| Dorsal Attention | -0.274193548 | 0.725 | 0.984 |
| Default Mode | 0.153439153439153 | 0.897 | 0.984 |
| Somatomotor | -0.358333333 | 0.265 | 0.75133333333 |

**Table S21. Network enrichment of regional CT deviation-PPS multivariate mappings.** Enrichment score reaching statistical significance for a given network is in bold.

| Brain network | Enrichment Scores | P value | BH Q value |
|---|---|---|---|
| Ventral Attention | 0.208333333333333 | 0.71 | 0.973 |
| Limbic | -0.452380952 | 0.019 | 0.133 |
| Frontoparietal | -0.268817204 | 0.725 | 0.973 |
| Visual | -0.165517241 | 0.945 | 0.973 |
| Dorsal Attention | -0.182795699 | 0.973 | 0.973 |
| Default Mode | 0.335978835978836 | 0.135 | 0.4725 |
| Somatomotor | 0.3 | 0.435 | 0.973 |

**Table S22. Network enrichment of regional SA deviation-PPS multivariate mappings.** Enrichment score reaching statistical significance for a given network is in bold.

**7. Neurobiological Mappings from Neural Representation Patterns of PPS**

To interpret the neurobiological significance of the representation similarity (RS) patterns derived from cortical thickness (CT) and surface area (SA) deviations, we aligned the neural representation maps generated from RSA into the Neuromap framework (Markello et al., 2022). This involved transforming the whole-brain deviation maps into standardized coordinate systems compatible with existing neurobiological atlases. The resulting RS maps were spatially correlated with over 40 reference neurobiological datasets, including cortical functional gradients, neurotransmitter receptor densities, and molecular gene expression patterns. These maps were processed using Neuromaps' spatial enrichment tool to evaluate the alignment of PPS-related deviation patterns with established biological ontologies. Statistical significance was determined spatial autocorrelation-preserving null model ("spin test"). Alignment of the CT or SA-derived RS maps with Neuromap revealed significant spatial enrichment in several neurobiological features. Notably, PPS-specific deviation patterns in CT were significantly associated with cortical functional gradients, particularly the fourth gradient (G4, $r = 0.20$, $p_{spin} = 0.003$) **(Table S23)**. Furthermore, these patterns in the SA deviations showed strong correlations with key neurotransmitter receptor systems, including the serotonin transporter (5-HTT, r = -

0.34, p < 0.001), dopamine transporter (DAT, r = -0.24, p = 0.01), and dopamine D1 receptor (D1, r = -0.19, p = 0.03) **(Table S24)**.

| R value | $P_{spin}$ value | Source space | Neurobiological map |
|---|---|---|---|
| **0.196516976** | **0.002997003** | **fsLR** | **Functional gradient04** |
| 0.146218013 | 0.058941059 | fsaverage | 5-HTT |
| 0.106184445 | 0.062937063 | fsLR | Functional gradient05 |
| 0.100804327 | 0.111888112 | fsLR | Cerebral blood volume |
| 0.083016261 | 0.156843157 | MNI152 | NET |
| 0.102356017 | 0.161838162 | fsaverage | PC1 gene expression |
| -0.084915251 | 0.164835165 | fsLR | Functional gradient03 |
| -0.152379259 | 0.204795205 | MNI152 | MOR |
| -0.06002821 | 0.282717283 | fsLR | Functional gradient10 |
| -0.058427165 | 0.342657343 | MNI152 | KOR |
| 0.069631886 | 0.350649351 | fsaverage | 5-HT2a |
| 0.069943406 | 0.385614386 | fsLR | Functional gradient02 |
| 0.039181687 | 0.440559441 | fsLR | Functional gradient06 |
| 0.069737237 | 0.488511489 | MNI152 | DAT |
| 0.053994161 | 0.504495504 | fsaverage | 5-HT4 |
| -0.044396102 | 0.525474525 | MNI152 | PC1NeuroSynth |
| -0.031212203 | 0.628371628 | fsLR | Glucose metabolism |
| 0.034196168 | 0.637362637 | fsaverage | 5-HT1b |
| 0.027515913 | 0.709290709 | MNI152 | GABAa |
| -0.030021952 | 0.755244755 | MNI152 | D1 |
| 0.019945649 | 0.761238761 | fsLR | Cortical thickness |
| 0.01524494 | 0.807192807 | fsaverage | 5-HT1a |
| -0.012502133 | 0.828171828 | fsLR | Functional gradient09 |
| -0.010760773 | 0.842157842 | fsLR | Functional gradient08 |
| 0.015402127 | 0.85014985 | fsLR | Oxygen metabolism |
| 0.012783373 | 0.852147852 | MNI152 | D2 |
| 0.009699854 | 0.87012987 | fsLR | Functional gradient07 |
| -0.01200426 | 0.885114885 | fsLR | Functional gradient01 |
| 0.011429689 | 0.892107892 | fsLR | T1w/T2w ratio |
| -0.007622814 | 0.894105894 | fsLR | Cross-species functional homology |
| 0.008038268 | 0.909090909 | fsLR | Cerebral blood flow |
| 0.008625772 | 0.925074925 | fsLR | Intersubject variability |

**Table S23. Neuromap spatial correlation of neural representation patterns curated by CT to source maps.** Enrichment reaching statistical significance for a given neurobiological map is in bold. The presentation of these maps was reordered by spatial correlation strengths.

| R value | $P_{spin}$ value | Source space | Neurobiological map |
|---|---|---|---|
| -0.343647344 | 0.000999001 | fsaverage | 5-HTT |
| -0.24212462 | 0.010989011 | MNI152 | DAT |
| -0.191209696 | 0.031968032 | MNI152 | D1 |

| | | | |
|---|---|---|---|
| -0.110937789 | 0.05994006 | fsLR | Functional gradient07 |
| -0.11210223 | 0.120879121 | MNI152 | GABAa |
| -0.103279941 | 0.151848152 | fsLR | Functional gradient04 |
| -0.088677964 | 0.183816184 | MNI152 | D2 |
| -0.10000158 | 0.217782218 | fsLR | Cerebral blood flow |
| 0.095831178 | 0.293706294 | fsLR | Functional gradient01 |
| -0.105629653 | 0.301698302 | fsaverage | 5-HT1a |
| 0.065163784 | 0.341658342 | fsLR | Functional gradient10 |
| 0.056309144 | 0.34965035 | MNI152 | NET |
| -0.067274563 | 0.358641359 | fsLR | Functional gradient09 |
| -0.083991679 | 0.363636364 | fsLR | Oxygen metabolism |
| 0.057951295 | 0.400599401 | MNI152 | PC1NeuroSynth |
| -0.04943113 | 0.491508492 | fsLR | Cross-species functional homology |
| -0.055723656 | 0.528471528 | fsLR | T1w/T2w ratio |
| 0.054556471 | 0.555444555 | fsLR | Glucose metabolism |
| -0.035123949 | 0.575424575 | fsLR | Cerebral blood volume |
| -0.032890826 | 0.582417582 | MNI152 | KOR |
| 0.044947229 | 0.588411588 | fsLR | Cortical thickness |
| -0.038279444 | 0.591408591 | fsLR | Functional gradient03 |
| -0.063418342 | 0.603396603 | MNI152 | MOR |
| 0.030583085 | 0.646353646 | fsLR | Functional gradient06 |
| -0.042548206 | 0.678321678 | fsaverage | 5-HT4 |
| 0.023293193 | 0.739260739 | fsLR | Functional gradient08 |
| 0.028464315 | 0.791208791 | fsaverage | PC1 gene expression |
| 0.021341015 | 0.845154845 | fsaverage | 5-HT2a |
| 0.017821754 | 0.88011988 | fsLR | Intersubject variability |
| -0.012843532 | 0.909090909 | fsaverage | 5-HT1b |
| -0.001507102 | 0.981018981 | fsLR | Functional gradient05 |
| 0.001424118 | 0.986013986 | fsLR | Functional gradient02 |

**Table S24. Neuromap spatial correlation of neural representation patterns curated by SA to source maps.** Enrichment reaching statistical significance for a given neurobiological map is in bold. The presentation of these maps was reordered by spatial correlation strengths.

**8. Transcriptomic Markers of PPS Phenotyped by SA Deviations**

Given significant enrichment of molecular pathways from the SA-derived deviation representation patterns, we used the PLS model to regress transcriptomic profiles (10,027 genes × 68 brain parcels) to PPS-specific deviation pattern (1 × 68 brain parcels). Results showed that the PLS2 significantly explained this prediction model ($p < .01$, Bootstrapping method). By predefined statistical significance criterion ($q < .0005$, Benjamini-Hochberg correction), we identified 78 (226) genes overexpressed (underexpressed) **(Table S25-26)**.

| Gene Symbol | Entrez ID | Z scores | P values | BH Q values |
|---|---|---|---|---|
| *FRMD8* | 7758 | 5.408269 | 3.18E-08 | 0.00016 |

| | | | | |
|---|---|---|---|---|
| *MPLKIP* | 8806 | 5.112447 | 1.59E-07 | 0.000199 |
| *DEPP1* | 4441 | 5.006578 | 2.77E-07 | 0.000231 |
| *SRPK2* | 2689 | 4.945422 | 3.80E-07 | 0.000272 |
| *SERTAD3* | 5477 | 4.910827 | 4.53E-07 | 0.000284 |
| *AATK* | 3729 | 4.85189 | 6.11E-07 | 0.000307 |
| *MIGA1* | 9724 | 4.752128 | 1.01E-06 | 0.000374 |
| *STARD9* | 6762 | 4.635907 | 1.78E-06 | 0.000457 |
| *PCGF2* | 3045 | 4.628221 | 1.84E-06 | 0.000462 |
| *DEPDC5* | 3765 | 4.579486 | 2.33E-06 | 0.000497 |
| *NEK9* | 8265 | 4.570279 | 2.44E-06 | 0.000498 |
| *CCNYL1* | 8996 | 4.550405 | 2.68E-06 | 0.000537 |
| *RPL24* | 2432 | 4.409902 | 5.17E-06 | 0.000762 |
| *AFTPH* | 6016 | 4.407423 | 5.23E-06 | 0.00076 |
| *CDK5R2* | 3431 | 4.405449 | 5.28E-06 | 0.000745 |
| *TRUB1* | 8863 | 4.391312 | 5.63E-06 | 0.000753 |
| *PAGR1* | 7296 | 4.368861 | 6.24E-06 | 0.000783 |
| *ANKIB1* | 5923 | 4.362326 | 6.43E-06 | 0.000777 |
| *HLF* | 1241 | 4.353163 | 6.71E-06 | 0.000791 |
| *NELFA* | 2997 | 4.349543 | 6.82E-06 | 0.000786 |
| *NPHP4* | 9460 | 4.338711 | 7.17E-06 | 0.000798 |
| *GNPDA2* | 8771 | 4.331577 | 7.40E-06 | 0.000816 |
| *TCF7* | 2775 | 4.309194 | 8.19E-06 | 0.000893 |
| *NPPB* | 1862 | 4.300349 | 8.53E-06 | 0.0009 |
| *NUDT9P1* | 8567 | 4.295021 | 8.73E-06 | 0.000912 |
| *PITPNM2* | 6809 | 4.284281 | 9.17E-06 | 0.000928 |
| *MLYCD* | 4880 | 4.283591 | 9.20E-06 | 0.000922 |
| *JAZF1* | 9358 | 4.267344 | 9.89E-06 | 0.000982 |
| *C2CD5* | 3848 | 4.229956 | 1.17E-05 | 0.001028 |
| *UMODL1-AS1* | 8972 | 4.216461 | 1.24E-05 | 0.001073 |
| *TRIO* | 2895 | 4.212023 | 1.27E-05 | 0.001075 |
| *CAND2* | 4698 | 4.200935 | 1.33E-05 | 0.001111 |
| *CCT6B* | 4286 | 4.183456 | 1.44E-05 | 0.001133 |
| *TARSL2* | 8613 | 4.182883 | 1.44E-05 | 0.001127 |
| *TBC1D1* | 4769 | 4.156837 | 1.61E-05 | 0.001198 |
| *SSFA2* | 2693 | 4.13192 | 1.80E-05 | 0.001261 |
| *ANKS6* | 9275 | 4.123118 | 1.87E-05 | 0.001292 |
| *SP100* | 2668 | 4.078973 | 2.26E-05 | 0.001502 |
| *SLC38A2* | 5907 | 4.076437 | 2.29E-05 | 0.001508 |
| *SYDE2* | 7827 | 4.074478 | 2.31E-05 | 0.001511 |
| *JUND* | 1437 | 4.065785 | 2.39E-05 | 0.001509 |
| *ACTR6* | 7076 | 4.031732 | 2.77E-05 | 0.001652 |
| *ZEB2* | 3842 | 4.028821 | 2.80E-05 | 0.001653 |
| *MED30* | 8178 | 4.024523 | 2.85E-05 | 0.001674 |
| *INTS6L* | 9278 | 4.015626 | 2.96E-05 | 0.001689 |
| *PEX11A* | 3361 | 3.99259 | 3.27E-05 | 0.00181 |
| *SLC10A4* | 9258 | 3.984899 | 3.38E-05 | 0.001849 |

| | | | | |
|---|---|---|---|---|
| *CD34* | 410 | 3.974911 | 3.52E-05 | 0.001878 |
| *G3BP1* | 4003 | 3.971178 | 3.58E-05 | 0.001897 |
| *PRB1* | 2153 | 3.965462 | 3.66E-05 | 0.001903 |
| *ICE2* | 7351 | 3.944885 | 3.99E-05 | 0.002032 |
| *SLC38A3* | 4415 | 3.918004 | 4.46E-05 | 0.002173 |
| *NEURL3* | 8328 | 3.879491 | 5.23E-05 | 0.002385 |
| *ETV5* | 870 | 3.864152 | 5.57E-05 | 0.002506 |
| *FZD5* | 3093 | 3.859498 | 5.68E-05 | 0.002532 |
| *DENND6A* | 9257 | 3.856299 | 5.76E-05 | 0.002531 |
| *PLEKHA1* | 6928 | 3.856197 | 5.76E-05 | 0.002521 |
| *CETN3* | 474 | 3.851466 | 5.87E-05 | 0.002537 |
| *EIF1B* | 4077 | 3.821735 | 6.63E-05 | 0.002657 |
| *IPO5* | 1501 | 3.820305 | 6.66E-05 | 0.002662 |
| *IDI1* | 1335 | 3.819424 | 6.69E-05 | 0.002661 |
| *KIF19* | 8633 | 3.819248 | 6.69E-05 | 0.002653 |
| *PHF2* | 2023 | 3.81564 | 6.79E-05 | 0.002671 |
| *TOGARAM2* | 9140 | 3.807103 | 7.03E-05 | 0.002743 |
| *PCDHAC2* | 6518 | 3.798898 | 7.27E-05 | 0.002771 |
| *FNBP4* | 4846 | 3.793124 | 7.44E-05 | 0.002814 |
| *SEC24D* | 3863 | 3.791271 | 7.49E-05 | 0.002814 |
| *C7* | 320 | 3.779519 | 7.86E-05 | 0.002854 |
| *UBE2B* | 2932 | 3.779108 | 7.87E-05 | 0.002849 |
| *MKRN3* | 3040 | 3.777122 | 7.93E-05 | 0.002841 |
| *ARIH2* | 4142 | 3.769368 | 8.18E-05 | 0.002879 |
| *HEMK1* | 5734 | 3.762094 | 8.42E-05 | 0.002933 |
| *AQP6* | 159 | 3.759504 | 8.51E-05 | 0.002953 |
| *SRRT* | 5788 | 3.757059 | 8.60E-05 | 0.002972 |
| *RTF1* | 4743 | 3.752378 | 8.76E-05 | 0.003018 |
| *ATP6V1G1* | 3696 | 3.740324 | 9.19E-05 | 0.003134 |
| *ATP9B* | 9717 | 3.736904 | 9.32E-05 | 0.003166 |
| *UBE2E2* | 2937 | 3.736644 | 9.32E-05 | 0.003159 |

**Table S25. Gene list for statistically significant associations with PPS-specific deviation patterns in the PLS2+ component.** P values were estimated by using two-sided Z sampling distribution test without corrections. Q values presented corrected P values after the Benjamini-Hochberg method.

| Gene Symbol | Entrez ID | Z scores | P values | BH Q values |
|---|---|---|---|---|
| *CSTB* | 616 | -5.75755 | 4.27E-09 | 4.28E-05 |
| *SCRN1* | 3826 | -5.40566 | 3.23E-08 | 0.000108 |
| *C9orf16* | 7256 | -5.27351 | 6.69E-08 | 0.000168 |
| *PSME3* | 4029 | -5.24079 | 7.99E-08 | 0.00016 |
| *CEP19* | 8074 | -5.20039 | 9.94E-08 | 0.000166 |
| *SELENON* | 6686 | -5.18893 | 1.06E-07 | 0.000151 |
| *MANEAL* | 8961 | -5.10793 | 1.63E-07 | 0.000181 |
| *YIF1B* | 8187 | -5.05567 | 2.14E-07 | 0.000215 |

| | | | | |
|---|---|---|---|---|
| *PPP5C* | 2151 | -5.03732 | 2.36E-07 | 0.000215 |
| *EHD3* | 5513 | -4.98454 | 3.11E-07 | 0.00024 |
| *COG1* | 3611 | -4.92999 | 4.11E-07 | 0.000275 |
| *MT1A* | 1710 | -4.89569 | 4.90E-07 | 0.000289 |
| *POLR2E* | 2099 | -4.88374 | 5.20E-07 | 0.00029 |
| *RAB11FIP2* | 4589 | -4.86554 | 5.71E-07 | 0.000301 |
| *MARK4* | 6872 | -4.84726 | 6.26E-07 | 0.000299 |
| *DCHS1* | 3296 | -4.83375 | 6.70E-07 | 0.000305 |
| *FAM160B1* | 6855 | -4.83255 | 6.74E-07 | 0.000294 |
| *CNOT1* | 4675 | -4.81724 | 7.28E-07 | 0.000304 |
| *PDE9A* | 1973 | -4.79452 | 8.15E-07 | 0.000327 |
| *VAC14* | 6384 | -4.75509 | 9.92E-07 | 0.000382 |
| *RNF208* | 9976 | -4.74688 | 1.03E-06 | 0.00037 |
| *AMIGO1* | 6731 | -4.74688 | 1.03E-06 | 0.000357 |
| *CDH4* | 437 | -4.74125 | 1.06E-06 | 0.000355 |
| *RITA1* | 8050 | -4.70433 | 1.27E-06 | 0.000412 |
| *AGPAT5* | 6271 | -4.69528 | 1.33E-06 | 0.000417 |
| *MLLT11* | 4397 | -4.69171 | 1.35E-06 | 0.000412 |
| *KRTCAP2* | 9228 | -4.6898 | 1.37E-06 | 0.000403 |
| *HBA2* | 1206 | -4.68512 | 1.40E-06 | 0.000401 |
| *WNK2* | 7182 | -4.67286 | 1.49E-06 | 0.000414 |
| *FMNL1* | 325 | -4.66486 | 1.54E-06 | 0.000418 |
| *DDX51* | 9604 | -4.64935 | 1.66E-06 | 0.000439 |
| *INTS14* | 7645 | -4.60903 | 2.02E-06 | 0.000495 |
| *SLC1A4* | 2580 | -4.60444 | 2.07E-06 | 0.000494 |
| *MICU1* | 4110 | -4.60319 | 2.08E-06 | 0.000485 |
| *RASSF7* | 3123 | -4.60054 | 2.11E-06 | 0.00048 |
| *ARL2* | 180 | -4.58184 | 2.30E-06 | 0.000513 |
| *DSTYK* | 5024 | -4.58109 | 2.31E-06 | 0.000504 |
| *MAP6* | 1610 | -4.577 | 2.36E-06 | 0.000493 |
| *HBA1* | 1205 | -4.5448 | 2.75E-06 | 0.000541 |
| *FBXL18* | 7499 | -4.53367 | 2.90E-06 | 0.000559 |
| *SSSCA1* | 4201 | -4.52952 | 2.96E-06 | 0.000559 |
| *DEF8* | 6030 | -4.52827 | 2.97E-06 | 0.000552 |
| *CYFIP2* | 5253 | -4.49689 | 3.45E-06 | 0.000629 |
| *MRPL49* | 322 | -4.48569 | 3.63E-06 | 0.000651 |
| *CCAR2* | 6877 | -4.46846 | 3.94E-06 | 0.000693 |
| *NUTF2* | 4033 | -4.46301 | 4.04E-06 | 0.000699 |
| *LIMD2* | 7604 | -4.46094 | 4.08E-06 | 0.000693 |
| *GLT8D1* | 6455 | -4.45785 | 4.14E-06 | 0.000692 |
| *FOXP4* | 8500 | -4.45495 | 4.20E-06 | 0.00069 |
| *ADGRL1* | 4599 | -4.45097 | 4.27E-06 | 0.000691 |
| *LOC642852* | 9937 | -4.43423 | 4.62E-06 | 0.000735 |
| *CAPRIN2* | 7186 | -4.43239 | 4.66E-06 | 0.00073 |
| *NFS1* | 3467 | -4.42895 | 4.73E-06 | 0.00073 |
| *NDUFA9* | 1782 | -4.41164 | 5.13E-06 | 0.000779 |
| *COA6* | 9785 | -4.40999 | 5.17E-06 | 0.000774 |

| | | | | |
|---|---|---|---|---|
| *PAFAH1B3* | 1925 | -4.40613 | 5.26E-06 | 0.000754 |
| *PSMB10* | 2234 | -4.39362 | 5.57E-06 | 0.000776 |
| *OAZ2* | 1892 | -4.39308 | 5.59E-06 | 0.000768 |
| *ACACA* | 13 | -4.39272 | 5.60E-06 | 0.000758 |
| *BCAN* | 6972 | -4.37989 | 5.94E-06 | 0.000783 |
| *RILPL1* | 9698 | -4.37577 | 6.05E-06 | 0.000788 |
| *MCHR1* | 1112 | -4.37225 | 6.15E-06 | 0.00079 |
| *STX4* | 2717 | -4.36977 | 6.22E-06 | 0.000789 |
| *BOP1* | 4787 | -4.36691 | 6.30E-06 | 0.00078 |
| *GRAMD1A* | 6834 | -4.36415 | 6.38E-06 | 0.00078 |
| *IDS* | 1336 | -4.35992 | 6.51E-06 | 0.000777 |
| *METRN* | 7217 | -4.34956 | 6.82E-06 | 0.000795 |
| *C11orf1* | 7103 | -4.34847 | 6.85E-06 | 0.000781 |
| *SOS1* | 2654 | -4.34763 | 6.88E-06 | 0.000775 |
| *C16orf70* | 7556 | -4.30268 | 8.44E-06 | 0.00091 |
| *MPZL1* | 3451 | -4.30139 | 8.49E-06 | 0.000905 |
| *TRIM4* | 8133 | -4.29125 | 8.88E-06 | 0.000918 |
| *GRK6* | 1121 | -4.28477 | 9.15E-06 | 0.000936 |
| *SPIRE1* | 6586 | -4.26726 | 9.89E-06 | 0.000973 |
| *AHCYL2* | 4859 | -4.25737 | 1.03E-05 | 0.001007 |
| *SPTAN1* | 2676 | -4.25632 | 1.04E-05 | 0.001002 |
| *LRRC61* | 7197 | -4.25083 | 1.06E-05 | 0.001017 |
| *AP2S1* | 514 | -4.25082 | 1.06E-05 | 0.001007 |
| *GTF2F1* | 1179 | -4.25008 | 1.07E-05 | 0.001001 |
| *NDUFS4* | 1793 | -4.24716 | 1.08E-05 | 0.001005 |
| *FAM213B* | 8681 | -4.24619 | 1.09E-05 | 0.001 |
| *PLD3* | 4983 | -4.24034 | 1.12E-05 | 0.001017 |
| *RPL13A* | 4927 | -4.23296 | 1.15E-05 | 0.001042 |
| *TMEM199* | 8919 | -4.23222 | 1.16E-05 | 0.001036 |
| *LRP1* | 1563 | -4.23148 | 1.16E-05 | 0.00103 |
| *ATF5* | 4574 | -4.21794 | 1.23E-05 | 0.001075 |
| *ARF3* | 164 | -4.21497 | 1.25E-05 | 0.00107 |
| *MALL* | 3091 | -4.21111 | 1.27E-05 | 0.001071 |
| *SF3A3* | 4387 | -4.19621 | 1.36E-05 | 0.001125 |
| *KCNQ2* | 1478 | -4.19106 | 1.39E-05 | 0.001141 |
| *TTPAL* | 7285 | -4.19052 | 1.39E-05 | 0.001134 |
| *STXBP1* | 2718 | -4.1893 | 1.40E-05 | 0.001131 |
| *LRRN2* | 4154 | -4.18897 | 1.40E-05 | 0.001124 |
| *STYXL1* | 5809 | -4.18427 | 1.43E-05 | 0.001138 |
| *SYT3* | 7865 | -4.17791 | 1.47E-05 | 0.001143 |
| *ILF2* | 1375 | -4.16995 | 1.52E-05 | 0.001175 |
| *PIGX* | 6087 | -4.16583 | 1.55E-05 | 0.001187 |
| *FAHD1* | 7697 | -4.16451 | 1.56E-05 | 0.001185 |
| *MYH3* | 1745 | -4.16113 | 1.58E-05 | 0.001194 |
| *SAMD15* | 9104 | -4.15932 | 1.60E-05 | 0.001194 |
| *MEST* | 1655 | -4.15495 | 1.63E-05 | 0.001199 |
| *NOS1AP* | 3784 | -4.14936 | 1.67E-05 | 0.00122 |

| | | | | |
|---|---|---|---|---|
| *PCBD1* | 1944 | -4.14085 | 1.73E-05 | 0.001257 |
| *RNFT2* | 8034 | -4.14033 | 1.73E-05 | 0.001251 |
| *PCK2* | 1950 | -4.13715 | 1.76E-05 | 0.001259 |
| *ZER1* | 4152 | -4.13292 | 1.79E-05 | 0.001274 |
| *SORBS3* | 4019 | -4.13223 | 1.80E-05 | 0.001268 |
| *ZNF599* | 8942 | -4.13018 | 1.81E-05 | 0.001262 |
| *RPL7L1* | 9579 | -4.10261 | 2.04E-05 | 0.001403 |
| *DARS* | 665 | -4.10196 | 2.05E-05 | 0.001397 |
| *NDFIP1* | 7600 | -4.10121 | 2.05E-05 | 0.001392 |
| *CABYR* | 5199 | -4.09836 | 2.08E-05 | 0.0014 |
| *NRSN2* | 7496 | -4.08824 | 2.17E-05 | 0.001453 |
| *DYNLRB1* | 7736 | -4.07227 | 2.33E-05 | 0.001516 |
| *DCUN1D5* | 7866 | -4.07107 | 2.34E-05 | 0.001514 |
| *DLG4* | 708 | -4.06977 | 2.35E-05 | 0.001512 |
| *ELOVL6* | 7241 | -4.06905 | 2.36E-05 | 0.001507 |
| *EEF1A2* | 782 | -4.06631 | 2.39E-05 | 0.001516 |
| *PSMD7* | 2245 | -4.05866 | 2.47E-05 | 0.001547 |
| *FAM174B* | 9829 | -4.05791 | 2.48E-05 | 0.001542 |
| *TMSB10* | 3515 | -4.05607 | 2.50E-05 | 0.001544 |
| *CFD* | 689 | -4.0557 | 2.50E-05 | 0.001537 |
| *LEMD2* | 9351 | -4.05023 | 2.56E-05 | 0.001564 |
| *EPS8L2* | 7110 | -4.04705 | 2.59E-05 | 0.001576 |
| *PWWP2B* | 9172 | -4.0469 | 2.60E-05 | 0.001567 |
| *RFXANK* | 3288 | -4.03875 | 2.69E-05 | 0.001613 |
| *RAB12* | 9252 | -4.03073 | 2.78E-05 | 0.00165 |
| *NEUROD2* | 1812 | -4.02211 | 2.88E-05 | 0.001681 |
| *PTPRS* | 2288 | -4.02199 | 2.89E-05 | 0.001672 |
| *YWHAB* | 3009 | -4.01737 | 2.94E-05 | 0.001696 |
| *MAP4K2* | 2313 | -4.01639 | 2.95E-05 | 0.001693 |
| *PDK2* | 1984 | -4.01507 | 2.97E-05 | 0.001683 |
| *S100A1* | 2493 | -4.00603 | 3.09E-05 | 0.001739 |
| *RAMP1* | 4068 | -4.00341 | 3.12E-05 | 0.001749 |
| *GINM1* | 8515 | -3.99457 | 3.24E-05 | 0.001805 |
| *TNFAIP8L3* | 9766 | -3.98697 | 3.35E-05 | 0.001844 |
| *PIGF* | 2036 | -3.98071 | 3.44E-05 | 0.001872 |
| *GRIN2B* | 1144 | -3.98014 | 3.44E-05 | 0.001867 |
| *THY1* | 2826 | -3.97805 | 3.47E-05 | 0.001873 |
| *POLR1A* | 5071 | -3.97775 | 3.48E-05 | 0.001865 |
| *ADCY5* | 50 | -3.96977 | 3.60E-05 | 0.001898 |
| *PTPRU* | 3966 | -3.9697 | 3.60E-05 | 0.001889 |
| *THOP1* | 2823 | -3.96949 | 3.60E-05 | 0.001881 |
| *FABP3* | 889 | -3.95276 | 3.86E-05 | 0.001996 |
| *KMT5A* | 9762 | -3.95258 | 3.87E-05 | 0.001988 |
| *SNCG* | 2641 | -3.94868 | 3.93E-05 | 0.00201 |
| *RTL8C* | 3424 | -3.93562 | 4.15E-05 | 0.002101 |
| *C9orf116* | 8819 | -3.93106 | 4.23E-05 | 0.002131 |
| *TRADD* | 3328 | -3.92421 | 4.35E-05 | 0.002181 |

| | | | | |
|---|---|---|---|---|
| *PPDPF* | 7268 | -3.92269 | 4.38E-05 | 0.002184 |
| *WDR82* | 7582 | -3.91886 | 4.45E-05 | 0.002208 |
| *KRT17* | 1504 | -3.9187 | 4.45E-05 | 0.002199 |
| *ADCY1* | 48 | -3.91828 | 4.46E-05 | 0.002192 |
| *RNF123* | 6976 | -3.91801 | 4.46E-05 | 0.002184 |
| *TMEM191A* | 7849 | -3.90811 | 4.65E-05 | 0.002253 |
| *CEP83* | 5618 | -3.90781 | 4.66E-05 | 0.002245 |
| *PLBD2* | 9200 | -3.90714 | 4.67E-05 | 0.00224 |
| *PTPRM* | 2284 | -3.90706 | 4.67E-05 | 0.00223 |
| *PHF24* | 4842 | -3.90205 | 4.77E-05 | 0.002266 |
| *ACTR3* | 3976 | -3.9019 | 4.77E-05 | 0.002257 |
| *ARPC2* | 3984 | -3.9002 | 4.81E-05 | 0.002262 |
| *CCM2L* | 8845 | -3.89692 | 4.87E-05 | 0.002282 |
| *ADAMTS10* | 7684 | -3.89579 | 4.89E-05 | 0.002282 |
| *COPG2* | 5245 | -3.89448 | 4.92E-05 | 0.002284 |
| *MMGT1* | 8338 | -3.88503 | 5.12E-05 | 0.002364 |
| *EVL* | 5753 | -3.88404 | 5.14E-05 | 0.002363 |
| *MAX* | 1618 | -3.88368 | 5.14E-05 | 0.002355 |
| *RIIAD1* | 9531 | -3.87378 | 5.36E-05 | 0.002431 |
| *EME1* | 8918 | -3.86474 | 5.56E-05 | 0.002511 |
| *HCP5* | 4347 | -3.86052 | 5.66E-05 | 0.002532 |
| *TBC1D2B* | 4714 | -3.85872 | 5.70E-05 | 0.002529 |
| *IRF2BP1* | 5180 | -3.85675 | 5.75E-05 | 0.002538 |
| *BASP1* | 4130 | -3.85206 | 5.86E-05 | 0.002553 |
| *RESP18* | 9794 | -3.85147 | 5.87E-05 | 0.002548 |
| *PTDSS1* | 3821 | -3.85035 | 5.90E-05 | 0.002538 |
| *CIDECP* | 9022 | -3.84967 | 5.91E-05 | 0.002534 |
| *FAM210B* | 8506 | -3.84872 | 5.94E-05 | 0.002533 |
| *APBA1* | 140 | -3.84759 | 5.96E-05 | 0.002534 |
| *TMEM38A* | 7232 | -3.84657 | 5.99E-05 | 0.002534 |
| *STAT2* | 2704 | -3.84356 | 6.06E-05 | 0.002554 |
| *DNAJC30* | 7879 | -3.84223 | 6.10E-05 | 0.002558 |
| *NNAT* | 1839 | -3.83933 | 6.17E-05 | 0.002577 |
| *ASTN2* | 4786 | -3.83833 | 6.19E-05 | 0.002577 |
| *SHISAL1* | 8094 | -3.83749 | 6.21E-05 | 0.002575 |
| *ARMCX6* | 5926 | -3.83531 | 6.27E-05 | 0.002587 |
| *ZNF415* | 6430 | -3.83473 | 6.29E-05 | 0.002583 |
| *DTNA* | 747 | -3.83379 | 6.31E-05 | 0.002582 |
| *CREB3L3* | 7979 | -3.83369 | 6.31E-05 | 0.002573 |
| *VTA1* | 5772 | -3.8294 | 6.42E-05 | 0.002607 |
| *FARSB* | 3953 | -3.82713 | 6.48E-05 | 0.002621 |
| *CACNB1* | 342 | -3.82612 | 6.51E-05 | 0.002621 |
| *HMG20A* | 4108 | -3.81817 | 6.72E-05 | 0.002654 |
| *NOS1* | 1847 | -3.81397 | 6.84E-05 | 0.002678 |
| *TTC3P1* | 9596 | -3.80689 | 7.04E-05 | 0.002735 |
| *LRP3* | 1565 | -3.80654 | 7.05E-05 | 0.002728 |
| *C2CD6* | 8998 | -3.80376 | 7.13E-05 | 0.002748 |

| | | | | |
|---|---|---|---|---|
| *CPSF4* | 4362 | -3.80294 | 7.15E-05 | 0.002747 |
| *JOSD1* | 3892 | -3.80026 | 7.23E-05 | 0.002766 |
| *YWHAZ* | 3012 | -3.79398 | 7.41E-05 | 0.002815 |
| *GPNMB* | 4161 | -3.79286 | 7.45E-05 | 0.002807 |
| *MAP2* | 1608 | -3.78988 | 7.54E-05 | 0.00282 |
| *TMEM178A* | 8743 | -3.78936 | 7.55E-05 | 0.002815 |
| *MRPS33* | 5806 | -3.78931 | 7.55E-05 | 0.002805 |
| *ANGPT2* | 123 | -3.78792 | 7.60E-05 | 0.00281 |
| *FLOT2* | 954 | -3.78744 | 7.61E-05 | 0.002806 |
| *LDAH* | 6956 | -3.78354 | 7.73E-05 | 0.002839 |
| *UBE2L3* | 2938 | -3.78282 | 7.75E-05 | 0.002837 |
| *FHOD1* | 5420 | -3.78241 | 7.77E-05 | 0.002832 |
| *TROVE2* | 2691 | -3.77905 | 7.87E-05 | 0.002839 |
| *DLL3* | 4283 | -3.77722 | 7.93E-05 | 0.00285 |
| *CCND2* | 394 | -3.77517 | 7.99E-05 | 0.002853 |
| *FCGBP* | 3388 | -3.77306 | 8.06E-05 | 0.002867 |
| *ZKSCAN1* | 3033 | -3.77284 | 8.07E-05 | 0.002859 |
| *LPIN1* | 4748 | -3.77022 | 8.16E-05 | 0.002879 |
| *TTC9B* | 8939 | -3.76726 | 8.25E-05 | 0.002893 |
| *DYNLRB2* | 7735 | -3.76339 | 8.38E-05 | 0.002928 |
| *ZNF821* | 6326 | -3.751 | 8.81E-05 | 0.003024 |
| *NAE1* | 3402 | -3.74227 | 9.12E-05 | 0.00312 |
| *GNG5* | 1086 | -3.7346 | 9.40E-05 | 0.003174 |
| *TADA2B* | 8349 | -3.73284 | 9.47E-05 | 0.003185 |
| *PQLC1* | 7517 | -3.73251 | 9.48E-05 | 0.003179 |
| *ZNF233* | 9701 | -3.73192 | 9.50E-05 | 0.003176 |
| *GFOD1* | 5915 | -3.72948 | 9.59E-05 | 0.003196 |
| *MRPL34* | 7152 | -3.72105 | 9.92E-05 | 0.003294 |
| *LYNX1* | 7199 | -3.7208 | 9.93E-05 | 0.003286 |
| *PARD3B* | 8550 | -3.72079 | 9.93E-05 | 0.003275 |

**Table S26. Gene list for statistically significant associations with PPS-specific deviation patterns in the PLS2- component.** P values were estimated by using two-sided Z sampling distribution test without corrections. Q values presented corrected P values after the Benjamini-Hochberg method.

**9. Longitudinal Trajectories of Gene Expression for PPS-specific Markers**
For investigating whether these longitudinal gene expression profiles posed statistically significant trajectories, we built upon the Random Forest model to regress gene expression profiles across 30 developmental periods to specific developmental stage of brain regions identified in the RSA. The gene expression data, spanning 30 developmental periods, were organized into a 273 × 30 matrix (genes × periods), region-by-region. Using R package titled "randomForest" (Version 4.7-1.1), the random forest model was trained to predict the developmental periods based on gene expression levels, using 1,000 trees and bootstrapping with a mean square error

(MSE) criterion to evaluate predictive accuracy. Full results have been tabulated into the **Table S27**.

| Brain Parcels | Mean of squared residuals | Explained variance, % | P value at Permutation |
|---|---|---|---|
| Dorsolateral prefrontal cortex, DLPFC | 13.93 | 77.02 | < .0001 |
| Inferior temporal cortex, ITC | 14.97 | 69.33 | < .0001 |
| Caudal anterior cingulate cortex, cauACC | 11.48 | 81.26 | < .0001 |
| Orbital frontal cortex, OFC | 11.16 | 79.76 | < .0001 |
| Superior temporal cortex, STC | 13.98 | 79.31 | < .0001 |
| Rostral middle frontal cortex, rosMFC | 15.54 | 73.01 | < .0001 |

**Table S27. Model fits of the random forest regressing longitudinal gene expressions to neurodevelopmental periods.**

To further understand the contribution of individual genes to the neurodevelopmental period prediction model, we employed the R package "randomForestExplainer" to perform dominance analysis. This tool allowed us to assess the relative importance of each gene in predicting the neurodevelopmental periods. Using the model generated by the randomForest package, we calculated increased mean square error from the model by dropping out each gene. These analyses were performed by examining the permutation-based changes across all the genes to quantify importance (dominance). Full results for all the six brain parcels have been tabulated into the **Table S28-33**.

| Entrez ID | Gene Symbol | Mean_min_depth | IncrMSE, % | node_purity_increase | P value |
|---|---|---|---|---|---|
| 272 | *ZNF599* | 0.279069767 | 10.15220461 | 177.3412142 | 1.07E-43 |
| 165 | *PAFAH1B3* | 2.257790698 | 3.633761398 | 78.33153462 | 9.57E-11 |
| 21 | *ARMCX6* | 2.042930233 | 3.354612312 | 80.18262604 | 1.54E-13 |
| 57 | *DCHS1* | 2.324023256 | 3.166793848 | 60.22952066 | 3.98E-14 |
| 174 | *PHF2* | 2.372046512 | 3.144634052 | 56.88619843 | 2.77E-11 |
| 169 | *PCGF2* | 2.710255814 | 2.838129629 | 52.86734864 | 3.65E-05 |
| 227 | *SRRT* | 2.733511628 | 2.618729382 | 40.6717362 | 3.65E-05 |
| 105 | *HLF* | 2.397325581 | 2.559380209 | 47.15984744 | 2.77E-11 |
| 210 | *S100A1* | 3.050488372 | 2.307827389 | 38.45255453 | 0.047232436 |

| | | | | | |
|---|---|---|---|---|---|
| 246 | *TMEM38A* | 2.718348837 | 2.250556263 | 32.67216696 | 7.90E-07 |
| 171 | *PDE9A* | 2.482255814 | 1.984108508 | 48.75069113 | 1.07E-08 |
| 180 | *PLEKHA1* | 2.269930233 | 1.744130232 | 33.84472564 | 3.98E-14 |
| 155 | *NEUROD2* | 3.075767442 | 1.704755665 | 33.96173751 | 0.015386276 |
| 273 | *ZNF821* | 3.161209302 | 1.703584885 | 22.19433737 | 0.004249476 |
| 143 | *MPZL1* | 2.511581395 | 1.352953713 | 28.93413543 | 2.77E-11 |
| 70 | *EHD3* | 3.099023256 | 1.286951458 | 33.59894513 | 0.027536766 |
| 124 | *LRRC61* | 3.079813953 | 1.239461174 | 32.0774632 | 0.004249476 |
| 157 | *NNAT* | 3.095488372 | 1.125816569 | 27.57869267 | 0.002105567 |
| 97 | *GRAMD1A* | 3.248162791 | 0.973936444 | 24.2219164 | 0.027536766 |
| 86 | *FMNL1* | 3.022186047 | 0.60124787 | 21.79479712 | 8.80E-05 |
| 63 | *DLL3* | 2.898325581 | 0.583450364 | 14.28668701 | 2.16E-06 |
| 92 | *GFOD1* | 3.048976744 | 0.581610472 | 16.02567963 | 0.002105567 |
| 120 | *LIMD2* | 3.099023256 | 0.568774086 | 28.81611706 | 0.027536766 |
| 43 | *CDH4* | 3.515093023 | 0.560088892 | 9.389120081 | 0.46550996 |
| 106 | *HMG20A* | 3.128348837 | 0.532290406 | 8.70049722 | 0.004249476 |
| 126 | *LYNX1* | 3.205697674 | 0.510589287 | 11.32169928 | 0.008249565 |
| 135 | *MED30* | 3.323488372 | 0.499468406 | 11.11196631 | 0.121761504 |
| 121 | *LPIN1* | 3.203674419 | 0.493426981 | 11.9602164 | 0.027536766 |
| 102 | *HBA2* | 3.511046512 | 0.468179728 | 10.96835258 | 0.690033999 |
| 66 | *DTNA* | 3.358372093 | 0.415273002 | 3.975672254 | 0.182618492 |
| 239 | *TBC1D1* | 3.114697674 | 0.391021246 | 6.890704261 | 0.008249565 |
| 109 | *ILF2* | 3.254232558 | 0.388564361 | 11.02002701 | 0.008249565 |
| 238 | *TARSL2* | 3.381627907 | 0.385816329 | 8.423491514 | 0.182618492 |
| 96 | *GPNMB* | 3.536325581 | 0.365017014 | 4.947016226 | 0.46550996 |
| 24 | *ATF5* | 3.352813953 | 0.364996167 | 7.335389965 | 0.047232436 |
| 258 | *UBE2E2* | 3.233 | 0.359581667 | 9.052146018 | 0.004249476 |

| | | | | | |
|---|---|---|---|---|---|
| 103 | *HCP5* | 3.317930233 | 0.349025399 | 6.588752945 | 0.0275367 66 |
| 248 | *TNFAIP8 L3* | 3.325511628 | 0.31937649 | 7.306187032 | 0.1217615 04 |
| 3 | *ACTR3* | 3.567162791 | 0.311738925 | 5.23997172 | 0.7883461 09 |
| 5 | *ADAMTS 10* | 3.545930233 | 0.290415991 | 5.53346719 | 0.6900339 99 |
| 156 | *NFS1* | 3.263837209 | 0.280889977 | 4.689376747 | 0.0153862 76 |
| 190 | *PTDSS1* | 3.654116279 | 0.275416515 | 6.687424807 | 0.9628685 03 |
| 170 | *PCK2* | 3.287093023 | 0.274489652 | 3.969548206 | 0.0082495 65 |
| 134 | *MCHR1* | 3.402860465 | 0.260753561 | 6.09154597 | 0.2614427 81 |
| 243 | *THY1* | 3.306302326 | 0.259773179 | 7.245878965 | 0.0275367 66 |
| 117 | *KRT17* | 3.604069767 | 0.253922823 | 6.93576455 | 0.6900339 99 |
| 11 | *AMIGO1* | 3.619232558 | 0.252298 | 4.136276263 | 0.9628685 03 |
| 261 | *VTA1* | 3.613674419 | 0.233910942 | 4.428460005 | 0.7883461 09 |
| 244 | *TMEM19 1A* | 3.283046512 | 0.224223491 | 2.793006901 | 0.0472324 36 |
| 119 | *LEMD2* | 3.497906977 | 0.215745377 | 3.620875041 | 0.1217615 04 |
| 88 | *FOXP4* | 2.792162791 | 0.208207394 | 8.75044473 | 9.73E-08 |
| 29 | *BOP1* | 3.449372093 | 0.2049491 | 5.962520558 | 0.2614427 81 |
| 32 | *C7* | 3.228953488 | 0.199413372 | 3.454927646 | 0.0153862 76 |
| 17 | *AQP6* | 3.439767442 | 0.19879219 | 5.46713872 | 0.1826184 92 |
| 74 | *EPS8L2* | 3.543906977 | 0.196561049 | 4.782355797 | 0.7883461 09 |
| 100 | *GTF2F1* | 3.515093023 | 0.191807749 | 5.699627992 | 0.4655099 6 |
| 27 | *BASP1* | 3.652093023 | 0.182788135 | 2.415395082 | 0.9839588 07 |
| 91 | *G3BP1* | 3.432186047 | 0.17642214 | 1.818698324 | 0.0775533 24 |
| 270 | *ZNF233* | 3.569186047 | 0.171038221 | 2.331798453 | 0.6900339 99 |
| 138 | *MICU1* | 3.367976744 | 0.15132608 | 4.172518326 | 0.2614427 81 |
| 215 | *SF3A3* | 3.230976744 | 0.149922494 | 4.548926046 | 0.0021055 67 |

| | | | | | |
|---|---|---|---|---|---|
| 19 | *ARIH2* | 3.400837209 | 0.145601788 | 2.998255028 | 0.357129511 |
| 116 | *KIF19* | 3.487790698 | 0.142110425 | 9.797355026 | 0.690033999 |
| 229 | *SSSCA1* | 3.66372093 | 0.140930926 | 5.271385533 | 0.983958807 |
| 249 | *TRADD* | 3.696581395 | 0.13979214 | 0.909078571 | 0.994172272 |
| 218 | *SLC38A2* | 3.632883721 | 0.139100444 | 1.354203221 | 0.925550627 |
| 35 | *CABYR* | 3.221372093 | 0.137117081 | 3.698480694 | 0.004249476 |
| 1 | *AATK* | 3.549976744 | 0.135347112 | 2.63504121 | 0.46550996 |
| 217 | *SLC1A4* | 3.29872093 | 0.127565861 | 2.769867591 | 0.015386276 |
| 228 | *SSFA2* | 3.673325581 | 0.124567475 | 3.524095308 | 0.994172272 |
| 87 | *FNBP4* | 3.578790698 | 0.122845993 | 1.445533333 | 0.788346109 |
| 162 | *NRSN2* | 3.458976744 | 0.122348945 | 1.628881974 | 0.357129511 |
| 211 | *SAMD15* | 3.451395349 | 0.119155221 | 3.429007931 | 0.182618492 |
| 61 | *DEPDC5* | 3.600023256 | 0.11730575 | 1.155452381 | 0.867790238 |
| 223 | *SP100* | 3.43572093 | 0.116618351 | 3.197090866 | 0.357129511 |
| 71 | *EIF1B* | 3.590418605 | 0.110601864 | 2.599059476 | 0.788346109 |
| 251 | *TRIO* | 3.567162791 | 0.110349198 | 3.129811294 | 0.788346109 |
| 154 | *NEURL3* | 3.447348837 | 0.109432915 | 6.355760886 | 0.357129511 |
| 14 | *ANKS6* | 3.456953488 | 0.109053666 | 2.264325236 | 0.46550996 |
| 47 | *CFD* | 3.484255814 | 0.108571067 | 1.277985981 | 0.261442781 |
| 68 | *DYNLRB2* | 3.675348837 | 0.103825695 | 2.393915873 | 0.983958807 |
| 44 | *CDK5R2* | 3.331581395 | 0.099739252 | 2.437829421 | 0.027536766 |
| 262 | *WDR82* | 3.520651163 | 0.099202001 | 11.80873535 | 0.788346109 |
| 77 | *FABP3* | 3.571209302 | 0.09901091 | 1.035790332 | 0.579465733 |
| 187 | *PSMB10* | 3.327534884 | 0.098729303 | 3.873064561 | 0.047232436 |
| 159 | *NOS1AP* | 3.540372093 | 0.098645697 | 0.72787619 | 0.357129511 |

| | | | | | |
|---|---|---|---|---|---|
| 137 | *METRN* | 3.62327907 | 0.096404995 | 0.828708333 | 0.867790238 |
| 233 | *STXBP1* | 3.569186047 | 0.095118775 | 4.413993497 | 0.690033999 |
| 23 | *ASTN2* | 3.505488372 | 0.093944223 | 0.680281097 | 0.357129511 |
| 64 | *DNAJC30* | 3.602046512 | 0.091731636 | 0.822371861 | 0.788346109 |
| 191 | *PTPRM* | 3.534302326 | 0.091608393 | 2.119289057 | 0.690033999 |
| 205 | *RNFT2* | 3.588395349 | 0.09096016 | 2.313567582 | 0.867790238 |
| 108 | *IDS* | 3.580813953 | 0.08853958 | 0.808805647 | 0.690033999 |
| 220 | *SNCG* | 3.53227907 | 0.083702959 | 2.869199235 | 0.788346109 |
| 131 | *MAP6* | 3.565139535 | 0.083525652 | 6.614292633 | 0.788346109 |
| 65 | *DSTYK* | 3.360395349 | 0.08198674 | 3.441546758 | 0.121761504 |
| 193 | *PTPRU* | 3.406906977 | 0.080049615 | 2.197272733 | 0.121761504 |
| 38 | *CAPRIN2* | 3.611651163 | 0.076959167 | 1.004934626 | 0.867790238 |
| 18 | *ARF3* | 3.524697674 | 0.075288609 | 1.741720391 | 0.579465733 |
| 230 | *STARD9* | 3.634906977 | 0.074192074 | 1.399207287 | 0.867790238 |
| 250 | *TRIM4* | 3.559581395 | 0.069150107 | 1.002087302 | 0.46550996 |
| 252 | *TROVE2* | 3.600023256 | 0.063984198 | 1.150301893 | 0.867790238 |
| 139 | *MKRN3* | 3.083860465 | 0.06364268 | 8.331196179 | 0.002105567 |
| 189 | *PSME3* | 3.646534884 | 0.061034531 | 0.593606566 | 0.867790238 |
| 36 | *CACNB1* | 3.503465116 | 0.059422111 | 1.912654473 | 0.46550996 |
| 192 | *PTPRS* | 3.354837209 | 0.057257224 | 2.159439977 | 0.027536766 |
| 130 | *MAP4K2* | 3.285069767 | 0.055514263 | 10.17367912 | 0.027536766 |
| 53 | *CREB3L3* | 3.172837209 | 0.05443019 | 7.936060919 | 0.008249565 |
| 31 | *C16orf70* | 3.62327907 | 0.054312049 | 3.824877931 | 0.867790238 |
| 73 | *EME1* | 3.733488372 | 0.053786768 | 1.717033489 | 0.983958807 |
| 234 | *STYXL1* | 3.563627907 | 0.052516035 | 0.585163492 | 0.357129511 |

| | | | | | |
|---|---|---|---|---|---|
| 267 | *ZEB2* | 3.62327907 | 0.05153679 | 1.043213292 | 0.867790238 |
| 62 | *DLG4* | 3.712255814 | 0.051210256 | 0.381729493 | 0.962868503 |
| 204 | *RNF208* | 3.532790698 | 0.050840841 | 0.702336203 | 0.121761504 |
| 255 | *TTC9B* | 3.385674419 | 0.050377975 | 2.821513111 | 0.077553324 |
| 168 | *PCDHAC2* | 3.611651163 | 0.050007963 | 1.284758442 | 0.867790238 |
| 206 | *RPL13A* | 3.661697674 | 0.050001818 | 3.797336606 | 0.994172272 |
| 226 | *SRPK2* | 3.578790698 | 0.04904725 | 1.5746805 | 0.690033999 |
| 265 | *YWHAB* | 3.48627907 | 0.048869172 | 0.821173443 | 0.182618492 |
| 28 | *BCAN* | 3.366465116 | 0.048318499 | 2.030904669 | 0.027536766 |
| 148 | *MYH3* | 3.733488372 | 0.047703471 | 0.213821212 | 0.983958807 |
| 93 | *GLT8D1* | 3.710232558 | 0.047380056 | 0.2627557 | 0.983958807 |
| 216 | *SLC10A4* | 3.667767442 | 0.046116235 | 0.537697985 | 0.925550627 |
| 221 | *SORBS3* | 3.719837209 | 0.045224697 | 0.309101299 | 0.994172272 |
| 202 | *RILPL1* | 3.644511628 | 0.044684649 | 0.60351034 | 0.925550627 |
| 39 | *CCND2* | 3.700627907 | 0.041925019 | 0.235987302 | 0.962868503 |
| 41 | *CCT6B* | 3.650581395 | 0.038645103 | 0.38632619 | 0.690033999 |
| 271 | *ZNF415* | 3.686976744 | 0.037998939 | 0.64607619 | 0.983958807 |
| 9 | *AGPAT5* | 3.517116279 | 0.037165152 | 2.172910007 | 0.357129511 |
| 207 | *RPL24* | 3.762302326 | 0.037020833 | 0.146369048 | 0.999622323 |
| 237 | *TADA2B* | 3.538348837 | 0.0366923 | 1.084649206 | 0.46550996 |
| 136 | *MEST* | 3.658162791 | 0.036052083 | 0.308002381 | 0.867790238 |
| 10 | *AHCYL2* | 3.721860465 | 0.035469378 | 0.530240867 | 0.983958807 |
| 197 | *RAMP1* | 3.350790698 | 0.035456735 | 2.314062932 | 0.077553324 |
| 122 | *LRP1* | 3.721860465 | 0.033993333 | 0.239128355 | 0.962868503 |
| 72 | *ELOVL6* | 3.611651163 | 0.032847833 | 0.698576611 | 0.788346109 |

| | | | | | |
|---|---|---|---|---|---|
| 113 | *JOSD1* | 3.584860465 | 0.031689026 | 0.647301243 | 0.4655099<br>6 |
| 145 | *MRPL49* | 3.752697674 | 0.031472222 | 0.095457143 | 0.9982928<br>04 |
| 51 | *COPG2* | 3.540372093 | 0.031410347 | 0.917631457 | 0.3571295<br>11 |
| 141 | *MLYCD* | 3.741069767 | 0.031316857 | 0.281404762 | 0.9982928<br>04 |
| 179 | *PLD3* | 3.575255814 | 0.028797278 | 0.454880087 | 0.3571295<br>11 |
| 128 | *MANEAL* | 3.731465116 | 0.027922659 | 0.101006349 | 0.9941722<br>72 |
| 209 | *RTF1* | 3.652093023 | 0.027873384 | 2.293239562 | 0.9839588<br>07 |
| 178 | *PLBD2* | 3.64855814 | 0.027637753 | 0.388974315 | 0.7883461<br>09 |
| 196 | *RAB12* | 3.660186047 | 0.026968119 | 0.340770758 | 0.7883461<br>09 |
| 80 | *FAM174<br>B* | 3.675348837 | 0.026942955 | 0.729642569 | 0.9839588<br>07 |
| 15 | *AP2S1* | 3.721860465 | 0.024245926 | 0.153445238 | 0.9839588<br>07 |
| 167 | *PCBD1* | 3.708209302 | 0.023425926 | 0.66687619 | 0.9941722<br>72 |
| 112 | *JAZF1* | 3.710232558 | 0.022064667 | 0.730041811 | 0.9839588<br>07 |
| 49 | *CNOT1* | 3.646534884 | 0.021798467 | 0.957445665 | 0.8677902<br>38 |
| 151 | *NDUFA9* | 3.511046512 | 0.021050778 | 8.927639754 | 0.6900339<br>99 |
| 147 | *MT1A* | 3.677372093 | 0.020529321 | 0.594627289 | 0.9255506<br>27 |
| 55 | *CYFIP2* | 3.66372093 | 0.020395025 | 2.722696072 | 0.9839588<br>07 |
| 254 | *TTC3P1* | 3.743093023 | 0.02036065 | 0.105214286 | 0.9941722<br>72 |
| 111 | *IRF2BP1* | 3.636930233 | 0.020117741 | 0.227477778 | 0.7883461<br>09 |
| 224 | *SPIRE1* | 3.441790698 | 0.018163107 | 1.827703019 | 0.1217615<br>04 |
| 236 | *SYT3* | 3.549976744 | 0.017811846 | 0.793313925 | 0.4655099<br>6 |
| 2 | *ACACA* | 3.72944186 | 0.015924374 | 0.432410823 | 0.9982928<br>04 |
| 107 | *IDI1* | 3.731465116 | 0.015692708 | 0.2225 | 0.9941722<br>72 |
| 79 | *FAM160<br>B1* | 3.700627907 | 0.015202764 | 0.222530952 | 0.9628685<br>03 |
| 232 | *STX4* | 3.59244186 | 0.014920303 | 0.960239916 | 0.6900339<br>99 |

| | | | | | |
|---|---|---|---|---|---|
| 98 | *GRIN2B* | 3.775953488 | 0.014778875 | 0.100942857 | 0.998292804 |
| 259 | *UBE2L3* | 3.517116279 | 0.01446512 | 1.275051854 | 0.357129511 |
| 37 | *CAND2* | 3.625302326 | 0.014329008 | 0.317815873 | 0.788346109 |
| 214 | *SERTAD3* | 3.658162791 | 0.013592662 | 0.264559524 | 0.867790238 |
| 225 | *SPTAN1* | 3.752697674 | 0.013357576 | 0.186207143 | 0.998292804 |
| 160 | *NPHP4* | 3.689 | 0.012329242 | 0.680478571 | 0.962868503 |
| 104 | *HEMK1* | 3.611651163 | 0.012224711 | 0.881469048 | 0.867790238 |
| 163 | *NUTF2* | 3.700627907 | 0.011987514 | 0.178503968 | 0.962868503 |
| 194 | *PWWP2B* | 3.55755814 | 0.011928028 | 1.147654168 | 0.690033999 |
| 84 | *FHOD1* | 3.538348837 | 0.010691369 | 4.454371693 | 0.46550996 |
| 132 | *MARK4* | 3.634906977 | 0.010644444 | 0.400233333 | 0.867790238 |
| 264 | *YIF1B* | 3.658162791 | 0.010009616 | 0.46229011 | 0.867790238 |
| 25 | *ATP6V1G1* | 3.762302326 | 0.00942 | 0.469329004 | 0.999622323 |
| 144 | *MRPL34* | 3.681418605 | 0.009249231 | 0.250629365 | 0.867790238 |
| 182 | *POLR2E* | 3.731465116 | 0.00911875 | 0.128642857 | 0.994172272 |
| 260 | *VAC14* | 3.689 | 0.008038611 | 0.243216667 | 0.962868503 |
| 42 | *CD34* | 3.681418605 | 0.0080125 | 0.264107143 | 0.867790238 |
| 199 | *RESP18* | 3.717813953 | 0.007257956 | 0.632756227 | 0.998292804 |
| 75 | *ETV5* | 3.735511628 | 0.007083333 | 0.126452381 | 0.962868503 |
| 118 | *KRTCAP2* | 3.749162791 | 0.006972972 | 0.057628571 | 0.925550627 |
| 161 | *NPPB* | 3.78555814 | 0.006666667 | 0.076242857 | 0.999622323 |
| 34 | *C9orf16* | 3.771906977 | 0.006545455 | 0.352075483 | 0.999943962 |
| 245 | *TMEM199* | 3.773930233 | 0.005973333 | 0.072433333 | 0.999622323 |
| 85 | *FLOT2* | 3.719837209 | 0.005827273 | 0.173809524 | 0.994172272 |
| 54 | *CSTB* | 3.719837209 | 0.005438718 | 0.417442641 | 0.994172272 |

| | | | | | |
|---|---|---|---|---|---|
| 16 | *APBA1* | 3.698604651 | 0.00540235 | 0.176296753 | 0.983958807 |
| 200 | *RFXANK* | 3.717813953 | 0.005013418 | 0.399377778 | 0.998292804 |
| 184 | *PPP5C* | 3.710232558 | 0.004272412 | 0.187902381 | 0.983958807 |
| 149 | *NAE1* | 3.602046512 | 0.004194536 | 0.928340049 | 0.788346109 |
| 99 | *GRK6* | 3.764325581 | 0.003765152 | 0.208935065 | 0.998292804 |
| 4 | *ACTR6* | 3.766348837 | 0.00375 | 0.170416667 | 0.994172272 |
| 7 | *ADCY5* | 3.625302326 | 0.003717929 | 0.339892063 | 0.788346109 |
| 185 | *PQLC1* | 3.721860465 | 0.003716599 | 0.175425397 | 0.983958807 |
| 166 | *PARD3B* | 3.708209302 | 0.003624514 | 0.481227778 | 0.994172272 |
| 164 | *OAZ2* | 3.675348837 | 0.003395611 | 0.799953824 | 0.983958807 |
| 133 | *MAX* | 3.25827907 | 0.003357125 | 5.228807311 | 0.000462216 |
| 127 | *MALL* | 3.656139535 | 0.002896603 | 0.745432143 | 0.925550627 |
| 146 | *MRPS33* | 3.644511628 | 0.002762832 | 0.556072176 | 0.925550627 |
| 173 | *PEX11A* | 3.636930233 | 0.002566325 | 0.441237518 | 0.788346109 |
| 183 | *PPDPF* | 3.733488372 | 0.002557365 | 0.198742857 | 0.983958807 |
| 81 | *FARSB* | 3.689 | 0.002252747 | 0.208996825 | 0.962868503 |
| 69 | *EEF1A2* | 3.752697674 | 0.002008519 | 0.178568182 | 0.998292804 |
| 129 | *MAP2* | 3.773930233 | 0.002 | 0.034828571 | 0.999622323 |
| 22 | *ARPC2* | 3.708209302 | 0.001801481 | 0.234620779 | 0.994172272 |
| 177 | *PITPNM2* | 3.437744186 | 0.001516534 | 1.859238312 | 0.261442781 |
| 208 | *RPL7L1* | 3.667767442 | 0.001493333 | 0.738348718 | 0.925550627 |
| 110 | *IPO5* | 3.677372093 | 0.000680505 | 0.52364445 | 0.962868503 |
| 45 | *CEP19* | 3.000953488 | 0.000666303 | 6.905845946 | 1.47E-05 |
| 186 | *PRB1* | 3.783534884 | 0.000458333 | 0.03275 | 0.999943962 |
| 150 | *NDFIP1* | 3.731465116 | 0.000284299 | 0.385323932 | 0.994172272 |

| | | | | | |
|---|---|---|---|---|---|
| 240 | *TBC1D2B* | 3.686976744 | 0.000281905 | 0.55691746 | 0.983958807 |
| 266 | *YWHAZ* | 3.764325581 | 0 | 0.081742857 | 0.998292804 |
| 56 | *DARS* | 3.658162791 | -0.00020297 | 0.479207648 | 0.867790238 |
| 78 | *FAHD1* | 3.731465116 | -0.000360795 | 0.125058608 | 0.994172272 |
| 76 | *EVL* | 3.627325581 | -0.001451583 | 0.357711905 | 0.690033999 |
| 152 | *NDUFS4* | 3.804767442 | -0.0016 | 0.021 | 0.999995821 |
| 172 | *PDK2* | 3.667767442 | -0.002245463 | 0.266664286 | 0.925550627 |
| 115 | *KCNQ2* | 3.700627907 | -0.002336746 | 0.158907143 | 0.962868503 |
| 58 | *DCUN1D5* | 3.741069767 | -0.003143732 | 0.265699206 | 0.998292804 |
| 33 | *C9orf116* | 3.646534884 | -0.00318461 | 0.36684127 | 0.867790238 |
| 213 | *SEC24D* | 3.783534884 | -0.0035 | 0.112777778 | 0.999943962 |
| 222 | *SOS1* | 3.816395349 | -0.003818182 | 0.01225 | 0.999995821 |
| 94 | *GNG5* | 3.691023256 | -0.004058308 | 0.296499784 | 0.925550627 |
| 123 | *LRP3* | 3.530767442 | -0.004131486 | 0.694041631 | 0.261442781 |
| 40 | *CCNYL1* | 3.717813953 | -0.00529053 | 0.606013348 | 0.998292804 |
| 52 | *CPSF4* | 3.613674419 | -0.005418318 | 0.796935736 | 0.788346109 |
| 253 | *TRUB1* | 3.743093023 | -0.005613889 | 0.13055873 | 0.994172272 |
| 181 | *POLR1A* | 3.698604651 | -0.006149383 | 0.270961905 | 0.983958807 |
| 60 | *DEF8* | 3.708209302 | -0.006636857 | 0.322457576 | 0.994172272 |
| 188 | *PSMD7* | 3.708209302 | -0.006733686 | 0.235091198 | 0.994172272 |
| 212 | *SCRN1* | 3.721860465 | -0.007450617 | 0.242273016 | 0.983958807 |
| 50 | *COG1* | 3.698604651 | -0.008009773 | 0.457294805 | 0.983958807 |
| 140 | *MLLT11* | 3.768372093 | -0.00819321 | 0.098583333 | 0.983958807 |
| 26 | *ATP9B* | 3.588395349 | -0.008855566 | 1.145681136 | 0.867790238 |
| 175 | *PIGF* | 3.689 | -0.009273611 | 0.162713636 | 0.962868503 |

| | | | | | |
|---|---|---|---|---|---|
| 201 | *RIIAD1* | 3.698604651 | -0.009319815 | 0.600724875 | 0.983958807 |
| 241 | *TCF7* | 3.632883721 | -0.010333086 | 0.843228283 | 0.925550627 |
| 90 | *FZD5* | 3.549976744 | -0.012583522 | 1.587818398 | 0.46550996 |
| 153 | *NEK9* | 3.733488372 | -0.01518 | 0.386194872 | 0.983958807 |
| 269 | *ZKSCAN1* | 3.451395349 | -0.016359199 | 1.56262381 | 0.182618492 |
| 198 | *RASSF7* | 3.762302326 | -0.016983165 | 0.154657576 | 0.999622323 |
| 8 | *AFTPH* | 3.686976744 | -0.0178 | 0.593579487 | 0.983958807 |
| 125 | *LRRN2* | 3.686976744 | -0.017837556 | 0.421971429 | 0.983958807 |
| 89 | *FRMD8* | 3.721860465 | -0.018085601 | 0.239849639 | 0.983958807 |
| 13 | *ANKIB1* | 3.656139535 | -0.018099747 | 0.262331746 | 0.925550627 |
| 257 | *UBE2B* | 3.677372093 | -0.018449532 | 0.57057381 | 0.962868503 |
| 6 | *ADCY1* | 3.609627907 | -0.019986505 | 1.263623684 | 0.925550627 |
| 242 | *THOP1* | 3.741069767 | -0.021987866 | 0.495139977 | 0.998292804 |
| 235 | *SYDE2* | 3.698604651 | -0.022093213 | 0.642468254 | 0.983958807 |
| 82 | *FBXL18* | 3.762302326 | -0.023567937 | 0.173511111 | 0.999622323 |
| 203 | *RNF123* | 3.698604651 | -0.023914396 | 0.29245 | 0.983958807 |
| 231 | *STAT2* | 3.578790698 | -0.02543107 | 1.048837302 | 0.788346109 |
| 12 | *ANGPT2* | 3.625302326 | -0.025735522 | 0.59344127 | 0.788346109 |
| 176 | *PIGX* | 3.634906977 | -0.025864154 | 1.125330464 | 0.867790238 |
| 219 | *SLC38A3* | 3.580813953 | -0.028395297 | 1.235978716 | 0.690033999 |
| 158 | *NOS1* | 3.540372093 | -0.028575968 | 1.218967372 | 0.261442781 |
| 20 | *ARL2* | 3.621255814 | -0.034453467 | 0.787929926 | 0.925550627 |
| 268 | *ZER1* | 3.561604651 | -0.034598067 | 0.7543345 | 0.46550996 |
| 195 | *RAB11FIP2* | 3.636930233 | -0.037425551 | 0.314619336 | 0.788346109 |
| 101 | *HBA1* | 3.461 | -0.03768152 | 1.437530808 | 0.261442781 |

| | | | | | |
|---|---|---|---|---|---|
| 114 | *JUND* | 3.689 | -0.037714328 | 0.452954762 | 0.962868503 |
| 67 | *DYNLRB1* | 3.698604651 | -0.038158877 | 0.718423016 | 0.962868503 |
| 48 | *CIDECP* | 3.590418605 | -0.044166389 | 1.719912725 | 0.788346109 |
| 95 | *GNPDA2* | 3.372023256 | -0.045561975 | 6.238131636 | 0.077553324 |
| 247 | *TMSB10* | 3.669790698 | -0.058368162 | 1.07418602 | 0.788346109 |
| 83 | *FCGBP* | 3.569186047 | -0.066747082 | 2.960797358 | 0.690033999 |
| 263 | *WNK2* | 3.457465116 | -0.074601109 | 1.406038384 | 0.027536766 |
| 46 | *CETN3* | 3.667767442 | -0.077938215 | 2.588884275 | 0.925550627 |
| 142 | *MMGT1* | 3.62327907 | -0.0801394 | 1.242292857 | 0.867790238 |
| 256 | *TTPAL* | 3.580813953 | -0.089789167 | 3.574477147 | 0.690033999 |
| 30 | *C11orf1* | 3.677372093 | -0.10285375 | 0.706803263 | 0.962868503 |
| 59 | *DDX51* | 3.578790698 | -0.295594348 | 3.50481969 | 0.690033999 |

**Table S28. Importance (Dominance) of each gene in the random forest model for dorsolateral prefrontal cortex (DLPFC).** Mean_min_depth represents the average minimum depth at which a variable is used as a split node across all trees, with lower values indicating higher importance as the variable is used earlier in the decision process. IncrMSE, % (Increase in Mean Squared Error) quantifies the relative increase in prediction error when the variable is permuted, reflecting its contribution to the model's predictive accuracy; higher values signify greater importance. Node_purity_increase measures the improvement in node purity (e.g., Gini index or variance reduction) achieved by splitting on the variable, where larger values indicate a stronger impact on classification or regression performance. Finally, the P value assesses the statistical significance of a variable's importance, typically derived from permutation tests, with lower values suggesting that the variable's contribution is unlikely to be due to random chance. This presentation was in descending order by the IncrMSE.

| Entrez ID | Gene Symbol | Mean_min_depth | IncrMSE, % | node_purity_increase | P value |
|---|---|---|---|---|---|
| 121 | *LPIN1* | 0.333333333 | 6.328232532 | 115.5919012 | 3.11E-46 |
| 210 | *S100A1* | 1.178857143 | 5.130646657 | 82.69278327 | 3.39E-29 |
| 238 | *TARSL2* | 1.90847619 | 2.320165893 | 50.94331432 | 1.71E-17 |

| | | | | | |
|---|---|---|---|---|---|
| 157 | *NNAT* | 2.240857143 | 1.781239643 | 35.20557291 | 9.96E-11 |
| 262 | *WDR82* | 2.663619048 | 1.547717108 | 26.06112024 | 9.35E-06 |
| 70 | *EHD3* | 2.585142857 | 1.51432766 | 32.52857666 | 9.35E-06 |
| 228 | *SSFA2* | 2.428190476 | 1.468837929 | 39.2553179 | 1.21E-06 |
| 272 | *ZNF599* | 2.203428571 | 1.385385446 | 23.9569785 | 3.55E-16 |
| 63 | *DLL3* | 2.798952381 | 1.31388851 | 23.18122472 | 0.000849002 |
| 65 | *DSTYK* | 2.26247619 | 1.060323185 | 25.72542893 | 7.24E-12 |
| 136 | *MEST* | 3.045809524 | 0.971063739 | 17.43194534 | 0.134596469 |
| 27 | *BASP1* | 2.941333333 | 0.920310916 | 19.2274631 | 0.01539031 |
| 53 | *CREB3L3* | 2.925047619 | 0.914918284 | 17.50478967 | 0.001866014 |
| 126 | *LYNX1* | 2.756666667 | 0.857476033 | 10.25575541 | 3.41E-06 |
| 248 | *TNFAIP8L3* | 2.851428571 | 0.830128613 | 18.96990098 | 0.000371221 |
| 147 | *MT1A* | 2.877428571 | 0.802575775 | 15.91780176 | 0.001866014 |
| 251 | *TRIO* | 3.300190476 | 0.659162884 | 7.079593038 | 0.5218103 |
| 143 | *MPZL1* | 2.932095238 | 0.6274236 | 16.32075312 | 0.000849002 |
| 273 | *ZNF821* | 2.837333333 | 0.609523839 | 14.72791116 | 6.33E-05 |
| 140 | *MLLT11* | 3.366761905 | 0.442176758 | 6.389815152 | 0.642940286 |
| 171 | *PDE9A* | 3.041428571 | 0.43305162 | 10.79180801 | 0.01539031 |
| 165 | *PAFAH1B3* | 2.97752381 | 0.400961065 | 11.59622159 | 0.000371221 |
| 139 | *MKRN3* | 2.887142857 | 0.389008004 | 13.26809294 | 0.000371221 |
| 116 | *KIF19* | 3.152952381 | 0.386482859 | 12.62962224 | 0.134596469 |
| 124 | *LRRC61* | 3.240666667 | 0.366362715 | 9.800822722 | 0.5218103 |
| 135 | *MED30* | 3.174571429 | 0.357179328 | 5.412997958 | 0.050154344 |
| 215 | *SF3A3* | 3.340285714 | 0.35596989 | 5.300223413 | 0.91522955 |
| 206 | *RPL13A* | 3.058190476 | 0.325277004 | 7.859054459 | 0.00795636 |
| 220 | *SNCG* | 2.751809524 | 0.311279698 | 10.32016066 | 1.21E-06 |
| 26 | *ATP9B* | 3.357047619 | 0.285444444 | 5.229102181 | 0.8470741 |
| 92 | *GFOD1* | 3.376 | 0.282516343 | 5.09671317 | 0.91522955 |
| 59 | *DDX51* | 3.150761905 | 0.26283952 | 5.703827935 | 0.050154344 |
| 222 | *SOS1* | 3.283428571 | 0.259630273 | 6.568999675 | 0.642940286 |
| 180 | *PLEKHA1* | 3.380857143 | 0.246082808 | 3.558786797 | 0.8470741 |

| | | | | | |
|---|---|---|---|---|---|
| 57 | *DCHS1* | 3.250380952 | 0.23621669 | 8.686809617 | 0.294363209 |
| 169 | *PCGF2* | 3.380857143 | 0.230003497 | 2.555249258 | 0.8470741 |
| 247 | *TMSB10* | 3.328857143 | 0.226031302 | 2.95964254 | 0.402340183 |
| 103 | *HCP5* | 3.007904762 | 0.222787544 | 14.0622689 | 0.050154344 |
| 29 | *BOP1* | 3.347809524 | 0.209639673 | 4.427190476 | 0.5218103 |
| 192 | *PTPRS* | 3.392761905 | 0.209178437 | 3.017117582 | 0.8470741 |
| 131 | *MAP6* | 3.283428571 | 0.194380049 | 4.193710262 | 0.642940286 |
| 209 | *RTF1* | 3.23847619 | 0.188002778 | 7.138392191 | 0.294363209 |
| 21 | *ARMCX6* | 3.447428571 | 0.184886649 | 0.775052394 | 0.91522955 |
| 73 | *EME1* | 2.927714286 | 0.182537177 | 4.989211491 | 0.000156191 |
| 102 | *HBA2* | 3.326190476 | 0.172947506 | 4.910230952 | 0.754550572 |
| 120 | *LIMD2* | 3.331047619 | 0.170382817 | 1.234478283 | 0.642940286 |
| 243 | *THY1* | 3.387904762 | 0.167234735 | 4.44218254 | 0.91522955 |
| 227 | *SRRT* | 3.361904762 | 0.165599414 | 1.50447361 | 0.642940286 |
| 202 | *RILPL1* | 3.385714286 | 0.161379048 | 4.103891181 | 0.754550572 |
| 230 | *STARD9* | 3.260095238 | 0.15079908 | 4.741751123 | 0.134596469 |
| 235 | *SYDE2* | 3.354857143 | 0.141196084 | 2.526234182 | 0.642940286 |
| 163 | *NUTF2* | 3.283904762 | 0.139441562 | 1.527220635 | 0.134596469 |
| 134 | *MCHR1* | 3.361904762 | 0.135337484 | 1.945227922 | 0.754550572 |
| 190 | *PTDSS1* | 3.300190476 | 0.132310541 | 3.496683442 | 0.5218103 |
| 170 | *PCK2* | 3.319142857 | 0.125367623 | 2.969683242 | 0.642940286 |
| 159 | *NOS1AP* | 3.269333333 | 0.125183866 | 3.785831471 | 0.402340183 |
| 17 | *AQP6* | 2.956380952 | 0.124802358 | 4.844879293 | 6.33E-05 |
| 174 | *PHF2* | 3.338095238 | 0.123855371 | 3.430097619 | 0.754550572 |
| 96 | *GPNMB* | 3.319142857 | 0.122589792 | 5.194300901 | 0.642940286 |
| 254 | *TTC3P1* | 3.19352381 | 0.121943991 | 2.994964516 | 0.084241286 |
| 119 | *LEMD2* | 3.366761905 | 0.120119521 | 1.145744206 | 0.642940286 |

| | | | | | |
|---|---|---|---|---|---|
| 167 | *PCBD1* | 3.138857143 | 0.119804496 | 4.747344078 | 0.050154344 |
| 164 | *OAZ2* | 3.046285714 | 0.11742269 | 9.508498269 | 0.00795636 |
| 28 | *BCAN* | 3.065238095 | 0.115765754 | 3.527959693 | 0.01539031 |
| 181 | *POLR1A* | 3.29047619 | 0.107662391 | 6.949303968 | 0.642940286 |
| 11 | *AMIGO1* | 3.423619048 | 0.106624778 | 2.854878571 | 0.91522955 |
| 8 | *AFTPH* | 3.46152381 | 0.105105 | 0.894759341 | 0.983389462 |
| 50 | *COG1* | 3.298 | 0.103501724 | 0.848304834 | 0.294363209 |
| 24 | *ATF5* | 3.069619048 | 0.103175253 | 19.81216113 | 0.134596469 |
| 105 | *HLF* | 3.354857143 | 0.095029342 | 2.345765904 | 0.642940286 |
| 39 | *CCND2* | 3.210285714 | 0.093221248 | 1.772600042 | 0.050154344 |
| 133 | *MAX* | 3.314761905 | 0.092584206 | 0.80769127 | 0.204311145 |
| 246 | *TMEM38A* | 3.411714286 | 0.091996951 | 2.675482051 | 0.8470741 |
| 132 | *MARK4* | 3.492380952 | 0.0915375 | 0.3655 | 0.99454045 |
| 187 | *PSMB10* | 3.347809524 | 0.091159015 | 1.509202151 | 0.5218103 |
| 151 | *NDUFA9* | 3.368952381 | 0.088334937 | 4.787776881 | 0.8470741 |
| 249 | *TRADD* | 3.385714286 | 0.085908153 | 2.121236905 | 0.754550572 |
| 37 | *CAND2* | 3.309904762 | 0.084133972 | 0.566343001 | 0.294363209 |
| 10 | *AHCYL2* | 3.43552381 | 0.082109999 | 0.716978584 | 0.91522955 |
| 38 | *CAPRIN2* | 3.162666667 | 0.079181768 | 4.17060575 | 0.050154344 |
| 87 | *FNBP4* | 3.309904762 | 0.077790499 | 2.217430087 | 0.294363209 |
| 43 | *CDH4* | 3.42847619 | 0.075762794 | 0.838670766 | 0.8470741 |
| 155 | *NEUROD2* | 3.324 | 0.075553111 | 1.842360944 | 0.5218103 |
| 225 | *SPTAN1* | 3.430666667 | 0.074894325 | 2.219922872 | 0.959144262 |
| 261 | *VTA1* | 3.469047619 | 0.072520432 | 2.429871279 | 0.754550572 |
| 72 | *ELOVL6* | 3.333238095 | 0.072259167 | 6.124806066 | 0.8470741 |
| 156 | *NFS1* | 3.376 | 0.066525833 | 4.383497619 | 0.91522955 |
| 45 | *CEP19* | 3.293142857 | 0.063491466 | 1.477032684 | 0.402340183 |

| | | | | | |
|---|---|---|---|---|---|
| 31 | *C16orf70* | 3.404666667 | 0.061108165 | 2.38672378 | 0.8470741 |
| 214 | *SERTAD3* | 3.354857143 | 0.059756944 | 0.960819481 | 0.642940286 |
| 86 | *FMNL1* | 3.331047619 | 0.059242582 | 0.934569786 | 0.642940286 |
| 9 | *AGPAT5* | 3.452285714 | 0.058815377 | 0.250197619 | 0.8470741 |
| 66 | *DTNA* | 3.366761905 | 0.058778211 | 1.047836075 | 0.642940286 |
| 13 | *ANKIB1* | 3.485333333 | 0.057125062 | 0.538433766 | 0.983389462 |
| 75 | *ETV5* | 3.392761905 | 0.056267063 | 2.300046465 | 0.8470741 |
| 234 | *STYXL1* | 3.319619048 | 0.055500873 | 0.590546609 | 0.084241286 |
| 61 | *DEPDC5* | 3.452285714 | 0.053624296 | 0.521955556 | 0.8470741 |
| 82 | *FBXL18* | 3.279047619 | 0.053075823 | 1.103531979 | 0.204311145 |
| 49 | *CNOT1* | 3.366761905 | 0.052140999 | 1.182333542 | 0.642940286 |
| 178 | *PLBD2* | 3.528095238 | 0.051503704 | 0.153857143 | 0.99454045 |
| 19 | *ARIH2* | 3.335904762 | 0.051317 | 1.18686746 | 0.5218103 |
| 109 | *ILF2* | 3.328857143 | 0.050779994 | 1.143679293 | 0.402340183 |
| 76 | *EVL* | 3.399809524 | 0.048964997 | 0.988230736 | 0.91522955 |
| 150 | *NDFIP1* | 3.447428571 | 0.048651375 | 0.310038095 | 0.91522955 |
| 204 | *RNF208* | 3.378666667 | 0.048408698 | 1.816531469 | 0.642940286 |
| 138 | *MICU1* | 3.298 | 0.048043038 | 1.066618681 | 0.204311145 |
| 197 | *RAMP1* | 3.43552381 | 0.046928182 | 1.00415772 | 0.91522955 |
| 97 | *GRAMD1A* | 3.504285714 | 0.046554653 | 0.740708333 | 0.99454045 |
| 2 | *ACACA* | 3.48047619 | 0.04560787 | 0.273255556 | 0.99454045 |
| 48 | *CIDECP* | 3.473428571 | 0.043405247 | 0.376518254 | 0.983389462 |
| 108 | *IDS* | 3.274190476 | 0.043059162 | 1.263957504 | 0.134596469 |
| 15 | *AP2S1* | 3.478285714 | 0.042296296 | 0.243057143 | 0.959144262 |
| 193 | *PTPRU* | 3.293142857 | 0.042089411 | 1.566996581 | 0.402340183 |
| 7 | *ADCY5* | 3.42847619 | 0.041966915 | 0.488042857 | 0.8470741 |
| 94 | *GNG5* | 3.478285714 | 0.039965556 | 0.257516667 | 0.959144262 |
| 68 | *DYNLRB2* | 3.324 | 0.039808808 | 3.814439416 | 0.5218103 |

| | | | | | |
|---|---|---|---|---|---|
| 60 | *DEF8* | 3.392761905 | 0.039027309 | 0.584318651 | 0.8470741 |
| 64 | *DNAJC30* | 3.430666667 | 0.037713188 | 2.015331254 | 0.959144262 |
| 113 | *JOSD1* | 3.335904762 | 0.037558667 | 1.385381044 | 0.402340183 |
| 141 | *MLYCD* | 3.037047619 | 0.036128712 | 3.544246774 | 0.000849002 |
| 154 | *NEURL3* | 3.416571429 | 0.03602287 | 0.5357557 | 0.8470741 |
| 162 | *NRSN2* | 3.423619048 | 0.033770833 | 0.390311905 | 0.91522955 |
| 69 | *EEF1A2* | 3.40247619 | 0.033670626 | 0.918394444 | 0.642940286 |
| 258 | *UBE2E2* | 3.281238095 | 0.03345442 | 1.397401443 | 0.402340183 |
| 41 | *CCT6B* | 3.459333333 | 0.032365239 | 0.559181061 | 0.91522955 |
| 218 | *SLC38A2* | 3.392761905 | 0.031585708 | 0.718663187 | 0.8470741 |
| 78 | *FAHD1* | 3.380857143 | 0.029447751 | 1.023218498 | 0.8470741 |
| 80 | *FAM174B* | 3.490190476 | 0.029138222 | 0.235929293 | 0.959144262 |
| 130 | *MAP4K2* | 3.373809524 | 0.028211905 | 0.829636719 | 0.754550572 |
| 191 | *PTPRM* | 2.844857143 | 0.027993919 | 4.444960744 | 1.21E-06 |
| 257 | *UBE2B* | 3.466380952 | 0.027268233 | 0.311816667 | 0.959144262 |
| 255 | *TTC9B* | 3.295333333 | 0.026670747 | 3.045615201 | 0.642940286 |
| 115 | *KCNQ2* | 3.504285714 | 0.025927167 | 0.400421062 | 0.99454045 |
| 260 | *VAC14* | 3.509142857 | 0.024734722 | 0.164472222 | 0.983389462 |
| 242 | *THOP1* | 3.459333333 | 0.024603039 | 0.51548456 | 0.91522955 |
| 240 | *TBC1D2B* | 3.380857143 | 0.023631515 | 0.778414652 | 0.8470741 |
| 177 | *PITPNM2* | 3.347809524 | 0.02311617 | 0.963523449 | 0.5218103 |
| 44 | *CDK5R2* | 3.547047619 | 0.021550833 | 0.096833333 | 0.998641963 |
| 16 | *APBA1* | 3.492380952 | 0.020962 | 0.280593651 | 0.99454045 |
| 183 | *PPDPF* | 3.478285714 | 0.020922222 | 0.11109697 | 0.959144262 |
| 118 | *KRTCAP2* | 3.395428571 | 0.020597383 | 0.40565873 | 0.5218103 |
| 5 | *ADAMTS10* | 3.442571429 | 0.020198651 | 0.707258242 | 0.959144262 |
| 233 | *STXBP1* | 3.469047619 | 0.020124848 | 0.399283333 | 0.754550572 |

| | | | | | |
|---|---|---|---|---|---|
| 117 | *KRT17* | 3.267142857 | 0.019925123 | 1.556634349 | 0.134596469 |
| 40 | *CCNYL1* | 3.497238095 | 0.019648194 | 0.549745238 | 0.959144262 |
| 245 | *TMEM199* | 3.48047619 | 0.0191775 | 1.452504762 | 0.99454045 |
| 172 | *PDK2* | 3.521047619 | 0.019067497 | 0.152516667 | 0.983389462 |
| 25 | *ATP6V1G1* | 3.476095238 | 0.017988889 | 0.230242063 | 0.8470741 |
| 236 | *SYT3* | 3.359714286 | 0.017898716 | 0.494388023 | 0.5218103 |
| 266 | *YWHAZ* | 3.566 | 0.017777778 | 0.017047619 | 0.999773032 |
| 95 | *GNPDA2* | 3.504285714 | 0.017767295 | 0.197052381 | 0.99454045 |
| 267 | *ZEB2* | 3.236285714 | 0.017717546 | 2.654859979 | 0.134596469 |
| 219 | *SLC38A3* | 3.445238095 | 0.017222167 | 0.825184704 | 0.754550572 |
| 195 | *RAB11FIP2* | 3.404666667 | 0.017170561 | 0.388683333 | 0.8470741 |
| 184 | *PPP5C* | 3.433333333 | 0.016575778 | 0.571144078 | 0.754550572 |
| 199 | *RESP18* | 3.38352381 | 0.016343984 | 0.447103319 | 0.5218103 |
| 265 | *YWHAB* | 3.509142857 | 0.015873016 | 0.140445238 | 0.983389462 |
| 107 | *IDI1* | 3.502095238 | 0.015823056 | 0.100166667 | 0.959144262 |
| 229 | *SSSCA1* | 3.495047619 | 0.015370754 | 0.163814286 | 0.91522955 |
| 42 | *CD34* | 3.497238095 | 0.014458486 | 0.462585714 | 0.959144262 |
| 207 | *RPL24* | 3.485333333 | 0.013789956 | 0.440107143 | 0.983389462 |
| 268 | *ZER1* | 3.447428571 | 0.013731508 | 0.345145455 | 0.91522955 |
| 194 | *PWWP2B* | 3.516190476 | 0.013568889 | 0.227511111 | 0.99454045 |
| 123 | *LRP3* | 3.395428571 | 0.013554329 | 0.506042935 | 0.402340183 |
| 269 | *ZKSCAN1* | 3.274190476 | 0.01345247 | 1.712530536 | 0.204311145 |
| 112 | *JAZF1* | 3.404666667 | 0.013271517 | 0.831794567 | 0.8470741 |
| 176 | *PIGX* | 3.485333333 | 0.012910222 | 0.411088853 | 0.959144262 |
| 129 | *MAP2* | 3.426285714 | 0.012125248 | 0.239780952 | 0.5218103 |
| 111 | *IRF2BP1* | 3.395428571 | 0.012031702 | 0.758047619 | 0.5218103 |
| 221 | *SORBS3* | 3.547047619 | 0.011946667 | 0.102955556 | 0.998641963 |

| | | | | | |
|---|---|---|---|---|---|
| 205 | *RNFT2* | 3.45447619 | 0.011902381 | 0.388968926 | 0.959144262 |
| 188 | *PSMD7* | 3.40952381 | 0.011799603 | 0.876466667 | 0.754550572 |
| 212 | *SCRN1* | 3.034380952 | 0.011576441 | 5.42959479 | 0.00795636 |
| 23 | *ASTN2* | 3.476095238 | 0.010900476 | 0.121344444 | 0.8470741 |
| 1 | *AATK* | 3.103619048 | 0.010445914 | 1.974760373 | 0.001866014 |
| 56 | *DARS* | 3.558952381 | 0.00908642 | 0.076444444 | 0.998641963 |
| 77 | *FABP3* | 3.440380952 | 0.008551022 | 0.620055861 | 0.8470741 |
| 168 | *PCDHAC2* | 3.452285714 | 0.008061514 | 0.330638817 | 0.8470741 |
| 158 | *NOS1* | 3.471238095 | 0.007577166 | 0.322127171 | 0.91522955 |
| 232 | *STX4* | 3.483142857 | 0.007125 | 0.24202381 | 0.91522955 |
| 263 | *WNK2* | 3.509142857 | 0.00708642 | 0.603252381 | 0.983389462 |
| 244 | *TMEM191A* | 3.361904762 | 0.006788165 | 1.342643656 | 0.754550572 |
| 127 | *MALL* | 3.371619048 | 0.006722659 | 0.58858254 | 0.5218103 |
| 106 | *HMG20A* | 3.43552381 | 0.006666741 | 0.387873016 | 0.91522955 |
| 55 | *CYFIP2* | 3.457142857 | 0.006487405 | 0.808529604 | 0.754550572 |
| 160 | *NPHP4* | 3.504285714 | 0.00575 | 0.296621795 | 0.99454045 |
| 104 | *HEMK1* | 3.502095238 | 0.004962963 | 0.161488889 | 0.959144262 |
| 144 | *MRPL34* | 3.566 | 0.004909091 | 0.029880952 | 0.999773032 |
| 34 | *C9orf16* | 3.46152381 | 0.004790617 | 0.753466023 | 0.983389462 |
| 237 | *TADA2B* | 3.267142857 | 0.004708182 | 1.253133039 | 0.204311145 |
| 152 | *NDUFS4* | 3.333714286 | 0.004258714 | 0.596025325 | 0.204311145 |
| 85 | *FLOT2* | 3.504285714 | 0.0042 | 0.392628205 | 0.99454045 |
| 198 | *RASSF7* | 3.577904762 | 0.004 | 0.028714286 | 0.999773032 |
| 203 | *RNF123* | 3.528095238 | 0.004 | 0.08402381 | 0.99454045 |
| 196 | *RAB12* | 3.421428571 | 0.003878162 | 0.571278272 | 0.754550572 |
| 253 | *TRUB1* | 3.399809524 | 0.003398827 | 1.04367664 | 0.91522955 |

| | | | | | |
|---|---|---|---|---|---|
| 47 | *CFD* | 3.514 | 0.003215724 | 0.205638889 | 0.959144262 |
| 211 | *SAMD15* | 3.430666667 | 0.002439473 | 0.513058608 | 0.959144262 |
| 122 | *LRP1* | 3.464190476 | 0.002406984 | 0.284077778 | 0.8470741 |
| 46 | *CETN3* | 3.504285714 | 0.001737913 | 0.354361111 | 0.99454045 |
| 81 | *FARSB* | 3.570857143 | 0.000833333 | 0.014452381 | 0.998641963 |
| 110 | *IPO5* | 3.43552381 | 0.00064784 | 0.557834127 | 0.91522955 |
| 99 | *GRK6* | 3.509142857 | 0.000390804 | 0.168344444 | 0.983389462 |
| 18 | *ARF3* | 3.397619048 | 0.000355491 | 0.628983838 | 0.754550572 |
| 128 | *MANEAL* | 3.516190476 | 0.000242222 | 0.565173856 | 0.99454045 |
| 36 | *CACNB1* | 3.577904762 | 0 | 0.011 | 0.999773032 |
| 161 | *NPPB* | 3.584952381 | 0 | 0.002666667 | 0.999980905 |
| 186 | *PRB1* | NA | 0 | 0 | 1 |
| 259 | *UBE2L3* | 3.573047619 | 0 | 0.0225 | 0.999980905 |
| 270 | *ZNF233* | 3.528095238 | 0 | 0.120847619 | 0.99454045 |
| 137 | *METRN* | 3.404666667 | -0.000212639 | 0.748718681 | 0.8470741 |
| 90 | *FZD5* | 3.40952381 | -0.000621235 | 0.51666241 | 0.754550572 |
| 83 | *FCGBP* | 3.490190476 | -0.001519605 | 0.057257143 | 0.959144262 |
| 142 | *MMGT1* | 3.521047619 | -0.001742985 | 0.228019048 | 0.983389462 |
| 173 | *PEX11A* | 3.495047619 | -0.001812467 | 0.124589255 | 0.91522955 |
| 22 | *ARPC2* | 3.497238095 | -0.002032182 | 0.194802381 | 0.983389462 |
| 3 | *ACTR3* | 3.423619048 | -0.002638794 | 2.762083061 | 0.91522955 |
| 93 | *GLT8D1* | 3.309904762 | -0.002916415 | 1.065829726 | 0.294363209 |
| 153 | *NEK9* | 3.547047619 | -0.0036 | 0.043357143 | 0.998641963 |
| 256 | *TTPAL* | 3.488 | -0.003601111 | 0.138447619 | 0.8470741 |
| 216 | *SLC10A4* | 3.203238095 | -0.004587514 | 1.204470418 | 0.02844456 |
| 33 | *C9orf116* | 3.324 | -0.004834722 | 0.931461508 | 0.5218103 |

| | | | | | |
|---|---|---|---|---|---|
| 149 | *NAE1* | 3.478285714 | -0.005268571 | 0.130933333 | 0.959144262 |
| 88 | *FOXP4* | 3.354857143 | -0.005665366 | 1.052130447 | 0.642940286 |
| 6 | *ADCY1* | 3.509142857 | -0.005822711 | 0.383659957 | 0.983389462 |
| 145 | *MRPL49* | 3.551904762 | -0.010822222 | 0.051128571 | 0.99454045 |
| 71 | *EIF1B* | 3.528095238 | -0.012604248 | 0.100983333 | 0.99454045 |
| 114 | *JUND* | 3.414380952 | -0.013277778 | 0.386348557 | 0.642940286 |
| 148 | *MYH3* | 3.279047619 | -0.014304095 | 2.825472799 | 0.204311145 |
| 224 | *SPIRE1* | 3.385714286 | -0.01513357 | 2.1034571 | 0.642940286 |
| 35 | *CABYR* | 3.42847619 | -0.017380125 | 0.507745937 | 0.8470741 |
| 217 | *SLC1A4* | 3.457142857 | -0.017509505 | 0.246845238 | 0.754550572 |
| 182 | *POLR2E* | 3.473428571 | -0.017771111 | 0.385193651 | 0.983389462 |
| 98 | *GRIN2B* | 3.316952381 | -0.017965749 | 0.911292063 | 0.294363209 |
| 51 | *COPG2* | 3.385714286 | -0.01815084 | 2.598931349 | 0.754550572 |
| 12 | *ANGPT2* | 3.492380952 | -0.018439394 | 0.543211538 | 0.99454045 |
| 125 | *LRRN2* | 3.497238095 | -0.019299603 | 0.174119048 | 0.983389462 |
| 241 | *TCF7* | 3.497238095 | -0.02018 | 0.211256349 | 0.983389462 |
| 264 | *YIF1B* | 3.423619048 | -0.02075342 | 0.896480952 | 0.8470741 |
| 89 | *FRMD8* | 3.18647619 | -0.020835903 | 1.988472923 | 0.050154344 |
| 208 | *RPL7L1* | 3.440380952 | -0.021232014 | 0.401380952 | 0.8470741 |
| 52 | *CPSF4* | 3.43552381 | -0.022563605 | 0.392766667 | 0.91522955 |
| 213 | *SEC24D* | 3.421428571 | -0.024698951 | 0.687384127 | 0.642940286 |
| 189 | *PSME3* | 3.547047619 | -0.024933333 | 0.087695238 | 0.998641963 |
| 58 | *DCUN1D5* | 3.390571429 | -0.03090646 | 0.705653175 | 0.642940286 |
| 231 | *STAT2* | 3.442571429 | -0.032700278 | 0.700740404 | 0.959144262 |
| 101 | *HBA1* | 3.378666667 | -0.033145979 | 2.587101515 | 0.642940286 |

| | | | | | |
|---|---|---|---|---|---|
| 146 | *MRPS33* | 3.416571429 | -0.033358416 | 0.439817827 | 0.8470741 |
| 226 | *SRPK2* | 3.46152381 | -0.035385247 | 0.938238095 | 0.983389462 |
| 250 | *TRIM4* | 3.466380952 | -0.036324416 | 0.281719048 | 0.959144262 |
| 239 | *TBC1D1* | 3.053333333 | -0.037588404 | 4.750019081 | 0.00795636 |
| 185 | *PQLC1* | 3.404666667 | -0.037911722 | 2.321261905 | 0.8470741 |
| 67 | *DYNLRB1* | 3.485333333 | -0.038152778 | 0.377789683 | 0.983389462 |
| 79 | *FAM160B1* | 3.485333333 | -0.038506448 | 0.369896825 | 0.983389462 |
| 166 | *PARD3B* | 3.471238095 | -0.038978774 | 0.201961905 | 0.91522955 |
| 200 | *RFXANK* | 3.466380952 | -0.039962963 | 0.182561905 | 0.959144262 |
| 20 | *ARL2* | 3.397619048 | -0.042627778 | 0.806817965 | 0.754550572 |
| 252 | *TROVE2* | 3.473428571 | -0.043538667 | 0.486579739 | 0.983389462 |
| 271 | *ZNF415* | 3.530285714 | -0.045090909 | 0.276352814 | 0.999773032 |
| 100 | *GTF2F1* | 3.316952381 | -0.049112782 | 2.455763728 | 0.402340183 |
| 91 | *G3BP1* | 3.40247619 | -0.052134722 | 0.597924747 | 0.642940286 |
| 30 | *C11orf1* | 3.21952381 | -0.054330302 | 2.973488836 | 0.204311145 |
| 84 | *FHOD1* | 3.40952381 | -0.055687727 | 2.312311305 | 0.754550572 |
| 54 | *CSTB* | 3.324 | -0.056758372 | 2.554789805 | 0.5218103 |
| 4 | *ACTR6* | 3.535142857 | -0.059816905 | 0.131233333 | 0.998641963 |
| 62 | *DLG4* | 3.447428571 | -0.063359759 | 0.825939683 | 0.91522955 |
| 32 | *C7* | 3.359714286 | -0.06647252 | 1.222721465 | 0.5218103 |
| 179 | *PLD3* | 3.516190476 | -0.066881944 | 0.337043651 | 0.99454045 |
| 14 | *ANKS6* | 3.404666667 | -0.071438422 | 1.014695238 | 0.754550572 |
| 74 | *EPS8L2* | 3.430666667 | -0.074053357 | 2.955715734 | 0.959144262 |
| 223 | *SP100* | 3.45447619 | -0.088434815 | 0.78048663 | 0.959144262 |
| 175 | *PIGF* | 3.368952381 | -0.09274316 | 5.087844972 | 0.8470741 |
| 201 | *RIIAD1* | 3.119904762 | -0.101989687 | 5.557361598 | 0.02844456 |

**Table S29. Importance (Dominance) of each gene in the random forest model for inferior temporal cortex (ITC).** Mean_min_depth represents the average minimum depth at which a variable is used as a split node across all trees, with lower values indicating higher importance as the variable is used earlier in the decision process. IncrMSE, % (Increase in Mean Squared Error) quantifies the relative increase in prediction error when the variable is permuted, reflecting its contribution to the model's predictive accuracy; higher values signify greater importance. Node_purity_increase measures the improvement in node purity (e.g., Gini index or variance reduction) achieved by splitting on the variable, where larger values indicate a stronger impact on classification or regression performance. Finally, the P value assesses the statistical significance of a variable's importance, typically derived from permutation tests, with lower values suggesting that the variable's contribution is unlikely to be due to random chance. This presentation was in descending order by the IncrMSE.

| Entrez ID | Gene Symbol | Mean_min_depth | IncrMSE, % | node_purity_increase | P value |
|---|---|---|---|---|---|
| 57 | *DCHS1* | 0.497142857 | 8.143890182 | 132.452106 | 2.38E-35 |
| 272 | *ZNF599* | 0.493506494 | 6.467941544 | 108.2950501 | 7.78E-44 |
| 220 | *SNCG* | 1.749090909 | 3.649047637 | 69.19369172 | 5.16E-15 |
| 165 | *PAFAH1B3* | 2.357818182 | 2.638101134 | 39.60053128 | 2.51E-08 |
| 174 | *PHF2* | 2.289246753 | 2.396762684 | 36.09853959 | 6.27E-11 |
| 97 | *GRAMD1A* | 2.301402597 | 2.276609912 | 52.59915347 | 2.51E-08 |
| 169 | *PCGF2* | 2.650805195 | 2.185506116 | 40.15165126 | 0.000454621 |
| 143 | *MPZL1* | 2.448311688 | 1.937770901 | 39.65290063 | 6.60E-07 |
| 124 | *LRRC61* | 2.767272727 | 1.89844814 | 40.95415539 | 0.004395841 |
| 155 | *NEUROD2* | 2.685714286 | 1.895869987 | 31.33352513 | 3.37E-05 |
| 126 | *LYNX1* | 2.521766234 | 1.714603974 | 32.33486944 | 7.68E-08 |
| 120 | *LIMD2* | 2.866701299 | 1.499187152 | 28.03395575 | 0.004395841 |
| 157 | *NNAT* | 3.156051948 | 1.299747483 | 15.37592117 | 0.132556825 |
| 247 | *TMSB10* | 2.444675325 | 1.233198296 | 23.38866851 | 2.47E-09 |
| 105 | *HLF* | 2.935688312 | 1.20835532 | 24.1326229 | 0.016322288 |
| 70 | *EHD3* | 3.043636364 | 1.175262622 | 23.15400174 | 0.084167123 |
| 238 | *TARSL2* | 2.880103896 | 1.162081907 | 15.64416119 | 0.000454621 |
| 26 | *ATP9B* | 2.897142857 | 1.126340035 | 24.75868631 | 0.004395841 |

| | | | | | |
|---|---|---|---|---|---|
| 246 | *TMEM38 A* | 3.207584416 | 1.119837776 | 17.48711218 | 0.3867373 96 |
| 243 | *THY1* | 3.061090909 | 1.071540642 | 18.64033599 | 0.0294972 37 |
| 171 | *PDE9A* | 2.871584416 | 0.947926049 | 19.63487564 | 0.0001979 13 |
| 27 | *BASP1* | 3.259532468 | 0.834176486 | 12.07176671 | 0.3867373 96 |
| 180 | *PLEKHA 1* | 3.198649351 | 0.758706523 | 16.18616575 | 0.6180304 88 |
| 222 | *SOS1* | 2.866701299 | 0.722615675 | 26.73246763 | 0.0086498 17 |
| 134 | *MCHR1* | 3.341090909 | 0.658737854 | 10.31088368 | 0.8956835 98 |
| 108 | *IDS* | 2.897142857 | 0.625445214 | 16.39827781 | 0.0021451 57 |
| 17 | *AQP6* | 3.164987013 | 0.552491808 | 8.791303769 | 0.0509829 23 |
| 210 | *S100A1* | 3.289558442 | 0.52176873 | 8.48046332 | 0.6180304 88 |
| 61 | *DEPDC5* | 2.897558442 | 0.521022506 | 5.470007749 | 0.0001979 13 |
| 273 | *ZNF821* | 2.992103896 | 0.502856526 | 14.01897977 | 0.0163222 88 |
| 197 | *RAMP1* | 3.156467532 | 0.493900002 | 4.449308519 | 0.0294972 37 |
| 132 | *MARK4* | 3.307428571 | 0.49252527 | 4.09668486 | 0.1989477 4 |
| 24 | *ATF5* | 3.182025974 | 0.426375072 | 9.608029699 | 0.1325568 25 |
| 248 | *TNFAIP8 L3* | 3.255064935 | 0.384377894 | 14.97591176 | 0.3867373 96 |
| 76 | *EVL* | 3.276571429 | 0.379690093 | 9.356794505 | 0.6180304 88 |
| 219 | *SLC38A3* | 3.031064935 | 0.373768368 | 4.653072976 | 0.0163222 88 |
| 263 | *WNK2* | 2.945038961 | 0.364808639 | 4.167178745 | 0.0010065 04 |
| 237 | *TADA2B* | 2.927584416 | 0.350800613 | 5.067622743 | 0.0021451 57 |
| 206 | *RPL13A* | 3.419012987 | 0.349590774 | 5.765147253 | 0.8956835 98 |
| 140 | *MLLT11* | 3.311480519 | 0.341322496 | 8.083892974 | 0.3867373 96 |
| 262 | *WDR82* | 3.358545455 | 0.320835571 | 6.405511321 | 0.8229211 64 |
| 170 | *PCK2* | 2.846441558 | 0.314234281 | 4.567849204 | 1.85E-06 |
| 139 | *MKRN3* | 3.238441558 | 0.290154526 | 9.244272097 | 0.0509829 23 |

| | | | | | |
|---|---|---|---|---|---|
| 86 | *FMNL1* | 3.376 | 0.283202417 | 2.146738428 | 0.728431059 |
| 151 | *NDUFA9* | 3.436467532 | 0.279768998 | 4.483384715 | 0.822921164 |
| 207 | *RPL24* | 3.354493506 | 0.251375988 | 2.912180445 | 0.618030488 |
| 1 | *AATK* | 3.113454545 | 0.250358126 | 3.000426993 | 0.008649817 |
| 121 | *LPIN1* | 2.867116883 | 0.247044732 | 13.47744462 | 0.001006504 |
| 18 | *ARF3* | 3.276987013 | 0.241993585 | 2.730586851 | 0.284316455 |
| 191 | *PTPRM* | 3.018493506 | 0.240874184 | 4.42901894 | 0.002145157 |
| 99 | *GRK6* | 3.414961039 | 0.238501959 | 3.350209219 | 0.728431059 |
| 201 | *RIIAD1* | 3.264 | 0.237986451 | 8.41238199 | 0.284316455 |
| 96 | *GPNMB* | 3.130493506 | 0.235827497 | 2.939545882 | 0.029497237 |
| 101 | *HBA1* | 3.294441558 | 0.23021285 | 5.636707187 | 0.19894774 |
| 135 | *MED30* | 3.203948052 | 0.227551648 | 3.161321615 | 0.050982923 |
| 48 | *CIDECP* | 3.083012987 | 0.215687884 | 4.095414109 | 0.016322288 |
| 215 | *SF3A3* | 3.264 | 0.214620642 | 5.387089966 | 0.284316455 |
| 73 | *EME1* | 3.397506494 | 0.213015086 | 4.104248696 | 0.822921164 |
| 255 | *TTC9B* | 3.272519481 | 0.207568642 | 2.829490186 | 0.386737396 |
| 116 | *KIF19* | 3.6 | 0.197568182 | 1.690089044 | 0.999894045 |
| 193 | *PTPRU* | 2.975064935 | 0.178935039 | 5.206627784 | 0.004395841 |
| 209 | *RTF1* | 3.208 | 0.178809375 | 3.949485095 | 0.132556825 |
| 63 | *DLL3* | 3.216935065 | 0.169301466 | 5.011414405 | 0.050982923 |
| 21 | *ARMCX6* | 3.457974026 | 0.161859008 | 3.619095721 | 0.822921164 |
| 270 | *ZNF233* | 3.432 | 0.160745859 | 2.8653819 | 0.895683598 |
| 38 | *CAPRIN2* | 3.035532468 | 0.156960735 | 4.033158592 | 0.008649817 |
| 75 | *ETV5* | 3.376 | 0.145822733 | 1.167631746 | 0.728431059 |
| 136 | *MEST* | 3.311064935 | 0.141388434 | 12.10863836 | 0.728431059 |

| | | | | | |
|---|---|---|---|---|---|
| 233 | *STXBP1* | 3.367480519 | 0.138908809 | 3.927929154 | 0.618030488 |
| 227 | *SRRT* | 3.247376623 | 0.131140179 | 2.676861876 | 0.008649817 |
| 264 | *YIF1B* | 3.513974026 | 0.124769302 | 1.814383272 | 0.975275022 |
| 216 | *SLC10A4* | 3.156467532 | 0.123758733 | 2.141904847 | 0.029497237 |
| 192 | *PTPRS* | 3.371532468 | 0.12162162 | 4.682189893 | 0.822921164 |
| 117 | *KRT17* | 3.397922078 | 0.118783687 | 0.726642063 | 0.386737396 |
| 29 | *BOP1* | 3.246961039 | 0.117662242 | 2.324023366 | 0.132556825 |
| 51 | *COPG2* | 3.363012987 | 0.117660496 | 1.813247222 | 0.728431059 |
| 66 | *DTNA* | 2.962493506 | 0.103163701 | 3.922369736 | 0.000454621 |
| 31 | *C16orf70* | 3.488 | 0.10295436 | 4.034459921 | 0.975275022 |
| 49 | *CNOT1* | 3.160519481 | 0.094790884 | 3.292435714 | 0.084167123 |
| 183 | *PPDPF* | 3.354493506 | 0.093548923 | 7.066522699 | 0.618030488 |
| 152 | *NDUFS4* | 3.457974026 | 0.093474818 | 2.112240139 | 0.895683598 |
| 198 | *RASSF7* | 3.324467532 | 0.090966392 | 1.92370772 | 0.386737396 |
| 269 | *ZKSCAN1* | 3.264 | 0.090484977 | 2.320921784 | 0.284316455 |
| 128 | *MANEAL* | 3.509506494 | 0.090300556 | 2.106139182 | 0.990559909 |
| 258 | *UBE2E2* | 3.479480519 | 0.090094937 | 0.982590476 | 0.945473238 |
| 177 | *PITPNM2* | 3.440519481 | 0.08982505 | 6.731187295 | 0.945473238 |
| 254 | *TTC3P1* | 3.324467532 | 0.088552015 | 1.485837246 | 0.386737396 |
| 10 | *AHCYL2* | 3.470961039 | 0.088364729 | 0.578449206 | 0.895683598 |
| 30 | *C11orf1* | 3.345974026 | 0.081698571 | 1.689381173 | 0.500803679 |
| 28 | *BCAN* | 3.272519481 | 0.076863355 | 2.05966859 | 0.386737396 |
| 162 | *NRSN2* | 3.449454545 | 0.076384546 | 0.975300354 | 0.822921164 |
| 106 | *HMG20A* | 3.423480519 | 0.074100083 | 1.153699351 | 0.822921164 |
| 205 | *RNFT2* | 3.432 | 0.073519192 | 2.520728921 | 0.895683598 |

| | | | | | |
|---|---|---|---|---|---|
| 190 | *PTDSS1* | 3.380467532 | 0.072785985 | 0.974622799 | 0.618030488 |
| 65 | *DSTYK* | 3.242077922 | 0.070873152 | 6.974456183 | 0.500803679 |
| 144 | *MRPL34* | 3.397506494 | 0.069799653 | 2.131088217 | 0.822921164 |
| 94 | *GNG5* | 3.225454545 | 0.065569942 | 1.688964358 | 0.084167123 |
| 53 | *CREB3L3* | 3.384519481 | 0.065076878 | 4.126865074 | 0.822921164 |
| 159 | *NOS1AP* | 3.341506494 | 0.062888027 | 1.928413855 | 0.618030488 |
| 7 | *ADCY5* | 3.281454545 | 0.061671402 | 2.229859327 | 0.19894774 |
| 59 | *DDX51* | 3.401974026 | 0.061215745 | 1.73912521 | 0.728431059 |
| 231 | *STAT2* | 3.457974026 | 0.059894848 | 0.592290224 | 0.895683598 |
| 35 | *CABYR* | 3.354493506 | 0.059824802 | 1.597697131 | 0.500803679 |
| 249 | *TRADD* | 3.444987013 | 0.057670045 | 1.228035714 | 0.895683598 |
| 212 | *SCRN1* | 3.471376623 | 0.056419921 | 0.611231746 | 0.618030488 |
| 118 | *KRTCAP2* | 3.509922078 | 0.056010902 | 0.454958608 | 0.895683598 |
| 245 | *TMEM199* | 3.40238961 | 0.055989446 | 0.713931335 | 0.386737396 |
| 81 | *FARSB* | 3.414961039 | 0.055849985 | 0.696603968 | 0.728431059 |
| 200 | *RFXANK* | 3.604467532 | 0.054877922 | 0.262266667 | 0.999322301 |
| 228 | *SSFA2* | 3.556987013 | 0.054363413 | 0.560853463 | 0.997090681 |
| 130 | *MAP4K2* | 3.229506494 | 0.054045971 | 2.482733578 | 0.19894774 |
| 172 | *PDK2* | 3.526961039 | 0.0529194 | 0.34153961 | 0.975275022 |
| 78 | *FAHD1* | 3.561454545 | 0.052786891 | 0.395588562 | 0.990559909 |
| 36 | *CACNB1* | 3.350441558 | 0.052750056 | 3.394556899 | 0.386737396 |
| 11 | *AMIGO1* | 3.535480519 | 0.052337721 | 2.388269356 | 0.975275022 |
| 223 | *SP100* | 3.384519481 | 0.051388212 | 2.051562384 | 0.822921164 |
| 131 | *MAP6* | 3.466909091 | 0.051383182 | 0.714897619 | 0.728431059 |
| 189 | *PSME3* | 3.496935065 | 0.051206219 | 0.416220635 | 0.895683598 |

| | | | | | |
|---|---|---|---|---|---|
| 62 | *DLG4* | 3.483948052 | 0.051015783 | 0.248189683 | 0.895683598 |
| 251 | *TRIO* | 3.544 | 0.049777576 | 0.635390909 | 0.997090681 |
| 119 | *LEMD2* | 3.500987013 | 0.048787682 | 2.878838711 | 0.975275022 |
| 163 | *NUTF2* | 3.315948052 | 0.044980175 | 1.436722095 | 0.284316455 |
| 154 | *NEURL3* | 3.440935065 | 0.044805386 | 0.279257143 | 0.728431059 |
| 77 | *FABP3* | 3.496935065 | 0.044801389 | 0.515197619 | 0.895683598 |
| 100 | *GTF2F1* | 3.337454545 | 0.044737758 | 1.417985498 | 0.386737396 |
| 109 | *ILF2* | 3.285506494 | 0.044597426 | 2.020621482 | 0.386737396 |
| 98 | *GRIN2B* | 3.496935065 | 0.042685358 | 0.382154185 | 0.895683598 |
| 199 | *RESP18* | 3.376 | 0.040771081 | 1.748274085 | 0.728431059 |
| 47 | *CFD* | 3.513974026 | 0.040216121 | 0.466737734 | 0.975275022 |
| 187 | *PSMB10* | 3.531428571 | 0.038965957 | 0.623819048 | 0.945473238 |
| 234 | *STYXL1* | 3.578493506 | 0.037781463 | 0.537477656 | 0.999322301 |
| 74 | *EPS8L2* | 3.152 | 0.037200641 | 4.31973794 | 0.050982923 |
| 236 | *SYT3* | 3.462441558 | 0.037019534 | 0.259347619 | 0.822921164 |
| 14 | *ANKS6* | 3.475428571 | 0.036312556 | 1.110540837 | 0.822921164 |
| 242 | *THOP1* | 3.513974026 | 0.035850256 | 1.361304029 | 0.975275022 |
| 167 | *PCBD1* | 3.539948052 | 0.035083039 | 0.487071429 | 0.975275022 |
| 6 | *ADCY1* | 3.436467532 | 0.034891559 | 1.081970448 | 0.822921164 |
| 267 | *ZEB2* | 3.350441558 | 0.03475546 | 1.296145116 | 0.386737396 |
| 103 | *HCP5* | 3.246545455 | 0.034472298 | 6.384469353 | 0.284316455 |
| 110 | *IPO5* | 3.397922078 | 0.033723657 | 1.020930159 | 0.500803679 |
| 149 | *NAE1* | 3.496935065 | 0.032138095 | 0.493304762 | 0.895683598 |
| 239 | *TBC1D1* | 3.526961039 | 0.031470889 | 0.216965801 | 0.975275022 |
| 160 | *NPHP4* | 3.389402597 | 0.031421108 | 0.440759524 | 0.284316455 |

| | | | | | |
|---|---|---|---|---|---|
| 50 | *COG1* | 3.531428571 | 0.031402286 | 0.323264286 | 0.945473238 |
| 111 | *IRF2BP1* | 3.376415584 | 0.030938048 | 0.753656349 | 0.386737396 |
| 72 | *ELOVL6* | 3.522493506 | 0.030674603 | 0.442687179 | 0.990559909 |
| 82 | *FBXL18* | 3.061922078 | 0.029963277 | 3.630166305 | 0.001006504 |
| 127 | *MALL* | 3.406441558 | 0.029738333 | 1.334194517 | 0.618030488 |
| 240 | *TBC1D2B* | 3.483948052 | 0.029470966 | 0.610923413 | 0.895683598 |
| 181 | *POLR1A* | 3.380467532 | 0.029383139 | 1.184684127 | 0.500803679 |
| 79 | *FAM160B1* | 3.556987013 | 0.027677 | 0.347109957 | 0.997090681 |
| 229 | *SSSCA1* | 3.414961039 | 0.027516364 | 1.127553247 | 0.728431059 |
| 64 | *DNAJC30* | 3.483948052 | 0.027123611 | 0.311259524 | 0.895683598 |
| 164 | *OAZ2* | 3.505454545 | 0.026523333 | 0.579832353 | 0.945473238 |
| 156 | *NFS1* | 3.195012987 | 0.025955109 | 8.298323982 | 0.084167123 |
| 182 | *POLR2E* | 3.484363636 | 0.025453157 | 0.2425 | 0.618030488 |
| 224 | *SPIRE1* | 3.328935065 | 0.025041991 | 1.937637396 | 0.284316455 |
| 40 | *CCNYL1* | 3.509922078 | 0.024433384 | 0.465033333 | 0.895683598 |
| 34 | *C9orf16* | 3.427948052 | 0.023631763 | 1.294925224 | 0.728431059 |
| 22 | *ARPC2* | 3.419012987 | 0.023363443 | 2.40677094 | 0.895683598 |
| 95 | *GNPDA2* | 3.483948052 | 0.023282716 | 0.578403968 | 0.895683598 |
| 266 | *YWHAZ* | 3.535480519 | 0.023058333 | 0.356014286 | 0.990559909 |
| 257 | *UBE2B* | 3.505454545 | 0.018644667 | 0.350111255 | 0.945473238 |
| 185 | *PQLC1* | 3.565922078 | 0.018642113 | 0.134470635 | 0.975275022 |
| 166 | *PARD3B* | 3.561454545 | 0.015638232 | 0.104004762 | 0.990559909 |
| 175 | *PIGF* | 3.470961039 | 0.015424671 | 0.788087179 | 0.895683598 |
| 259 | *UBE2L3* | 3.578493506 | 0.014651515 | 0.394686946 | 0.999322301 |
| 104 | *HEMK1* | 3.470961039 | 0.013880999 | 0.451690476 | 0.895683598 |

| | | | | | |
|---|---|---|---|---|---|
| 146 | *MRPS33* | 3.475428571 | 0.013566919 | 0.54730873 | 0.822921164 |
| 5 | *ADAMTS10* | 3.496935065 | 0.012912044 | 0.369015873 | 0.895683598 |
| 142 | *MMGT1* | 3.518441558 | 0.01271708 | 0.183406926 | 0.945473238 |
| 44 | *CDK5R2* | 3.500987013 | 0.01160899 | 0.521683766 | 0.975275022 |
| 60 | *DEF8* | 3.587428571 | 0.011566667 | 0.085968254 | 0.990559909 |
| 241 | *TCF7* | 3.513974026 | 0.011136389 | 0.501204762 | 0.975275022 |
| 153 | *NEK9* | 3.561454545 | 0.009756211 | 0.113053968 | 0.990559909 |
| 179 | *PLD3* | 3.500987013 | 0.008564726 | 0.690490476 | 0.975275022 |
| 225 | *SPTAN1* | 3.264 | 0.00818287 | 1.89714062 | 0.284316455 |
| 41 | *CCT6B* | 3.539948052 | 0.008091683 | 0.561760101 | 0.975275022 |
| 161 | *NPPB* | 3.591480519 | 0.007531746 | 0.10818254 | 0.999322301 |
| 268 | *ZER1* | 3.595948052 | 0.007387619 | 0.072007143 | 0.997090681 |
| 122 | *LRP1* | 3.414961039 | 0.006846595 | 1.05293779 | 0.500803679 |
| 226 | *SRPK2* | 3.574441558 | 0.006672123 | 0.113206349 | 0.990559909 |
| 173 | *PEX11A* | 3.526961039 | 0.00652346 | 0.586948535 | 0.975275022 |
| 244 | *TMEM191A* | 3.436467532 | 0.006013605 | 0.812136652 | 0.822921164 |
| 37 | *CAND2* | 3.604467532 | 0.006 | 0.10602381 | 0.999322301 |
| 45 | *CEP19* | 3.561454545 | 0.005953125 | 0.468474747 | 0.990559909 |
| 253 | *TRUB1* | 3.582961039 | 0.00572125 | 0.224930525 | 0.997090681 |
| 68 | *DYNLRB2* | 3.276987013 | 0.005624376 | 2.872435426 | 0.284316455 |
| 230 | *STARD9* | 3.414961039 | 0.003914968 | 0.626440981 | 0.728431059 |
| 129 | *MAP2* | 3.582961039 | 0.0036375 | 0.06625 | 0.997090681 |
| 67 | *DYNLRB1* | 3.587428571 | 0.003502906 | 0.114790476 | 0.990559909 |
| 107 | *IDI1* | 3.561454545 | 0.002753395 | 0.149321429 | 0.990559909 |
| 188 | *PSMD7* | 3.518441558 | 0.002338889 | 0.265579365 | 0.945473238 |

| | | | | | |
|---|---|---|---|---|---|
| 221 | *SORBS3* | 3.604467532 | 0.0015 | 0.018933333 | 0.999322301 |
| 19 | *ARIH2* | 3.384935065 | 0.001292244 | 1.490562879 | 0.500803679 |
| 46 | *CETN3* | 3.604467532 | 0.001181235 | 0.058290476 | 0.999322301 |
| 112 | *JAZF1* | 3.488415584 | 0.001176031 | 0.294414286 | 0.822921164 |
| 80 | *FAM174B* | 3.401974026 | 0.001128086 | 0.84283254 | 0.728431059 |
| 8 | *AFTPH* | 3.578493506 | 0.000697143 | 0.196419048 | 0.999322301 |
| 232 | *STX4* | 3.470961039 | 0.000571506 | 0.533316991 | 0.895683598 |
| 90 | *FZD5* | 3.453922078 | 0.000522727 | 0.185595022 | 0.728431059 |
| 56 | *DARS* | 3.569974026 | 0.000395062 | 0.146904762 | 0.997090681 |
| 102 | *HBA2* | 3.604467532 | 0 | 0.075066667 | 0.999322301 |
| 141 | *MLYCD* | 3.591480519 | 0 | 0.153616667 | 0.999322301 |
| 186 | *PRB1* | 3.578493506 | -9.09E-05 | 0.270149206 | 0.999322301 |
| 54 | *CSTB* | 3.526961039 | -0.000204017 | 0.328924908 | 0.975275022 |
| 178 | *PLBD2* | 3.556987013 | -0.000265556 | 0.179070635 | 0.997090681 |
| 203 | *RNF123* | 3.569974026 | -0.000334267 | 0.183081385 | 0.997090681 |
| 125 | *LRRN2* | 3.539948052 | -0.000906944 | 0.209391847 | 0.975275022 |
| 218 | *SLC38A2* | 3.505454545 | -0.001090287 | 0.42039127 | 0.945473238 |
| 9 | *AGPAT5* | 3.625974026 | -0.0015 | 0.010166667 | 0.999894045 |
| 113 | *JOSD1* | 3.505454545 | -0.001736526 | 0.355344444 | 0.945473238 |
| 87 | *FNBP4* | 3.414961039 | -0.002316153 | 4.338462607 | 0.728431059 |
| 271 | *ZNF415* | 3.526961039 | -0.002813034 | 0.235014286 | 0.975275022 |
| 202 | *RILPL1* | 3.371948052 | -0.003017885 | 0.56503254 | 0.500803679 |
| 217 | *SLC1A4* | 3.259948052 | -0.003501026 | 2.084823933 | 0.132556825 |
| 85 | *FLOT2* | 3.621506494 | -0.00365625 | 0.096571429 | 0.999991668 |
| 196 | *RAB12* | 3.535480519 | -0.004026667 | 0.6027 | 0.990559909 |

| | | | | | |
|---|---|---|---|---|---|
| 32 | *C7* | 3.393454545 | -0.004343622 | 0.704284921 | 0.618030488 |
| 91 | *G3BP1* | 3.470961039 | -0.00449688 | 0.519963492 | 0.895683598 |
| 33 | *C9orf116* | 3.388987013 | -0.004742539 | 1.038617971 | 0.728431059 |
| 25 | *ATP6V1G1* | 3.488415584 | -0.004825617 | 0.178759524 | 0.822921164 |
| 235 | *SYDE2* | 3.526961039 | -0.004905097 | 0.437671429 | 0.975275022 |
| 204 | *RNF208* | 3.380467532 | -0.005558111 | 1.02149884 | 0.618030488 |
| 39 | *CCND2* | 3.544 | -0.005788444 | 0.561921429 | 0.997090681 |
| 145 | *MRPL49* | 3.569974026 | -0.006008333 | 0.236266667 | 0.997090681 |
| 194 | *PWWP2B* | 3.625974026 | -0.006700247 | 0.076861905 | 0.999894045 |
| 168 | *PCDHAC2* | 3.505454545 | -0.006726599 | 0.379842063 | 0.945473238 |
| 3 | *ACTR3* | 3.513974026 | -0.007103419 | 3.414977273 | 0.975275022 |
| 211 | *SAMD15* | 3.561454545 | -0.007446667 | 0.354681457 | 0.990559909 |
| 184 | *PPP5C* | 3.595948052 | -0.007489899 | 0.114038095 | 0.997090681 |
| 93 | *GLT8D1* | 3.29038961 | -0.007894722 | 3.22910575 | 0.084167123 |
| 12 | *ANGPT2* | 3.401974026 | -0.007980818 | 1.047146465 | 0.728431059 |
| 114 | *JUND* | 3.557402597 | -0.008 | 0.192809524 | 0.945473238 |
| 176 | *PIGX* | 3.518441558 | -0.008611883 | 0.133551587 | 0.945473238 |
| 148 | *MYH3* | 3.492467532 | -0.009257633 | 0.686044084 | 0.945473238 |
| 252 | *TROVE2* | 3.501402597 | -0.010055952 | 1.580791209 | 0.822921164 |
| 213 | *SEC24D* | 3.569974026 | -0.01090963 | 0.39235368 | 0.997090681 |
| 55 | *CYFIP2* | 3.354493506 | -0.011373442 | 8.715244811 | 0.618030488 |
| 84 | *FHOD1* | 3.569974026 | -0.013421042 | 0.314206227 | 0.997090681 |
| 4 | *ACTR6* | 3.522909091 | -0.013762006 | 0.221927706 | 0.895683598 |
| 265 | *YWHAB* | 3.574441558 | -0.013857338 | 0.083711111 | 0.990559909 |
| 58 | *DCUN1D5* | 3.432 | -0.014237273 | 1.059405067 | 0.895683598 |

| | | | | | |
|---|---|---|---|---|---|
| 195 | *RAB11FIP2* | 3.414961039 | -0.015372381 | 0.682251227 | 0.728431059 |
| 43 | *CDH4* | 3.449454545 | -0.015455503 | 0.819182107 | 0.822921164 |
| 71 | *EIF1B* | 3.492467532 | -0.015501667 | 0.460776696 | 0.945473238 |
| 150 | *NDFIP1* | 3.548467532 | -0.016783463 | 0.169777778 | 0.990559909 |
| 83 | *FCGBP* | 3.492467532 | -0.016790741 | 0.553677489 | 0.945473238 |
| 123 | *LRP3* | 3.410493506 | -0.018302918 | 0.789828087 | 0.822921164 |
| 158 | *NOS1* | 3.238441558 | -0.019107689 | 1.740705556 | 0.050982923 |
| 16 | *APBA1* | 3.604467532 | -0.0195572 | 0.112990476 | 0.999322301 |
| 2 | *ACACA* | 3.440935065 | -0.02076358 | 1.017230952 | 0.728431059 |
| 13 | *ANKIB1* | 3.591480519 | -0.022881944 | 0.148 | 0.999322301 |
| 69 | *EEF1A2* | 3.612987013 | -0.02413838 | 0.303418651 | 0.999322301 |
| 15 | *AP2S1* | 3.436467532 | -0.024944286 | 0.660763492 | 0.822921164 |
| 133 | *MAX* | 3.324467532 | -0.025657926 | 1.253886919 | 0.386737396 |
| 137 | *METRN* | 3.488 | -0.026117246 | 1.114704271 | 0.975275022 |
| 214 | *SERTAD3* | 3.45838961 | -0.033719899 | 0.38289596 | 0.618030488 |
| 88 | *FOXP4* | 3.539948052 | -0.034193333 | 0.295890476 | 0.975275022 |
| 115 | *KCNQ2* | 3.453506494 | -0.035541563 | 1.122837063 | 0.945473238 |
| 89 | *FRMD8* | 3.143480519 | -0.040529807 | 2.216330517 | 0.029497237 |
| 23 | *ASTN2* | 3.427948052 | -0.044957037 | 0.347163492 | 0.728431059 |
| 92 | *GFOD1* | 3.350025974 | -0.050852953 | 4.759001099 | 0.728431059 |
| 256 | *TTPAL* | 3.393454545 | -0.054603737 | 2.640047824 | 0.618030488 |
| 147 | *MT1A* | 3.246545455 | -0.055814708 | 5.11177785 | 0.386737396 |
| 208 | *RPL7L1* | 3.143896104 | -0.056387522 | 1.675412446 | 0.004395841 |
| 250 | *TRIM4* | 3.457974026 | -0.059100481 | 0.819887085 | 0.895683598 |
| 52 | *CPSF4* | 3.345974026 | -0.062425615 | 1.123289377 | 0.500803679 |

| | | | | | |
|---|---|---|---|---|---|
| 138 | *MICU1* | 3.496935065 | -0.077799383 | 0.459109524 | 0.895683598 |
| 20 | *ARL2* | 3.414545455 | -0.083852475 | 1.323656643 | 0.945473238 |
| 261 | *VTA1* | 3.444987013 | -0.10294795 | 1.218147619 | 0.895683598 |
| 42 | *CD34* | 3.436467532 | -0.113426481 | 0.55371337 | 0.822921164 |
| 260 | *VAC14* | 3.548467532 | -0.169795056 | 1.90174359 | 0.990559909 |

**Table S30. Importance (Dominance) of each gene in the random forest model for caudal anterior cingulate cortex (cauACC).** Mean_min_depth represents the average minimum depth at which a variable is used as a split node across all trees, with lower values indicating higher importance as the variable is used earlier in the decision process. IncrMSE, % (Increase in Mean Squared Error) quantifies the relative increase in prediction error when the variable is permuted, reflecting its contribution to the model's predictive accuracy; higher values signify greater importance. Node_purity_increase measures the improvement in node purity (e.g., Gini index or variance reduction) achieved by splitting on the variable, where larger values indicate a stronger impact on classification or regression performance. Finally, the P value assesses the statistical significance of a variable's importance, typically derived from permutation tests, with lower values suggesting that the variable's contribution is unlikely to be due to random chance. This presentation was in descending order by the IncrMSE.

| Entrez ID | Gene Symbol | Mean_min_depth | IncrMSE, % | node_purity_increase | P value |
|---|---|---|---|---|---|
| 63 | *DLL3* | 0.795862069 | 6.356103476 | 84.68112697 | 7.28E-17 |
| 126 | *LYNX1* | 0.672413793 | 4.352442091 | 63.35850735 | 1.60E-24 |
| 247 | *TMSB10* | 1.668689655 | 3.241346008 | 52.38646925 | 1.05E-08 |
| 157 | *NNAT* | 1.671310345 | 3.156511117 | 46.76660213 | 9.61E-10 |
| 272 | *ZNF599* | 1.727034483 | 3.014573731 | 46.08992794 | 3.22E-09 |
| 210 | *S100A1* | 2.146206897 | 2.241634245 | 42.38218391 | 1.81E-05 |
| 273 | *ZNF821* | 2.440689655 | 2.221565123 | 27.83537121 | 0.001370367 |
| 124 | *LRRC61* | 1.972413793 | 2.134814673 | 41.46974916 | 3.33E-08 |
| 116 | *KIF19* | 2.077241379 | 2.028199609 | 44.29051684 | 2.51E-06 |
| 207 | *RPL24* | 2.361103448 | 1.518447743 | 32.394446 | 0.000621142 |
| 70 | *EHD3* | 2.627724138 | 1.438684003 | 22.01024166 | 0.011546289 |
| 121 | *LPIN1* | 2.078482759 | 1.304886729 | 21.38983171 | 9.61E-10 |
| 27 | *BASP1* | 2.56537931 | 1.211626788 | 27.49808554 | 0.002906131 |

| | | | | | |
|---|---|---|---|---|---|
| 97 | *GRAMD1A* | 2.676827586 | 1.184274995 | 24.025127 | 0.06567788 |
| 143 | *MPZL1* | 1.998896552 | 1.179520569 | 20.28007712 | 9.61E-10 |
| 165 | *PAFAH1B3* | 2.342482759 | 1.121453249 | 24.65187008 | 6.85E-06 |
| 73 | *EME1* | 1.898068966 | 1.114527461 | 14.07518056 | 5.91E-12 |
| 206 | *RPL13A* | 2.617103448 | 1.006193514 | 13.72325071 | 0.005916289 |
| 1 | *AATK* | 2.755034483 | 0.849122432 | 14.32790095 | 0.005916289 |
| 109 | *ILF2* | 2.737793103 | 0.736907355 | 12.56981494 | 0.005916289 |
| 134 | *MCHR1* | 2.58262069 | 0.734189437 | 24.89439737 | 0.005916289 |
| 65 | *DSTYK* | 2.679448276 | 0.599251489 | 17.16419904 | 0.011546289 |
| 24 | *ATF5* | 2.931448276 | 0.566211242 | 11.03040798 | 0.02157156 |
| 245 | *TMEM199* | 3.053517241 | 0.543266433 | 10.53027971 | 0.243029624 |
| 177 | *PITPNM2* | 2.952689655 | 0.534678095 | 14.8351195 | 0.06567788 |
| 53 | *CREB3L3* | 2.491034483 | 0.466844912 | 11.79122653 | 1.81E-05 |
| 261 | *VTA1* | 3.023034483 | 0.445477647 | 14.1969725 | 0.451805576 |
| 135 | *MED30* | 2.699310345 | 0.437239105 | 12.96439722 | 0.001370367 |
| 201 | *RIIAD1* | 2.922206897 | 0.432219059 | 11.4998142 | 0.16514715 |
| 120 | *LIMD2* | 3.092 | 0.392185171 | 11.6713157 | 0.451805576 |
| 248 | *TNFAIP8L3* | 2.932827586 | 0.362488818 | 16.6209826 | 0.243029624 |
| 21 | *ARMCX6* | 3.08537931 | 0.350042441 | 10.12405828 | 0.571035035 |
| 11 | *AMIGO1* | 3.036275862 | 0.341037803 | 8.958541668 | 0.16514715 |
| 258 | *UBE2E2* | 3.019034483 | 0.338570892 | 3.072314042 | 0.243029624 |
| 66 | *DTNA* | 2.824 | 0.330275709 | 4.013845429 | 0.005916289 |
| 2 | *ACACA* | 2.546758621 | 0.323343751 | 5.035583937 | 4.62E-05 |
| 159 | *NOS1AP* | 2.703310345 | 0.308800123 | 12.15929187 | 0.005916289 |
| 93 | *GLT8D1* | 3.109241379 | 0.303658389 | 11.03248142 | 0.451805576 |
| 92 | *GFOD1* | 3.060137931 | 0.297108681 | 6.674934979 | 0.16514715 |
| 17 | *AQP6* | 2.907586207 | 0.277310516 | 4.427729907 | 0.038525291 |

| | | | | | |
|---|---|---|---|---|---|
| 202 | *RILPL1* | 3.306896552 | 0.267722305 | 4.785363095 | 0.873281465 |
| 162 | *NRSN2* | 3.015034483 | 0.262856687 | 2.49086998 | 0.106735294 |
| 96 | *GPNMB* | 3.115862069 | 0.239654414 | 7.413991032 | 0.339918373 |
| 39 | *CCND2* | 2.828 | 0.23372981 | 3.798566989 | 0.02157156 |
| 55 | *CYFIP2* | 3.352 | 0.232952274 | 5.338904615 | 0.931811557 |
| 155 | *NEUROD2* | 3.292275862 | 0.210121416 | 1.378229949 | 0.571035035 |
| 190 | *PTDSS1* | 3.171586207 | 0.207081221 | 6.315493519 | 0.451805576 |
| 187 | *PSMB10* | 2.987172414 | 0.204470714 | 3.143467431 | 0.06567788 |
| 86 | *FMNL1* | 3.264413793 | 0.204394834 | 1.288098375 | 0.339918373 |
| 101 | *HBA1* | 3.261793103 | 0.19978353 | 5.962609078 | 0.790785238 |
| 57 | *DCHS1* | 3.379862069 | 0.188039838 | 4.536651169 | 0.968135316 |
| 46 | *CETN3* | 3.198068966 | 0.187026855 | 1.246884799 | 0.16514715 |
| 117 | *KRT17* | 3.313517241 | 0.18627518 | 4.120305873 | 0.790785238 |
| 200 | *RFXANK* | 3.257793103 | 0.184528341 | 2.054494012 | 0.571035035 |
| 243 | *THY1* | 3.007034483 | 0.17746231 | 3.581860865 | 0.011546289 |
| 26 | *ATP9B* | 3.36262069 | 0.169672293 | 2.467145599 | 0.968135316 |
| 169 | *PCGF2* | 3.36262069 | 0.168870014 | 4.546770909 | 0.968135316 |
| 106 | *HMG20A* | 3.386482759 | 0.167834497 | 1.017170563 | 0.931811557 |
| 174 | *PHF2* | 3.261793103 | 0.167495968 | 1.788245563 | 0.790785238 |
| 199 | *RESP18* | 3.251172414 | 0.167260468 | 1.197558208 | 0.687462757 |
| 103 | *HCP5* | 3.015034483 | 0.153989402 | 5.702156073 | 0.106735294 |
| 102 | *HBA2* | 3.317517241 | 0.153826123 | 5.119702079 | 0.931811557 |
| 61 | *DEPDC5* | 2.802758621 | 0.150692991 | 3.130757287 | 0.001370367 |
| 18 | *ARF3* | 3.017655172 | 0.149651567 | 1.950000991 | 0.02157156 |
| 22 | *ARPC2* | 3.407724138 | 0.145880432 | 3.622114935 | 0.987451177 |

| | | | | | |
|---|---|---|---|---|---|
| 130 | *MAP4K2* | 3.435586207 | 0.136732171 | 2.735971099 | 0.99600804 |
| 171 | *PDE9A* | 3.154344828 | 0.128055668 | 6.406808107 | 0.571035035 |
| 3 | *ACTR3* | 3.326758621 | 0.125399596 | 4.382590056 | 0.571035035 |
| 238 | *TARSL2* | 3.302896552 | 0.124947778 | 2.432243797 | 0.687462757 |
| 44 | *CDK5R2* | 2.83862069 | 0.124520658 | 3.601449681 | 0.038525291 |
| 35 | *CABYR* | 3.174206897 | 0.124021395 | 1.662333655 | 0.243029624 |
| 227 | *SRRT* | 3.105241379 | 0.11873205 | 3.422459921 | 0.243029624 |
| 220 | *SNCG* | 3.285655172 | 0.114209685 | 2.047219908 | 0.687462757 |
| 105 | *HLF* | 3.302896552 | 0.113346222 | 1.803729717 | 0.687462757 |
| 94 | *GNG5* | 2.924827586 | 0.112429002 | 2.977213636 | 0.038525291 |
| 77 | *FABP3* | 3.306896552 | 0.108815166 | 1.075974603 | 0.873281465 |
| 75 | *ETV5* | 3.255172414 | 0.104192259 | 4.545307363 | 0.873281465 |
| 47 | *CFD* | 3.011034483 | 0.104057257 | 1.496555628 | 0.038525291 |
| 180 | *PLEKHA1* | 3.184827586 | 0.095838827 | 4.2498142 | 0.339918373 |
| 241 | *TCF7* | 3.33737931 | 0.094845556 | 0.384554335 | 0.687462757 |
| 140 | *MLLT11* | 3.379862069 | 0.094519093 | 3.220175549 | 0.968135316 |
| 246 | *TMEM38A* | 3.272413793 | 0.09184853 | 3.078656441 | 0.873281465 |
| 228 | *SSFA2* | 3.452827586 | 0.091425687 | 1.871821713 | 0.99600804 |
| 269 | *ZKSCAN1* | 3.251172414 | 0.089937589 | 1.512520411 | 0.687462757 |
| 231 | *STAT2* | 3.34137931 | 0.085721033 | 2.307251349 | 0.873281465 |
| 133 | *MAX* | 3.375862069 | 0.085355333 | 2.585424805 | 0.873281465 |
| 249 | *TRADD* | 3.348 | 0.081111746 | 1.102212821 | 0.790785238 |
| 215 | *SF3A3* | 3.261793103 | 0.07930878 | 3.296607106 | 0.790785238 |
| 164 | *OAZ2* | 2.678068966 | 0.078033294 | 3.653779412 | 0.000270976 |
| 72 | *ELOVL6* | 3.264413793 | 0.07490887 | 0.875844322 | 0.339918373 |

| | | | | | |
|---|---|---|---|---|---|
| 100 | *GTF2F1* | 3.493931034 | 0.074817778 | 2.154320476 | 0.987451177 |
| 74 | *EPS8L2* | 3.255172414 | 0.074397444 | 5.549113794 | 0.873281465 |
| 226 | *SRPK2* | 3.369241379 | 0.072769394 | 1.004793001 | 0.931811557 |
| 49 | *CNOT1* | 3.223310345 | 0.072678173 | 1.601858561 | 0.571035035 |
| 141 | *MLYCD* | 2.639586207 | 0.071498912 | 3.027578355 | 4.62E-05 |
| 31 | *C16orf70* | 3.476689655 | 0.068340644 | 2.169031538 | 0.987451177 |
| 193 | *PTPRU* | 3.371862069 | 0.068144109 | 0.528625108 | 0.687462757 |
| 219 | *SLC38A3* | 3.302896552 | 0.067319718 | 0.829838939 | 0.687462757 |
| 48 | *CIDECP* | 3.320137931 | 0.066783444 | 0.901643723 | 0.687462757 |
| 112 | *JAZF1* | 2.806758621 | 0.065864694 | 3.075882018 | 0.002906131 |
| 191 | *PTPRM* | 2.886344828 | 0.064664856 | 3.342324675 | 0.011546289 |
| 262 | *WDR82* | 3.442206897 | 0.064059921 | 0.673662338 | 0.987451177 |
| 168 | *PCDHAC2* | 2.955310345 | 0.063258137 | 1.865210184 | 0.011546289 |
| 19 | *ARIH2* | 3.268413793 | 0.056573221 | 1.781421337 | 0.687462757 |
| 68 | *DYNLRB2* | 3.236551724 | 0.056381727 | 1.607284719 | 0.339918373 |
| 12 | *ANGPT2* | 3.316137931 | 0.054909021 | 0.735094827 | 0.451805576 |
| 99 | *GRK6* | 3.386482759 | 0.054445687 | 1.96179039 | 0.931811557 |
| 267 | *ZEB2* | 3.427586207 | 0.054235853 | 0.319184704 | 0.873281465 |
| 221 | *SORBS3* | 3.420965517 | 0.052809672 | 0.379530952 | 0.931811557 |
| 6 | *ADCY1* | 2.626344828 | 0.052178528 | 4.574108548 | 0.000113916 |
| 108 | *IDS* | 3.070758621 | 0.050268948 | 2.212093715 | 0.243029624 |
| 192 | *PTPRS* | 3.302896552 | 0.048610287 | 0.702864069 | 0.687462757 |
| 218 | *SLC38A2* | 3.348 | 0.047888013 | 0.519228355 | 0.790785238 |
| 9 | *AGPAT5* | 3.281655172 | 0.047415551 | 0.958221623 | 0.451805576 |
| 59 | *DDX51* | 3.275034483 | 0.045842378 | 2.979786841 | 0.451805576 |

| | | | | | |
|---|---|---|---|---|---|
| 62 | *DLG4* | 3.34137931 | 0.045692354 | 0.875559524 | 0.873281465 |
| 34 | *C9orf16* | 2.959310345 | 0.042309444 | 3.105512876 | 0.038525291 |
| 179 | *PLD3* | 3.448827586 | 0.040050741 | 0.634107143 | 0.968135316 |
| 216 | *SLC10A4* | 3.139724138 | 0.039361828 | 1.615306349 | 0.243029624 |
| 203 | *RNF123* | 3.515172414 | 0.0392 | 0.172433333 | 0.999039582 |
| 197 | *RAMP1* | 3.212689655 | 0.037177525 | 1.400561905 | 0.451805576 |
| 123 | *LRP3* | 3.386482759 | 0.036031328 | 0.433380303 | 0.931811557 |
| 232 | *STX4* | 3.219310345 | 0.035758241 | 1.107401604 | 0.339918373 |
| 214 | *SERTAD3* | 3.038896552 | 0.03444078 | 3.106183744 | 0.06567788 |
| 85 | *FLOT2* | 3.532413793 | 0.032753788 | 0.288478571 | 0.999039582 |
| 13 | *ANKIB1* | 3.528413793 | 0.032585354 | 0.103690476 | 0.987451177 |
| 128 | *MANEAL* | 3.493931034 | 0.032489946 | 0.244428571 | 0.987451177 |
| 213 | *SEC24D* | 3.292275862 | 0.03232716 | 0.878779709 | 0.571035035 |
| 144 | *MRPL34* | 3.285655172 | 0.031831423 | 0.940687729 | 0.687462757 |
| 237 | *TADA2B* | 3.167586207 | 0.028981855 | 2.520886344 | 0.243029624 |
| 33 | *C9orf116* | 3.206068966 | 0.027945098 | 1.493672386 | 0.451805576 |
| 198 | *RASSF7* | 3.459448276 | 0.027325476 | 0.266285931 | 0.987451177 |
| 217 | *SLC1A4* | 3.042896552 | 0.025086066 | 1.672669841 | 0.16514715 |
| 41 | *CCT6B* | 3.382482759 | 0.025076444 | 0.316742063 | 0.790785238 |
| 69 | *EEF1A2* | 3.386482759 | 0.024451618 | 0.506256061 | 0.873281465 |
| 250 | *TRIM4* | 3.33737931 | 0.023947338 | 0.661744949 | 0.687462757 |
| 139 | *MKRN3* | 3.178206897 | 0.023677421 | 4.465669581 | 0.339918373 |
| 4 | *ACTR6* | 3.539034483 | 0.023465606 | 0.098677778 | 0.99600804 |
| 7 | *ADCY5* | 3.184827586 | 0.02342485 | 1.397385498 | 0.339918373 |
| 255 | *TTC9B* | 3.167586207 | 0.023180376 | 1.648942569 | 0.339918373 |

| | | | | | |
|---|---|---|---|---|---|
| 212 | *SCRN1* | 3.223310345 | 0.02254837 | 1.393007018 | 0.571035035 |
| 156 | *NFS1* | 3.330758621 | 0.022055952 | 0.571032179 | 0.790785238 |
| 50 | *COG1* | 3.414344828 | 0.02196201 | 0.807456349 | 0.968135316 |
| 71 | *EIF1B* | 3.448827586 | 0.021552169 | 0.22587619 | 0.968135316 |
| 67 | *DYNLRB1* | 3.382482759 | 0.019996465 | 0.377069841 | 0.790785238 |
| 64 | *DNAJC30* | 3.386482759 | 0.019524444 | 0.848066089 | 0.931811557 |
| 271 | *ZNF415* | 3.521793103 | 0.018785859 | 0.109851732 | 0.99600804 |
| 127 | *MALL* | 3.289655172 | 0.018722937 | 1.329311446 | 0.873281465 |
| 251 | *TRIO* | 3.369241379 | 0.01839972 | 0.82132381 | 0.931811557 |
| 16 | *APBA1* | 3.504551724 | 0.01818625 | 0.325617424 | 0.99600804 |
| 29 | *BOP1* | 3.34137931 | 0.01627947 | 2.243656353 | 0.873281465 |
| 79 | *FAM160B1* | 3.515172414 | 0.015688333 | 0.294584127 | 0.999039582 |
| 252 | *TROVE2* | 3.219310345 | 0.01469784 | 0.464973449 | 0.339918373 |
| 208 | *RPL7L1* | 3.466068966 | 0.01455 | 0.136160317 | 0.968135316 |
| 185 | *PQLC1* | 3.403724138 | 0.01441418 | 0.226983333 | 0.931811557 |
| 14 | *ANKS6* | 3.302896552 | 0.013746936 | 0.869348612 | 0.687462757 |
| 30 | *C11orf1* | 3.504551724 | 0.013697778 | 0.192556349 | 0.99600804 |
| 36 | *CACNB1* | 3.309517241 | 0.013611065 | 0.611000794 | 0.571035035 |
| 263 | *WNK2* | 3.233931034 | 0.012564444 | 1.804941533 | 0.687462757 |
| 265 | *YWHAB* | 3.420965517 | 0.01212 | 0.169448124 | 0.931811557 |
| 131 | *MAP6* | 3.459448276 | 0.011697222 | 0.247175325 | 0.987451177 |
| 236 | *SYT3* | 3.414344828 | 0.010613757 | 0.409382684 | 0.968135316 |
| 114 | *JUND* | 3.326758621 | 0.010088757 | 0.463546104 | 0.571035035 |
| 51 | *COPG2* | 3.126482759 | 0.009972467 | 4.360987726 | 0.339918373 |
| 82 | *FBXL18* | 3.334758621 | 0.009803896 | 3.428099841 | 0.931811557 |

| | | | | | |
|---|---|---|---|---|---|
| 205 | *RNFT2* | 3.33737931 | 0.008290415 | 0.596416667 | 0.687462757 |
| 45 | *CEP19* | 3.420965517 | 0.008081995 | 0.648687764 | 0.931811557 |
| 256 | *TTPAL* | 3.560275862 | 0.007919192 | 0.071083333 | 0.999844836 |
| 23 | *ASTN2* | 3.493931034 | 0.0078945 | 0.192311111 | 0.987451177 |
| 194 | *PWWP2B* | 3.420965517 | 0.007754969 | 0.372026984 | 0.931811557 |
| 80 | *FAM174B* | 3.393103448 | 0.007749369 | 0.335956044 | 0.873281465 |
| 60 | *DEF8* | 3.455448276 | 0.007067 | 0.225469048 | 0.931811557 |
| 43 | *CDH4* | 3.438206897 | 0.006921333 | 0.596616667 | 0.931811557 |
| 132 | *MARK4* | 3.504551724 | 0.006692346 | 0.238620635 | 0.99600804 |
| 223 | *SP100* | 3.455448276 | 0.005636364 | 0.271561255 | 0.931811557 |
| 254 | *TTC3P1* | 3.257793103 | 0.005411145 | 1.460836369 | 0.571035035 |
| 110 | *IPO5* | 3.268413793 | 0.005339375 | 1.009252283 | 0.687462757 |
| 153 | *NEK9* | 3.528413793 | 0.004458923 | 0.058811111 | 0.987451177 |
| 20 | *ARL2* | 3.386482759 | 0.003855884 | 0.509596825 | 0.931811557 |
| 150 | *NDFIP1* | 3.549655172 | 0.003594192 | 0.108183333 | 0.999039582 |
| 240 | *TBC1D2B* | 3.438206897 | 0.003292328 | 0.235061039 | 0.931811557 |
| 152 | *NDUFS4* | 3.455448276 | 0.003161806 | 0.227792641 | 0.931811557 |
| 111 | *IRF2BP1* | 3.35862069 | 0.0027975 | 0.574671429 | 0.873281465 |
| 146 | *MRPS33* | 3.386482759 | 0.00278127 | 0.285215657 | 0.931811557 |
| 244 | *TMEM191A* | 3.272413793 | 0.002500611 | 1.720179257 | 0.873281465 |
| 173 | *PEX11A* | 3.511172414 | 0.002044444 | 0.077680952 | 0.987451177 |
| 172 | *PDK2* | 3.386482759 | 0.001990159 | 0.679792735 | 0.931811557 |
| 137 | *METRN* | 3.448827586 | 0.001971852 | 0.260378571 | 0.968135316 |
| 230 | *STARD9* | 3.448827586 | 0.0014375 | 0.265369048 | 0.968135316 |
| 239 | *TBC1D1* | 3.045517241 | 0.001304131 | 1.747805922 | 0.02157156 |

| | | | | | |
|---|---|---|---|---|---|
| 8 | *AFTPH* | 3.60537931 | 0 | 0.001666667 | 0.999987387 |
| 32 | *C7* | 3.476689655 | 0 | 0.118980952 | 0.987451177 |
| 129 | *MAP2* | 3.539034483 | 0 | 0.10752381 | 0.99600804 |
| 186 | *PRB1* | 3.577517241 | 0 | 0.036457143 | 0.999844836 |
| 253 | *TRUB1* | 3.515172414 | 0 | 0.293641558 | 0.999039582 |
| 183 | *PPDPF* | 3.438206897 | -0.000287556 | 0.620909524 | 0.931811557 |
| 125 | *LRRN2* | 3.504551724 | -0.000430769 | 0.596761233 | 0.99600804 |
| 178 | *PLBD2* | 3.532413793 | -0.000552 | 0.205292641 | 0.999039582 |
| 40 | *CCNYL1* | 3.382482759 | -0.001230327 | 0.321438095 | 0.790785238 |
| 91 | *G3BP1* | 3.375862069 | -0.001248 | 0.425940049 | 0.873281465 |
| 204 | *RNF208* | 3.539034483 | -0.001508838 | 0.202628571 | 0.99600804 |
| 15 | *AP2S1* | 3.511172414 | -0.001791667 | 0.168206349 | 0.987451177 |
| 25 | *ATP6V1G1* | 3.448827586 | -0.002991667 | 0.236011111 | 0.968135316 |
| 211 | *SAMD15* | 3.414344828 | -0.003541667 | 0.501335137 | 0.968135316 |
| 233 | *STXBP1* | 3.188827586 | -0.003739185 | 1.271517172 | 0.571035035 |
| 95 | *GNPDA2* | 3.594758621 | -0.004181818 | 0.006 | 0.999844836 |
| 175 | *PIGF* | 3.348 | -0.004702679 | 1.006461294 | 0.790785238 |
| 181 | *POLR1A* | 3.233931034 | -0.004835039 | 1.25500522 | 0.687462757 |
| 5 | *ADAMTS10* | 3.455448276 | -0.004929745 | 0.388580586 | 0.931811557 |
| 76 | *EVL* | 3.487310345 | -0.005067481 | 0.351846753 | 0.99600804 |
| 54 | *CSTB* | 3.483310345 | -0.005111111 | 0.123384921 | 0.968135316 |
| 158 | *NOS1* | 3.375862069 | -0.005399391 | 0.756718853 | 0.873281465 |
| 88 | *FOXP4* | 3.296275862 | -0.005476268 | 1.493339683 | 0.790785238 |
| 81 | *FARSB* | 3.438206897 | -0.005489815 | 0.301720635 | 0.931811557 |
| 195 | *RAB11FIP2* | 3.399724138 | -0.006560354 | 0.222980952 | 0.790785238 |

| | | | | | |
|---|---|---|---|---|---|
| 119 | *LEMD2* | 3.476689655 | -0.007356554 | 0.329706349 | 0.987451177 |
| 260 | *VAC14* | 3.549655172 | -0.007506173 | 0.053434921 | 0.999039582 |
| 160 | *NPHP4* | 3.302896552 | -0.007509292 | 1.244980736 | 0.687462757 |
| 163 | *NUTF2* | 3.493931034 | -0.007921875 | 0.132992063 | 0.987451177 |
| 87 | *FNBP4* | 3.223310345 | -0.008109416 | 1.148298268 | 0.571035035 |
| 170 | *PCK2* | 3.476689655 | -0.008792222 | 0.187768254 | 0.987451177 |
| 235 | *SYDE2* | 3.208689655 | -0.009028162 | 0.950263112 | 0.243029624 |
| 136 | *MEST* | 3.253793103 | -0.009343086 | 1.129292857 | 0.339918373 |
| 264 | *YIF1B* | 3.397103448 | -0.009707346 | 1.035054119 | 0.968135316 |
| 38 | *CAPRIN2* | 3.309517241 | -0.009813485 | 0.737189033 | 0.571035035 |
| 184 | *PPP5C* | 3.532413793 | -0.00987 | 0.228244444 | 0.999039582 |
| 89 | *FRMD8* | 3.448827586 | -0.01013963 | 0.129587302 | 0.968135316 |
| 122 | *LRP1* | 3.442206897 | -0.010755354 | 0.373348629 | 0.987451177 |
| 151 | *NDUFA9* | 3.35862069 | -0.010807716 | 0.992226984 | 0.790785238 |
| 42 | *CD34* | 3.511172414 | -0.010861208 | 0.076827778 | 0.987451177 |
| 78 | *FAHD1* | 3.493931034 | -0.011534882 | 0.222306349 | 0.968135316 |
| 189 | *PSME3* | 3.539034483 | -0.012114626 | 0.074213636 | 0.99600804 |
| 182 | *POLR2E* | 3.455448276 | -0.015676229 | 0.232539683 | 0.931811557 |
| 115 | *KCNQ2* | 3.487310345 | -0.016715909 | 0.371087302 | 0.99600804 |
| 234 | *STYXL1* | 3.493931034 | -0.017510101 | 0.227381868 | 0.987451177 |
| 52 | *CPSF4* | 3.403724138 | -0.017604444 | 0.545040188 | 0.931811557 |
| 37 | *CAND2* | 3.448827586 | -0.018496667 | 0.411359524 | 0.968135316 |
| 83 | *FCGBP* | 3.448827586 | -0.018638556 | 0.344061255 | 0.968135316 |
| 176 | *PIGX* | 3.399724138 | -0.018681688 | 0.353290476 | 0.790785238 |
| 84 | *FHOD1* | 3.386482759 | -0.019407923 | 0.485016545 | 0.931811557 |

| | | | | | |
|---|---|---|---|---|---|
| 224 | *SPIRE1* | 2.983172414 | -0.020979523 | 3.052330204 | 0.02157156 |
| 266 | *YWHAZ* | 3.424965517 | -0.021497352 | 0.546686203 | 0.987451177 |
| 149 | *NAE1* | 3.160965517 | -0.022077668 | 1.859889993 | 0.451805576 |
| 142 | *MMGT1* | 3.500551724 | -0.02222716 | 0.114528571 | 0.968135316 |
| 225 | *SPTAN1* | 3.150344828 | -0.022384098 | 2.420210958 | 0.339918373 |
| 188 | *PSMD7* | 3.560275862 | -0.022410101 | 0.582988889 | 0.999844836 |
| 107 | *IDI1* | 3.470068966 | -0.022600778 | 0.335359829 | 0.99600804 |
| 138 | *MICU1* | 3.511172414 | -0.02522 | 0.108957143 | 0.987451177 |
| 90 | *FZD5* | 3.375862069 | -0.025434745 | 0.729521429 | 0.873281465 |
| 270 | *ZNF233* | 3.459448276 | -0.025768581 | 0.529081638 | 0.987451177 |
| 229 | *SSSCA1* | 3.133103448 | -0.026878975 | 7.094439746 | 0.339918373 |
| 118 | *KRTCAP2* | 3.348 | -0.027733333 | 0.651724242 | 0.790785238 |
| 56 | *DARS* | 3.271034483 | -0.029795981 | 0.489345238 | 0.243029624 |
| 222 | *SOS1* | 3.386482759 | -0.029943611 | 0.980436386 | 0.931811557 |
| 104 | *HEMK1* | 3.382482759 | -0.030262833 | 0.159094444 | 0.790785238 |
| 242 | *THOP1* | 3.410344828 | -0.030272737 | 0.34883254 | 0.790785238 |
| 113 | *JOSD1* | 3.257793103 | -0.032679167 | 1.430109524 | 0.571035035 |
| 10 | *AHCYL2* | 3.285655172 | -0.036273591 | 0.839276929 | 0.687462757 |
| 166 | *PARD3B* | 3.504551724 | -0.0373 | 0.137607143 | 0.99600804 |
| 196 | *RAB12* | 3.302896552 | -0.03795042 | 0.567223504 | 0.687462757 |
| 167 | *PCBD1* | 3.202068966 | -0.040192762 | 1.232223687 | 0.243029624 |
| 268 | *ZER1* | 3.244551724 | -0.042796667 | 1.434246212 | 0.790785238 |
| 145 | *MRPL49* | 3.442206897 | -0.044179 | 1.997898557 | 0.987451177 |
| 154 | *NEURL3* | 3.466068966 | -0.04556381 | 0.301795288 | 0.968135316 |
| 98 | *GRIN2B* | 3.208689655 | -0.047272312 | 0.738553175 | 0.243029624 |

| | | | | | |
|---|---|---|---|---|---|
| 209 | *RTF1* | 3.253793103 | -0.049907556 | 0.465591758 | 0.339918373 |
| 58 | *DCUN1D5* | 3.483310345 | -0.050677778 | 0.259118132 | 0.968135316 |
| 257 | *UBE2B* | 3.403724138 | -0.05312658 | 0.884612759 | 0.931811557 |
| 148 | *MYH3* | 3.386482759 | -0.054431259 | 2.301048139 | 0.931811557 |
| 147 | *MT1A* | 3.334758621 | -0.054726647 | 1.302104401 | 0.931811557 |
| 28 | *BCAN* | 3.154344828 | -0.059983675 | 2.047665554 | 0.451805576 |
| 161 | *NPPB* | 3.261793103 | -0.081105854 | 1.715280428 | 0.790785238 |
| 259 | *UBE2L3* | 3.386482759 | -0.093874424 | 1.892764843 | 0.931811557 |

**Table S31. Importance (Dominance) of each gene in the random forest model for orbital frontal cortex (OFC).** Mean_min_depth represents the average minimum depth at which a variable is used as a split node across all trees, with lower values indicating higher importance as the variable is used earlier in the decision process. IncrMSE, % (Increase in Mean Squared Error) quantifies the relative increase in prediction error when the variable is permuted, reflecting its contribution to the model's predictive accuracy; higher values signify greater importance. Node_purity_increase measures the improvement in node purity (e.g., Gini index or variance reduction) achieved by splitting on the variable, where larger values indicate a stronger impact on classification or regression performance. Finally, the P value assesses the statistical significance of a variable's importance, typically derived from permutation tests, with lower values suggesting that the variable's contribution is unlikely to be due to random chance. This presentation was in descending order by the IncrMSE.

| Entrez ID | Gene Symbol | Mean_min_depth | IncrMSE, % | node_purity_increase | P value |
|---|---|---|---|---|---|
| 157 | *NNAT* | 1.58753684210526 | 6.9990305511248 | 139.311689567184 | 2.88019166047824e-21 |
| 273 | *ZNF821* | 2.21894736842105 | 4.71626229752718 | 97.1074530934752 | 1.47377836051495e-12 |
| 220 | *SNCG* | 2.48353684210526 | 4.50917928065761 | 81.2891219104597 | 2.16401947522466e-10 |
| 143 | *MPZL1* | 0.926315789473684 | 4.38141961537391 | 83.7162280144863 | 2.66322111830972e-53 |

| | | | | | |
|---|---|---|---|---|---|
| 1 | *AATK* | 2.168 | 2.8457475734127 | 43.9288377326664 | 1.22742452828412e-22 |
| 165 | *PAFAH1B3* | 2.47452631578947 | 2.69614604079408 | 65.9673145970644 | 7.10638370272217e-10 |
| 24 | *ATF5* | 2.84884210526316 | 2.42790078146237 | 49.2620827537959 | 3.7957575255518e-06 |
| 27 | *BASP1* | 3.15402105263158 | 1.92936899524087 | 36.6075146226323 | 0.00586158103804021 |
| 228 | *SSFA2* | 3.24572631578947 | 1.54749553198653 | 23.8083148515228 | 0.0201423318626269 |
| 74 | *EPS8L2* | 3.13145263157895 | 1.48206575789026 | 38.7944252291826 | 0.0110840307247928 |
| 21 | *ARMCX6* | 2.912 | 1.39893193728956 | 44.4066566595771 | 2.44532063529431e-05 |
| 155 | *NEUROD2* | 2.8384 | 1.31957725689788 | 20.5670500835766 | 2.16912113543905e-08 |
| 171 | *PDE9A* | 3.27124210526316 | 1.28557933429688 | 30.9905144930233 | 0.144251494432929 |
| 210 | *S100A1* | 3.39452631578947 | 1.15822420216049 | 17.5627004931343 | 0.211337646582481 |
| 57 | *DCHS1* | 2.94509473684211 | 1.05175639423077 | 25.1225168258041 | 3.7957575255518e-06 |
| 124 | *LRRC61* | 3.26526315789474 | 1.01577097065897 | 29.1148693511425 | 0.0351353820218004 |
| 180 | *PLEKHA1* | 3.27427368421053 | 1.00139101268176 | 28.5013689075287 | 0.0351353820218004 |
| 116 | *KIF19* | 3.06534736842105 | 0.990919882222716 | 32.3369605807935 | 0.000315119780720919 |
| 197 | *RAMP1* | 3.37802105263158 | 0.929028992226184 | 13.8888407428743 | 0.0941254875824312 |
| 139 | *MKRN3* | 3.17204210526316 | 0.912346781144781 | 31.4527202075702 | 0.0201423318626269 |
| 251 | *TRIO* | 3.34341052631579 | 0.870830704405163 | 30.8469811293969 | 0.211337646582481 |
| 215 | *SF3A3* | 3.44867368421053 | 0.832741802987136 | 17.9867134024902 | 0.211337646582481 |
| 63 | *DLL3* | 3.42012631578947 | 0.773149560649227 | 9.94496476856477 | 0.0587629002332443 |
| 190 | *PTDSS1* | 3.50425263157895 | 0.726840835460835 | 19.7598923880213 | 0.723433475193249 |
| 97 | *GRAMD1A* | 3.4336 | 0.688121064714606 | 15.2527002345608 | 0.505314187710712 |

| | | | | | |
|---|---|---|---|---|---|
| 128 | *MANEAL* | 3.4456421052 6316 | 0.680032962 962963 | 22.15067600732 6 | 0.3955196 4100646 |
| 247 | *TMSB10* | 3.2667789473 6842 | 0.673641933 458517 | 18.56655759714 14 | 0.0110840 307247928 |
| 262 | *WDR82* | 3.5253052631 5789 | 0.650793773 828024 | 14.42707091633 86 | 0.6175528 45457526 |
| 120 | *LIMD2* | 3.2171789473 6842 | 0.614396789 212331 | 10.82956267474 36 | 0.0058615 810380402 1 |
| 249 | *TRADD* | 3.2788210526 3158 | 0.607059698 492865 | 9.504032319314 67 | 0.0110840 307247928 |
| 121 | *LPIN1* | 2.8805052631 5789 | 0.604296508 388217 | 14.95780584641 54 | 7.1189053 9062112e-09 |
| 70 | *EHD3* | 3.4877473684 2105 | 0.528007359 848485 | 9.522401076555 02 | 0.3955196 4100646 |
| 272 | *ZNF599* | 3.4862315789 4737 | 0.525966612 012987 | 15.92731668103 14 | 0.5053141 87710712 |
| 238 | *TARSL2* | 3.1886315789 4737 | 0.517948526 936027 | 12.21497297058 81 | 0.0014616 384094140 5 |
| 119 | *LEMD2* | 3.4877473684 2105 | 0.466935787 237454 | 8.215103052503 05 | 0.3955196 4100646 |
| 156 | *NFS1* | 3.5944421052 6316 | 0.458733939 393939 | 12.68248887764 28 | 0.9377177 25859144 |
| 10 | *AHCYL2* | 3.4005894736 8421 | 0.458685412 004662 | 11.05516502517 95 | 0.0351353 820218004 |
| 248 | *TNFAIP8 L3* | 3.4666947368 4211 | 0.449119972 643098 | 14.67464156497 1 | 0.3955196 4100646 |
| 66 | *DTNA* | 3.5493894736 8421 | 0.444379287 749288 | 7.179682921172 39 | 0.3955196 4100646 |
| 230 | *STARD9* | 2.7031578947 3684 | 0.435433193 800027 | 8.868697354279 71 | 4.0038062 3533397e-13 |
| 174 | *PHF2* | 3.4802526315 7895 | 0.435398430 523181 | 10.74196204481 79 | 0.2113376 46582481 |
| 14 | *ANKS6* | 3.1991578947 3684 | 0.432931949 245199 | 6.265798143849 61 | 0.0014616 384094140 5 |
| 160 | *NPHP4* | 3.1706105263 1579 | 0.424270785 834536 | 6.074192089197 35 | 0.0003151 197807209 19 |
| 126 | *LYNX1* | 3.3419789473 6842 | 0.386136432 561266 | 9.435533129005 5 | 0.0110840 307247928 |
| 209 | *RTF1* | 3.4396631578 9474 | 0.381013043 430335 | 14.55145873491 83 | 0.1442514 94432929 |
| 201 | *RIIAD1* | 3.5584 | 0.371117055 555556 | 4.451836668886 67 | 0.6175528 45457526 |
| 136 | *MEST* | 3.5944421052 6316 | 0.370895584 693085 | 11.76010394833 92 | 0.9377177 25859144 |

| | | | | | |
|---|---|---|---|---|---|
| 105 | *HLF* | 3.52682105263158 | 0.36950351377018 | 16.1013131949578 | 0.617552845457526 |
| 130 | *MAP4K2* | 3.49827368421053 | 0.366615271302771 | 9.98782297979798 | 0.39551964100646 |
| 100 | *GTF2F1* | 3.43966315789474 | 0.364729621212121 | 5.734255001 | 0.144251494432929 |
| 87 | *FNBP4* | 3.25178947368421 | 0.36253031559644 | 8.44050750972454 | 0.00146163840941405 |
| 17 | *AQP6* | 3.43966315789474 | 0.360486702926703 | 8.82630253684224 | 0.144251494432929 |
| 134 | *MCHR1* | 3.63806315789474 | 0.357249683911767 | 10.4446797506587 | 0.886881035593066 |
| 131 | *MAP6* | 3.43814736842105 | 0.348436442760943 | 8.38790354090354 | 0.211337646582481 |
| 2 | *ACACA* | 3.0864 | 0.314217732919858 | 8.67940834312746 | 0.000315119780720919 |
| 192 | *PTPRS* | 3.56589473684211 | 0.308092298881674 | 5.86041025641026 | 0.814966224158659 |
| 127 | *MALL* | 3.60648421052632 | 0.296009894586895 | 6.71161425567896 | 0.886881035593066 |
| 169 | *PCGF2* | 3.59898947368421 | 0.293549701113701 | 4.52859484126984 | 0.617552845457526 |
| 65 | *DSTYK* | 3.42913684210526 | 0.289901026936027 | 5.55063612849139 | 0.144251494432929 |
| 43 | *CDH4* | 3.34652631578947 | 0.25115400964776 | 3.26276720893486 | 0.000315119780720919 |
| 261 | *VTA1* | 3.69818947368421 | 0.241149277777778 | 2.77630996944232 | 0.969652090183331 |
| 235 | *SYDE2* | 3.52985263157895 | 0.240590071428571 | 2.86085514485514 | 0.39551964100646 |
| 185 | *PQLC1* | 3.59747368421053 | 0.228189033929034 | 11.6204066188713 | 0.814966224158659 |
| 246 | *TMEM38A* | 3.52985263157895 | 0.219509235542236 | 3.04534606782107 | 0.39551964100646 |
| 3 | *ACTR3* | 3.67115789473684 | 0.216613709401709 | 5.87819434682964 | 0.814966224158659 |
| 117 | *KRT17* | 3.54341052631579 | 0.210411202020202 | 1.63098283938284 | 0.211337646582481 |
| 5 | *ADAMTS10* | 3.54189473684211 | 0.207310676767677 | 4.33990675932255 | 0.211337646582481 |
| 68 | *DYNLRB2* | 3.44867368421053 | 0.204359418328585 | 6.108875 | 0.211337646582481 |
| 204 | *RNF208* | 3.67713684210526 | 0.200723694207028 | 6.73318738091973 | 0.969652090183331 |
| 28 | *BCAN* | 3.50130526315789 | 0.186695987567988 | 2.13047860750361 | 0.211337646582481 |

| | | | | | |
|---|---|---|---|---|---|
| 92 | *GFOD1* | 3.58989473684 21 | 0.186647518037518 | 3.39486190476191 | 0.723433475193249 |
| 191 | *PTPRM* | 3.43814736842105 | 0.180281710858586 | 7.39369665276244 | 0.211337646582481 |
| 268 | *ZER1* | 3.66063157894737 | 0.176425096181596 | 3.81744482797904 | 0.814966224158659 |
| 229 | *SSSCA1* | 3.71621052631579 | 0.175574170465337 | 3.52008871800714 | 0.995462402129288 |
| 46 | *CETN3* | 3.71772631578947 | 0.159080015194682 | 6.94961238336827 | 0.969652090183331 |
| 167 | *PCBD1* | 3.44117894736842 | 0.148694430415264 | 5.88373197194962 | 0.0941254875824312 |
| 31 | *C16orf70* | 3.69667368421053 | 0.145054820512821 | 4.36743687423687 | 0.969652090183331 |
| 48 | *CIDECP* | 3.62753684210526 | 0.144239804023137 | 1.36939963924964 | 0.886881035593066 |
| 206 | *RPL13A* | 3.55991578947368 | 0.142349090779591 | 5.28272334066003 | 0.505314187710712 |
| 163 | *NUTF2* | 3.42012631578947 | 0.139092864265364 | 2.97000401820402 | 0.0941254875824312 |
| 146 | *MRPS33* | 3.59149473684211 | 0.13749117003367 | 2.03934920634921 | 0.505314187710712 |
| 55 | *CYFIP2* | 3.66063157894737 | 0.134681541375291 | 1.73640453908836 | 0.814966224158659 |
| 172 | *PDK2* | 3.68614736842105 | 0.131027925084175 | 7.16123300963742 | 0.987199862094804 |
| 257 | *UBE2B* | 3.39907368421053 | 0.125580373479099 | 3.65095245310245 | 0.0941254875824312 |
| 148 | *MYH3* | 3.40656842105263 | 0.122526555250305 | 11.7397462190458 | 0.211337646582481 |
| 231 | *STAT2* | 3.66812631578947 | 0.119143475869809 | 2.53720384615385 | 0.937717725859144 |
| 38 | *CAPRIN2* | 3.36606315789474 | 0.115091592509342 | 3.1870717948718 | 0.00146163840941405 |
| 104 | *HEMK1* | 3.56294736842105 | 0.114958664502164 | 1.15097974247974 | 0.295816504844879 |
| 59 | *DDX51* | 3.37355789473684 | 0.114227422077922 | 4.88000801192272 | 0.00586158103804021 |
| 224 | *SPIRE1* | 3.608 | 0.113594941822067 | 2.50250523594053 | 0.814966224158659 |
| 147 | *MT1A* | 3.73726315789474 | 0.110541777777778 | 1.13762936507936 | 0.995462402129288 |
| 50 | *COG1* | 3.65010526315789 | 0.109725195847363 | 1.05330518207283 | 0.723433475193249 |
| 269 | *ZKSCAN1* | 3.09094736842105 | 0.109273642316017 | 7.51200812608444 | 9.78264217265471e-06 |

| | | | | | |
|---|---|---|---|---|---|
| 108 | *IDS* | 3.7072 | 0.109208080808081 | 1.06653651348651 | 0.987199862094804 |
| 69 | *EEF1A2* | 3.59301052631579 | 0.101929836829837 | 1.81960184176848 | 0.39551964100646 |
| 263 | *WNK2* | 3.52682105263158 | 0.100708518842269 | 3.22563671557201 | 0.617552845457526 |
| 94 | *GNG5* | 3.66063157894737 | 0.0992907121212121 | 1.5324025974026 | 0.814966224158659 |
| 77 | *FABP3* | 3.46821052631579 | 0.0929783389450056 | 1.90514975302475 | 0.295816504844879 |
| 49 | *CNOT1* | 3.62905263157895 | 0.0912949740999741 | 3.06452463910359 | 0.814966224158659 |
| 239 | *TBC1D1* | 3.49684210526316 | 0.0856071771238021 | 1.60961306144835 | 0.0351353820218004 |
| 35 | *CABYR* | 3.74778947368421 | 0.0849790909090909 | 0.893970699475962 | 0.995462402129288 |
| 61 | *DEPDC5* | 3.63957894736842 | 0.0840568650793651 | 1.53032575757576 | 0.814966224158659 |
| 12 | *ANGPT2* | 3.67267368421053 | 0.0820531992313242 | 1.22021611721612 | 0.723433475193249 |
| 53 | *CREB3L3* | 3.37052631578947 | 0.0804639197530864 | 3.38723785658786 | 0.0351353820218004 |
| 103 | *HCP5* | 3.56741052631579 | 0.0792544940322024 | 6.81016785714286 | 0.723433475193249 |
| 205 | *RNFT2* | 3.75528421052632 | 0.0747801226551227 | 3.52312417582418 | 0.999720378726235 |
| 225 | *SPTAN1* | 3.66964210526316 | 0.0735853598484848 | 1.65906971916972 | 0.886881035593066 |
| 99 | *GRK6* | 3.8064 | 0.0699987373737374 | 0.309419047619048 | 0.99995956392929 |
| 223 | *SP100* | 3.35705263157895 | 0.0676647598096349 | 2.15940566884096 | 0.00146163840941405 |
| 7 | *ADCY5* | 3.68917894736842 | 0.0648512121212121 | 1.04309492729493 | 0.937717725859144 |
| 18 | *ARF3* | 3.61103157894737 | 0.0647663247863248 | 0.538215584415584 | 0.617552845457526 |
| 182 | *POLR2E* | 3.75983157894737 | 0.0644057614607614 | 0.359917948717949 | 0.987199862094804 |
| 245 | *TMEM199* | 3.77936842105263 | 0.0620176481481482 | 0.298661255411255 | 0.995462402129288 |
| 253 | *TRUB1* | 3.68766315789474 | 0.056780303030303 | 1.32884126984127 | 0.969652090183331 |
| 237 | *TADA2B* | 3.62753684210526 | 0.056768127946128 | 1.815963816 | 0.886881035593066 |
| 207 | *RPL24* | 3.50130526315789 | 0.0553058580246914 | 3.48437127366098 | 0.211337646582481 |
| 29 | *BOP1* | 3.55393684210526 | 0.0536823106060607 | 1.10354567099567 | 0.211337646582481 |

| | | | | | |
|---|---|---|---|---|---|
| 258 | *UBE2E2* | 3.33145263157895 | 0.0499885801066218 | 2.94739859846036 | 0.0110840307247928 |
| 162 | *NRSN2* | 3.66812631578947 | 0.0498848849607183 | 0.79810722055722 | 0.937717725859144 |
| 33 | *C9orf116* | 3.40362105263158 | 0.04890544332211 | 2.46295422077922 | 0.0201423318626269 |
| 137 | *METRN* | 3.63056842105263 | 0.0457126843434343 | 1.51455737595738 | 0.723433475193249 |
| 20 | *ARL2* | 3.57793684210526 | 0.0450555499438833 | 3.90765427350427 | 0.723433475193249 |
| 243 | *THY1* | 3.57195789473684 | 0.0448987474747475 | 7.39941416083916 | 0.39551964100646 |
| 76 | *EVL* | 3.73877894736842 | 0.0440790895061728 | 0.449466666666667 | 0.987199862094804 |
| 8 | *AFTPH* | 3.74029473684211 | 0.0421391913580247 | 0.422659951159951 | 0.969652090183331 |
| 254 | *TTC3P1* | 3.62155789473684 | 0.0416395575412242 | 1.06458589466089 | 0.39551964100646 |
| 252 | *TROVE2* | 3.63208421052632 | 0.0399988531144781 | 1.6103670995671 | 0.617552845457526 |
| 81 | *FARSB* | 3.65010526315789 | 0.0386503308481642 | 1.04546505063564 | 0.723433475193249 |
| 19 | *ARIH2* | 3.65162105263158 | 0.037836835016835 | 0.634300505050505 | 0.723433475193249 |
| 189 | *PSME3* | 3.64109473684211 | 0.0363810222863556 | 0.890712842712842 | 0.723433475193249 |
| 138 | *MICU1* | 3.32993684210526 | 0.0363369157601658 | 3.32980209365798 | 0.0201423318626269 |
| 193 | *PTPRU* | 3.72976842105263 | 0.035850218013468 | 0.665815728715729 | 0.969652090183331 |
| 175 | *PIGF* | 3.63208421052632 | 0.0348274883449883 | 0.542671001221001 | 0.617552845457526 |
| 78 | *FAHD1* | 3.74778947368421 | 0.0337167500400834 | 0.89400233426704 | 0.995462402129288 |
| 151 | *NDUFA9* | 3.40808421052632 | 0.0332831440781441 | 5.74433088023088 | 0.144251494432929 |
| 152 | *NDUFS4* | 3.68168421052632 | 0.0323721212121212 | 0.593121428571429 | 0.814966224158659 |
| 102 | *HBA2* | 3.72976842105263 | 0.0318584781144781 | 0.326134920634921 | 0.969652090183331 |
| 267 | *ZEB2* | 3.584 | 0.0310993703703704 | 1.7225862026862 | 0.295816504844879 |
| 90 | *FZD5* | 3.67115789473684 | 0.0297755694444445 | 0.868276190476191 | 0.814966224158659 |
| 159 | *NOS1AP* | 3.50282105263158 | 0.0290901334776335 | 4.76725384615385 | 0.0941254875824312 |
| 154 | *NEURL3* | 3.52538947368421 | 0.0288021813140564 | 0.82337215007215 | 0.0941254875824312 |
| 216 | *SLC10A4* | 3.65162105263158 | 0.0284148653198653 | 3.38313183483183 | 0.723433475193249 |

| | | | | | |
|---|---|---|---|---|---|
| 158 | *NOS1* | 3.61254736842105 | 0.0251225992618493 | 1.12371601731602 | 0.39551964100646 |
| 4 | *ACTR6* | 3.68917894736842 | 0.0246284788051455 | 0.736511038961039 | 0.937717725859144 |
| 244 | *TMEM191A* | 3.69970526315789 | 0.0243068537511871 | 0.477317094017094 | 0.937717725859144 |
| 71 | *EIF1B* | 3.7898947368421 | 0.0240237477337477 | 0.175630303030303 | 0.995462402129288 |
| 178 | *PLBD2* | 3.74181052631579 | 0.0234021842355176 | 0.445480952380953 | 0.937717725859144 |
| 266 | *YWHAZ* | 3.64858947368421 | 0.0234001262626262 | 0.850307142857143 | 0.886881035593066 |
| 140 | *MLLT11* | 3.7824 | 0.0230618181818182 | 0.137028571428572 | 0.969652090183331 |
| 109 | *ILF2* | 3.54037894736842 | 0.0214477769499019 | 1.73145313580468 | 0.39551964100646 |
| 30 | *C11orf1* | 3.71772631578947 | 0.0203738528138528 | 0.711233333333334 | 0.987199862094804 |
| 133 | *MAX* | 3.68766315789474 | 0.0203404463437797 | 1.90866024531025 | 0.969652090183331 |
| 86 | *FMNL1* | 3.66964210526316 | 0.0198650540123457 | 0.763097768897769 | 0.886881035593066 |
| 44 | *CDK5R2* | 3.71023157894737 | 0.0198379938271605 | 0.489675885225885 | 0.937717725859144 |
| 112 | *JAZF1* | 3.68766315789474 | 0.0196862676767677 | 1.3744221001221 | 0.969652090183331 |
| 256 | *TTPAL* | 3.49381052631579 | 0.0194280763989098 | 1.21472508047508 | 0.0941254875824312 |
| 79 | *FAM160B1* | 3.73128421052632 | 0.0191640296431963 | 0.268657142857143 | 0.937717725859144 |
| 202 | *RILPL1* | 3.60050526315789 | 0.0189595183982684 | 1.31331551226551 | 0.617552845457526 |
| 98 | *GRIN2B* | 3.75831578947368 | 0.0183636790986791 | 0.384233333333333 | 0.995462402129288 |
| 186 | *PRB1* | 3.80791578947368 | 0.0179204545454545 | 0.153826556776557 | 0.999720378726235 |
| 47 | *CFD* | 3.66812631578947 | 0.0173720590520591 | 0.997217965367965 | 0.937717725859144 |
| 255 | *TTC9B* | 3.64109473684211 | 0.0170614848484848 | 0.782776984126984 | 0.723433475193249 |
| 67 | *DYNLRB1* | 3.62905263157895 | 0.016930707070707 | 0.920671123321124 | 0.814966224158659 |
| 56 | *DARS* | 3.65465263157895 | 0.0166882626262626 | 0.925286363636364 | 0.505314187710712 |
| 233 | *STXBP1* | 3.67865263157895 | 0.0160915364357864 | 0.861870779220779 | 0.937717725859144 |
| 91 | *G3BP1* | 3.67115789473684 | 0.0156708154191487 | 1.17384603174603 | 0.814966224158659 |
| 183 | *PPDPF* | 3.71924210526316 | 0.0147951388888889 | 0.636511760461761 | 0.969652090183331 |

| | | | | | |
|---|---|---|---|---|---|
| 149 | *NAE1* | 3.64261052631579 | 0.0143606275144608 | 0.874212698412698 | 0.617552845457526 |
| 132 | *MARK4* | 3.75528421052632 | 0.0142098535353536 | 5.75243640473733 | 0.999720378726235 |
| 111 | *IRF2BP1* | 3.81844210526316 | 0.0136481481481481 | 0.0456666666666669 | 0.999720378726235 |
| 96 | *GPNMB* | 3.69970526315789 | 0.01299980995856 | 1.82519043062201 | 0.937717725859144 |
| 106 | *HMG20A* | 3.69221052631579 | 0.0124871693121693 | 0.371685714285714 | 0.814966224158659 |
| 181 | *POLR1A* | 3.66964210526316 | 0.0124432954545455 | 0.893825396825397 | 0.886881035593066 |
| 188 | *PSMD7* | 3.71023157894737 | 0.0121722222222223 | 0.341664285714286 | 0.937717725859144 |
| 22 | *ARPC2* | 3.66964210526316 | 0.0121008984071484 | 0.821129797979798 | 0.886881035593066 |
| 110 | *IPO5* | 3.73128421052632 | 0.0118374962691629 | 0.275711111111111 | 0.937717725859144 |
| 236 | *SYT3* | 3.75983157894737 | 0.0114285714285714 | 0.233256349206349 | 0.987199862094804 |
| 58 | *DCUN1D5* | 3.73877894736842 | 0.0113131313131313 | 0.864662937062937 | 0.987199862094804 |
| 271 | *ZNF415* | 3.81692631578947 | 0.0111111111111111 | 0.178 | 0.99995956392929 |
| 101 | *HBA1* | 3.72075789473684 | 0.0108449074074074 | 0.259042857142857 | 0.937717725859144 |
| 218 | *SLC38A2* | 3.66214736842105 | 0.00998614102564105 | 0.45666235986236 | 0.723433475193249 |
| 221 | *SORBS3* | 3.74930526315789 | 0.00931887442804106 | 0.182423809523809 | 0.987199862094804 |
| 11 | *AMIGO1* | 3.66063157894737 | 0.00831927272727272 | 0.828664141414141 | 0.814966224158659 |
| 194 | *PWWP2B* | 3.7989052631579 | 0.0082962962962963 | 0.157166666666667 | 0.998703548494128 |
| 72 | *ELOVL6* | 3.76884210526316 | 0.00822651515151514 | 0.2261 | 0.995462402129288 |
| 89 | *FRMD8* | 3.75983157894737 | 0.00781163636363637 | 0.435960173160173 | 0.987199862094804 |
| 23 | *ASTN2* | 3.82745263157895 | 0.00711333333333334 | 0.0527142857142859 | 0.99995956392929 |
| 222 | *SOS1* | 3.7328 | 0.00708229166666665 | 0.462347402597403 | 0.886881035593066 |
| 54 | *CSTB* | 3.66366315789474 | 0.00691818588818588 | 0.536272222222222 | 0.617552845457526 |
| 36 | *CACNB1* | 3.71174736842105 | 0.00606888111888113 | 0.699252380952381 | 0.886881035593066 |
| 199 | *RESP18* | 3.71023157894737 | 0.00575428869895542 | 0.222675396825396 | 0.937717725859144 |
| 135 | *MED30* | 3.71023157894737 | 0.00537226851851851 | 0.550905677655678 | 0.937717725859144 |

| | | | | | |
|---|---|---|---|---|---|
| 226 | *SRPK2* | 3.69221052631579 | 0.0050219670206337 | 0.327274603174603 | 0.814966224158659 |
| 25 | *ATP6V1G1* | 3.56597894736842 | 0.00479510732323233 | 0.656959862359862 | 0.144251494432929 |
| 187 | *PSMB10* | 3.77785263157895 | 0.00311111111111111 | 0.489724458874459 | 0.998703548494128 |
| 170 | *PCK2* | 3.47275789473684 | 0.00293965263748598 | 1.64867130647131 | 0.0587629002332443 |
| 95 | *GNPDA2* | 3.72075789473684 | 0.00218606177156177 | 0.312225396825397 | 0.937717725859144 |
| 150 | *NDFIP1* | 3.58694736842105 | 0.00167184848484851 | 2.37340525259055 | 0.814966224158659 |
| 16 | *APBA1* | 3.72227368421053 | 0.00157195809591643 | 0.180592063492064 | 0.886881035593066 |
| 9 | *AGPAT5* | 3.72075789473684 | 0.00153782517482517 | 0.223280952380952 | 0.937717725859144 |
| 195 | *RAB11FIP2* | 3.71924210526316 | 0.00076218795093796 | 0.886310989010989 | 0.969652090183331 |
| 214 | *SERTAD3* | 3.68766315789474 | 0.000730959595959597 | 0.866100072150072 | 0.969652090183331 |
| 213 | *SEC24D* | 3.69221052631579 | 0.000283232323232341 | 0.378695238095238 | 0.814966224158659 |
| 26 | *ATP9B* | 3.54938947368421 | 0.359855833 | 6.72415184815185 | 0.505314187710712 |
| 64 | *DNAJC30* | 3.63208421052632 | 0.1027205 | 0.836881854256854 | 0.617552845457526 |
| 227 | *SRRT* | 3.67865263157895 | 0.057976 | 1.3259913421374 | 0.937717725859144 |
| 161 | *NPPB* | 3.83646315789474 | 0.0147 | 0.049 | 0.999997061551901 |
| 260 | *VAC14* | 3.7989052631579 | 0.01085 | 0.111566666666667 | 0.998703548494128 |
| 113 | *JOSD1* | 3.78837894736842 | 0.0104395 | 0.748197113997114 | 0.998703548494128 |
| 15 | *AP2S1* | 3.69970526315789 | 0.0053125 | 0.351918253968254 | 0.937717725859144 |
| 37 | *CAND2* | 3.83646315789474 | 0 | 0.0201666666666667 | 0.999997061551901 |
| 107 | *IDI1* | 3.80791578947368 | 0 | 0.0860952380952381 | 0.999720378726235 |
| 142 | *MMGT1* | 3.80791578947368 | 0 | 0.0942000000000001 | 0.999720378726235 |
| 176 | *PIGX* | 3.78837894736842 | 0 | 0.160211904761905 | 0.998703548494128 |
| 184 | *PPP5C* | 3.84698947368421 | 0 | 0.00240000000000001 | 0.999997061551901 |
| 264 | *YIF1B* | 3.82745263157895 | 0 | 0.0342619047619048 | 0.99995956392929 |
| 13 | *ANKIB1* | 3.81844210526316 | -0.000507716 | 0.0852619047619047 | 0.999720378726235 |

| | | | | | |
|---|---|---|---|---|---|
| 217 | *SLC1A4* | 3.7989052631579 | -0.000769231 | 0.190595238095238 | 0.998703548494128 |
| 114 | *JUND* | 3.4922947368421 | -0.001067311 | 1.35815007215007 | 0.144251494432929 |
| 144 | *MRPL34* | 3.62004210526316 | -0.001225812 | 4.63147087912088 | 0.723433475193249 |
| 80 | *FAM174B* | 3.74930526315789 | -0.002024949 | 0.224796825396826 | 0.987199862094804 |
| 129 | *MAP2* | 3.69970526315789 | -0.002432945 | 0.323827777777777 | 0.886881035593066 |
| 75 | *ETV5* | 3.71621052631579 | -0.002562146 | 3.62634978354978 | 0.995462402129288 |
| 200 | *RFXANK* | 3.64261052631579 | -0.002640102 | 1.02392761682762 | 0.617552845457526 |
| 60 | *DEF8* | 3.7989052631579 | -0.00306 | 0.186107142857143 | 0.998703548494128 |
| 196 | *RAB12* | 3.78837894736842 | -0.003083333 | 0.225468253968254 | 0.998703548494128 |
| 85 | *FLOT2* | 3.82745263157895 | -0.003166667 | 0.128035714285714 | 0.99995956392929 |
| 84 | *FHOD1* | 3.68016842105263 | -0.003241538 | 6.22477142857143 | 0.886881035593066 |
| 125 | *LRRN2* | 3.74778947368421 | -0.003618547 | 0.365284126984127 | 0.995462402129288 |
| 93 | *GLT8D1* | 3.71924210526316 | -0.004321731 | 0.503572035480859 | 0.969652090183331 |
| 198 | *RASSF7* | 3.75082105263158 | -0.004506845 | 0.426551282051282 | 0.969652090183331 |
| 141 | *MLYCD* | 3.80943157894737 | -0.004625 | 0.0653999999999999 | 0.998703548494128 |
| 240 | *TBC1D2B* | 3.67418947368421 | -0.005238468 | 0.770654401154401 | 0.617552845457526 |
| 88 | *FOXP4* | 3.32993684210526 | -0.005942757 | 4.72668464035964 | 0.0110840307247928 |
| 166 | *PARD3B* | 3.7989052631579 | -0.006068636 | 0.120669841269841 | 0.998703548494128 |
| 242 | *THOP1* | 3.7989052631579 | -0.006071111 | 0.0646666666666667 | 0.998703548494128 |
| 153 | *NEK9* | 3.74778947368421 | -0.006099327 | 0.534551282051282 | 0.995462402129288 |
| 208 | *RPL7L1* | 3.80791578947368 | -0.006755556 | 0.0676952380952379 | 0.999720378726235 |
| 219 | *SLC38A3* | 3.74029473684211 | -0.007338 | 2.71819050165521 | 0.969652090183331 |
| 34 | *C9orf16* | 3.77936842105263 | -0.007470444 | 0.156946825396825 | 0.995462402129288 |
| 45 | *CEP19* | 3.70122105263158 | -0.007934844 | 0.568258823529411 | 0.886881035593066 |
| 39 | *CCND2* | 3.62155789473684 | -0.007956777 | 0.736383838383839 | 0.617552845457526 |

| | | | | | |
|---|---|---|---|---|---|
| 203 | *RNF123* | 3.72976842105263 | -0.008905808 | 0.432719047619048 | 0.937717725859144 |
| 164 | *OAZ2* | 3.69818947368421 | -0.009023902 | 3.46913412698413 | 0.969652090183331 |
| 234 | *STYXL1* | 3.61103157894737 | -0.009451835 | 0.94018253968254 | 0.617552845457526 |
| 115 | *KCNQ2* | 3.68917894736842 | -0.00952919 | 0.727691847041847 | 0.937717725859144 |
| 168 | *PCDHAC2* | 3.65162105263158 | -0.01013571 | 0.711888888888888 | 0.617552845457526 |
| 40 | *CCNYL1* | 3.73128421052632 | -0.010266667 | 0.218352380952381 | 0.886881035593066 |
| 122 | *LRP1* | 3.66366315789474 | -0.012749262 | 0.844791774891775 | 0.505314187710712 |
| 259 | *UBE2L3* | 3.72075789473684 | -0.015720313 | 0.307988095238095 | 0.937717725859144 |
| 123 | *LRP3* | 3.65162105263158 | -0.016783763 | 1.41559065233013 | 0.723433475193249 |
| 6 | *ADCY1* | 3.67865263157895 | -0.017337697 | 0.757438528138528 | 0.937717725859144 |
| 212 | *SCRN1* | 3.58997894736842 | -0.017427481 | 1.47393095238095 | 0.617552845457526 |
| 51 | *COPG2* | 3.64261052631579 | -0.017461941 | 1.05930230880231 | 0.617552845457526 |
| 42 | *CD34* | 3.72075789473684 | -0.018082315 | 0.258433261183261 | 0.937717725859144 |
| 145 | *MRPL49* | 3.65313684210526 | -0.019020088 | 0.714385714285714 | 0.617552845457526 |
| 41 | *CCT6B* | 3.73128421052632 | -0.021094809 | 0.175246031746032 | 0.886881035593066 |
| 265 | *YWHAB* | 3.63056842105263 | -0.023004506 | 0.763020562770562 | 0.723433475193249 |
| 83 | *FCGBP* | 3.62004210526316 | -0.027160687 | 0.763925108225108 | 0.723433475193249 |
| 211 | *SAMD15* | 3.73877894736842 | -0.028179293 | 0.44965 | 0.987199862094804 |
| 250 | *TRIM4* | 3.68168421052632 | -0.028922127 | 0.985802530802531 | 0.814966224158659 |
| 232 | *STX4* | 3.66366315789474 | -0.030836797 | 0.372011544011544 | 0.617552845457526 |
| 177 | *PITPNM2* | 3.69970526315789 | -0.031937611 | 0.527939682539682 | 0.937717725859144 |
| 118 | *KRTCAP2* | 3.66063157894737 | -0.033912603 | 0.542353174603174 | 0.814966224158659 |
| 62 | *DLG4* | 3.54189473684211 | -0.034367336 | 1.16781092796093 | 0.295816504844879 |
| 82 | *FBXL18* | 3.64109473684211 | -0.035089462 | 0.898162324929972 | 0.723433475193249 |
| 179 | *PLD3* | 3.70871578947368 | -0.045132552 | 0.436275396825397 | 0.969652090183331 |

| | | | | | |
|---|---|---|---|---|---|
| 241 | *TCF7* | 3.63208421052632 | -0.048217738 | 0.741714285714286 | 0.39551964100646 |
| 52 | *CPSF4* | 3.65313684210526 | -0.049535175 | 0.406093650793651 | 0.617552845457526 |
| 73 | *EME1* | 3.68168421052632 | -0.05806076 | 0.656982112332112 | 0.814966224158659 |
| 173 | *PEX11A* | 3.76732631578947 | -0.074052727 | 0.484206349206349 | 0.998703548494128 |
| 32 | *C7* | 3.71772631578947 | -0.080208106 | 0.78189292929293 | 0.987199862094804 |
| 270 | *ZNF233* | 3.62602105263158 | -0.104836478 | 7.22369907265937 | 0.937717725859144 |

**Table S32. Importance (Dominance) of each gene in the random forest model for superior temporal cortex (STC).** Mean_min_depth represents the average minimum depth at which a variable is used as a split node across all trees, with lower values indicating higher importance as the variable is used earlier in the decision process. IncrMSE, % (Increase in Mean Squared Error) quantifies the relative increase in prediction error when the variable is permuted, reflecting its contribution to the model's predictive accuracy; higher values signify greater importance. Node_purity_increase measures the improvement in node purity (e.g., Gini index or variance reduction) achieved by splitting on the variable, where larger values indicate a stronger impact on classification or regression performance. Finally, the P value assesses the statistical significance of a variable's importance, typically derived from permutation tests, with lower values suggesting that the variable's contribution is unlikely to be due to random chance. This presentation was in descending order by the IncrMSE.

| Entrez ID | Gene Symbol | Mean_min_depth | IncrMSE, % | node_purity_increase | P value |
|---|---|---|---|---|---|
| 124 | *LRRC61* | 0.333333333 | 6.352490946 | 95.85451459 | 2.57E-23 |
| 27 | *BASP1* | 1.175578947 | 3.199210407 | 64.56379957 | 1.61E-14 |
| 272 | *ZNF599* | 1.253894737 | 3.19276767 | 67.49819392 | 1.52E-11 |
| 273 | *ZNF821* | 1.407719298 | 3.04860079 | 55.95521165 | 6.63E-14 |
| 157 | *NNAT* | 1.785684211 | 2.615081407 | 48.11208414 | 7.76E-08 |
| 63 | *DLL3* | 2.268912281 | 2.403365007 | 36.04153849 | 0.000524348 |
| 165 | *PAFAH1B3* | 1.429333333 | 2.402057979 | 52.72240389 | 4.05E-12 |
| 210 | *S100A1* | 1.689824561 | 2.291404511 | 47.34520233 | 1.99E-10 |
| 121 | *LPIN1* | 0.923087719 | 2.051665826 | 32.78699528 | 1.37E-22 |
| 120 | *LIMD2* | 2.58877193 | 1.32996261 | 21.60104658 | 0.001170712 |
| 247 | *TMSB10* | 2.586105263 | 1.184530763 | 31.37789327 | 0.019311298 |
| 126 | *LYNX1* | 2.033964912 | 1.06939521 | 23.32151444 | 6.92E-10 |

| | | | | | |
|---|---|---|---|---|---|
| 174 | *PHF2* | 2.553684211 | 1.059198545 | 23.56229685 | 0.0025123 34 |
| 97 | *GRAMD1 A* | 2.592842105 | 1.046994906 | 20.08960026 | 0.0102181 36 |
| 109 | *ILF2* | 2.673824561 | 1.033672873 | 18.83533881 | 0.0193112 98 |
| 143 | *MPZL1* | 1.898947368 | 0.975124648 | 20.09806784 | 1.05E-12 |
| 17 | *AQP6* | 2.062315789 | 0.972183054 | 15.14488639 | 7.75E-09 |
| 215 | *SF3A3* | 2.560421053 | 0.842114511 | 13.33818712 | 0.0005243 48 |
| 171 | *PDE9A* | 2.658947368 | 0.836604177 | 21.41730472 | 0.0011707 12 |
| 21 | *ARMCX6* | 2.746666667 | 0.827380746 | 18.41079354 | 0.0025123 34 |
| 24 | *ATF5* | 2.951859649 | 0.822332662 | 10.46500126 | 0.0987984 53 |
| 209 | *RTF1* | 2.938385965 | 0.792361137 | 11.24742561 | 0.1544972 13 |
| 103 | *HCP5* | 2.687298246 | 0.688100074 | 14.25973048 | 0.0051751 09 |
| 105 | *HLF* | 2.870877193 | 0.62329312 | 12.27161349 | 0.0601344 |
| 134 | *MCHR1* | 2.83845614 | 0.605467837 | 6.533663067 | 0.0102181 36 |
| 92 | *GFOD1* | 1.944842105 | 0.551773083 | 9.987827716 | 4.05E-12 |
| 248 | *TNFAIP8 L3* | 2.835789474 | 0.549763358 | 12.29965402 | 0.0601344 |
| 190 | *PTDSS1* | 2.772350877 | 0.515527595 | 15.80048193 | 0.0348823 59 |
| 155 | *NEUROD 2* | 3.043649123 | 0.456198184 | 8.958995765 | 0.1544972 13 |
| 100 | *GTF2F1* | 2.892491228 | 0.441518316 | 12.0618477 | 0.1544972 13 |
| 70 | *EHD3* | 2.888421053 | 0.41557225 | 9.929130775 | 0.0601344 |
| 66 | *DTNA* | 2.866807018 | 0.38758563 | 5.007262477 | 0.0102181 36 |
| 237 | *TADA2B* | 3.124631579 | 0.338559071 | 6.005063736 | 0.3244222 51 |
| 268 | *ZER1* | 2.729122807 | 0.322085553 | 3.710461479 | 0.0025123 34 |
| 136 | *MEST* | 3.185403509 | 0.281238866 | 9.598378901 | 0.7790631 74 |
| 177 | *PITPNM 2* | 2.81277193 | 0.271716613 | 2.434183239 | 0.0005243 48 |
| 223 | *SP100* | 3.297403509 | 0.269846559 | 1.506279365 | 0.6728754 24 |
| 229 | *SSSCA1* | 3.347368421 | 0.268810948 | 2.878953333 | 0.9265833 5 |
| 261 | *VTA1* | 3.288 | 0.253294728 | 7.53769012 | 0.9653371 14 |

| | | | | | |
|---|---|---|---|---|---|
| 101 | *HBA1* | 3.393263158 | 0.247491389 | 4.293770476 | 0.965337114 |
| 262 | *WDR82* | 3.340631579 | 0.246883494 | 6.063305788 | 0.965337114 |
| 1 | *AATK* | 3.078736842 | 0.23632253 | 3.96247207 | 0.22969607 |
| 239 | *TBC1D1* | 2.171649123 | 0.230113808 | 9.53447051 | 7.76E-08 |
| 135 | *MED30* | 3.24477193 | 0.221920253 | 3.984233983 | 0.672875424 |
| 206 | *RPL13A* | 3.050385965 | 0.220970379 | 4.531768356 | 0.098798453 |
| 119 | *LEMD2* | 3.308210526 | 0.214151497 | 3.461570537 | 0.672875424 |
| 162 | *NRSN2* | 2.775017544 | 0.201661141 | 3.428696784 | 0.005175109 |
| 140 | *MLLT11* | 3.308210526 | 0.199041471 | 5.065010808 | 0.779063174 |
| 246 | *TMEM38A* | 3.347368421 | 0.198835899 | 5.339720656 | 0.92658335 |
| 139 | *MKRN3* | 3.216421053 | 0.19390653 | 5.287750625 | 0.554702513 |
| 86 | *FMNL1* | 2.980210526 | 0.191898424 | 3.039314413 | 0.154497213 |
| 169 | *PCGF2* | 3.192140351 | 0.190108728 | 7.917825649 | 0.672875424 |
| 91 | *G3BP1* | 2.834385965 | 0.189599575 | 3.471436708 | 0.001170712 |
| 18 | *ARF3* | 2.954526316 | 0.182838805 | 2.431085672 | 0.019311298 |
| 158 | *NOS1* | 3.02877193 | 0.179966162 | 1.820319368 | 0.019311298 |
| 233 | *STXBP1* | 2.965333333 | 0.179923286 | 2.551974695 | 0.034882359 |
| 222 | *SOS1* | 3.308210526 | 0.179517399 | 3.021856623 | 0.779063174 |
| 159 | *NOS1AP* | 2.528 | 0.164543617 | 6.249616787 | 9.39E-05 |
| 102 | *HBA2* | 2.703438596 | 0.156698268 | 5.651270188 | 9.39E-05 |
| 220 | *SNCG* | 3.194807018 | 0.139323662 | 3.895355402 | 0.324422251 |
| 48 | *CIDECP* | 3.142175439 | 0.137347515 | 2.297405937 | 0.22969607 |
| 50 | *COG1* | 3.138105263 | 0.130210431 | 1.805916135 | 0.154497213 |
| 74 | *EPS8L2* | 3.301473684 | 0.125176083 | 2.225316681 | 0.86491113 |
| 72 | *ELOVL6* | 3.360842105 | 0.123297424 | 1.598213652 | 0.779063174 |
| 180 | *PLEKHA1* | 3.354105263 | 0.122738352 | 0.494328571 | 0.86491113 |
| 199 | *RESP18* | 2.958596491 | 0.118921331 | 2.581704484 | 0.0601344 |

| | | | | | |
|---|---|---|---|---|---|
| 224 | *SPIRE1* | 3.312280702 | 0.117211199 | 5.075805455 | 0.92658335 |
| 111 | *IRF2BP1* | 3.031438596 | 0.117178895 | 2.267590418 | 0.010218136 |
| 51 | *COPG2* | 3.157052632 | 0.114706975 | 5.702759141 | 0.435186727 |
| 112 | *JAZF1* | 3.314947368 | 0.113148627 | 0.88622381 | 0.672875424 |
| 53 | *CREB3L3* | 2.993684211 | 0.111608681 | 2.501223676 | 0.0601344 |
| 228 | *SSFA2* | 3.279859649 | 0.107018586 | 3.765699394 | 0.672875424 |
| 62 | *DLG4* | 3.456701754 | 0.101072294 | 0.601483333 | 0.986203542 |
| 187 | *PSMB10* | 3.100350877 | 0.09943298 | 2.010851937 | 0.435186727 |
| 192 | *PTPRS* | 3.395929825 | 0.093247338 | 2.437153413 | 0.779063174 |
| 216 | *SLC10A4* | 3.013894737 | 0.091158584 | 1.710052975 | 0.010218136 |
| 2 | *ACACA* | 2.608982456 | 0.090640896 | 4.069385702 | 0.000226019 |
| 78 | *FAHD1* | 3.177263158 | 0.086932685 | 2.247729957 | 0.324422251 |
| 29 | *BOP1* | 3.144842105 | 0.081964196 | 2.265378995 | 0.0601344 |
| 96 | *GPNMB* | 3.216421053 | 0.081754256 | 4.690999807 | 0.554702513 |
| 117 | *KRT17* | 3.364912281 | 0.079595726 | 2.550055642 | 0.92658335 |
| 245 | *TMEM199* | 3.30554386 | 0.075892047 | 8.130971449 | 0.965337114 |
| 218 | *SLC38A2* | 3.375719298 | 0.075537446 | 0.969746376 | 0.965337114 |
| 251 | *TRIO* | 3.152982456 | 0.07192062 | 6.363498098 | 0.435186727 |
| 254 | *TTC3P1* | 3.513403509 | 0.070853227 | 0.152165873 | 0.998920641 |
| 258 | *UBE2E2* | 3.24477193 | 0.068732875 | 1.494518464 | 0.554702513 |
| 167 | *PCBD1* | 3.017964912 | 0.065854198 | 3.011903544 | 0.034882359 |
| 106 | *HMG20A* | 3.467508772 | 0.064762987 | 0.582633117 | 0.995563154 |
| 133 | *MAX* | 3.170526316 | 0.063923989 | 3.62455162 | 0.435186727 |
| 10 | *AHCYL2* | 3.223157895 | 0.063199051 | 0.905252381 | 0.435186727 |
| 59 | *DDX51* | 3.456701754 | 0.062521556 | 0.600729004 | 0.986203542 |
| 175 | *PIGF* | 3.520140351 | 0.061634556 | 0.17883254 | 0.995563154 |

| | | | | | |
|---|---|---|---|---|---|
| 77 | *FABP3* | 3.329824561 | 0.060508681 | 1.243290772 | 0.92658335 |
| 47 | *CFD* | 3.117894737 | 0.060011262 | 2.045355506 | 0.435186727 |
| 68 | *DYNLRB2* | 3.238035088 | 0.059891047 | 6.332961501 | 0.779063174 |
| 9 | *AGPAT5* | 3.314947368 | 0.056515039 | 2.878188425 | 0.672875424 |
| 219 | *SLC38A3* | 3.428350877 | 0.05637 | 0.697994877 | 0.965337114 |
| 207 | *RPL24* | 2.973473684 | 0.055629828 | 7.159129676 | 0.154497213 |
| 108 | *IDS* | 3.216421053 | 0.055412313 | 3.585809993 | 0.554702513 |
| 26 | *ATP9B* | 3.439157895 | 0.055243056 | 0.95101917 | 0.986203542 |
| 14 | *ANKS6* | 3.343298246 | 0.053206914 | 0.554765007 | 0.779063174 |
| 147 | *MT1A* | 2.934315789 | 0.048613182 | 3.707740481 | 0.098798453 |
| 57 | *DCHS1* | 3.142175439 | 0.047129134 | 1.974988354 | 0.324422251 |
| 19 | *ARIH2* | 3.393263158 | 0.045125736 | 0.946885714 | 0.965337114 |
| 12 | *ANGPT2* | 2.900491228 | 0.044961802 | 2.338847124 | 0.000524348 |
| 44 | *CDK5R2* | 3.389192982 | 0.043478125 | 0.87523956 | 0.86491113 |
| 212 | *SCRN1* | 3.286596491 | 0.041047554 | 1.024850144 | 0.554702513 |
| 69 | *EEF1A2* | 3.319017544 | 0.040295 | 0.864010317 | 0.86491113 |
| 90 | *FZD5* | 3.301473684 | 0.040103166 | 1.671184343 | 0.86491113 |
| 265 | *YWHAB* | 3.456701754 | 0.03894254 | 0.385598413 | 0.986203542 |
| 123 | *LRP3* | 3.38245614 | 0.037899716 | 0.461780159 | 0.92658335 |
| 186 | *PRB1* | 3.491789474 | 0.037793081 | 0.267109524 | 0.986203542 |
| 243 | *THY1* | 3.24477193 | 0.037743136 | 3.318717863 | 0.672875424 |
| 138 | *MICU1* | 3.229894737 | 0.034868126 | 0.720063853 | 0.324422251 |
| 114 | *JUND* | 3.485052632 | 0.033910895 | 0.291391392 | 0.995563154 |
| 238 | *TARSL2* | 3.378385965 | 0.03387324 | 0.813438095 | 0.779063174 |
| 179 | *PLD3* | 3.290666667 | 0.033577868 | 1.44671751 | 0.779063174 |

| | | | | | |
|---|---|---|---|---|---|
| 8 | *AFTPH* | 3.445894737 | 0.03352381 | 0.330021429 | 0.965337114 |
| 22 | *ARPC2* | 3.413473684 | 0.031038673 | 0.238254762 | 0.779063174 |
| 80 | *FAM174B* | 3.354105263 | 0.030129282 | 0.920669625 | 0.86491113 |
| 23 | *ASTN2* | 3.456701754 | 0.02934101 | 0.209336291 | 0.986203542 |
| 255 | *TTC9B* | 3.360842105 | 0.028226889 | 0.604484055 | 0.779063174 |
| 110 | *IPO5* | 3.328421053 | 0.027946291 | 0.614052381 | 0.435186727 |
| 227 | *SRRT* | 3.371649123 | 0.027539971 | 0.771525397 | 0.86491113 |
| 36 | *CACNB1* | 3.312280702 | 0.027102081 | 1.613809951 | 0.92658335 |
| 142 | *MMGT1* | 3.480982456 | 0.025777778 | 0.109885714 | 0.965337114 |
| 45 | *CEP19* | 2.947789474 | 0.025548035 | 3.632731548 | 0.034882359 |
| 168 | *PCDHAC2* | 3.297403509 | 0.025544136 | 1.24954067 | 0.672875424 |
| 197 | *RAMP1* | 3.354105263 | 0.025244444 | 0.515764286 | 0.86491113 |
| 71 | *EIF1B* | 3.485052632 | 0.024428403 | 0.0847 | 0.995563154 |
| 131 | *MAP6* | 3.439157895 | 0.023302222 | 0.474744078 | 0.986203542 |
| 213 | *SEC24D* | 3.389192982 | 0.022716481 | 0.258128571 | 0.86491113 |
| 235 | *SYDE2* | 3.456701754 | 0.022496736 | 0.532671429 | 0.986203542 |
| 164 | *OAZ2* | 3.212350877 | 0.02227037 | 1.322546753 | 0.324422251 |
| 43 | *CDH4* | 3.345964912 | 0.021587523 | 0.386961688 | 0.435186727 |
| 61 | *DEPDC5* | 3.184 | 0.021363333 | 1.407317274 | 0.22969607 |
| 16 | *APBA1* | 3.393263158 | 0.021258117 | 0.479052381 | 0.965337114 |
| 154 | *NEURL3* | 3.4 | 0.021021728 | 0.470620635 | 0.92658335 |
| 38 | *CAPRIN2* | 3.24477193 | 0.020866241 | 1.245826557 | 0.672875424 |
| 191 | *PTPRM* | 3.209684211 | 0.020675001 | 1.843466933 | 0.672875424 |
| 205 | *RNFT2* | 3.485052632 | 0.019970417 | 0.272735714 | 0.995563154 |
| 249 | *TRADD* | 3.410807018 | 0.019828008 | 2.792110159 | 0.965337114 |

| | | | | | |
|---|---|---|---|---|---|
| 196 | *RAB12* | 3.520140351 | 0.0194475 | 0.107909524 | 0.995563154 |
| 153 | *NEK9* | 3.325754386 | 0.019367268 | 1.083062915 | 0.779063174 |
| 39 | *CCND2* | 3.332491228 | 0.017795101 | 0.982493258 | 0.672875424 |
| 181 | *POLR1A* | 3.474245614 | 0.01779 | 0.131871429 | 0.986203542 |
| 146 | *MRPS33* | 3.480982456 | 0.01770303 | 0.184028571 | 0.965337114 |
| 3 | *ACTR3* | 3.474245614 | 0.017418232 | 0.186780952 | 0.986203542 |
| 225 | *SPTAN1* | 3.360842105 | 0.016853304 | 0.37918254 | 0.779063174 |
| 256 | *TTPAL* | 3.49445614 | 0.016311111 | 0.177977922 | 0.86491113 |
| 230 | *STARD9* | 3.548491228 | 0.016282 | 0.038257143 | 0.998920641 |
| 160 | *NPHP4* | 3.410807018 | 0.016156813 | 0.353146125 | 0.965337114 |
| 132 | *MARK4* | 3.364912281 | 0.01523625 | 0.959677417 | 0.92658335 |
| 170 | *PCK2* | 3.41754386 | 0.013240741 | 0.422478838 | 0.86491113 |
| 34 | *C9orf16* | 3.4 | 0.012459677 | 0.681380478 | 0.92658335 |
| 137 | *METRN* | 3.526877193 | 0.012424242 | 0.097571429 | 0.986203542 |
| 253 | *TRUB1* | 3.526877193 | 0.012214074 | 0.103554762 | 0.986203542 |
| 64 | *DNAJC30* | 3.188070175 | 0.01220004 | 3.889251587 | 0.435186727 |
| 79 | *FAM160B1* | 3.41754386 | 0.01175463 | 0.16367619 | 0.92658335 |
| 156 | *NFS1* | 3.198877193 | 0.011714866 | 3.023503292 | 0.554702513 |
| 125 | *LRRN2* | 3.485052632 | 0.01156 | 0.442313908 | 0.995563154 |
| 198 | *RASSF7* | 3.537684211 | 0.009965833 | 0.097852381 | 0.995563154 |
| 35 | *CABYR* | 3.325754386 | 0.009670525 | 0.92180873 | 0.779063174 |
| 189 | *PSME3* | 3.487719298 | 0.00951358 | 0.136540476 | 0.92658335 |
| 122 | *LRP1* | 3.38245614 | 0.009183056 | 0.369802381 | 0.92658335 |
| 236 | *SYT3* | 3.520140351 | 0.009025778 | 0.0814 | 0.995563154 |
| 150 | *NDFIP1* | 3.474245614 | 0.00825 | 0.489282051 | 0.965337114 |

| | | | | | |
|---|---|---|---|---|---|
| 55 | *CYFIP2* | 3.513403509 | 0.008162222 | 0.225204396 | 0.998920641 |
| 13 | *ANKIB1* | 3.530947368 | 0.007111111 | 0.1608 | 0.998920641 |
| 129 | *MAP2* | 3.513403509 | 0.007095486 | 0.196297619 | 0.998920641 |
| 95 | *GNPDA2* | 3.513403509 | 0.006483025 | 0.353945887 | 0.998920641 |
| 148 | *MYH3* | 3.059789474 | 0.006338677 | 0.930028427 | 0.019311298 |
| 116 | *KIF19* | 3.432421053 | 0.005828571 | 1.723344176 | 0.995563154 |
| 56 | *DARS* | 3.480982456 | 0.005065972 | 0.271740404 | 0.965337114 |
| 269 | *ZKSCAN1* | 3.410807018 | 0.004604848 | 0.697522206 | 0.965337114 |
| 94 | *GNG5* | 3.410807018 | 0.004079403 | 0.803627872 | 0.965337114 |
| 46 | *CETN3* | 3.297403509 | 0.003501192 | 0.684974242 | 0.672875424 |
| 49 | *CNOT1* | 2.930245614 | 0.003297969 | 4.449915158 | 0.019311298 |
| 89 | *FRMD8* | 3.378385965 | 0.003292929 | 0.404868398 | 0.779063174 |
| 257 | *UBE2B* | 3.240701754 | 0.003150056 | 0.851284726 | 0.435186727 |
| 84 | *FHOD1* | 3.474245614 | 0.002804222 | 0.171287302 | 0.986203542 |
| 185 | *PQLC1* | 3.485052632 | 0.0027935 | 0.262468687 | 0.995563154 |
| 183 | *PPDPF* | 3.587649123 | 0.002765432 | 0.021333333 | 0.999985499 |
| 6 | *ADCY1* | 3.435087719 | 0.001989553 | 0.36895873 | 0.92658335 |
| 263 | *WNK2* | 3.371649123 | 0.001786592 | 0.594650366 | 0.86491113 |
| 145 | *MRPL49* | 3.491789474 | 0.001459753 | 0.142862698 | 0.986203542 |
| 161 | *NPPB* | 3.463438596 | 0.000687556 | 0.137907143 | 0.965337114 |
| 107 | *IDI1* | 3.452631579 | 0.000625 | 0.168647619 | 0.86491113 |
| 28 | *BCAN* | 3.197473684 | 0.000472643 | 0.94886221 | 0.098798453 |
| 242 | *THOP1* | 3.587649123 | 0.000411429 | 0.0216 | 0.999985499 |
| 201 | *RIIAD1* | 3.421614035 | 0.000308444 | 0.54273254 | 0.986203542 |
| 176 | *PIGX* | 3.502596491 | 0.000261333 | 0.266240171 | 0.995563154 |

| | | | | | |
|---|---|---|---|---|---|
| 144 | *MRPL34* | 3.541754386 | 0.000125 | 0.247669697 | 0.999823643 |
| 58 | *DCUN1D5* | 3.559298246 | 0 | 0.06475 | 0.999823643 |
| 182 | *POLR2E* | 3.587649123 | 0 | 0.0375 | 0.999985499 |
| 260 | *VAC14* | 3.587649123 | 0 | 0.004166667 | 0.999985499 |
| 266 | *YWHAZ* | 3.576842105 | 0 | 0.076083333 | 0.999823643 |
| 271 | *ZNF415* | 3.605192982 | 0 | 0.007714286 | 0.999985499 |
| 231 | *STAT2* | 3.456701754 | -3.13E-05 | 0.303686203 | 0.986203542 |
| 75 | *ETV5* | 3.148912281 | -0.000418485 | 1.394616545 | 0.154497213 |
| 5 | *ADAMTS10* | 3.485052632 | -0.001344444 | 0.213816667 | 0.995563154 |
| 33 | *C9orf116* | 3.02877193 | -0.001706552 | 1.654521429 | 0.0601344 |
| 40 | *CCNYL1* | 3.389192982 | -0.001825879 | 0.307830159 | 0.86491113 |
| 31 | *C16orf70* | 3.38245614 | -0.002033455 | 1.946910303 | 0.92658335 |
| 200 | *RFXANK* | 3.474245614 | -0.002140354 | 0.469586501 | 0.986203542 |
| 172 | *PDK2* | 3.485052632 | -0.002929443 | 0.261547403 | 0.995563154 |
| 81 | *FARSB* | 3.594385965 | -0.002965909 | 0.0195 | 0.999823643 |
| 82 | *FBXL18* | 3.364912281 | -0.003244881 | 0.693208449 | 0.92658335 |
| 244 | *TMEM191A* | 3.08954386 | -0.003328107 | 1.61999011 | 0.324422251 |
| 264 | *YIF1B* | 3.520140351 | -0.003424444 | 0.083980952 | 0.995563154 |
| 194 | *PWWP2B* | 3.321684211 | -0.003529938 | 0.345773016 | 0.554702513 |
| 259 | *UBE2L3* | 3.124631579 | -0.003914459 | 1.865250011 | 0.324422251 |
| 20 | *ARL2* | 3.343298246 | -0.004145455 | 1.06505656 | 0.779063174 |
| 217 | *SLC1A4* | 3.096280702 | -0.004325257 | 1.802056133 | 0.22969607 |
| 195 | *RAB11FIP2* | 3.435087719 | -0.004517519 | 0.133563492 | 0.92658335 |
| 60 | *DEF8* | 3.4 | -0.004723333 | 0.947624459 | 0.92658335 |
| 173 | *PEX11A* | 3.520140351 | -0.005949009 | 0.154061905 | 0.995563154 |

| | | | | | |
|---|---|---|---|---|---|
| 52 | *CPSF4* | 3.530947368 | -0.006735556 | 0.329833333 | 0.998920641 |
| 115 | *KCNQ2* | 3.439157895 | -0.006794048 | 0.595814453 | 0.986203542 |
| 252 | *TROVE2* | 3.109754386 | -0.006915673 | 1.048476237 | 0.098798453 |
| 151 | *NDUFA9* | 3.410807018 | -0.006944444 | 0.575909158 | 0.965337114 |
| 184 | *PPP5C* | 3.389192982 | -0.007156358 | 2.011191667 | 0.86491113 |
| 211 | *SAMD15* | 3.509333333 | -0.007305397 | 0.0738 | 0.986203542 |
| 128 | *MANEAL* | 3.304140351 | -0.007436529 | 0.75765944 | 0.554702513 |
| 178 | *PLBD2* | 3.537684211 | -0.007646605 | 0.08502381 | 0.995563154 |
| 83 | *FCGBP* | 3.502596491 | -0.007679012 | 0.266547619 | 0.995563154 |
| 54 | *CSTB* | 3.41754386 | -0.008165041 | 0.380328355 | 0.92658335 |
| 98 | *GRIN2B* | 3.474245614 | -0.008439955 | 0.17232265 | 0.986203542 |
| 85 | *FLOT2* | 3.474245614 | -0.00872484 | 0.148629365 | 0.986203542 |
| 32 | *C7* | 3.371649123 | -0.008870667 | 0.395697619 | 0.86491113 |
| 204 | *RNF208* | 3.310877193 | -0.009465432 | 0.499750866 | 0.435186727 |
| 30 | *C11orf1* | 3.509333333 | -0.010758173 | 0.115064286 | 0.986203542 |
| 221 | *SORBS3* | 3.456701754 | -0.010818673 | 0.300406349 | 0.986203542 |
| 163 | *NUTF2* | 3.576842105 | -0.011832 | 0.038385714 | 0.999823643 |
| 93 | *GLT8D1* | 3.485052632 | -0.012819764 | 0.394216667 | 0.995563154 |
| 65 | *DSTYK* | 3.41754386 | -0.0131 | 0.369426618 | 0.92658335 |
| 127 | *MALL* | 3.269052632 | -0.014132153 | 1.369182493 | 0.554702513 |
| 152 | *NDUFS4* | 3.495859649 | -0.01586553 | 0.559370996 | 0.998920641 |
| 130 | *MAP4K2* | 3.485052632 | -0.018079444 | 1.886391795 | 0.995563154 |
| 232 | *STX4* | 3.502596491 | -0.019358025 | 0.13152381 | 0.995563154 |
| 37 | *CAND2* | 3.456701754 | -0.021819259 | 0.3197663 | 0.986203542 |
| 203 | *RNF123* | 3.513403509 | -0.022152778 | 0.255817893 | 0.998920641 |

| | | | | | |
|---|---|---|---|---|---|
| 193 | *PTPRU* | 3.428350877 | -0.023235043 | 0.847206746 | 0.965337114 |
| 141 | *MLYCD* | 3.445894737 | -0.023514881 | 0.614738095 | 0.965337114 |
| 202 | *RILPL1* | 3.279859649 | -0.024877769 | 0.792015651 | 0.672875424 |
| 226 | *SRPK2* | 3.463438596 | -0.026456667 | 0.29135098 | 0.965337114 |
| 166 | *PARD3B* | 3.485052632 | -0.026760605 | 0.617507143 | 0.995563154 |
| 104 | *HEMK1* | 3.530947368 | -0.02718084 | 0.169502381 | 0.998920641 |
| 4 | *ACTR6* | 3.441824561 | -0.028340965 | 0.178052381 | 0.86491113 |
| 15 | *AP2S1* | 3.502596491 | -0.028462747 | 0.190222222 | 0.995563154 |
| 113 | *JOSD1* | 3.485052632 | -0.028605556 | 0.485190476 | 0.995563154 |
| 99 | *GRK6* | 3.445894737 | -0.029486515 | 0.484812512 | 0.965337114 |
| 250 | *TRIM4* | 3.389192982 | -0.031635732 | 0.301013492 | 0.86491113 |
| 87 | *FNBP4* | 3.350035088 | -0.031983626 | 0.432415873 | 0.672875424 |
| 41 | *CCT6B* | 3.148912281 | -0.032687425 | 0.886479004 | 0.22969607 |
| 11 | *AMIGO1* | 3.467508772 | -0.035222778 | 0.393174237 | 0.995563154 |
| 118 | *KRTCAP2* | 3.445894737 | -0.036060185 | 0.288689316 | 0.965337114 |
| 188 | *PSMD7* | 3.367578947 | -0.03691054 | 0.987534921 | 0.672875424 |
| 7 | *ADCY5* | 3.078736842 | -0.037011328 | 1.950659468 | 0.154497213 |
| 208 | *RPL7L1* | 3.463438596 | -0.037291667 | 0.396076496 | 0.965337114 |
| 234 | *STYXL1* | 3.424280702 | -0.037377989 | 0.16347619 | 0.86491113 |
| 241 | *TCF7* | 3.459368421 | -0.045545051 | 0.237540482 | 0.86491113 |
| 88 | *FOXP4* | 3.329824561 | -0.0484985 | 1.331177381 | 0.92658335 |
| 73 | *EME1* | 3.428350877 | -0.052483301 | 0.324921212 | 0.965337114 |
| 149 | *NAE1* | 3.146245614 | -0.065938941 | 9.750266244 | 0.554702513 |
| 214 | *SERTAD3* | 2.941052632 | -0.072323382 | 2.953244455 | 0.0601344 |
| 267 | *ZEB2* | 3.116491228 | -0.084890814 | 0.952029426 | 0.0601344 |

| | | | | | |
|---|---|---|---|---|---|
| 42 | *CD34* | 3.314947368 | -0.089495779 | 2.93329381 | 0.672875424 |
| 67 | *DYNLRB1* | 3.233964912 | -0.092625126 | 1.22104475 | 0.554702513 |
| 240 | *TBC1D2B* | 3.393263158 | -0.093603818 | 0.909644444 | 0.965337114 |
| 270 | *ZNF233* | 3.255578947 | -0.097082655 | 1.526100799 | 0.779063174 |
| 25 | *ATP6V1G1* | 3.233964912 | -0.104411144 | 1.556727587 | 0.554702513 |
| 76 | *EVL* | 3.354105263 | -0.131030222 | 3.3616929 | 0.86491113 |

**Table S33. Importance (Dominance) of each gene in the random forest model for rostral middle frontal cortex (rosMFC).** Mean_min_depth represents the average minimum depth at which a variable is used as a split node across all trees, with lower values indicating higher importance as the variable is used earlier in the decision process. IncrMSE, % (Increase in Mean Squared Error) quantifies the relative increase in prediction error when the variable is permuted, reflecting its contribution to the model's predictive accuracy; higher values signify greater importance. Node_purity_increase measures the improvement in node purity (e.g., Gini index or variance reduction) achieved by splitting on the variable, where larger values indicate a stronger impact on classification or regression performance. Finally, the P value assesses the statistical significance of a variable's importance, typically derived from permutation tests, with lower values suggesting that the variable's contribution is unlikely to be due to random chance. This presentation was in descending order by the IncrMSE.

**10. Biological Ontology Annotation to Cluster Longitudinal Gene Expression Trajectory**

We utilized the R package "Mfuzz" to perform fuzzy clustering on the gene × neurodevelopmental matrix (273 genes × 30 periods), with gene expression data averaged across six brain regions to minimize regional variability. Prior to clustering, gene expression values were normalized for consistency across developmental stages. The Mfuzz package was used to conduct fuzzy c-means clustering, specifying two clusters as the optimal cluster number (*see Supplemental Methods 4*). By doing so, 93 (180) genes were identified into the cluster 1 (2) **(Table S34-35)**.

| Gene Symbol | Cluster Number # |
|---|---|
| *CREB3L3* | Cluster#1 |
| *SP100* | Cluster#1 |
| *TMEM38A* | Cluster#1 |
| *TCF7* | Cluster#1 |
| *AQP6* | Cluster#1 |
| *NOS1* | Cluster#1 |

| | |
|---|---|
| *FCGBP* | Cluster#1 |
| *PITPNM2* | Cluster#1 |
| *ANGPT2* | Cluster#1 |
| *SYDE2* | Cluster#1 |
| *RASSF7* | Cluster#1 |
| *DEPDC5* | Cluster#1 |
| *SAMD15* | Cluster#1 |
| *PCK2* | Cluster#1 |
| *TRADD* | Cluster#1 |
| *MLYCD* | Cluster#1 |
| *METRN* | Cluster#1 |
| *PLEKHA1* | Cluster#1 |
| *HLF* | Cluster#1 |
| *MYH3* | Cluster#1 |
| *CAPRIN2* | Cluster#1 |
| *C7* | Cluster#1 |
| *HEMK1* | Cluster#1 |
| *PARD3B* | Cluster#1 |
| *NEK9* | Cluster#1 |
| *SORBS3* | Cluster#1 |
| *NPPB* | Cluster#1 |
| *TTPAL* | Cluster#1 |
| *FRMD8* | Cluster#1 |
| *MCHR1* | Cluster#1 |
| *KRT17* | Cluster#1 |
| *NPHP4* | Cluster#1 |
| *CCT6B* | Cluster#1 |
| *RAMP1* | Cluster#1 |
| *LPIN1* | Cluster#1 |
| *DTNA* | Cluster#1 |
| *FHOD1* | Cluster#1 |
| *GPNMB* | Cluster#1 |
| *C11orf1* | Cluster#1 |
| *SSFA2* | Cluster#1 |
| *MALL* | Cluster#1 |
| *CAND2* | Cluster#1 |
| *SLC10A4* | Cluster#1 |
| *TRIM4* | Cluster#1 |
| *GRIN2B* | Cluster#1 |
| *SEC24D* | Cluster#1 |
| *PIGF* | Cluster#1 |
| *CETN3* | Cluster#1 |
| *ZNF599* | Cluster#1 |
| *EME1* | Cluster#1 |

| | |
|---|---|
| *FBXL18* | Cluster#1 |
| *STARD9* | Cluster#1 |
| *ZNF233* | Cluster#1 |
| *C9orf116* | Cluster#1 |
| *S100A1* | Cluster#1 |
| *LEMD2* | Cluster#1 |
| *NEURL3* | Cluster#1 |
| *CCNYL1* | Cluster#1 |
| *FZD5* | Cluster#1 |
| *GNPDA2* | Cluster#1 |
| *KRTCAP2* | Cluster#1 |
| *ANKS6* | Cluster#1 |
| *ATP9B* | Cluster#1 |
| *PEX11A* | Cluster#1 |
| *TBC1D2B* | Cluster#1 |
| *SERTAD3* | Cluster#1 |
| *MAP4K2* | Cluster#1 |
| *DYNLRB2* | Cluster#1 |
| *BOP1* | Cluster#1 |
| *PWWP2B* | Cluster#1 |
| *SNCG* | Cluster#1 |
| *CEP19* | Cluster#1 |
| *CD34* | Cluster#1 |
| *EPS8L2* | Cluster#1 |
| *RIIAD1* | Cluster#1 |
| *MKRN3* | Cluster#1 |
| *LYNX1* | Cluster#1 |
| *AATK* | Cluster#1 |
| *RESP18* | Cluster#1 |
| *TNFAIP8L3* | Cluster#1 |
| *FMNL1* | Cluster#1 |
| *DDX51* | Cluster#1 |
| *FAM174B* | Cluster#1 |
| *CIDECP* | Cluster#1 |
| *RILPL1* | Cluster#1 |
| *SLC38A3* | Cluster#1 |
| *KIF19* | Cluster#1 |
| *CFD* | Cluster#1 |
| *NOS1AP* | Cluster#1 |
| *MT1A* | Cluster#1 |
| *HCP5* | Cluster#1 |
| *TMEM191A* | Cluster#1 |
| *PRB1* | Cluster#1 |
| *ANKIB1* | Cluster#2 |

| | |
|---|---|
| *PDK2* | Cluster#2 |
| *MARK4* | Cluster#2 |
| *VTA1* | Cluster#2 |
| *IDS* | Cluster#2 |
| *PPP5C* | Cluster#2 |
| *EHD3* | Cluster#2 |
| *GLT8D1* | Cluster#2 |
| *TMSB10* | Cluster#2 |
| *TRIO* | Cluster#2 |
| *AP2S1* | Cluster#2 |
| *NNAT* | Cluster#2 |
| *CYFIP2* | Cluster#2 |
| *PCGF2* | Cluster#2 |
| *PTPRU* | Cluster#2 |
| *RFXANK* | Cluster#2 |
| *IPO5* | Cluster#2 |
| *TBC1D1* | Cluster#2 |
| *IDI1* | Cluster#2 |
| *CACNB1* | Cluster#2 |
| *POLR1A* | Cluster#2 |
| *KCNQ2* | Cluster#2 |
| *ACTR6* | Cluster#2 |
| *MAP2* | Cluster#2 |
| *PAFAH1B3* | Cluster#2 |
| *SRRT* | Cluster#2 |
| *GRAMD1A* | Cluster#2 |
| *MRPS33* | Cluster#2 |
| *DLL3* | Cluster#2 |
| *POLR2E* | Cluster#2 |
| *JOSD1* | Cluster#2 |
| *EEF1A2* | Cluster#2 |
| *NUTF2* | Cluster#2 |
| *ZNF821* | Cluster#2 |
| *PSMD7* | Cluster#2 |
| *VAC14* | Cluster#2 |
| *STX4* | Cluster#2 |
| *PLD3* | Cluster#2 |
| *PTPRS* | Cluster#2 |
| *ZKSCAN1* | Cluster#2 |
| *MEST* | Cluster#2 |
| *APBA1* | Cluster#2 |
| *RAB11FIP2* | Cluster#2 |
| *MICU1* | Cluster#2 |
| *FNBP4* | Cluster#2 |
| *RPL24* | Cluster#2 |
| *EIF1B* | Cluster#2 |
| *ACTR3* | Cluster#2 |
| *DARS* | Cluster#2 |
| *SLC1A4* | Cluster#2 |
| *SOS1* | Cluster#2 |

| | |
|---|---|
| *FARSB* | Cluster#2 |
| *TROVE2* | Cluster#2 |
| *CCND2* | Cluster#2 |
| *UBE2B* | Cluster#2 |
| *AFTPH* | Cluster#2 |
| *FABP3* | Cluster#2 |
| *PQLC1* | Cluster#2 |
| *LRP1* | Cluster#2 |
| *CNOT1* | Cluster#2 |
| *C16orf70* | Cluster#2 |
| *PPDPF* | Cluster#2 |
| *GTF2F1* | Cluster#2 |
| *NRSN2* | Cluster#2 |
| *MAX* | Cluster#2 |
| *DYNLRB1* | Cluster#2 |
| *LRRC61* | Cluster#2 |
| *STYXL1* | Cluster#2 |
| *MRPL34* | Cluster#2 |
| *JUND* | Cluster#2 |
| *LRP3* | Cluster#2 |
| *PSME3* | Cluster#2 |
| *NDFIP1* | Cluster#2 |
| *ACACA* | Cluster#2 |
| *DLG4* | Cluster#2 |
| *FLOT2* | Cluster#2 |
| *BCAN* | Cluster#2 |
| *DSTYK* | Cluster#2 |
| *SPIRE1* | Cluster#2 |
| *ARF3* | Cluster#2 |
| *SLC38A2* | Cluster#2 |
| *RNFT2* | Cluster#2 |
| *SRPK2* | Cluster#2 |
| *SCRN1* | Cluster#2 |
| *LIMD2* | Cluster#2 |
| *STXBP1* | Cluster#2 |
| *ATP6V1G1* | Cluster#2 |
| *FOXP4* | Cluster#2 |
| *DCUN1D5* | Cluster#2 |
| *RTF1* | Cluster#2 |
| *NDUFA9* | Cluster#2 |
| *HMG20A* | Cluster#2 |
| *DEF8* | Cluster#2 |
| *ADAMTS10* | Cluster#2 |
| *RPL13A* | Cluster#2 |
| *ILF2* | Cluster#2 |
| *G3BP1* | Cluster#2 |
| *GFOD1* | Cluster#2 |
| *RPL7L1* | Cluster#2 |
| *ASTN2* | Cluster#2 |
| *MRPL49* | Cluster#2 |

| | |
|---|---|
| *PLBD2* | Cluster#2 |
| *FAM160B1* | Cluster#2 |
| *JAZF1* | Cluster#2 |
| *CABYR* | Cluster#2 |
| *THY1* | Cluster#2 |
| *AGPAT5* | Cluster#2 |
| *PTDSS1* | Cluster#2 |
| *AHCYL2* | Cluster#2 |
| *COPG2* | Cluster#2 |
| *NAE1* | Cluster#2 |
| *PDE9A* | Cluster#2 |
| *CSTB* | Cluster#2 |
| *ZER1* | Cluster#2 |
| *CPSF4* | Cluster#2 |
| *ARPC2* | Cluster#2 |
| *PIGX* | Cluster#2 |
| *RNF123* | Cluster#2 |
| *WDR82* | Cluster#2 |
| *NDUFS4* | Cluster#2 |
| *ADCY1* | Cluster#2 |
| *MED30* | Cluster#2 |
| *YWHAZ* | Cluster#2 |
| *WNK2* | Cluster#2 |
| *TRUB1* | Cluster#2 |
| *PCBD1* | Cluster#2 |
| *DCHS1* | Cluster#2 |
| *COG1* | Cluster#2 |
| *YWHAB* | Cluster#2 |
| *YIF1B* | Cluster#2 |
| *ATF5* | Cluster#2 |
| *MMGT1* | Cluster#2 |
| *ZEB2* | Cluster#2 |
| *LRRN2* | Cluster#2 |
| *ELOVL6* | Cluster#2 |
| *STAT2* | Cluster#2 |
| *IRF2BP1* | Cluster#2 |
| *ZNF415* | Cluster#2 |
| *C9orf16* | Cluster#2 |
| *CDK5R2* | Cluster#2 |
| *NEUROD2* | Cluster#2 |
| *MAP6* | Cluster#2 |
| *THOP1* | Cluster#2 |
| *TADA2B* | Cluster#2 |
| *ADCY5* | Cluster#2 |
| *SSSCA1* | Cluster#2 |
| *PTPRM* | Cluster#2 |
| *GNG5* | Cluster#2 |
| *TTC9B* | Cluster#2 |
| *DNAJC30* | Cluster#2 |
| *BASP1* | Cluster#2 |

| | |
|---|---|
| *ARIH2* | Cluster#2 |
| *CDH4* | Cluster#2 |
| *FAHD1* | Cluster#2 |
| *OAZ2* | Cluster#2 |
| *AMIGO1* | Cluster#2 |
| *UBE2E2* | Cluster#2 |
| *SF3A3* | Cluster#2 |
| *MANEAL* | Cluster#2 |
| *TARSL2* | Cluster#2 |
| *UBE2L3* | Cluster#2 |
| *HBA2* | Cluster#2 |
| *EVL* | Cluster#2 |
| *SPTAN1* | Cluster#2 |
| *PHF2* | Cluster#2 |
| *MPZL1* | Cluster#2 |
| *GRK6* | Cluster#2 |
| *ARMCX6* | Cluster#2 |
| *PSMB10* | Cluster#2 |
| *HBA1* | Cluster#2 |
| *RAB12* | Cluster#2 |
| *RNF208* | Cluster#2 |
| *SYT3* | Cluster#2 |
| *MLLT11* | Cluster#2 |
| *ARL2* | Cluster#2 |
| *TTC3P1* | Cluster#2 |
| *PCDHAC2* | Cluster#2 |
| *NFS1* | Cluster#2 |
| *TMEM199* | Cluster#2 |
| *ETV5* | Cluster#2 |

To further investigate the biological significance of the two gene clusters identified through fuzzy clustering, we used the R package "clusterProfiler" to perform Gene Ontology (GO) enrichment analysis. The two clusters of genes, identified from the Mfuzz clustering, were separately annotated with GO terms using the clusterProfiler package. GO enrichment analysis was performed for each cluster to determine overrepresented biological processes, molecular functions, and cellular components based on the org.Hs.eg.db annotation database for human genes. The enrichGO() function in clusterProfiler was used to identify significantly enriched GO terms, applying a false discovery rate (FDR) threshold of $< 0.05$ for statistical significance. Results converged into a line that these clusters enriched into biological functions of molecular transports **(Table S34-35)**.

| GO terms | Frequency |
|---|---|
| NO cGMP PKG mediated neuroprotection | 3 |
| Nuclear Envelope Breakdown | 3 |
| Pleural mesothelioma | 7 |
| microtubule-based movement | 7 |
| regulation of angiogenesis | 6 |
| Alcoholic liver disease | 4 |

| | |
|---|---|
| cellular catabolic process | 5 |
| Arrhythmogenic right ventricular cardiomyopathy | 3 |
| Ciliopathies | 4 |
| Phospholipid metabolism | 4 |
| Motor proteins | 4 |
| negative regulation of growth | 4 |
| TNF signaling pathway | 3 |
| NO cGMP PKG mediated neuroprotection | 3 |
| Nuclear Envelope Breakdown | 3 |
| Pleural mesothelioma | 7 |
| microtubule-based movement | 7 |
| regulation of angiogenesis | 6 |
| Alcoholic liver disease | 4 |
| cellular catabolic process | 5 |

**Table S34 GO terms of enriching into the Cluster #1 genes.**

| GO terms | Frequency |
|---|---|
| cell morphogenesis involved in neuron differentiation | 14 |
| Transport of small molecules | 18 |
| Vesicle-mediated transport | 17 |
| PID PDGFRB PATHWAY | 8 |
| Axon guidance | 15 |
| Neuronal System | 12 |
| organophosphate biosynthetic process | 13 |
| Programmed Cell Death | 8 |
| Scavenging of heme from plasma | 3 |
| Adaptive Immune System | 15 |
| Signaling by Receptor Tyrosine Kinases | 12 |
| amide biosynthetic process | 12 |
| regulation of neurogenesis | 10 |
| Protein-protein interactions at synapses | 5 |
| PID LKB1 PATHWAY | 4 |
| regulation of protein-containing complex assembly | 10 |
| regulation of cell projection organization | 13 |
| Golgi vesicle transport | 8 |
| Influenza Infection | 6 |
| actin nucleation | 3 |

**Table S35 GO terms of enriching into the Cluster #2 genes.**

**11. Neurobiological Enrichment to PPS-specific Gene Set**

To reveal the biological processes and enriched pathways associated with the PPS-specific gene set (i.e., PLS2), we conducted enrichment analysis using the Metascape (http://metascape.org/gp/index.html). The latest updated annotation datasets were integrated, including functional categories from Metascape's curated resources (last updated: 01-05-2024). All genes in the genome were used as the enrichment background to ensure comprehensive analysis, and Benjamini-Hochberg FDR correction was applied to adjust p-values for multiple comparisons. The full results of enrichment analysis for PLS2 have been structured into **Table S36**.

| GO | Category | Description | Count | % | Log10(P) |
|---|---|---|---|---|---|
| R-HSA-5653656 | Reactome Gene Sets | Vesicle-mediated transport | 21 | 6.91 | -5.27 |
| R-HSA-382551 | Reactome Gene Sets | Transport of small molecules | 22 | 7.24 | -5.19 |
| GO:0090407 | GO Biological Processes | organophosphate biosynthetic process | 18 | 5.92 | -4.94 |
| WP289 | WikiPathways | Myometrial relaxation and contraction pathways | 9 | 2.96 | -4.52 |
| R-HSA-983169 | Reactome Gene Sets | Class I MHC mediated antigen processing & presentation | 14 | 4.61 | -4.45 |
| GO:1990778 | GO Biological Processes | protein localization to cell periphery | 11 | 3.62 | -4.37 |
| GO:0048667 | GO Biological Processes | cell morphogenesis involved in neuron differentiation | 15 | 4.93 | -4.35 |
| GO:0007018 | GO Biological Processes | microtubule-based movement | 14 | 4.61 | -4.31 |
| M186 | Canonical Pathways | PID PDGFRB PATHWAY | 8 | 2.63 | -4.30 |
| R-HSA-8849932 | Reactome Gene Sets | Synaptic adhesion-like molecules | 4 | 1.32 | -4.28 |
| GO:0032784 | GO Biological Processes | regulation of DNA-templated transcription elongation | 7 | 2.30 | -4.13 |
| GO:0006868 | GO Biological Processes | glutamine transport | 3 | 0.99 | -4.09 |
| GO:0051660 | GO Biological Processes | establishment of centrosome localization | 3 | 0.99 | -3.94 |

| | | | | | |
|---|---|---|---|---|---|
| GO:0048193 | GO Biological Processes | Golgi vesicle transport | 11 | 3.62 | -3.86 |
| GO:0009152 | GO Biological Processes | purine ribonucleotide biosynthetic process | 9 | 2.96 | -3.86 |
| R-HSA-422475 | Reactome Gene Sets | Axon guidance | 16 | 5.26 | -3.82 |
| R-HSA-2168880 | Reactome Gene Sets | Scavenging of heme from plasma | 3 | 0.99 | -3.58 |
| GO:0031346 | GO Biological Processes | positive regulation of cell projection organization | 12 | 3.95 | -3.50 |

**Table S36. Top 20 clusters with their representative enriched terms (one per cluster) for PLS1 component.** "Count" is the number of genes in the user-provided lists with membership in the given ontology term. "%" is the percentage of all of the user-provided genes that are found in the given ontology term (only input genes with at least one ontology term annotation are included in the calculation). P values are estimated by two-sided cumulative hypergeometric distribution test, with Benjamini-Hochberg FDR correction. "Log10(P)" is the p-value in log base 10. "Log10(q)" is the multi-test adjusted p-value in log base 10.

For deeper insights into the functional relevance of PPS-specific PLS2 genes, we conducted protein-protein interaction (PPI) enrichment analysis using Metascape, leveraging interaction datasets from STRING, BioGrid, OmniPath, and InWeb_IM. Only physical interactions from STRING (physical score > 0.132) and BioGrid were retained to ensure high-confidence network construction. The resultant PPI network contained the subset of proteins forming physical interactions with at least one other gene from the list. Since the network included between 3 and 500 proteins, we applied the Molecular Complex Detection (MCODE) algorithm to identify densely connected functional modules. The PPI results for PLS2 have been summarized in **Table S37**.

| **MCODE** | **GO** | **Description** | **Log10(P)** |
|---|---|---|---|
| MCODE_1 | R-HSA-6791226 | Major pathway of rRNA processing in the nucleolus and cytosol | -5.7 |
| MCODE_1 | R-HSA-8868773 | rRNA processing in the nucleus and cytosol | -5.6 |
| MCODE_1 | R-HSA-72312 | rRNA processing | -5.5 |

| | | | |
|---|---|---|---|
| MCODE_2 | R-HSA-112382 | Formation of RNA Pol II elongation complex | -10.2 |
| MCODE_2 | R-HSA-75955 | RNA Polymerase II Transcription Elongation | -10.2 |
| MCODE_2 | R-HSA-674695 | RNA Polymerase II Pre-transcription Events | -9.6 |
| MCODE_3 | R-HSA-350562 | Regulation of ornithine decarboxylase (ODC) | -11.1 |
| MCODE_3 | R-HSA-351202 | Metabolism of polyamines | -10.9 |
| MCODE_3 | hsa03050 | Proteasome | -7.9 |
| MCODE_4 | CORUM:320 | 55S ribosome, mitochondrial | -7.2 |
| MCODE_4 | R-HSA-5419276 | Mitochondrial translation termination | -7.0 |
| MCODE_4 | R-HSA-5389840 | Mitochondrial translation elongation | -7.0 |
| MCODE_5 | GO:0052652 | cyclic purine nucleotide metabolic process | -8.7 |
| MCODE_5 | GO:0009187 | cyclic nucleotide metabolic process | -8.6 |
| MCODE_5 | WP4222 | Phosphodiesterases in neuronal function | -8.2 |

**Table S37. Protein-Protein Interaction (PPI) networks based on the Molecular Complex Detection (MCODE) algorithm.** *P* values are estimated by two-sided cumulative hypergeometric distribution test, with Benjamini-Hochberg FDR correction. "Log10(P)" is the p-value in log base 10. "Log10(q)" is the multi-test adjusted p-value in log base 10.

We have further analyzed neurobiological enrichment of PPS-specific gene set using both Metascape and Specific Expression Analysis (SEA). Statistical threshold was set to be $p < .05$ with Benjamini-Hochberg FDR corrections. All genes in the genome have been used as the enrichment background. Terms with a p-value < 0.01, a minimum count of 3, and an enrichment factor > 1.5 (the enrichment factor is the ratio between the observed counts and the counts expected by chance) are collected and grouped into clusters based on their membership similarities. Full results could be found in the **Table S38-39**.

| GO | Description | Count (%) | Log10(P) | Log10(q) |
|---|---|---|---|---|
| PGB:00065 | Cell-specific: DRG | 13 (4.3%) | -3.10 | -0.43 |

**Table S38. Tissue-specific enrichment in the PLS2 gene set.** *P* values are estimated by two-sided cumulative hypergeometric distribution test, with Benjamini-Hochberg FDR correction. "Log10(P)" is the p-value in log base 10.

| Go | Description | Count | Log10(P) | Log10(q) |
|---|---|---|---|---|
| M39269 | HU FETAL RETINA RGC | 19 | -6.8 | -2.3 |
| M39072 | MANNO MIDBRAIN NEUROTYPES HSERT | 17 | -5.4 | -1.5 |
| M39068 | MANNO MIDBRAIN NEUROTYPES HDA1 | 18 | -4.5 | -1.0 |
| M39104 | ZHONG PFC C3 MICROGLIA | 16 | -4.4 | -0.96 |
| M39069 | MANNO MIDBRAIN NEUROTYPES HDA2 | 15 | -3.6 | -0.60 |
| M41715 | FAN OVARY CL13 MONOCYTE MACROPHAGE | 14 | -3.6 | -0.60 |
| M39066 | MANNO MIDBRAIN NEUROTYPES HNBML5 | 14 | -3.5 | -0.58 |
| M41710 | FAN OVARY CL8 MATURE CUMULUS GRANULOSA CELL 2 | 17 | -3.5 | -0.58 |
| M41745 | RUBENSTEIN SKELETAL MUSCLE FAP CELLS | 8 | -3.3 | -0.58 |
| M40004 | BUSSLINGER ESOPHAGEAL LATE SUPRABASAL CELLS | 7 | -3.3 | -0.53 |

**Table S39. Cell-specific enrichment in the PLS2 gene set.** *P* values are estimated by two-sided cumulative hypergeometric distribution test, with Benjamini-Hochberg FDR correction. "Log10(P)" is the p-value in log base 10.

Finally, we decoded PPS-specific gene sets (i.e., PLS2+, PLS2-) at BrainMap to reveal the associations between gene expression patterns and risks of diseases/disorders in BrainMap dataset. BrainMap is the online meta-analytic model to estimate likelihood of linking both VBM and functional MRI datasets to gene-expression risks underlying diseases/disorders. Full results have been tabulated in the **Table S40-41 and Figure S6**.

| Modality | Item | Standardized beta |
|---|---|---|
| BrainMap_fMRI | ADHD | 0.003349164 |
| BrainMap_fMRI | ASD | 0.099988829 |
| BrainMap_fMRI | MCI | -0.126392876 |
| BrainMap_fMRI | MDD | -0.023048416 |
| BrainMap_fMRI | OCD | -0.068190847 |
| BrainMap_fMRI | PTSD | 0.008795849 |
| BrainMap_fMRI | Alzheimers | -0.068818454 |
| BrainMap_fMRI | Anxiety | -0.173441056 |
| BrainMap_fMRI | Asperger | 0.303313376 |
| BrainMap_fMRI | Bipolar | -0.18829482 |
| BrainMap_fMRI | Depression | -0.029424355 |
| BrainMap_fMRI | Dyslexia | 0.045251843 |
| BrainMap_fMRI | Obesity | 0.00081909 |
| BrainMap_fMRI | Parkinson | 0.040995484 |
| BrainMap_fMRI | Schizophrenia | -0.149307994 |
| BrainMap_fMRI | Stroke | -0.058457696 |
| BrainMap_VBM | ADHD | 0.074731819 |
| BrainMap_VBM | ALS | -0.336771983 |
| BrainMap_VBM | ASD | -0.124483571 |
| BrainMap_VBM | FTD | -0.229450457 |
| BrainMap_VBM | MCI | 0.083286756 |
| BrainMap_VBM | MS | 0.218288336 |
| BrainMap_VBM | OCD | -0.087479484 |
| BrainMap_VBM | PTSD | 0.082093065 |
| BrainMap_VBM | Alzheimers | 0.002625563 |
| BrainMap_VBM | Anxiety | -0.052654531 |
| BrainMap_VBM | Asperger | -0.082482915 |
| BrainMap_VBM | Bipolar | -0.162476462 |
| BrainMap_VBM | Dementia | -0.028252535 |
| BrainMap_VBM | Depression | 0.142062735 |
| BrainMap_VBM | Dyslexia | -0.019257034 |
| BrainMap_VBM | Huntington | 0.089740071 |

| | | |
|---|---|---|
| BrainMap_VBM | Obesity | 0.108192949 |
| BrainMap_VBM | Parkinson | -0.195433734 |
| BrainMap_VBM | Psychosis | -0.126658309 |
| BrainMap_VBM | Schizophrenia | -0.267994036 |
| BrainMap_VBM | SementicDementia | -5.50887E-05 |
| BrainMap_VBM | Stroke | 0.024866006 |

**Table S40. Association of gene set in the PLS2+ set for neurological and psychiatric diseases.** Statistical significance was set as two-sided $p < .05$ with Bonferroni-Holm FDR correction from $z$ test to linear regression model.

| Modality | Item | Standardized beta |
|---|---|---|
| BrainMap_fMRI | ADHD | -0.033460308 |
| BrainMap_fMRI | ASD | 0.098759776 |
| BrainMap_fMRI | MCI | 0.115434805 |
| BrainMap_fMRI | MDD | 0.269047707 |
| BrainMap_fMRI | OCD | 0.139783154 |
| BrainMap_fMRI | PTSD | 0.117294756 |
| BrainMap_fMRI | Alzheimers | 0.132357424 |
| BrainMap_fMRI | Anxiety | 0.09405709 |
| BrainMap_fMRI | Asperger | 0.02934885 |
| BrainMap_fMRI | Bipolar | 0.10523778 |
| BrainMap_fMRI | Depression | 0.143619621 |
| BrainMap_fMRI | Dyslexia | 0.109049514 |
| BrainMap_fMRI | Obesity | 0.165893627 |
| BrainMap_fMRI | Parkinson | -0.20614152 |
| BrainMap_fMRI | Schizophrenia | 0.088449435 |
| BrainMap_fMRI | Stroke | -0.002249885 |
| BrainMap_VBM | ADHD | 0.233211787 |
| BrainMap_VBM | ALS | -0.107876487 |
| BrainMap_VBM | ASD | -0.113231686 |
| BrainMap_VBM | FTD | 0.262264803 |
| BrainMap_VBM | MCI | 0.363106165 |
| BrainMap_VBM | MS | 0.013934664 |
| BrainMap_VBM | OCD | 0.029470959 |
| BrainMap_VBM | PTSD | 0.280620264 |
| BrainMap_VBM | Alzheimers | 0.540475387 |
| BrainMap_VBM | Anxiety | -0.087682949 |
| BrainMap_VBM | Asperger | -0.116367764 |
| BrainMap_VBM | Bipolar | 0.269895777 |
| BrainMap_VBM | Dementia | 0.628168933 |
| BrainMap_VBM | Depression | 0.005009292 |
| BrainMap_VBM | Dyslexia | 0.126141877 |
| BrainMap_VBM | Huntington | 0.127329089 |
| BrainMap_VBM | Obesity | 0.284518424 |

| | | |
|---|---|---|
| BrainMap_VBM | Parkinson | 0.362408151 |
| BrainMap_VBM | Psychosis | -0.267768215 |
| BrainMap_VBM | Schizophrenia | 0.214806527 |
| BrainMap_VBM | SementicDementia | 0.477517678 |
| BrainMap_VBM | Stroke | 0.022444893 |

**Table S41. Association of gene set in the PLS2- set for neurological and psychiatric diseases.** Statistical significance was set as two-sided $p < .05$ with Bonferroni-Holm FDR correction from $z$ test to linear regression model.

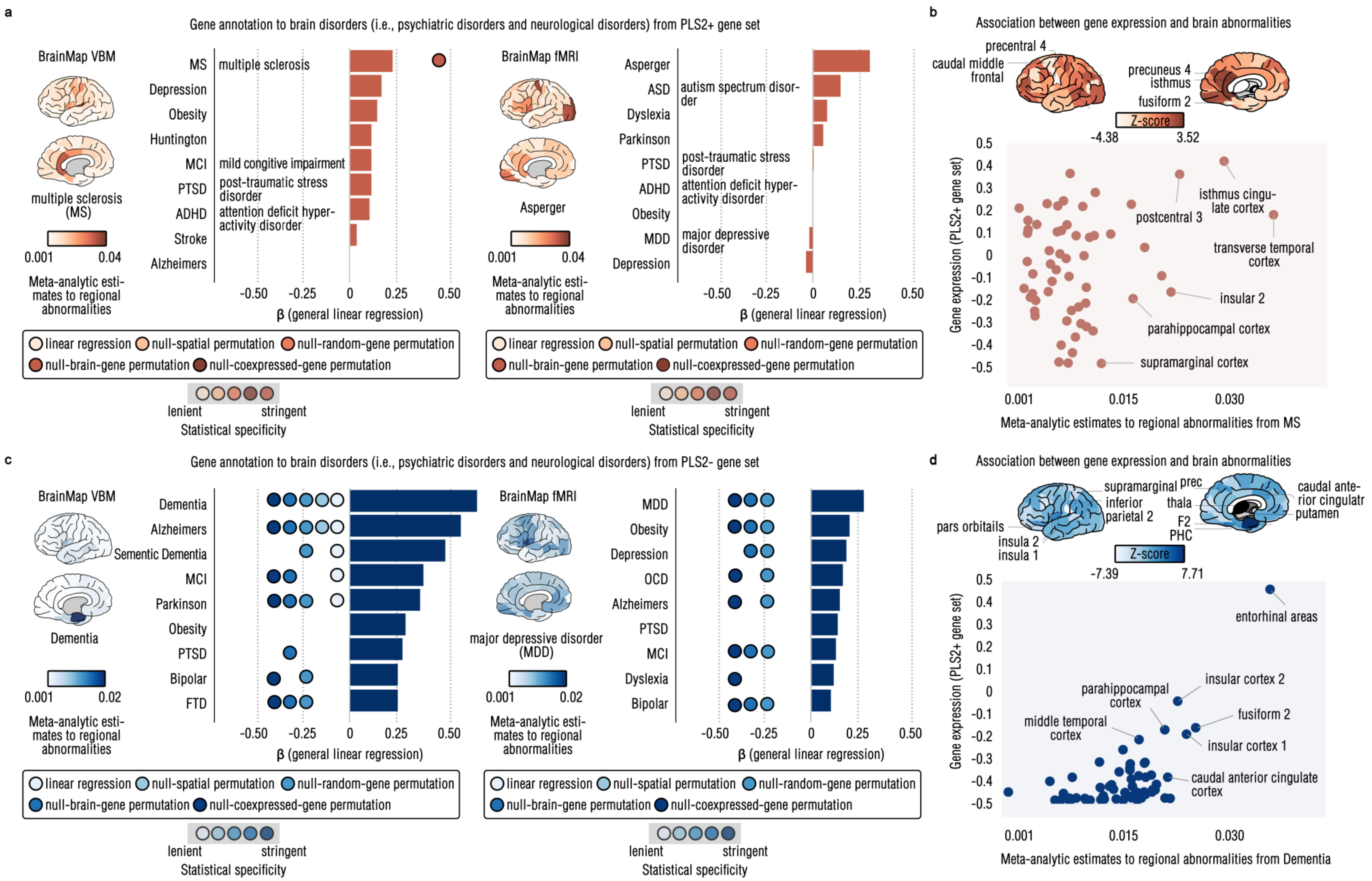


**Table S6. Brain disorders enrichment of PLS2 gene sets. a,** showed spatial similarity of regional structural (left) or functional abnormalities (right) that were estimated by meta-analysis in the BrainMap dataset for brain disorders and regional gene expression map for PLS2+ gene set. To reach high statistical specificity, the statistical significance was estimated by four nested spinned permutation tests. **b,** offered a scatter plot to show the linear association between brain structural abnormalities of multiple sclerosis (MS) and brain gene expression in the PLS2+. **c,** plot such spatial similarities from PLS2- gene set. **d,** drew a scatter plot to show the association between functional abnormalities of Dementia and gene expression of PLS2- set.
